# Computational inference, validation, and analysis of 5’UTR-leader sequences of alleles of immunoglobulin heavy chain variable genes

**DOI:** 10.1101/2021.06.10.447679

**Authors:** Yixun Huang, Linnea Thörnqvist, Mats Ohlin

**Author notes:** These authors have contributed equally to this work and share first authorship (Yixun Huang, Linnea Thörnqvist).

## Abstract

Upstream and downstream sequences of immunoglobulin genes may affect the expression of such genes. However, these sequences are rarely studied or characterized in most studies of immunoglobulin repertoires. Inference from large, rearranged immunoglobulin transcriptome data sets offers an opportunity to define the upstream regions (5’-untranslated regions and leader sequences). We have now established a new data pre-processing procedure to eliminate artifacts caused by a 5’-RACE library generation process, reanalyzed a previously studied data set defining human immunoglobulin heavy chain genes, and identified novel upstream regions, as well as previously identified upstream regions that may have been identified in error. Upstream sequences were also identified for a set of previously uncharacterized germline gene alleles. Several novel upstream region variants were validated, for instance by their segregation to a single haplotype in heterozygotic subjects. SNPs representing several sequence variants were identified from population data. Finally, based on the outcomes of the analysis, we define a set of testable hypotheses with respect to the placement of particular alleles in complex IGHV locus haplotypes, and discuss the evolutionary relatedness of particular heavy chain variable genes based on sequences of their upstream regions.

## 1 INTRODUCTION

Immunoglobulins play a vital role in recognition of pathogens, thereby enabling their removal or modification of their activities or functions. The typical antibody consists of two identical heavy (H) chains and two identical light chains, of which the H chain often plays a dominant role in determination of specificity (1). The diversity of antibody H chains is established by somatic recombination of immunoglobulin variable (IGHV), diversity (IGHD) and joining (IGHJ) genes, along with junction diversity and somatic hypermutation. Thanks to the development of next-generation sequencing (NGS), it has been possible to describe the nature of the adaptive immune receptor repertoire (AIRR), both in general terms and in relation to e.g. infectious disease, autoimmunity and allergy. Furthermore, it has been possible to approach features of AIRR at a personalized germline gene level as a key factor in the nature of developing immune responses (2). The importance of the personal germline gene repertoire for the development of specific antibodies may indeed be substantial, in particular in view of the importance of stereotyped (public) immune responses against a number of antigens (3).

The germline gene repertoire that encodes final, processed, complete antibody variable domains is extensively described and addressable by bioinformatic tools (4,5). The IMGT (the international ImMunoGeneTics information system) database (6) has developed into a recognized collection of germline genes for analysis of T and B cell AIRR. Despite the development of techniques visibly expanding our knowledge of germline gene variants, the reference database of such genes still cannot be considered to be complete and accurate (7). Importantly, however, long-read sequencing (8) and other NGS technologies and bioinformatics approaches (9–12) now allow us to generate extended, and personalized databases that in the future will enable better, high-quality analysis of AIRR as they develop in health and disease.

Features used to generate antibody repertoires, other than the nucleotide sequence of the product-encoding part of germline genes, are less well defined, studied and understood. Yet, they may play a role in gene expression and generation of a functional antibody repertoire. These include the 5’-untranslated region (5’UTR), the leader sequence encoding the signal peptide that play a vital role in protein transport (13–15), introns of immunoglobulin genes, 3’-non-coding regions including the recombination signal sequence, and more distant regulatory elements (16). Bioinformatic tools developed for studies of large transcriptomic repertoire data sets, such as IgDiscover (9) and IMGT/HighV-QUEST (17), are already able to capture parts of the 5’UTRs and the signal peptide-encoding part of the genes in many existing NGS data sets. Recent studies, however, have suggested that the diversities of 5’UTR and leader sequences are not well represented in the IMGT database (14,18), strongly arguing that such information ought to be updated to enable analysis of the role of these regions in gene expression and functionality. Heterozygosity in 5’UTR and leader sequences may also be used in sequence haplotyping efforts to assess gene expression from individual chromosomes (19,20) even in cases when their associated IGHV genes are identical, thereby allowing further development of our understanding of these genes in a broader context.

NGS-derived AIRR data generated from B cell lineage transcriptomes are now made available for analysis at a large scale. Many such data sets have been generated using 5’-RACE (rapid amplification of cDNA ends) technology and thus incorporate part of the 5’UTR and the entire leader sequence (but not its intron). Proper data pre-processing is important to accurately identify the 5’UTR and leader sequence of the genes. In this study, we developed a tool to remove 5’-barcodes and the homopolymeric tail introduced during the sequencing library generation process, to enable pre-processing of an NGS data set of antibody repertoires. Immunoglobulin germline gene repertoires were inferred by IgDiscover (9) and 5’UTR-leader sequences were subsequently used to infer consensus 5’UTR-leader sequences of each IGHV gene/allele. We explored haplotype analysis, third complementarity-determining region (CDR3) length distribution patterns, and population genome data to validate inferred sequences. Several novel examples of diversity in these upstream regions are described and discussed. Our findings extend the reference database of validated 5’UTR-leader sequences, and the study provides a new pipeline to infer and analyze the upstream sequences of IGHV-encoding genes, a pipeline that likely can be adapted to assess AIRR diversity in other transcriptome data sets.

## 2 MATERIALS AND METHODS

### 2.1 Data set

A publicly available NGS data set of antibody repertoires, first published in a study by Gidoni et.al (19), was analyzed in this study. The data set was obtained from the European Nucleotide Archive (ENA) under the accession number PRJEB26509. It contains reads of antibody heavy chain transcript data generated from naïve B-cells of 100 individuals in Norway. The 100 subjects comprise 52 patients with celiac disease and 48 healthy controls. Sequencing had been performed by a 300*2 paired-end kit by Illumina MiSeq. As in the study of Mikocziova et al. (18), data of two subjects (ERR2567273 & ERR2567275) were here excluded due to low sequencing depth.

### 2.2 Data pre-processing

The sequencing library of the used data set had been generated by 5’-RACE technology, where terminal deoxynucleotidyl transferase (TdT) typically is used to extend cDNA with a homopolymeric tail at its 3’-end, a tail that subsequently can be used to add an adaptor sequence. Details, such as what adaptor sequences that were used, were not stated in the original paper (19), but through inspection of a set of random raw reads, we identified that the sequences, upstream of the antibody genes, incorporated a random barcode sequence followed by a TAC-G_n_ adaptor. Using an in-house developed tool, we trimmed the 5’ end of the forward sequences, up to and including the homopolymeric tail (Figure 1). In addition, forward reads were removed entirely if the sequence TAC-G_3_ was not found within their first 40 bases. The PairSeq.py tool of the pRESTO 0.6.0 software (21) was subsequently used to combine remaining forward reads with reverse reads, into full-length sequences.

**Figure 1.**
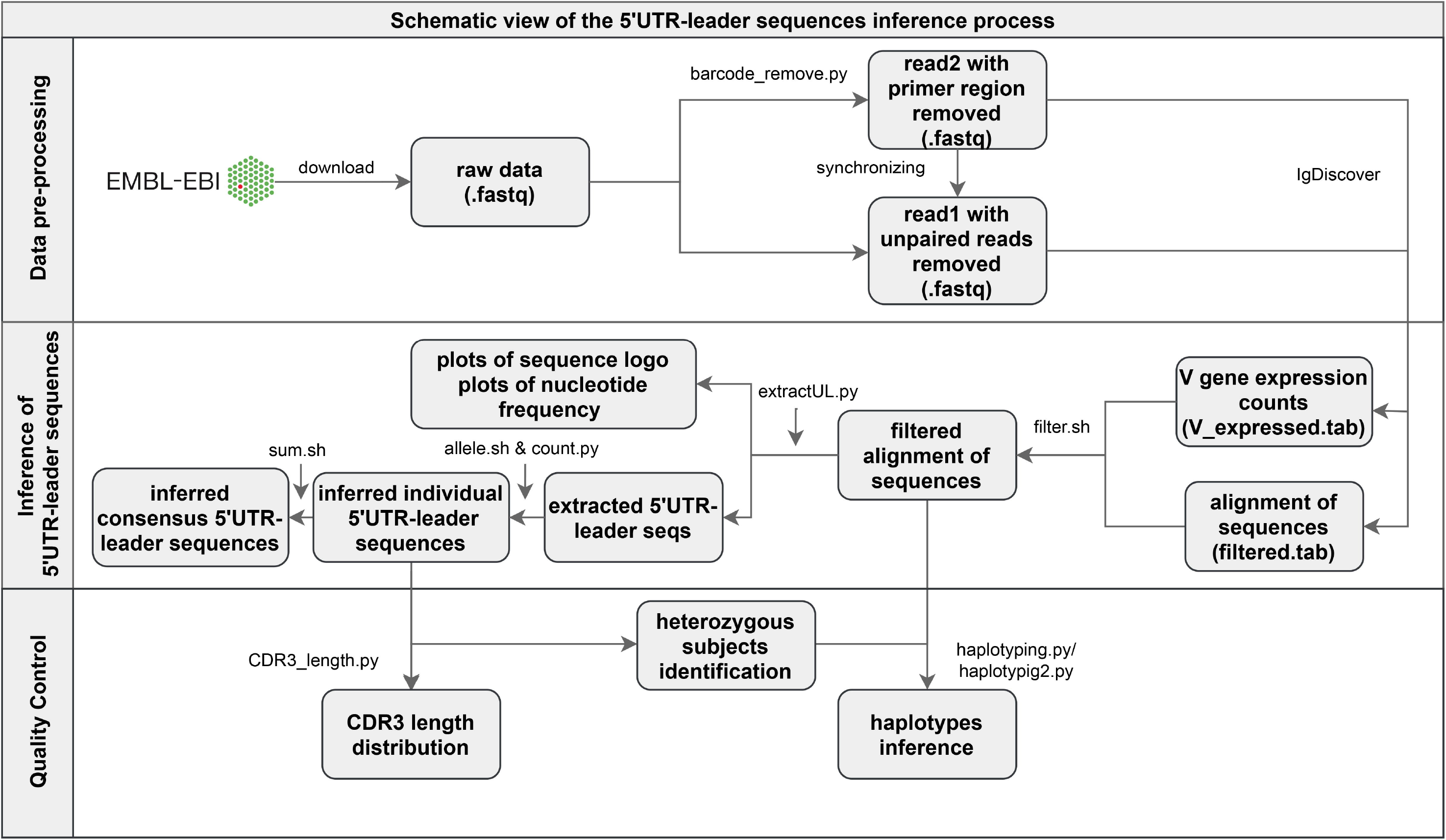
Schematic illustration of the pre-processing of Illumina MiSeq paired-end reads and of the pipeline of 5’UTR-leader sequences inference and validation process.

### 2.3 Germline gene inference and data filtering

The pre-processed sequences were analyzed by IgDiscover 0.12 (9), using default parameters, in order to infer personalized germline gene repertoires. Reference databases of human IGHV, IGHD and IGHJ genes were obtained from the IMGT database (release 202011-3) (6) (Supplementary Data 1). Thereafter, we filtered the IgDiscover processed reads by removing entries with V_errors > 0, as these either have been subject to sequencing errors or somatic hypermutations, or assigned to the wrong IGHV germline gene allele (Figure 1). Additionally, any germline gene allele with low diversity (defined as fewer than 75 unique CDR3s, all entries considered) or low number of assigned reads (less than 20, only entries with V_errors > 0 considered) were excluded, for each individual separately.

### 2.4 Extraction of 5’UTR-leader sequences

After data filtering, we extracted 5’UTR-leader reads, grouped them according to inferred IGHV germline gene allele, and inferred 5’UTR-leader sequences for each analyzed individual and allele (Figure 1). Sequences were built one position at a time, starting at the 3’ end, by extracting any nucleotide present in at least 30% of the reads. Whenever two nucleotides met this threshold, the reads were split accordingly and analyzed separately, starting over from the 3’ end. Length of inferred 5’UTR-leader sequences were set so that at least 50% of the underlying data covered the 5’ most inferred base.

We subsequently summarized 5’UTR-leader sequences of all analyzed individuals, each IGHV allele separately, and counted their frequencies (Figure 1). For sequences that were identical in different individuals, except with respect to how far in the 5’ direction they stretched, the length was set so that at least 80% of the inferred sequences would be of the same length or longer than the consensus sequence. Six upstream region sequences were removed from the output data. The majority of these expressed one extra base in a homopolymeric stretch, a type of region where sequencing insertion errors are not uncommonly seen (22), in either the leader region (three sequences), thus resulting in frame shift, or in the 5’UTR region (one sequence). One of the other two removed sequences showed remains of an adaptor sequence that had not been removed by the pre-processing trimming step, and the other showed low CDR3 diversity.

The upstream regions sequences are numbered in a way that assigns the last base of the leader sequence as base −1. Most upstream regions encode a 19 amino acids long signal peptide. Consequently, the initiation ATG codon will (with the exception of IGHV3-64*01 and IGHV6-1*01) be represented by bases −57 – −55, and the 5’UTR will extend beyond base −57.

### 2.5 Validation of alleles by haplotype inference and CDR3-length distribution analysis

Haplotype analysis was performed for all 35 of the 98 subjects that are heterozygous in the IGHJ6 gene, by calculating the frequency of 5’UTR-leader sequences found in transcripts derived from each allele of the IGHJ gene (Figure 1). The haplotype inference was conducted for alleles that have heterozygous 5’UTR-leader-IGHV allele sequences for the subjects mentioned above, as well as for some additional genes with novel inferred alleles. Clonal diversity of each inferred 5’UTR-leader sequence was examined, by plotting the distribution pattern of amino acid lengths of associated CDR3’s, using filtered IgDiscover data (Figure 1).

### 2.6 Base −93 of upstream regions of genes of the IGHV4 subgroup

To assess the ability of raw, assembled reads to correctly infer base −93 of upstream regions of genes of the IGHV4 subgroup we collected all reads of 8 subjects assembled by PEAR as part of an IgDiscover process (23). These reads were subjected to an IMGT HighV-QUEST analysis process (IMGT/V-QUEST program version: 3.5.18; IMGT/V-QUEST reference directory release: 202011-3). The reads in these data sets that perfectly matched bases −1 – −92 of the inferred upstream region(s), and that were unequivocally assigned by IMGT/HighV-QUEST to one allele of the gene in question, were collected. The occurrences of base T and G (that typically defined variant upstream regions in a recent study (18) of base −93) of reads associated to each haplotype (as defined by alleles of IGJ6) were counted and the ratio of these reads were calculated. For comparison, reads assigned to IGHV1-46*01/*03 were analyzed in the same manner as an example of variability in sets of reads that typically extend well beyond base −93.

### 2.7 Poorly expressed alleles of IGHV2-70

Assembled reads representing the naïve B cell repertoire of subjects that could be haplotyped based on heterozygocity of IGHJ6 had in the past been generated and subjected to IMGT/HighV-QUEST analysis (23). Rare reads of IGHV2-70 of alleles not directly inferred by IgDiscover were identified, a subset of which was associated to an IGHJ6-defined haplotype that did not express another, more highly expressed allele of this gene. A consensus sequence, also including the 5’UTR-leader sequence, of such reads was identified (23).

### 2.8 Identification of upstream region of IGHV4-59*12 in data sets SRR5471283 and SRR5471284

Raw read files SRR5471283 and SRR5471284, containing IgM library of donor LP08248, created by 5’-RACE technology and sequenced by 454 technology (24), were downloaded from ENA. Sequences of the two sets were merged, and subsequently converted using FASTQ Groomer (version 1.0.4) (25). Leading and trailing bases with a quality below 25 were discarded using Trimmomatic (version 0.32.3) (26). Reads were filtered by quality (quality cut-off value: 25; percent of bases ≥ quality cut-off: >95%), and reads were converted to FASTA files. The resulting reads were used to infer a germline gene repertoire using IgDiscover 0.12 (9). 89 bases of the upstream region of IGHV4-59*12 were identified from this output.

### 2.9 Comparison to upstream regions of IGHV genes of Rhesus macaques

Upstream regions of functional germline genes of the IGHV1 and IGHV3 subgroups of Rhesus monkey (*Macaca mulatta*) were retrieved from the assembled IMGT000064 entry (http://www.imgt.org/ligmdb/view?id=IMGT000064). These sequences were aligned to inferred upstream regions of human IGHV genes/alleles of the same subgroups. The similarity of IGHV gene coding regions of Rhesus monkeys to human IGHV germline genes was also assessed using IMGT V-QUEST (IMGT/V-QUEST program version: 3.5.24; IMGT/V-QUEST reference directory release: 202113-2).

### 2.10 SNPs and population data

VCF files describing single nucleotide polymorphisms (SNPs) in human population data of the 1000 Genomes project (27) were retrieved from the International Genome Sample Resource (Phase 3 release, https://www.internationalgenome.org), and data for any variants with global minor allele frequency (MAF) >1% within the analyzed regions were extracted. For genes not defined in the GRCh37 reference genome, but in the GRCh38 reference genome, data were obtained from the Ensembl Genome Browser (releases 102-103; http://www.ensembl.org) (28).

### 2.11 Linkage disequilibrium

Linkage disequilibrium of alleles of IGHV1-2, IGHV1-3, IGHV4-4, and IGHV7-4-1 has been studied in the past (23). The 5’UTR-leader sequences of several of the alleles that occupy these genes were determined in the present study. Some of the alleles of these genes, however, are very poorly expressed and thus cannot be inferred. The conventional haplotype inference was thus extended by past observations of rare transcripts in these transcriptomes, transcripts that suggest the presence of these poorly alleles in haplotypes that lack expression of other, highly expressed alleles (23). In order to extend the previous analysis of linkage disequilibrium of alleles of IGHV1-2, IGHV1-3, IGHV4-4, and IGHV74-4-1, the expected frequency of each haplotypic combination of these 4 genes was calculated, assuming random association, and compared with the observed frequency of the same combinations. Only haplotypic combinations observed in at least 2 of 70 haplotypes were considered. Calculation of expected frequencies was based on the separate occurrence frequency of each 5’UTR-leader-allelesequencew ithin the studied haplotypes.

## 3 RESULTS

### 3.1 Inference and validation of 166 5’UTR-leader sequences by a novel analysis pipeline

In order to address the incomplete representation of 5’UTR and leader sequences of antibody genes in the IMGT database (6), we have examined such sequences in a publicly available antibody transcript data set of 98 individuals (19), also analyzed for the same purpose in the study by Mikocziova et al. (18). Using a strict pre-processing and filtering pipeline followed by extraction of consensus 5’UTR-leader sequences (Figure 1), we identified 166 sequences, found in frequencies ranging from 1 individual to 98 individuals (Figure 2; Supplementary Table 1; Supplementary Data 2). A 5’UTR-leader sequence detected by an inference tool as defined in the present study should feature particular characteristics to be considered valid. Firstly, one would expect that these sequences should be present in a number of different rearrangements, for instance as evidenced by their association to a diversity of lengths of the third complementarity determining region (CDR3). Thus, for each 5’UTR-leader sequence we generated a plot of the number of unmutated reads vs. the length of CDR3 (Figure 3; Supplementary Figure 1), demonstrating that each inferred 5’UTR-leader sequence was associated to a diversity of rearrangements. Secondly, haplotyping offers an important tool to assess the outcome of an inference process (20); the inferred 5’UTR-leader sequences should typically be associated with a single haplotype in subjects that are heterozygous or hemizygous for a given 5’UTR-leader-IGHV gene combination. As illustrated for 5’UTR-leader sequence variants associated to IGHV4-4*02 and IGHV4-4*07 (Table 1), as well as for other 5’UTR-leader IGHV genes that were found in IGHJ6 heterozygous subjects (Supplementary Table 2), this proved to be the case. Thirdly, diversified positions in the 5’UTR-leader sequence of an IGHV gene could also be expected to be represented in genomic data. Population data as described in the Ensembl database (https://www.ensembl.org) has typically been generated by short read sequencing and thereby suffer from important technical caveats that may compromise the correct assembly of complex loci like those representing immunoglobulin germline genes (29). Nevertheless, such data may provide complementary information to other methods, like sequence inference. Analysis of population data of the 1000 Genome Project (27) confirmed that many of the variants seen in the inferred 5’UTR-leader sequences also were represented in the genomic data (Supplementary Table 1). Altogether these findings support the validity of the inferred 5’UTR-leader sequences.

**Figure 2.**
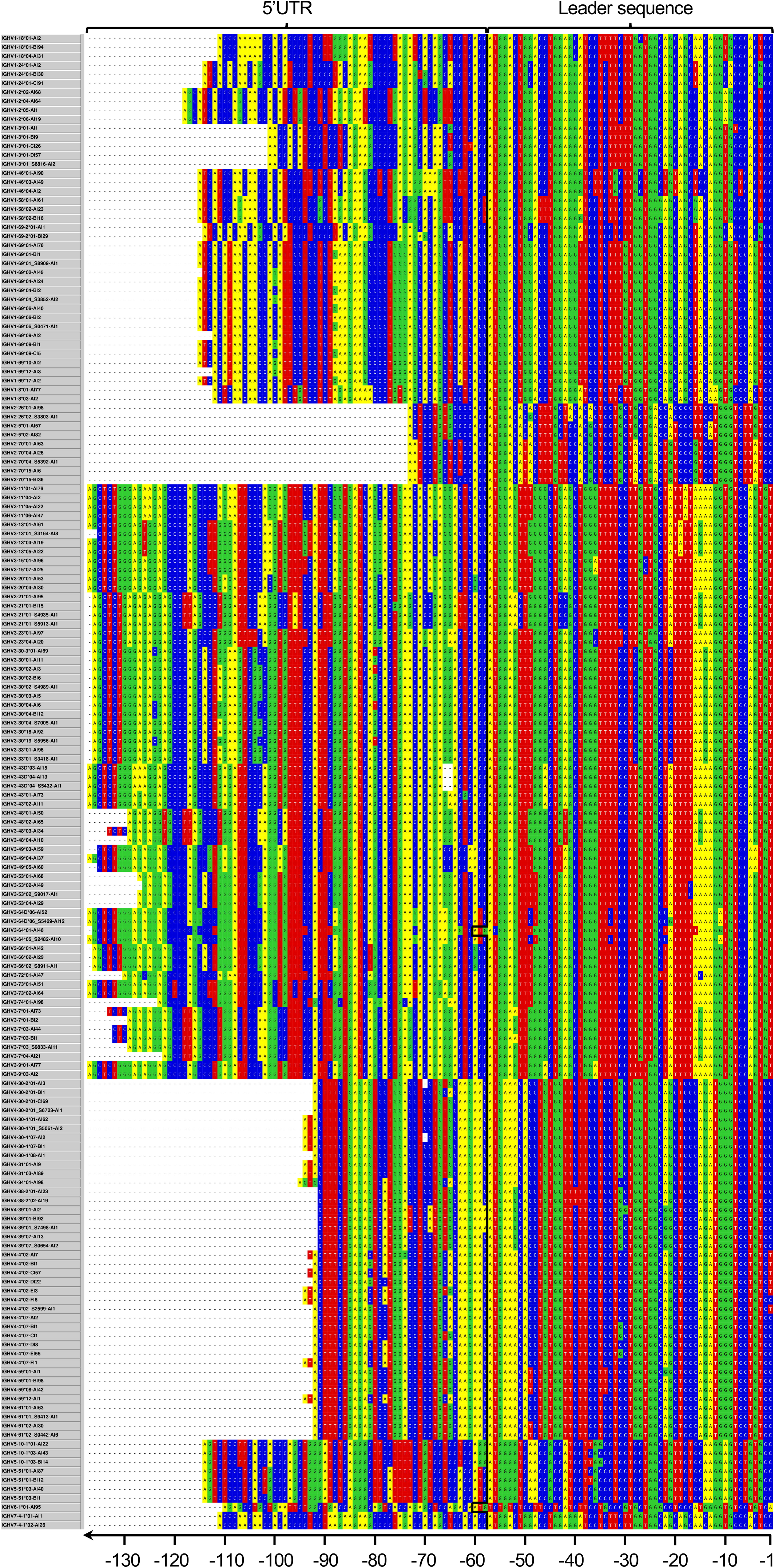
Overarching 5’UTR-leader sequence germline data set inferred in the present study. In addition, upstream regions of IGHV1-3*02 and IGHV4-4*01 have been identified in a separate study (23).

**Figure 3.**
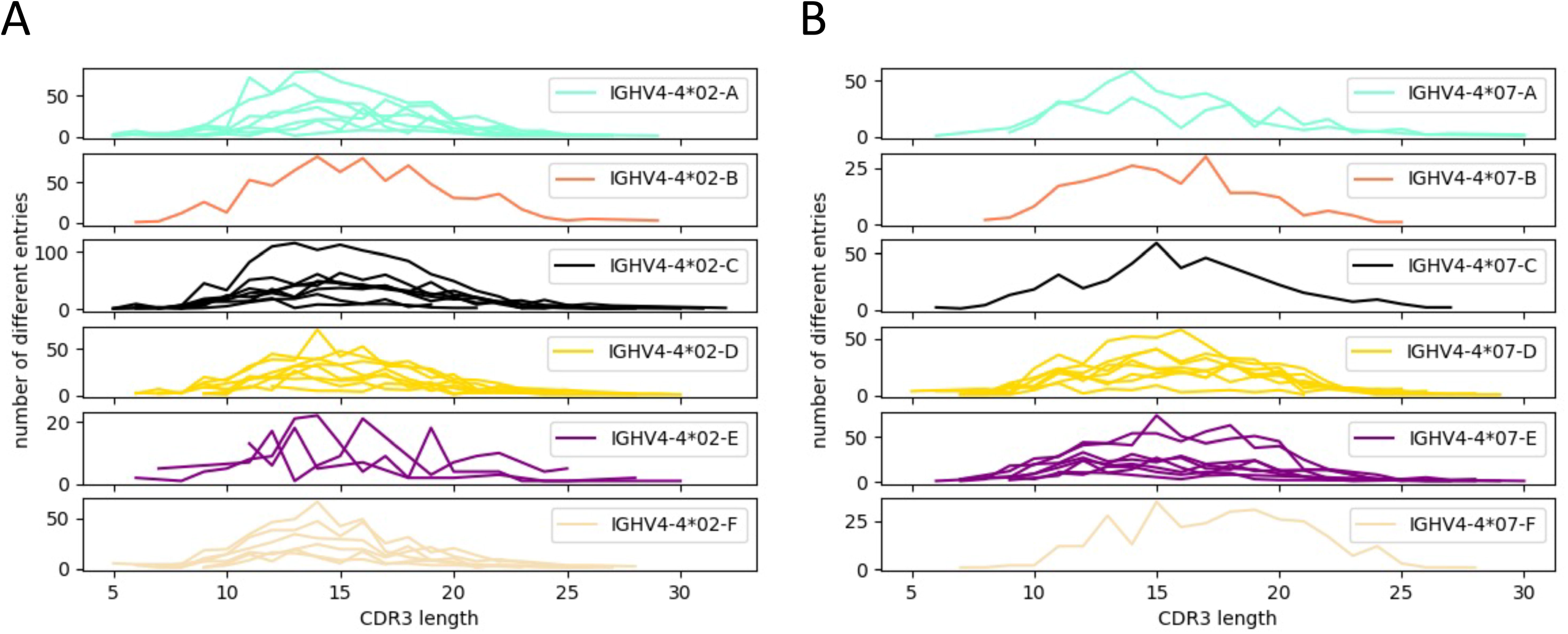
Distribution patterns of CDR3 length encoded by transcripts associated to 5’UTR-leader sequences of (A) IGHV4-4*02, (B) IGHV4-4*07. For each 5’UTR-leader sequence of a specific allele, the number of filtered reads in each length of CDR3 was counted to create the plots. Every line in the plots represents the 5’UTR-leader sequence from one subject (at maximum 8 subjects were included in each plot). Distribution patterns of CDR3 length for 5’UTR-leader sequences of other alleles are displayed in Supplementary Figure 1.

**Table 1.**
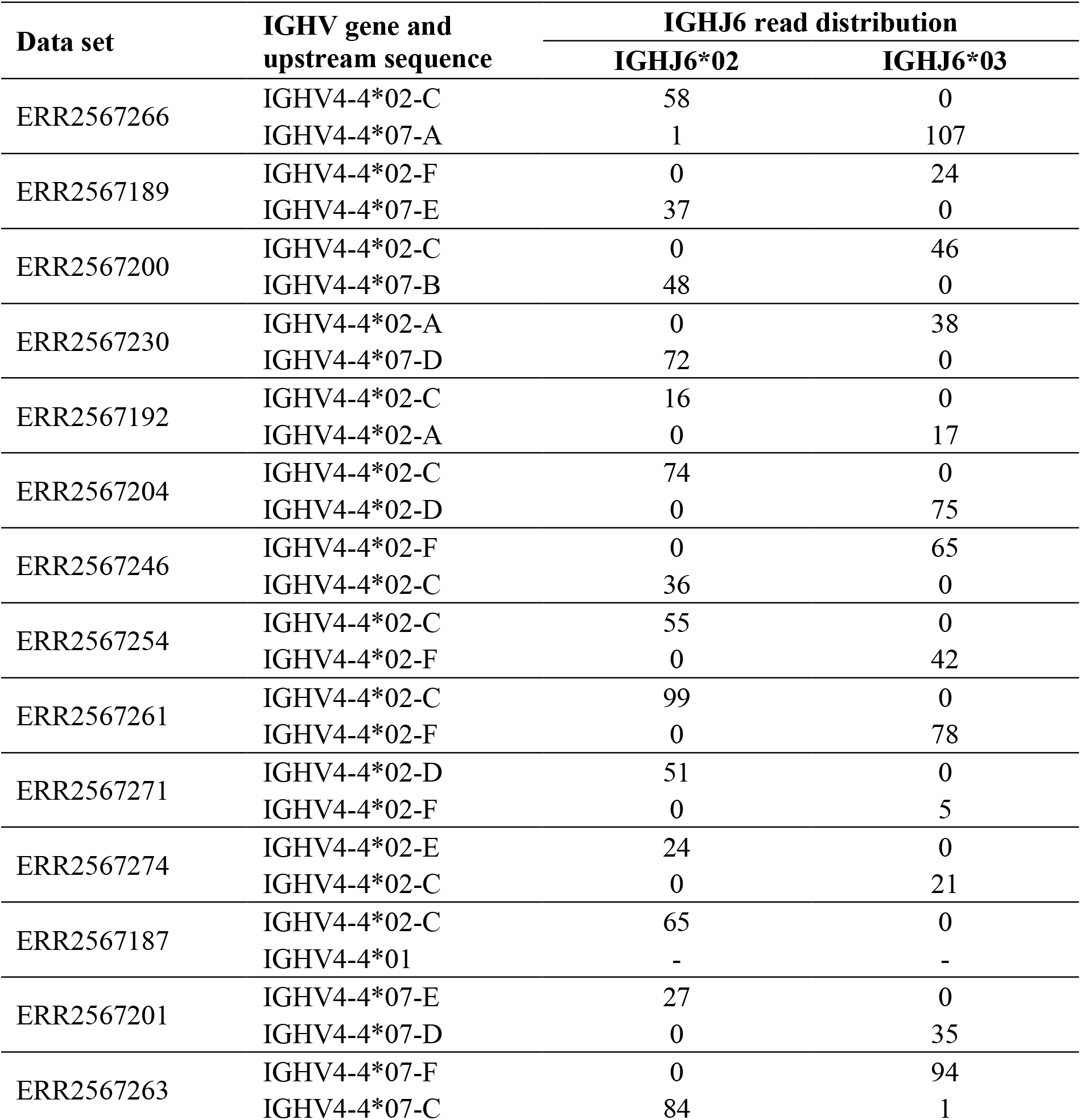
Haplotyping to support the validity of diverse 5’UTR-leader sequence of allele IGHV4-4*02 and IGHV4-4*07. The sequence counts of 5’UTR-leader sequences of alleles of IGHV4-4 associated to different alleles of IGHJ6 in rearranged sequences. Haplotyping data for other 5’UTR-leader sequences are available in Supplementary Table 2.

### 3.2 Novel IGHV alleles

Several novel IGHV alleles have been inferred from the present data set in the past and validated by sequencing of amplified genomic clones (18). These are now featured in more recent releases of the IMGT human IGHV database. Other alleles, some of which had also been identified in the past study but have not yet been entered into the IMGT database, were also identified in the present study. Some of these have independently been reviewed and provisionally accepted by the Inferred Allele Review Committee (https://www.antibodysociety.org/the-airr-community/airr-subcomittees/inferred-allele-review-committee-iarc/), while other alleles have not been identified in the past. The not yet reviewed inferences (IGHV2-70*04_S5392 [A14G], IGHV3-13*01_S3164 [G290A T300A], IGHV3-30*02_S4989 [G49A], IGHV3-30*04_S7005 [C201T G317A], IGHV3-43D*04_S5432 [G4A], IGHV3-53*02_S9017 [C259T], IGHV3-66*02_S8911 [G303A], and IGHV4-30-2*01_S6723 [G70A]) were validated by haplotyping (when possible), CDR3 length distribution, and frequency of unmutated reads based on VDJBase (30) and IgDiscover (9) analyses (Supplementary Table 3). Their upstream regions are now reported (Figure 2).

### 3.3 Conserved 5’UTR-leader sequences of multiple IGHV genes

For multiple genes, the inferred 5’UTR-leader sequences were highly conserved among the alleles of the respective gene. Some genes (such as IGHV3-64, IGHV3-72, IGHV3-74, IGHV4-34, and IGHV6-1) were represented by only one allele that all featured one and the same 5’UTR-leader sequence. Furthermore, all alleles of IGHV1-2, IGHV1-46, IGHV1-8, IGHV2-5, IGHV3-11, IGHV3-13, IGHV3-15, IGHV3-20, IGHV3-23, IGHV3-43D, IGHV3-48, IGHV3-49, IGHV3-66, IGHV3-73, IGHV3-9, IGHV4-31, IGHV4-38-2, and IGHV7-4-1 were associated to one, identical 5’UTR-leder sequence/gene in this cohort.

Assessment of population data (excluding IGHV3-30-3, IGHV3-43D, IGHV3-64D, IGHV4-30-2, IGHV4-30-4, and IGHV4-38-2, as these genes are not featured in any of the reference genomes GRCh37 or GRCh38) confirmed that IGHV1-2, IGHV1-46, IGHV1-8, IGHV2-5, IGHV3-13, IGHV3-48, IGHV3-49, IGHV3-66, IGHV3-72, IGHV4-34, IGHV6-1, and IGHV7-4-1 had no diverse residues (with an overall population MAF>1%) within the sequenced part of the 5’UTR-leader (*i.e*. excluding the leader sequence intron). IGHV3-11, IGHV3-15, IGHV3-20, IGHV3-23, IGHV3-73, and IGHV3-74 all had SNPs that carried variability at high frequency in some populations, although not in European populations (Supplementary Table 1). IGHV3-9 and IGHV3-64 however, expressed variants (−60 [A/G], −88 [A/G], −101 [G/C], and −127 [G/A]; and −56 [C/T], respectively) with MAF>1% also in European population, indicating that the 5’UTR-leader sequences of these genes may contain diversity not captured by our study. However, these genomic variants could potentially also be technical artefact resulting from incorrect assembly of the complex IGHV loci, which sometimes accompany short read sequencing (29). Base −56 of IGHV 5’UTR-leader sequence generally holds the T of the initiation ATG codon, but is represented by an C in the herein inferred 5’UTR-leader sequence of IGHV3-64 (as this gene’s ATG codon is located in position −60 – −58). Thus, incorrect mapping of reads derived from other IGHV genes, including the duplicate gene IGHV3-64D, to the IGHV3-64 region would indeed result in a technical artifact presented as a −56T variant. Likewise, the upstream region of IGHV3-9 is highly similar to e.g. those of IGHV3-20, IGHV3-43 and IGHV3-43D, the latter of which is not even present in the reference genome. It is certainly conceivable that improper assembly of short reads derived from these other genes to the upstream region of IGHV3-9 (Supplementary Figure 2) may contribute to precisely those sequence variants that were defined in Ensembl. Nevertheless, the population-based studies, despite their shortcomings (Watson et al., 2017), generally agreed with the observation of low diversity of these 5’UTR-leader sequences of the herein studied cohort. This analysis, furthermore, also suggested that differences may exist between populations with respect to diversity of the studied upstream region.

### 3.4 Highly diversified 5’UTR-leader sequences of multiple IGHV genes

The 5’UTR-leader sequences of several genes were diverse even after the stricter pre-processing procedure performed prior to the present analysis. Alleles of many genes (like IGHV1-18, IGHV1-24, IGHV1-3, IGHV1-58, IGHV1-69, IGHV2-26, IGHV2-70, IGHV3-21, IGHV3-7, IGHV3-30, IGHV3-43, IGHV3-53, IGHV3-64D, IGHV4-30-2, IGHV4-30-4, IGHV4-39, IGHV4-4, IGHV4-61, IGHV5-10-1, and IGHV5-51) were diverse in the population of this data set. Population-based studies addressing diversity in the 5’UTR-leader sequence was used to examine these variants further. For a majority of these genes (IGHV1-3, IGHV1-58, IGHV1-69, IGHV2-26, IGHV2-70, IGHV3-21, IGHV3-43, IGHV3-53, IGHV3-7, IGHV4-4, IGHV4-61, and IGHV5-51), all identified SNPs could also be observed in the population data (Supplementary Table 1). Many of these variants could also be further validated with haplotype analysis. Haplotyping could also be performed for two of the 5’UTR-leader sequence variants that could not be identified in population data. IGHV1-24 featured three different 5’UTR-leader sequences with diversity in two positions (−70 [A/G] and −71 [C/T]), with only the latter observed in analyzed population data. Yet, haplotyping of one individual, expressing both IGHV1-24*01-A (−70A, −71C) and IGHV1-24*01-C (−70G, −71C) showed appropriate segregation of these upstream regions between the two haplotypes, supporting the inferences (Supplementary Table 2). Similarly, the diversity of base −30 (G/C) in IGHV4-39 associated 5’UTR-leader sequences could not be confirmed by population studies but is supported by haplotype analysis of one individual expressing IGHV4-39*01-A and IGHV4-39*01-B on different haplotypes.

Diversification of 5’UTR-leader sequences can be limited to a single base (*e.g*. for IGHV1-18 and IGHV3-7) or include variability in multiple bases. One of the genes expressing the most diversified 5’UTR-leader sequences within the analyzed population is IGHV4-4, which is dominated by two quite different alleles, IGHV4-4*02 and IGHV4-4*07. These alleles together carry diversity located to seven positions (bases −1 [C/T], −31 [C/G], −65 [A/G], −66 [C/T], −74 [A/G], −78 [A/C], and −81 [C/G]), four of which are diverse in both alleles as defined by the present study. This diversity corresponded well to diversity seen in the 5’UTR-leader sequence of the gene as investigated in population studies (Supplementary Table 1). Additionally, haplotype analysis provides further evidence for most of the identified 5’UTR-leader sequences (Supplementary Table 2). Despite the substantial divergence of the two alleles’ coding regions, the 5’UTR-leader sequences are similar and several of their diversified 5’UTR-leader sequence residues carry similar type of diversification. In all, six different 5’UTR-leader sequences were found associated to each of these alleles of IGHV4-4, several of which were not identified in a previous study of the present data set (Supplementary Figure 3).

Some germline genes may, due to their high similarities, be hard to distinguish between in *e.g*. germline gene inferences and population-based studies. One example of such very similar germline genes is IGHV3-30, IGHV3-30-3, IGHV3-30-5, and IGHV3-33. We identified five different 5’UTR-leader sequences among the alleles of these genes with variability in bases −80 (G/T), −103 (G/C), −111 (G/A), and −124 (G/C). The 5’UTR-leader sequences of alleles like IGHV3-30*02, IGHV3-30*18, and IGHV3-33*01 share common sequence features while alleles like IGHV3-30*01, IGHV3-30*04, and IGHV3-30-3*01 shared another set of related sequence features (Supplementary Figure 4). Population-based studies using short read sequencing technology is complicated and error-prone (29), in particular in relation to sets of very similar genes, like these. In any case, analysis of data of these three genes from the 1000 Genome Project provides further evidence for two of the identified variable positions (−80 and −103) of the 5’UTR-leader sequence of IGHV3-30 (Supplementary Table 1). One additional SNP (−40 [G/T]) could be identified in the population data of IGHV3-30, but had a low MAF (<1%) in the European population. Another set of highly similar genes is IGHV4-30-2, IGHV4-30-4, and IGHV4-31. The 5’UTR-leader sequences of alleles of these genes are mostly identical. Only 2 and 1 rare sequence variants of these upstream regions were identified in IGHV4-30-2 and IGHV4-30-4, respectively. In contrast to other variants seen in this study two of these sequences represented base deletions, in both cases Δ-69C. Haplotyping of such upstream regions of IGHV4-30-2 was possible using one data set, in which case the haplotype with or without base −69 separated onto different haplotypes (Supplementary Table 2), supporting the validity of the inference. The frequency of these transcripts in the data sets suggested that they were expressed at similar levels as those alleles that had not deleted this particular base in the 5’UTR (0.39%±0.05% [n=3] and 0.45±0.16% [n=4], respectively). Population-based studies provided further validation of this deletion, as one such variant was identified for IGHV4-31 (Supplementary Table 1).

### 3.5 5’-terminal Gs in inferred 5’UTR-leader sequences

In contrast to the study by Mikocziova et al. (18), 5’UTR variants with a 5’-terminal G were largely eliminated in our analysis, a direct result of the strict 5’ trimming process used. As a consequence we, in several instances, inferred a 5’UTR that was shorter than that identified by Mikocziova et al. (18). For instance, in the case of alleles of IGHV2-5, only one common upstream sequence was identified in the present study, while Mikocziova et al. (18) identified two common, longer upstream sequences for each allele, with only a T/G difference in the 5’-most base (position −75) (Supplementary Figure 5). Population data suggest that base −75 is virtually invariant (T) in human populations (highest population minor allele frequency (MAF)<0.01%), suggesting that an inferred G variant may be a technical artifact. Similarly, in our hands and using a strict pre-processing protocol, 5’UTR-leader sequences with a length of 92 bases were typically inferred for many genes belonging to subgroup IGHV4, while alternative 5’UTR-leader sequence variants that carry either a G or a T at base −93 had been identified for many such genes in the past (18). Again, many of the alleles that had previously been suggested to carry a variant with a 5’-terminal G showed no evidence of such common SNPs in population studies (Supplementary Table 4).

To further study the matter of diversity in the 5’-most base of inferred 5’UTRs, we assessed the nature of the raw data generated in the sequencing process and its relation to a possible outcome of the inference process. Assessment through haplotyping of unprocessed reads associated to genes of subgroup IGHV4 frequently demonstrated that sequences carrying both bases at position −93 were in general associated to both haplotypes of each subject (Supplementary Table 5). Such observations indicate that the haplotyped individuals can only be heterozygous in position −93 if these genes are duplicated on both haplotypes, a requirement that is at odds with our current understanding of the locus. Altogether, these investigations suggest that further studies (such as long read sequencing) are required to provide evidence of the existence of many variants of 5’UTR-leader sequences with 5’-terminal Gs.

### 3.6 Uncommon 5’UTR-leader sequences in the IMGT germline database

It has previously been reported (14,18) that several 5’UTR-leader sequences associated to IGHV germline genes do not correspond to the sequence of the primary entry found in the IMGT database. We confirm this in several cases, such as for IGHV2-5*01, IGHV3-23*01, and IGHV5-51*01 (Supplementary Figure 5, Supplementary Table 6). Population data support that the sequences reported by us and others represent the real upstream sequences while the primary entries of the IMGT database are incorrect or represent vary rare sequence variants (MAF<0.01%) not representative of many populations. Interestingly, the common leader sequence of genes like IGHV2-5*01 and IGHV3-23*01 is represented in the IMGT database as secondary sequence entries. Such more representative 5’UTR-leader sequences are however not readily retrieved as one download upstream regions from the database.

### 3.7 5’UTR-leader sequences as a resource for defining genotype organization

Alleles of IGHV genes are commonly given a name associated to the closest known sequence even when the precise genomic location of these alleles might not be known. Some genes might thus be associated by name to a gene where it does not reside. Upstream regions might provide indications of gene relatedness beyond the sequence of the final product. 5’UTR-leader sequences of all identified alleles of the IGHV4 subgroup identified in the present study were consequently aligned to each other. The sequence of some alleles of IGHV4-4 are very similar to alleles of other genes (Supplementary Figure 6). One of the upstream regions, IGHV4-59*12-A identified in data set ERR2567237, was shown to be most similar to some of those of IGHV4-4. In fact, it was identical to IGHV4-4*02-F and IGHV4-4*07-D (Supplementary Figure 6). IGHV4-59*12 (https://ogrdb.airr-community.org/genotype/32) was originally identified in a data set (24) different from those assessed here. The 5’UTR-leader sequence of IGHV4-59*12 found is this genotype (donor LP08248) differed by one base from IGHV4-59*12-A, and it was identical to that of IGHV4-4*07-E (Supplementary Figure 6). Haplotyping of this genotype suggested that IGHV4-59*12 resided on a haplotype that apparently lacked an allele of IGHV4-4 but had alleles of IGHV4-59 and IGHV4-61. There are thus two instances of IGHV4-59*12 with leader sequences more similar to those of IGHV4-4 than to those of IGHV4-59, and circumstantial evidence through haplotyping that suggests that IGHV4-59*12 might very well be located in IGHV4-4.

5’UTR-leader sequence can also provide valuable information that can aid in the understanding how an individual’s IGHV loci are composed. For example, one genotype (defined by data set ERR2567264) carries IGHV1-69*02 and IGHV1-69*06, that through haplotyping were shown to segregate onto different haplotypes. Allele IGHV1-69*06 was, however, associated to two different upstream regions (Supplementary Table 2). This finding suggests that allele IGHV1-69*06 may occupy both gene location IGHV1-69 and IGHV1-69D. The inference of the germline gene repertoire of the data set ERR2567237 also demonstrated unusual features, in this case of IGHV4-30-2 and IGHV4-30-4. One allele of IGHV4-30-2 was inferred, but it was associated to three different 5’UTR-leader sequences, while three different alleles of IGHV4-30-4 were inferred. This suggests that both genes are duplicated, either both genes on one haplotype, or one gene on each haplotype. Alternatively, one of the alleles might be located at the site of another gene. Analysis of 5’UTR-leader sequences can thus provide additional evidence of genotype organization, in this case related to duplicated genes, not assessable by analysis of the coding region alone.

The part of the locus spanning from IGHV1-69 to IGHV2-70 is highly complex as it commonly harbors a large duplication (Watson et al, 2013) and numerous allelic variants of these genes. The present analysis inferred 8 alleles of IGHV1-69(D) and only 3 alleles of IGHV2-70(D) in 35 subjects in which the IGHV locus could be haplotyped based on heterozygocity of IGHJ6 (20). Assessment of the haplotypes identified in the present investigation identified four main types of expressed gene combinations in this part of the locus, as defined by the coding regions and their upstream sequences (Supplementary Figure 7A, C). IGHV2-70*15 was linked to two different 5’UTRs (Supplementary Figure 7D), that associated to different genomic contexts, with and without the duplication involving IGHV1-69D, IGHV1-69-2, and IGHV2-70D (Supplementary Figure 7A). Genomic sequencing has in the past identified haplotypes resembling some of the differences in upstream regions of genes in this part of the IGHV locus (Supplementary Figure 7B). Future descriptions of haplotypes of different populations will likely be required to understand the diversity of this complex part of the IGHV locus.

IGHV1-69-2 and IGHV2-70/70D are commonly expressed at relatively low levels (Gidoni et al, 2019), and may thus escape inference in samples with fewer reads and limited sequence complexity. In haplotypes expressing only a single copy of IGHV1-69/69D with upstream region sequence featuring −88A −100G, it was common not to infer an occurrence of IGHV2-70. Detailed analysis of IMGT/HighV-QUEST output of reads of IGHV2-70 nevertheless identified poorly expressed variants of IGHV2-70 in some cases, even alleles that are not currently defined or incomplete in the IMGT database (Supplementary Figure 7E). One of these alleles also carries a variant upstream region sequence (Supplementary Figure 7D) not seen in the other, more highly expressed alleles of this gene. Genomic sequencing has in the past identified a similar allelic variant of IGHV2-70 (GenBank accession number AC242528), an allele that is not yet featured in the IMGT database, that also encoded multiple unusual sequence modifications, including for instance an unusual cysteine in framework 3 (Supplementary Figure 7). Altogether, it is highly likely that at least some subjects carry an IGHV2-70 allele in their genotype that could not be efficiently detected by transcriptome-based sequencing and germline gene inference technology. Future identification and confirmation of such alleles and studies of their functionality will be required to allow us to understand their contribution to human functional antibody repertoires.

### 3.8 Role of the IGHV4-4*01 5’UTR-leader sequence in the poor expression of this allele

Allele IGHV4-4*01 has recently been identified as being very poorly expressed (23), and consequently difficult to infer using tools like IgDiscover. As a consequence of these technical aspects, it was not detected in the present study. The allele’s unusual protein sequence was proposed as the cause of its poor expression. Its 5’UTR-leader sequence (23) differs from the corresponding upstream regions of prototype highly expressed alleles IGHV4-4*02 and IGHV4-4*07 as defined in the IMGT database. With the present collection of novel 5’UTR-leader sequences of highly expressed alleles of IGHV4-4, it was possible to further assess the extent whereby these regions might also explain the poor expression of IGHV4-4*01. Indeed, the upstream region of IGHV4-4*01 (23) is identical to IGHV4-4*02-A, an upstream region identified in 7 subjects in the herein investigated data set. This upstream region, in combination with IGHV4-4*02 (0.92%±0.37%) expressed similarly with the other allele of IGHV4-4 (0.90%±0.13%) (n=7) in the same subject, suggesting that this upstream sequence is not responsible for the poor expression of IGHV4-4*01. Furthermore, while transcripts derived from IGHV4-4*01 are largely non-productive (23), transcripts derived from IGHV4-4*02 in combination with the IGHV4-4*02-A upstream region typically encoded an in-frame product. There is thus no evidence to suggest that the herein assessable upstream region of IGHV4-4*01 is responsible for the poor expression of this allele.

### 3.9 Length differences in the inferable part of the 5’UTR

Insertions and deletions (indels) may serve as markers to assess the evolution of genes (31). Inspection of the 5’UTR of genes belonging to the IGHV3 subgroup suggests that they have evolved by indels resulting in length differences in this region (Supplementary Figure 8A). For instance, alleles of gene IGHV3-43D (but not alleles of the related gene IGHV3-43) all lack bases −65 and −66 of the 5’UTR-leader sequence of other alleles. Similarly, alleles of IGHV3-23, IGHV3-30, IGHV3-30-3, IGHV3-53, and IGHV3-66 all lack base −121 of other 5’UTR-leader sequences, while alleles of IGHV3-7, IGHV3-21, and IGHV3-48 lack base −109 present in other 5’UTR-leader sequences. In contrast, all these bases are present in IGHV3-9, IGHV3-11, IGHV3-13, IGHV3-15, IGHV3-20, IGHV3-43, IGHV3-49, IGHV3-64, IGHV3-64D, IGHV3-72, IGHV3-73 and IGHV3-74. It is conceivable that these groups of genes have a common evolutionary history. We also compared the upstream regions of inferred human IGHV genes with the small set of functional genes of Rhesus macaques as defined in the IMGT database entry IMGT000064. We identified a number of length differences in the 5’UTR-leader sequences of such functional genes, including such identical to those found in human genes (Supplementary Figure 9A). Length differences were also observed in the 5’UTR of genes belonging to the IGHV1 subgroup (Supplementary Figure 8B) affecting for instance base −76. In similarity to the case of IGHV3, a similar indel event was observed in the upstream region of Rhesus macaque IGHV1 subgroup genes (Supplementary Figure 9B). The upstream region of macaque allele IGHV1-111*01 carried an indel event identical to that of the upstream region of human gene IGHV1-69-2*01. In addition, the human germline gene most similar to the V domain coding sequence of IGHV1-111*01 was IGHV1-69-2*01. In this case the close similarity of human and macaque IGHV germline genes in terms of indels in their upstream region sequences, was associated to a similarity of the coding sequences of these genes as well.

### 3.10 Linkage disequilibrium

We have previously identified a possible linkage disequilibrium in the IGHV locus that associates IGHV1-2*05 to IGHV4-4*01 (23). We now extended this finding by assessing the association of alleles and diverse upstream regions of these genes to each other and to alleles of IGHV1-3 and IGHV7-4-1, genes that are located close to each other on chromosome 14. This was made possible by analysis of 35 haplotypable data sets of the herein analyzed set of data. All cases of IGHV1-3*01 with upstream region C, and all cases of IGHV7-4-1*02 were associated to either all cases of poorly expressed alleles IGHV1-2*05 and IGHV4-4*01, or with all cases of IGHV1-2*06 and 6/7 cases of IGHV4-4*02 with upstream region D. These two gene combinations were found at a frequency >300-fold above those expected from the frequencies of these individual alleles/upstream regions, alone (Supplementary Figure 10). Similarly, IGHV1-2*04 and IGHV1-3*01 with upstream region D were mostly associated to IGHV4-4*02 with upstream region C or F, and poorly expressed allele IGHV7-4-1*01, at frequencies >10-fold higher than those excepted from random associations of the same alleles. Finally, IGHV1-2*02 was in most cases (>10 times more often than expected) linked to poorly expressed allele IGHV1-3*02, and IGHV4-4*07 with upstream region E or D while there was no evidence of expression of IGHV7-4-1 in these haplotypes. These conserved combinations of alleles were, however, found to be associated to a diverse set of alleles of more distal genes in the locus. For instance, the linked combination IGHV1-2*06 – IGHV1-3*01 (upstream region C) – IGHV4-4*02 (upstream region D) – IGHV7-4-1*02 was seen in haplotypes that carried IGHJ6*02 and IGHV1-69*02, or IGHV1-69*03 and IGHV1-69*02, or IGHJ6*03 and IGHV1-69*04, or IGHJ6*02 and IGHV1-69*10, or IGHJ6*02 and IGHV1-69*12). This strongly suggests that the observed linkage disequilibrium was not primarily an artifact caused by a close familiar relationship between several study subjects in the cohort, but rather that the gene combination exists in subjects with otherwise highly different IGHV loci. Altogether, although multiple alleles and upstream regions exist in IGHV1-2, IGHV1-3, IGHV4-4, and IGHV7-4-1, these are largely found only in a limited set of combinations in the herein investigated population (Supplementary Figure 10).

## 4 DISCUSSION

The present investigation has, inspired by recent studies (14,18), further investigated 5’UTR-leader sequences of IGHV genes. By exploring a strict 5’-trimming pre-processing procedure we eliminated strings of 5’-terminal Gs introduced during the sequencing library generation process, as these may result in technical inference artefacts. We also provide extensive validation of many inferred sequences in terms of haplotyping, association to rearrangement with a variety of CDR3 length, and genomic evidence. Several variants of the 5’UTR-sequences identified by Mikocziova et al. (18) are confirmed. We also report additional upstream sequences not identified in that study. Importantly, these studies ([18]; this study) collectively indicated that some primary sequences in the IMGT database do not represent common upstream regions of these genes. It is consequently suggested that this database is updated to better represent typical 5’UTR-leader sequences.

Past investigations in several cases suggested that alternative 5’UTR sequence variants with a 5’-G were proposed to be common in the investigated population (18). These variants could not be confirmed in the present study. Genomic data, generated largely by short read sequencing, further confirmed that these variants are at most rare in human populations. It is, however, certainly difficult to apply such sequencing technology on a highly repetitive locus like that encoding antibody H chain variable domains (29). This is in particular the case in a sequence discovery setting. Examples of particularly complicated cases, such as the upstream region of IGHV3-9, were identified, highlighting the need for caution when interpreting genomic population data. However, such data may also provide independent, supportive information for validation of more common sequence variants (SNPs), as we have demonstrated in this study. Genomic data indeed support many of the variants we have identified and indicate that additional upstream sequence variants, not identified in the present study of data sets collected in northern Europe, may exist in other populations.

Inference analysis cannot provide positional information on inferred sequences. However, inferred sequences may stimulate development of hypotheses that later has to be proven by alternative technologies, such as long-read genomic sequencing (32). In the present study, analysis of upstream regions identifies several genotypes that may represent unusual or previously not well-characterized structures of the IGHV locus. We identified one genotype (data set ERR2567237) that carried three copies of IGHV4-30-2 (as defined by different upstream regions) and three copies of IGHV4-30-4 (as defined by allelic differences in the coding region), suggesting that these closely linked genes may be present in two copies on one haplotype. Furthermore, we defined that allele IGHV1-69*06 may be present (data set ERR2567264) in two copies (with different upstream regions) within a single haplotype. This suggests that this allele, which frequently occurs in combination with IGHV1-69*01/IGHV1-69D*01, tentatively may occupy both the IGHV1-69 and the IGHV1-69D gene. Inference technology also allowed us to identify tentative linkage disequilibrium between alleles and their upstream regions from IGHV1-2 to IGHV7-4-1, genes that are located in close proximity to each other in the IGHV locus. Whenever particular genes/alleles are associated through such disequilibrium and are linked to particular (stereotyped) immune responses, these characteristics may thus be co-inherited. Finally, we found evidence (in data sets ERR2567237, and SRR5471283+SRR5471284) in its 5’UTR-leader sequence that IGHV4-59*12 may reside in a gene different from that suggested by its name, tentatively IGHV4-4. Indeed, haplotyping of data generated from a subject different from those primarily studied here suggest that IGHV4-59*12 is present on a haplotype that also carries IGHV4-59*01 but no allele of IGHV4-4 (https://ogrdb.airr-community.org/genotype/32). Similarly, IGHV4-4*09 has been discovered in a context in which it exists on the same haplotype as IGHV4-4*03 but in the perceived lack of an allele of IGHV4-61 (https://ogrdb.airr-community.org/genotype/51). These cases mimic the situation of IGHV4-59*08 that typically is present on the same haplotype as IGHV4-59*01. Such haplotypes commonly lack an allele of IGHV4-61. It has furthermore been proposed that IGHV4-59*08 is associated to non-coding regions more similar to those of alleles of IGHV4-61 than to those of other alleles of IGHV4-59 (33). Altogether these studies suggest that IGHV4-59*08 might be located to the IGHV4-61 gene location. Although not proof of these alleles’ location in the locus, findings through inference certainly stimulate debate on the organization of the locus, and the principles of allele naming that are currently in use (10,34,35).

Upstream regions of alleles/genes belonging to the same IGHV subgroup tend to be similar, suggesting a common origin. Different genes are though different in terms of diversity of the alleles and its upstream regions. For instance, IGHV1-2 of the present data set features a number of slightly different, highly expressed alleles (IGHV1-2*02, IGHV1-2*04, IGHV1-2*06, and IGHV1-2*07), and one poorly expressed allele (IGHV1-2*05) (23), but they are all associated to the very same upstream region. In contrast, IGHV4-4 is dominated by two quite different alleles (IGHV4-4*02 and IGHV4-4*07; only 92% base identity, and difference in the length of CDRH1) that show more similarity to alleles of other genes of the IGHV4 subgroup than to each other. These two very different alleles of IGHV4-4 are associated to six upstream regions each with similar sequence features, one of which are even shared between them (Supplementary Figure 3). Is the upstream region of some genes like IGHV1-2 less amendable to diversification than the upstream region of IGHV4-4? Was IGHV4-4 populated by independent duplications of other genes, or have processes like gene conversion contributed to the present diversity of this gene and its alleles, and the similarity of their associated upstream regions? Future phylogenetic and experimental studies are required to address these matters properly.

Although small differences in the expression of alleles of an IGHV gene have been identified, many alleles of a single gene tend to be expressed at similar levels (19,36). Indeed, such similarity is frequently used as a gatekeeper by germline gene inference tools to eliminate inference of sequence variants that are artifacts of PCR and sequencing errors, or somatic hypermutation (37). However, we recently described a number of very poorly expressed alleles (23). These alleles all encoded residues within the variable domain not found in other germline genes. We hypothesized that these alleles were poorly expressed as their encoded product in general would be non-functional. B cells encoding such antibodies would rarely be selected as their products would not be able to participate in a positive selection process. IGHV4-4*01 was one such poorly expressed allele. Assessment of its upstream region as detected in the few transcripts that were present in IGHV-encoding transcriptome suggested that it differed from upstream regions of other alleles of IGHV4-4, as defined in the IMGT database. It was thus plausible that this region associated to IGHV4-4*01 is responsible for the allele’s low level of expression. However, we herein demonstrated that the upstream region of IGHV4-4*01 assessable by analysis of IGHV transcripts amplified by 5’-RACE methodology is identical to that of a subset of the well-expressed allele IGHV4-4*02. Through this analysis approach we were able to extend the support for the hypothesis (23) that IGHV4-4*01 is poorly expressed, not as a consequence of the upstream region’ sequence, but as a consequence of a compromised ability of its encoded protein product to form a folded protein.

Interestingly, although similar within subgroups, the 5’UTR-leader sequences show some differences not only in sequence but also in terms of sequence length, differences that may relate to insertion and deletion events. For instance, the upstream regions of IGHV3-43 and its duplicated variant IGHV3-43D, differ by the absence of two bases in the 5’UTR of the latter gene. This difference has previously been used to support the naming of previously undefined, inferred allele IGHV3-43D*04, as an allele of IGHV3-43D and not of IGHV3-43 (38). This allele has independently been demonstrated to reside at IGHV3-43D through sequencing of a fosmid clone (http://www.imgt.org/ligmdb/view?id=AC242184). Upstream regions of some alleles of IGHV genes have thus been proven to contain information that can be used to build valid hypothesis about their location in the genome. Other genes of the IGHV3 subgroup differ in the length of their 5’UTR. Indeed, three other major sets of upstream regions that differ by the presence of perceived insertion/deletion events have been identified. Some of the genes grouped together based on the similarity of indels in their upstream regions are quite similar in their coding region while others are quite different in this respect (e.g. IGHV3-21 and IGHV3-48 vs. IGHV3-7, genes that all lack base −109 of the 5’UTR-leader sequence). We propose that the presence of these indels events may identify genes with a common evolutionary history. Intriguingly, identical insertion/deletion differences as those found in the upstream regions of human IGHV1 and IGHV3 genes were identified in a limited IMGT-database-defined set of functional germline genes of *Macaca mulatta* (Rhesus macaque). These findings suggest that either these positions are particularly sensitive to indel events, or that such events might have occurred prior to separation of linages (39) resulting in humans and Rhesus macaques, respectively. As IGHV germline gene repertoires of additional species become available, it might be possible to identify a line of events through which the human IGHV genes and their upstream regions have evolved.

In conclusion, we have generated a collection of validated 5’UTR-leader sequences associated to human IGHV genes in a European human population, a set that may be used for future studies of human IGHV genes. Through this effort we also identified SNPs that indicate diversity in these regions that may exist at high frequency in other populations. We also defined upstream region sequences that may have been identified in error in the past. We describe the extent of diversity of such regions in human germline genes, ranging from the invariable upstream regions of alleles of IGHV1-2 to the highly diversified upstream regions of IGHV4-4. Data on upstream regions were used to build hypotheses regarding for instance allele placement in the IGHV locus, in order to promote further studies of the locus’ structure. Finally, we used length differences in upstream regions of IGHV genes to postulate a model of the gene’s phylogenetic relatedness.

## AVAILABILITY

Raw sequence data files of IgM-encoding transcriptomes are available from the European Nucleotide Archive as project PRJEB26509. Raw sequence files that represent the transcriptome of subject LP08248 are available from the European Nucleotide Archive with accession numbers SRR5471283 and SRR5471284. Code developed in this study is available at https://github.com/yixun-h/5-UTR-leader_Infer.

## ACKNOWLEDGEMENT

The computations and data storage were enabled by resources provided by the Swedish National Infrastructure for Computing (SNIC) at LUNARC and Swestore, partially funded by the Swedish Research Council through grant agreement no. 2018-05973. Part of the study was conducted, presented, and defended by Yixun Huang as a MSc thesis project entitled “Computational inference and analysis of 5’UTR-leader sequence of alleles from immunoglobulin H chain genes”.

## AUTHOR CONTRIBUTIONS

Conception of study: LT, MO. Coding: YH, LT; Analysis: YH, LT, MO. Manuscript preparation and final approval: YH, LT, MO.

## ACKNOWLEDGEMENTS

This manuscript has been released as a pre-print at bioRxiv (https://www.biorxiv.org/) (40)

## REFERENCES

1. Xu, J.L. and Davis, M.M. (2000) Diversity in the CDR3 Region of VH Is Sufficient for Most Antibody Specificities. Immunity., 13, 37–45. doi: 10.1016/s1074-7613(00)00006-6

2. Avnir, Y., Watson, C.T., Glanville, J., Peterson, E.C., Tallarico, A.S., Bennett, A.S. et al. (2016) IGHV1-69 polymorphism modulates anti-influenza antibody repertoires, correlates with IGHV utilization shifts and varies by ethnicity. Sci Rep., 6, 20842. doi: 10.1038/srep20842

3. Sangesland, M., Yousif, A.S., Ronsard, L., Kazer, S.W., Zhu, A.L., Gatter, G.J. et al. (2020) A Single Human V(H)-gene Allows for a Broad-Spectrum Antibody Response Targeting Bacterial Lipopolysaccharides in the Blood. Cell Rep., 32, 108065. doi: 10.1016/j.celrep.2020.108065

4. Benichou, J., Ben-Hamo, R., Louzoun, Y. and Efroni, S. (2012) Rep-Seq: uncovering the immunological repertoire through next-generation sequencing. Immunology, 135, 183–191. doi: 10.1111/j.1365-2567.2011.03527.x

5. Yaari, G. and Kleinstein, S.H. (2015) Practical guidelines for B-cell receptor repertoire sequencing analysis. Genome Med., 7, 121. doi: 10.1186/s13073-015-0243-2

6. Giudicelli, V., Chaume, D. and Lefranc, M.-P. (2005) IMGT/GENE-DB: a comprehensive database for human and mouse immunoglobulin and T cell receptor genes. Nucleic Acids Res., 33, D256–D261. doi: 10.1093/nar/gki010

7. Wang, Y., Jackson, K.J.L., Sewell, W.A. and Collins, A.M. (2008) Many human immunoglobulin heavy-chain IGHV gene polymorphisms have been reported in error. Immunol Cell Biol., 86, 111–115. doi: 10.1038/sj.icb.7100144

8. Rodriguez, O.L., Gibson, W.S., Parks, T., Emery, M., Powell, J., Strahl, M. et al. (2020) A Novel Framework for Characterizing Genomic Haplotype Diversity in the Human Immunoglobulin Heavy Chain Locus. Front Immunol., 11, 2136. doi: 10.3389/fimmu.2020.02136

9. Corcoran, M.M., Phad, G.E., Bernat, N.V., Stahl-Hennig, C., Sumida, N., Persson, M.A.A. et al. (2016) Production of individualized V gene databases reveals high levels of immunoglobulin genetic diversity. Nat Commun., 7, 13642. doi: 10.1038/ncomms13642

10. Ohlin, M., Scheepers, C., Corcoran, M., Lees, W.D., Busse, C.E., Bagnara, D. et al. (2019) Inferred Allelic Variants of Immunoglobulin Receptor Genes: A System for Their Evaluation, Documentation, and Naming. Front Immunol., 10, 435. doi: 10.3389/fimmu.2019.00435

11. Ralph, D.K. and Matsen, F.A.I.V. (2019) Per-sample immunoglobulin germline inference from B cell receptor deep sequencing data. PLoS Comput Biol., 15, e1007133. doi: 10.1371/journal.pcbi.1007133

12. Gadala-Maria, D., Gidoni, M., Marquez, S., Vander Heiden, J.A., Kos, J.T., Watson, C.T. et al. (2019) Identification of Subject-Specific Immunoglobulin Alleles From Expressed Repertoire Sequencing Data. Front Immunol., 10, 129. doi: 10.3389/fimmu.2019.00129

13. Lovett, P.S. and Rogers, E.J. (1996) Ribosome regulation by the nascent peptide. Microbiological Rev., 60, 366–385. doi: 10.1128/mr.60.2.366-385.1996

14. Zhu, Y., Yang, X., Wu, J., Tang, H., Wang, Q., Guan, J. et al. (2020) Antibody Upstream Sequence Diversity and Its Biological Implications Revealed by Repertoire Sequencing. bioRxiv, doi: https://doi.org/10.1101/2020.09.02.280396, 3 September 2020, pre-print: not peer-reviewed.

15. Wellensiek, B.P., Larsen, A.C., Flores, J., Jacobs, B.L. and Chaput, J.C. (2013) A leader sequence capable of enhancing RNA expression and protein synthesis in mammalian cells. Protein Sci., 22, 1392–1398. doi: 10.1002/pro.2325

16. Saintamand, A., Vincent-Fabert, C., Marquet, M., Ghazzaui, N., Magnone, V., Pinaud, E. et al. (2017) Eμ and 3’RR IgH enhancers show hierarchic unilateral dependence in mature B-cells. Scientific Rep., 7, 442. doi: 10.1038/s41598-017-00575-0

17. Alamyar, E., Duroux, P., Lefranc, M-P. and Giudicelli, V. (2012) IMGT(®) tools for the nucleotide analysis of immunoglobulin (IG) and T cell receptor (TR) V-(D)-J repertoires, polymorphisms, and IG mutations: IMGT/V-QUEST and IMGT/HighV-QUEST for NGS. Methods Mol Biol., 882, 569–604. doi: 10.1007/978-1-61779-842-9_32

18. Mikocziova, I., Gidoni, M., Lindeman, I., Peres, A., Snir, O., Yaari, G. and Sollid, L.M. (2020) Polymorphisms in human immunoglobulin heavy chain variable genes and their upstream regions. Nucleic Acids Res., 48, 5499–5510. doi: 10.1093/nar/gkaa310

19. Gidoni, M., Snir, O., Peres, A., Polak, P., Lindeman, I., Mikocziova, I. et al. (2019) Mosaic deletion patterns of the human antibody heavy chain gene locus shown by Bayesian haplotyping. Nat Commun., 10, 628. doi: 10.1038/s41467-019-08489-3

20. Kirik, U., Greiff, L., Levander, F. and Ohlin, M. (2017) Parallel antibody germline gene and haplotype analyses support the validity of immunoglobulin germline gene inference and discovery. Mol Immunol., 87, 12–22. doi: 10.1016/j.molimm.2017.03.012

21. Vander Heiden, J.A., Yaari, G., Uduman, M., Stern, J.N., O’Connor, K.C., Hafler, D.A. et al. (2014) pRESTO: a toolkit for processing high-throughput sequencing raw reads of lymphocyte receptor repertoires. Bioinformatics., 30, 1930–1932. doi: 10.1093/bioinformatics/btu138

22. Minoche, A.E., Dohm, J.C. and Himmelbauer, H. (2011) Evaluation of genomic high-throughput sequencing data generated on Illumina HiSeq and genome analyzer systems. Genome Biol., 12, R112. doi: 10.1186/gb-2011-12-11-r112

23. Ohlin, M. (2021) Poorly Expressed Alleles of Several Human Immunoglobulin Heavy Chain Variable Genes are Common in the Human Population. Front Immunol., 11, 603980. doi: 10.3389/fimmu.2020.603980

24. Sheng, Z., Schramm, C.A., Kong, R., N.C.S.P., Mullikin, J.C., Mascola, J.R. et al. (2017) Gene-Specific Substitution Profiles Describe the Types and Frequencies of Amino Acid Changes during Antibody Somatic Hypermutation. Front Immunol., 8, 537. doi: 10.3389/fimmu.2017.00537

25. Blankenberg, D., Gordon, A., Von Kuster, G., Coraor, N., Taylor, J., Nekrutenko, A. and Galaxy, T. (2010) Manipulation of FASTQ data with Galaxy. Bioinformatics., 26, 1783–1785. doi: 10.1093/bioinformatics/btq281

26. Lohse, M., Bolger, A.M., Nagel, A., Fernie, A.R., Lunn, J.E., Stitt, M. and Usadel, B. (2012) RobiNA: a user-friendly, integrated software solution for RNA-Seq-based transcriptomics. Nucleic Acids Res., 40, W622–627. doi: 10.1093/nar/gks540

27. Auton, A., Brooks, L.D., Durbin, R.M., Garrison, E.P., Kang, H.M., Korbel, J.O. et al. (2015) A global reference for human genetic variation. Nature., 526, 68–74. doi: 10.1038/nature15393

28. Yates, A.D., Achuthan, P., Akanni, W., Allen, J., Allen, J., Alvarez-Jarreta, J. et al. (2020) Ensembl 2020. Nucleic Acids Res., 48, D682–D688. doi: 10.1093/nar/gkz966

29. Watson, C.T., Matsen, F.A., Jackson, K.J.L., Bashir, A., Smith, M.L., Glanville, J. et al. (2017) Comment on “A Database of Human Immune Receptor Alleles Recovered from Population Sequencing Data”. J Immunol., 198, 3371–3373. doi: 10.4049/jimmunol.1700306

30. Omer, A., Shemesh, O., Peres, A., Polak, P., Shepherd, A.J., Watson, C.T. et al. (2020) VDJbase: an adaptive immune receptor genotype and haplotype database. Nucleic Acids Res., 48, D1051–D1056. doi: 10.1093/nar/gkz872

31. Simmons, M.P., Ochoterena, H. and Carr, T.G. (2001) Incorporation, Relative Homoplasy, and Effect of Gap Characters in Sequence-Based Phylogenetic Analyses. Syst Biol., 50, 454–462.

32. Ford, M., Haghshenas, E., Watson, C.T. and Sahinalp, S.C. (2020) Genotyping and Copy Number Analysis of Immunoglobulin Heavy Chain Variable Genes Using Long Reads. iScience., 23, 100883. doi: 10.1016/j.isci.2020.100883

33. Parks, T., Mirabel, M.M., Kado, J., Auckland, K., Nowak, J., Rautanen, A. et al. (2017) Association between a common immunoglobulin heavy chain allele and rheumatic heart disease risk in Oceania. Nat Commun., 8, 14946. doi: 10.1038/ncomms14946

34. Busse, C.E., Jackson, K.J.L., Watson, C.T. and Collins, A.M. (2019) A Proposed New Nomenclature for the Immunoglobulin Genes of Mus musculus. Front Immunol., 10, 2961. doi: 10.3389/fimmu.2019.02961

35. Allele. IMGT^®^, the international ImMunoGeneTics information system^®^, http://www.imgt.org/IMGTindex/allele.php [Accessed June 8, 2021].

36. Boyd, S.D., Gaëta, B.A., Jackson, K.J., Fire, A.Z., Marshall, E.L., Merker, J.D. et al. (2010) Individual Variation in the Germline Ig Gene Repertoire Inferred from Variable Region Gene Rearrangements. J Immunol., 184, 6986–6992. doi: 10.4049/jimmunol.1000445

37. Gadala-Maria, D., Yaari, G., Uduman, M. and Kleinstein, S.H. (2015) Automated analysis of high-throughput B-cell sequencing data reveals a high frequency of novel immunoglobulin V gene segment alleles. Proc Natl Acad Sci U S A., 112, E862–E870. doi: 10.1073/pnas.1417683112

38. Thörnqvist, L. and Ohlin, M. (2018) Critical steps for computational inference of the 3’-end of novel alleles of immunoglobulin heavy chain variable genes - illustrated by an allele of IGHV3-7. Mol Immunol., 103, 1–6. doi: 10.1016/j.molimm.2018.08.018

39. Gibbs, R.A., Rogers, J., Katze, M.G., Bumgarner, R., Weinstock, G.M., Mardis, E.R. et al. (2007) Evolutionary and Biomedical Insights from the Rhesus Macaque Genome. Science., 316, 222–234. doi: 10.1126/science.1139247

40. Huang, Y., Thörnqvist, L. and Ohlin, M. (2021) Computational inference, validation, and analysis of 5’UTR-leader sequences of alleles of immunoglobulin heavy chain variable genes. bioRxiv, 2021.06.10.447679; doi: 10.1101/2021.06.10.447679

