## Supplementary data 1 for "Computational inference, validation, and analysis of 5’UTR-leader sequences of alleles of immunoglobulin heavy chain variable genes"

**Supplementary Data 1.** Database of IGHV, IGHD, and IGHJ genes, as FASTA entries, used for inference of germline genes from transcriptome data. The sequences were obtained were obtained from the IMGT database (release 202011-3) (6).

>IGHV1-18*01

caggttcagctggtgcagtctggagctgaggtgaagaagcctggggcctcagtgaag

gtctcctgcaaggcttctggttacacctttaccagctatggtatcagc

tgggtgcgacaggcccctggacaagggcttgagtggatgggatggatcagcgcttac

aatggtaacacaaactatgcacagaagctccagggcagagtcaccatgaccaca

gacacatccacgagcacagcctacatggagctgaggagcctgagatctgacgacacggcc

gtgtattactgtgcgagaga

>IGHV1-18*02

caggttcagctggtgcagtctggagctgaggtgaagaagcctggggcctcagtgaag

gtctcctgcaaggcttctggttacacctttaccagctatggtatcagc

tgggtgcgacaggcccctggacaagggcttgagtggatgggatggatcagcgcttac

aatggtaacacaaactatgcacagaagctccagggcagagtcaccatgaccaca

gacacatccacgagcacagcctacatggagctgaggagcctaagatctgacgacacggcc

>IGHV1-18*03

caggttcagctggtgcagtctggagctgaggtgaagaagcctggggcctcagtgaag

gtctcctgcaaggcttctggttacacctttaccagctatggtatcagc

tgggtgcgacaggcccctggacaagggcttgagtggatgggatggatcagcgcttac

aatggtaacacaaactatgcacagaagctccagggcagagtcaccatgaccaca

gacacatccacgagcacagcctacatggagctgaggagcctgagatctgacgacatggcc

gtgtattactgtgcgagaga

>IGHV1-18*04

caggttcagctggtgcagtctggagctgaggtgaagaagcctggggcctcagtgaag

gtctcctgcaaggcttctggttacacctttaccagctacggtatcagc

tgggtgcgacaggcccctggacaagggcttgagtggatgggatggatcagcgcttac

aatggtaacacaaactatgcacagaagctccagggcagagtcaccatgaccaca

gacacatccacgagcacagcctacatggagctgaggagcctgagatctgacgacacggcc

gtgtattactgtgcgagaga

>IGHV1-2*01

caggtgcagctggtgcagtctggggctgaggtgaagaagcctggggcctcagtgaag

gtctcctgcaaggcttctggatacaccttcaccggctactatatgcac

tgggtgcgacaggcccctggacaagggcttgagtggatgggacggatcaaccctaac

agtggtggcacaaactatgcacagaagtttcagggcagggtcaccagtaccagg

gacacgtccatcagcacagcctacatggagctgagcaggctgagatctgacgacacggtc

gtgtattactgtgcgagaga

>IGHV1-2*02

caggtgcagctggtgcagtctggggctgaggtgaagaagcctggggcctcagtgaag

gtctcctgcaaggcttctggatacaccttcaccggctactatatgcac

tgggtgcgacaggcccctggacaagggcttgagtggatgggatggatcaaccctaac

agtggtggcacaaactatgcacagaagtttcagggcagggtcaccatgaccagg

gacacgtccatcagcacagcctacatggagctgagcaggctgagatctgacgacacggcc

gtgtattactgtgcgagaga

>IGHV1-2*03

caggtgcagctggtgcagtctggggctgaggtgaagaagcttggggcctcagtgaag

gtctcctgcaaggcttctggatacaccttcaccggctactatatgcac

tgggtgcnacaggcccctggacaagggcttgagtggatgggatggatcaaccctaac

agtggtggcacaaactatgcacagaagtttcagggcagggtcaccatgaccagg

gacacgtccatcagcacagcctacatggagctgagcaggctgagatctgacgacacggcc

gtgtattactgtgcgagaga

>IGHV1-2*04

caggtgcagctggtgcagtctggggctgaggtgaagaagcctggggcctcagtgaag

gtctcctgcaaggcttctggatacaccttcaccggctactatatgcac

tgggtgcgacaggcccctggacaagggcttgagtggatgggatggatcaaccctaac

agtggtggcacaaactatgcacagaagtttcagggctgggtcaccatgaccagg

gacacgtccatcagcacagcctacatggagctgagcaggctgagatctgacgacacggcc

gtgtattactgtgcgagaga

>IGHV1-2*05

caggtgcagctggtgcagtctggggctgaggtgaagaagcctggggcctcagtgaag

gtctcctgcaaggcttctggatacaccttcaccggctactatatgcac

tgggtgcgacaggcccctggacaagggcttgagtggatgggacggatcaaccctaac

agtggtggcacaaactatgcacagaagtttcagggcagggtcaccatgaccagg

gacacgtccatcagcacagcctacatggagctgagcaggctgagatctgacgacacggtc

gtgtattactgtgcgagaga

>IGHV1-2*06

caggtgcagctggtgcagtctggggctgaggtgaagaagcctggggcctcagtgaag

gtctcctgcaaggcttctggatacaccttcaccggctactatatgcac

tgggtgcgacaggcccctggacaagggcttgagtggatgggacggatcaaccctaac

agtggtggcacaaactatgcacagaagtttcagggcagggtcaccatgaccagg

gacacgtccatcagcacagcctacatggagctgagcaggctgagatctgacgacacggcc

gtgtattactgtgcgagaga

>IGHV1-24*01

caggtccagctggtacagtctggggctgaggtgaagaagcctggggcctcagtgaag

gtctcctgcaaggtttccggatacaccctcactgaattatccatgcac

tgggtgcgacaggctcctggaaaagggcttgagtggatgggaggttttgatcctgaa

gatggtgaaacaatctacgcacagaagttccagggcagagtcaccatgaccgag

gacacatctacagacacagcctacatggagctgagcagcctgagatctgaggacacggcc

gtgtattactgtgcaacaga

>IGHV1-3*01

caggtccagcttgtgcagtctggggctgaggtgaagaagcctggggcctcagtgaag

gtttcctgcaaggcttctggatacaccttcactagctatgctatgcat

tgggtgcgccaggcccccggacaaaggcttgagtggatgggatggatcaacgctggc

aatggtaacacaaaatattcacagaagttccagggcagagtcaccattaccagg

gacacatccgcgagcacagcctacatggagctgagcagcctgagatctgaagacacggct

gtgtattactgtgcgagaga

>IGHV1-3*02

caggttcagctggtgcagtctggggctgaggtgaagaagcctggggcctcagtgaag

gtttcctgcaaggcttctggatacaccttcactagctatgctatgcat

tgggtgcgccaggcccccggacaaaggcttgagtggatgggatggagcaacgctggc

aatggtaacacaaaatattcacaggagttccagggcagagtcaccattaccagg

gacacatccgcgagcacagcctacatggagctgagcagcctgagatctgaggacatggct

gtgtattactgtgcgagaga

>IGHV1-3*03

caggtccagctggtgcagtctggggctgaggtgaagaagcctggggcctcagtgaag

gtttcctgcaaggcttctggatacaccttcactagctatgctatgcat

tgggtgcgccaggcccccggacaaaggcttgagtggatgggatggatcaacgctggc

aatggtaacacaaaatattcacaggagttccagggcagagtcaccattaccagg

gacacatccgcgagcacagcctacatggagctgagcagcctgagatctgaggacatggct

gtgtattactgtgcgagaga

>IGHV1-3*04

caggtccagcttgtgcagtctggggctgaggtgaagaagcctggggcctcagtgaag

gtttcctgcaaggcttctggatacaccttcactagctatgctatgcat

tgggtgcgccaggcccccggacaaaggcttgagtggatgggatggatcaacactggc

aatggtaacacaaaatattcacagaagttccagggcagagtcaccattaccagg

gacacatccgcgagcacagcctacatggagctgagcagcctgagatctgaagacacggct

gtgtattactgtgcgagag

>IGHV1-38-4*01

caggtccagctggtgcagtcttgggctgaggtgaggaagtctggggcctcagtgaaa

gtctcctgtagtttttctgggtttaccatcaccagctacggtatacat

tgggtgcaacagtcccctggacaagggcttgagtggatgggatggatcaaccctggc

aatggtagcccaagctatgccaagaagtttcagggcagattcaccatgaccagg

gacatgtccacaaccacagcctacacagacctgagcagcctgacatctgaggacatggct

gtgtattactatgcaagaca

>IGHV1-45*01

cagatgcagctggtgcagtctggggctgaggtgaagaagactgggtcctcagtgaag

gtttcctgcaaggcttccggatacaccttcacctaccgctacctgcac

tgggtgcgacaggcccccggacaagcgcttgagtggatgggatggatcacacctttc

aatggtaacaccaactacgcacagaaattccaggacagagtcaccattactagg

gacaggtctatgagcacagcctacatggagctgagcagcctgagatctgaggacacagcc

atgtattactgtgcaagana

>IGHV1-45*02

cagatgcagctggtgcagtctggggctgaggtgaagaagactgggtcctcagtgaag

gtttcctgcaaggcttccggatacaccttcacctaccgctacctgcac

tgggtgcgacaggcccccggacaagcgcttgagtggatgggatggatcacacctttc

aatggtaacaccaactacgcacagaaattccaggacagagtcaccattaccagg

gacaggtctatgagcacagcctacatggagctgagcagcctgagatctgaggacacagcc

atgtattactgtgcaagata

>IGHV1-45*03

cagatgcagctggtgcagtctggggctgaggtgaagaagactgggtcctcagtgaag

gtttcctgcaaggcttccggatacaccttcacctaccgctacctgcac

tgggtgcgacaggcccccagacaagcgcttgagtggatgggatggatcacacctttc

aatggtaacaccaactacgcacagaaattccaggacagagtcaccattaccagg

gacaggtctatgagcacagcctacatggagctgagcagcctgagatctgaggacacagcc

atgtattactgtgcaagata

>IGHV1-46*01

caggtgcagctggtgcagtctggggctgaggtgaagaagcctggggcctcagtgaag

gtttcctgcaaggcatctggatacaccttcaccagctactatatgcac

tgggtgcgacaggcccctggacaagggcttgagtggatgggaataatcaaccctagt

ggtggtagcacaagctacgcacagaagttccagggcagagtcaccatgaccagg

gacacgtccacgagcacagtctacatggagctgagcagcctgagatctgaggacacggcc

gtgtattactgtgcgagaga

>IGHV1-46*02

caggtgcagctggtgcagtctggggctgaggtgaagaagcctggggcctcagtgaag

gtttcctgcaaggcatctggatacaccttcaacagctactatatgcac

tgggtgcgacaggcccctggacaagggcttgagtggatgggaataatcaaccctagt

ggtggtagcacaagctacgcacagaagttccagggcagagtcaccatgaccagg

gacacgtccacgagcacagtctacatggagctgagcagcctgagatctgaggacacggcc

gtgtattactgtgcgagaga

>IGHV1-46*03

caggtgcagctggtgcagtctggggctgaggtgaagaagcctggggcctcagtgaag

gtttcctgcaaggcatctggatacaccttcaccagctactatatgcac

tgggtgcgacaggcccctggacaagggcttgagtggatgggaataatcaaccctagt

ggtggtagcacaagctacgcacagaagttccagggcagagtcaccatgaccagg

gacacgtccacgagcacagtctacatggagctgagcagcctgagatctgaggacacggcc

gtgtattactgtgctagaga

>IGHV1-46*04

caggtgcagctggtgcagtctggggctgaggtgaagaagcctggggcctcagtgaag

gtttcctgcaaggcatctggatacaccttcaccagctactatatgcac

tgggtgcgacaggcccctggacaagggcttgagtggatgggaataatcaaccctagt

ggtggtagcacaagctacgcacagaagttgcagggcagagtcaccatgaccagg

gacacgtccacgagcacagtctacatggagctgagcagcctgagatctgaggacacggcc

gtgtattactgtgcgagaga

>IGHV1-58*01

caaatgcagctggtgcagtctgggcctgaggtgaagaagcctgggacctcagtgaag

gtctcctgcaaggcttctggattcacctttactagctctgctgtgcag

tgggtgcgacaggctcgtggacaacgccttgagtggataggatggatcgtcgttggc

agtggtaacacaaactacgcacagaagttccaggaaagagtcaccattaccagg

gacatgtccacaagcacagcctacatggagctgagcagcctgagatccgaggacacggcc

gtgtattactgtgcggcaga

>IGHV1-58*02

caaatgcagctggtgcagtctgggcctgaggtgaagaagcctgggacctcagtgaag

gtctcctgcaaggcttctggattcacctttactagctctgctatgcag

tgggtgcgacaggctcgtggacaacgccttgagtggataggatggatcgtcgttggc

agtggtaacacaaactacgcacagaagttccaggaaagagtcaccattaccagg

gacatgtccacaagcacagcctacatggagctgagcagcctgagatccgaggacacggcc

gtgtattactgtgcggcaga

>IGHV1-58*03

caaatgcagctggtgcagtctgggcctgaagtgaagaagcctgggacctcagtgaag

gtctcctgcaaggcttctggattcacctttactagctctgctgtgcag

tgggtgcgacaggctcgtggacaacgccttgagtggataggatggatcgtcgttggc

agtggtaacacaaactacgcacagaagttccaggaaagagtcaccattaccagg

gacatgtccacaagcacagcctacatggagctgagcagcctgagatccgaggacacggcc

gtgtattactgtgcggcaga

>IGHV1-68*01

caggtgcagctggggcagtctgaggctgaggtaaagaagcctggggcctcagtgaag

gtctcctgcaaggcttccggatacaccttcacttgctgctccttgcac

tggttgcaacaggcccctggacaagggcttgaaaggatgagatggatcacactttac

aatggtaacaccaactatgcaaagaagttccagggcagagtcaccattaccagg

gacatgtccctgaggacagcctacatagagctgagcagcctgagatctgaggactcggct

gtgtattactgggcaagata

>IGHV1-68*02

caggtgcagctggggcagtctgaggctgaggtgaagaagcctggggcctcagtgaag

gtctcctgcaaggcttccggatacaccttcacctactgctccttgcac

tggttgcaacaggcccctggacaagggcttgaaaggatgagatggatcacactttac

aatggtaacatcaactatgcaaagaagttccagagcagagtcaccattaccagg

gacatgtccctgaggacagcctacatagagctgagcagcctgagatctgaggactcggct

gtgtattactgggcaagata

>IGHV1-69*01

caggtgcagctggtgcagtctggggctgaggtgaagaagcctgggtcctcggtgaag

gtctcctgcaaggcttctggaggcaccttcagcagctatgctatcagc

tgggtgcgacaggcccctggacaagggcttgagtggatgggagggatcatccctatc

tttggtacagcaaactacgcacagaagttccagggcagagtcacgattaccgcg

gacgaatccacgagcacagcctacatggagctgagcagcctgagatctgaggacacggcc

gtgtattactgtgcgagaga

>IGHV1-69*02

caggtccagctggtgcaatctggggctgaggtgaagaagcctgggtcctcggtgaag

gtctcctgcaaggcttctggaggcaccttcagcagctatactatcagc

tgggtgcgacaggcccctggacaagggcttgagtggatgggaaggatcatccctatc

cttggtatagcaaactacgcacagaagttccagggcagagtcacgattaccgcg

gacaaatccacgagcacagcctacatggagctgagcagcctgagatctgaggacacggcc

gtgtattactgtgcgaga

>IGHV1-69*03

caggtgcagctggtgcagtctggggctgaggtgaagaagcctgggtcctcggtgaag

gtctcctgcaaggcttctggaggcaccttcagcagctatgctatcagc

tgggtgcgacaggcccctggacaagggcttgagtggatgggagggatcatccctatc

tttggtacagcaaactacgcacagaagttccagggcagagtcacgattaccgcg

gacgaatccacgagcacagcctacatggagctgagcagcctgagatctgatgacacggc

>IGHV1-69*04

caggtccagctggtgcagtctggggctgaggtgaagaagcctgggtcctcggtgaag

gtctcctgcaaggcttctggaggcaccttcagcagctatgctatcagc

tgggtgcgacaggcccctggacaagggcttgagtggatgggaaggatcatccctatc

cttggtatagcaaactacgcacagaagttccagggcagagtcacgattaccgcg

gacaaatccacgagcacagcctacatggagctgagcagcctgagatctgaggacacggcc

gtgtattactgtgcgagaga

>IGHV1-69*05

caggtccagctggtgcagtctggggctgaggtgaagaagcctgggtcctcggtgaag

gtctcctgcaaggcttctggaggcaccttcagcagctatgctatcagc

tgggtgcgacaggcccctggacaagggcttgagtggatgggagggatcatccctatc

tttggtacagcaaactacgcacagaagttccagggcagagtcacgattaccacg

gacgaatccacgagcacagcctacatggagctgagcagcctgagatctgaggacacggcc

gtgtattactgtgcgaga

>IGHV1-69*06

caggtgcagctggtgcagtctggggctgaggtgaagaagcctgggtcctcggtgaag

gtctcctgcaaggcttctggaggcaccttcagcagctatgctatcagc

tgggtgcgacaggcccctggacaagggcttgagtggatgggagggatcatccctatc

tttggtacagcaaactacgcacagaagttccagggcagagtcacgattaccgcg

gacaaatccacgagcacagcctacatggagctgagcagcctgagatctgaggacacggcc

gtgtattactgtgcgagaga

>IGHV1-69*07

agaagcctgggtcctcggtgaag

gtctcctgcaaggcttctggaggcaccttcagcagctatgctatcagc

tgggtgcgacaggcccctggacaagggcttgagtggatgggaaggatcatccctatc

tttggtacagcaaactacgcacagaagttccagggcagagtcacgattaccgcg

gacgaatccacgagcacagcctacatggagctgagcagcctgagatctgag

>IGHV1-69*08

caggtccagctggtgcaatctggggctgaggtgaagaagcctgggtcctcggtgaag

gtctcctgcaaggcttctggaggcaccttcagcagctatactatcagc

tgggtgcgacaggcccctggacaagggcttgagtggatgggaaggatcatccctatc

cttggtacagcaaactacgcacagaagttccagggcagagtcacgattaccgcg

gacaaatccacgagcacagcctacatggagctgagcagcctgagatctgaggacacggcc

gtgtattactgtgcgagaga

>IGHV1-69*09

caggtgcagctggtgcagtctggggctgaggtgaagaagcctgggtcctcggtgaag

gtctcctgcaaggcttctggaggcaccttcagcagctatgctatcagc

tgggtgcgacaggcccctggacaagggcttgagtggatgggaaggatcatccctatc

cttggtatagcaaactacgcacagaagttccagggcagagtcacgattaccgcg

gacaaatccacgagcacagcctacatggagctgagcagcctgagatctgaggacacggcc

gtgtattactgtgcgagaga

>IGHV1-69*10

caggtccagctggtgcagtctggggctgaggtgaagaagcctgggtcctcagtgaag

gtctcctgcaaggcttctggaggcaccttcagcagctatgctatcagc

tgggtgcgacaggcccctggacaagggcttgagtggatgggagggatcatccctatc

cttggtatagcaaactacgcacagaagttccagggcagagtcacgattaccgcg

gacaaatccacgagcacagcctacatggagctgagcagcctgagatctgaggacacggcc

gtgtattactgtgcgagaga

>IGHV1-69*11

caggtccagctggtgcagtctggggctgaggtgaagaagcctgggtcctcggtgaag

gtctcctgcaaggcttctggaggcaccttcagcagctatgctatcagc

tgggtgcgacaggcccctggacaagggcttgagtggatgggaaggatcatccctatc

cttggtacagcaaactacgcacagaagttccagggcagagtcacgattaccgcg

gacgaatccacgagcacagcctacatggagctgagcagcctgagatctgaggacacggcc

gtgtattactgtgcgagaga

>IGHV1-69*12

caggtccagctggtgcagtctggggctgaggtgaagaagcctgggtcctcggtgaag

gtctcctgcaaggcttctggaggcaccttcagcagctatgctatcagc

tgggtgcgacaggcccctggacaagggcttgagtggatgggagggatcatccctatc

tttggtacagcaaactacgcacagaagttccagggcagagtcacgattaccgcg

gacgaatccacgagcacagcctacatggagctgagcagcctgagatctgaggacacggcc

gtgtattactgtgcgagaga

>IGHV1-69*13

caggtccagctggtgcagtctggggctgaggtgaagaagcctgggtcctcagtgaag

gtctcctgcaaggcttctggaggcaccttcagcagctatgctatcagc

tgggtgcgacaggcccctggacaagggcttgagtggatgggagggatcatccctatc

tttggtacagcaaactacgcacagaagttccagggcagagtcacgattaccgcg

gacgaatccacgagcacagcctacatggagctgagcagcctgagatctgaggacacggcc

gtgtattactgtgcgagaga

>IGHV1-69*14

caggtccagctggtgcagtctggggctgaggtgaagaagcctgggtcctcggtgaag

gtctcctgcaaggcttctggaggcaccttcagcagctatgctatcagc

tgggtgcgacaggcccctggacaagggcttgagtggatgggagggatcatccctatc

tttggtacagcaaactacgcacagaagttccagggcagagtcacgattaccgcg

gacaaatccacgagcacagcctacatggagctgagcagcctgagatctgaggacacggcc

gtgtattactgtgcgagaga

>IGHV1-69*15

caggtccagctggtgcagtctggggctgaggtgaagaagcctgggtcctcggtgaag

gtctcctgcaaggcttctggaggcaccttcagcagctatgctatcagc

tgggtgcgacaggcccctggacaagggcttgagtggatgggaaggatcatccctatc

tttggtacagcaaactacgcacagaagttccagggcagagtcacgattaccgcg

gacgaatccacgagcacagcctacatggagctgagcagcctgagatctgaggacacggcc

gtgtattactgtgcgagaga

>IGHV1-69*16

caggtccagctggtgcagtctggggctgaggtgaagaagcctgggtcctcggtgaag

gtctcctgcaaggcttctggaggcaccttcagcagctatactatcagc

tgggtgcgacaggcccctggacaagggcttgagtggatgggagggatcatccctatc

cttggtacagcaaactacgcacagaagttccagggcagagtcacgattaccacg

gacgaatccacgagcacagcctacatggagctgagcagcctgagatctgaggacacggcc

gtgtattactgtgcgagaga

>IGHV1-69*17

caggtgcagctggtgcagtctggggctgaggtgaagaagcctgggtcctcggtgaag

gtctcctgcaaggcttctggaggcaccttcagcagctatgctatcagc

tgggtgcgacaggcccctggacaagggcttgagtggatgggagggatcatccctatc

tttggtatagcaaactacgcacagaagttccagggcagagtcacgattaccgcg

gacaaatccacgagcacagcctacatggagctgagcagcctgagatctgaggacacggcc

gtgtattactgtgcgagaga

>IGHV1-69*18

caggtgcagctggtgcagtctggggctgaggtgaagaagcctgggtcctcggtgaag

gtctcctgcaaggcttctggaggcaccttcagcagctatgctatcagc

tgggtgcgacaggcccctggacaagggcttgagtggatgggaaggatcatccctatc

tttggtacagcaaactacgcacagaagttccagggcagagtcacgattaccgcg

gacgaatccacgagcacagcctacatggagctgagcagcctgagatctgaggacacggcc

gtgtattactgtgcgagag

>IGHV1-69-2*01

gaggtccagctggtacagtctggggctgaggtgaagaagcctggggctacagtgaaa

atctcctgcaaggtttctggatacaccttcaccgactactacatgcac

tgggtgcaacaggcccctggaaaagggcttgagtggatgggacttgttgatcctgaa

gatggtgaaacaatatacgcagagaagttccagggcagagtcaccataaccgcg

gacacgtctacagacacagcctacatggagctgagcagcctgagatctgaggacacggcc

gtgtattactgtgcaacaga

>IGHV1-69-2*02

agaagcctggggctacagtgaaa

atctcctgcaaggtttctggatacaccttcaccgactactacatgcac

tgggtgcaacaggcccctggaaaagggcttgagtggatgggacttgttgatcctgaa

gatggtgaaacaatatatgcagagaagttccagggcagagtcaccataaccgcg

gacacgtctacagacacagcctacatggagctgagcagcctgagatctgag

>IGHV1-69D*01

caggtgcagctggtgcagtctggggctgaggtgaagaagcctgggtcctcggtgaag

gtctcctgcaaggcttctggaggcaccttcagcagctatgctatcagc

tgggtgcgacaggcccctggacaagggcttgagtggatgggagggatcatccctatc

tttggtacagcaaactacgcacagaagttccagggcagagtcacgattaccgcg

gacgaatccacgagcacagcctacatggagctgagcagcctgagatctgaggacacggcc

gtgtattactgtgcgagaga

>IGHV1-8*01

caggtgcagctggtgcagtctggggctgaggtgaagaagcctggggcctcagtgaag

gtctcctgcaaggcttctggatacaccttcaccagttatgatatcaac

tgggtgcgacaggccactggacaagggcttgagtggatgggatggatgaaccctaac

agtggtaacacaggctatgcacagaagttccagggcagagtcaccatgaccagg

aacacctccataagcacagcctacatggagctgagcagcctgagatctgaggacacggcc

gtgtattactgtgcgagagg

>IGHV1-8*02

caggtgcagctggtgcagtctggggctgaggtgaagaagcctggggcctcagtgaag

gtctcctgcaaggcttctggatacaccttcaccagctatgatatcaac

tgggtgcgacaggccactggacaagggcttgagtggatgggatggatgaaccctaac

agtggtaacacaggctatgcacagaagttccagggcagagtcaccatgaccagg

aacacctccataagcacagcctacatggagctgagcagcctgagatctgaggacacggcc

gtgtattactgtgcgagagg

>IGHV1-8*03

caggtgcagctggtgcagtctggggctgaggtgaagaagcctggggcctcagtgaag

gtctcctgcaaggcttctggatacaccttcaccagctatgatatcaac

tgggtgcgacaggccactggacaagggcttgagtggatgggatggatgaaccctaac

agtggtaacacaggctatgcacagaagttccagggcagagtcaccattaccagg

aacacctccataagcacagcctacatggagctgagcagcctgagatctgaggacacggcc

gtgtattactgtgcgagagg

>IGHV1-NL1*01

caggttcagctgttgcagcctggggtccaggtgaagaagcctgggtcctcagtgaag

gtctcctgctaggcttccagatacaccttcaccaaatactttacacgg

tgggtgtgacaaagccctggacaagggcatnagtggatgggatgaatcaacccttac

aacgataacacacactacgcacagacgttctggggcagagtcaccattaccagt

gacaggtccatgagcacagcctacatggagctgagcngcctgagatccgaagacatggtc

gtgtattactgtgtgagaga

>IGHV1/OR15-1*01

caggtgcagctggtgcagtctggggctgaggtgaagaagcctggggcctcagtgaag

gtctcctgcaaggcttctggatacatcttcaccgactactatatgcac

tgggtgcgacaggcccctggacaagagcttgggtggatgggacggatcaaccctaac

agtggtggcacaaactatgcacagaagtttcagggcagagtcaccatgaccagg

gacacgtccatcagcacagcctacacggagctgagcagcctgagatctgaggacacggcc

acgtattactgtgcgaga

>IGHV1/OR15-1*02

caggtgcagctggtgcagtctggggctgaggtgaagaagcctggggcctcagtgaag

gtctcctgcaaggcttctggatacatcttcaccgactactatatgcac

tgggtgcgacaggcccctggacaagagcttgggtggatgggacggatcaaccctaac

agtggtggcacaaactatgcacagaagtttcagggcagagtcaccatgaccagg

gacacgtccatcagcacagcctgcacggagctgagcagcctgagatctgaggacacggcc

acgtattactgtgcgagaga

>IGHV1/OR15-1*03

caggtgcagctggtgcagtctggggctgaggtgaagaagcctggggcctcagtgaag

gtctcctgcaaggcttctggatacatcttcaccgactactatatgcac

tgggtgcgacaggcccctggacaagagcttgggtggatgggacggatcaaccctaac

agtggtggcacaaactatgcacagaagtttcagggcagagtcaccatgaccagg

gacacgtccatcagcacagcctacacggagctgagcagcctgagatctgaggacacagcc

acgtattactgtgcgagaga

>IGHV1/OR15-1*04

caggtgcagctggtgcagtctggggctgaggtgaagaagcctggggcctcagtgaag

gtctcctgcaaggcttctggatacatcttcaccgactactatatgcac

tgggtgcgacaggcccctggacaagagcttgggtggatgggacggatcaaccctaac

agtggtggcacaaactatgcacagaagtttcagggcagagtcaccatgaccagg

gacacgtccatcagcacagcctacatggagctgagcagcctgagatctgaggacacggcc

acgtattactgtgcgagaga

>IGHV1/OR15-2*01

caggtgcagctggtgcagtctggagctgaggtgaagaagcctagagcctcagtgaag

gtctcctgcaaggcttctggttacacctttaccagctactatatgcac

tgggtgtgacaggcccctgaacaagggcttgagtggatgggatggatcaacacttac

aatggtaacacaaactacccacagaagctccagggcagagtcaccatgaccaga

gacacatccacgagcacagcctacatggagctgagcaggctgagatctgacgacatggcc

gtgtattactgtgcgagaga

>IGHV1/OR15-2*02

caggtgcagctggtgcagtctggagctgaggtgaagaagcctggagcctcagtgaag

gtctcctgcaaggcttctggttacacctttaccagctactatatgcac

tgggtgtgacaggcccctgaacaagggcttgagtggatgggatggatcaacacttac

aatggtaacacaaactacccacagaagctccagggcagagtcaccatgaccaga

gacacatccacgagcacagcctacatggagctgagcagcctgagatctgacgacatggcc

gtgtattactgtgcgagaga

>IGHV1/OR15-2*03

caggtgcagctggtgcagtctggagctgaggtgaagaagcctagagcctcagtgaag

gtctcctgcaaggcttctggttacacctttaccagctactatatgcac

tgggtgtgacaggcccctgaacaagggcttgagtggatgggatggatcaacacttac

aatggtaacacaaactacccacagaagctccagggcagagtcaccatgaccaga

gacacatccacgagcacagcctacatggagctgagcagcctgagatctgacgacatggcc

gtgtattactgtgcgagaga

>IGHV1/OR15-3*01

caggtccaactggtgtagtctggagctgaggtgaagaagcctggggcctcagtgaag

gtctcctgcaaggcttctggatacaccttcaccgactactttatgaac

tggatgcgccaggcccctggacaaaggcttgagtggatgggatggatcaacgctggc

aatggtaacacaaaatattcacagaagctccagggcagagtcaccattaccagg

gacacatcttcgagcacagcctacatgcagctgagcagcctgagatctgaggacacggcc

gtgtattactgtgcgaga

>IGHV1/OR15-3*02

caggtccaactggtgtagtctggagctgaggtgaagaagcctggggcctcagtgaag

gtctcctgcaaggcttctggatacaccttcaccgactactttatgaac

tggatgcgccaggcccctggacaaaggcttgagtggatgggatggatcaacgctggc

aatggtaacacaaaatattcacagaagctccagggcagagtcaccattaccagg

gacacatctgcgagcacagcctacatgcagctgagcagcctgagatctgaggacacggcc

gtgtattactgtgcgagaga

>IGHV1/OR15-3*03

caggtccaactggtgtagtctggagctgaggtgaagaagcctggggcctcagtgaag

gtctcctgcaaggcttctggatacaccttcaccagctactatatgaac

tggatgcgccaggcccctggacaaggcttcgagtggatgggatggatcaacgctggc

aatggtaacacaaagtattcacagaagctccagggcagagtcaccattaccagg

gacacatctgcgagcacagcctacatgcagctgagcagcctgagatctgaggacacggcc

gtgtattactgtgcgagaga

>IGHV1/OR15-4*01

caggaccagttggtgcagtctggggctgaggtgaagaagcctctgtcctcagtgaag

gtctccttcaaggcttctggatacaccttcaccaacaactttatgcac

tgggtgtgacaggcccctggacaaggacttgagtggatgggatggatcaatgctggc

aatggtaacacaacatatgcacagaagttccagggcagagtcaccataaccagg

gacacgtccatgagcacagcctacacggagctgagcagcctgagatctgaggacacggcc

gtgtattactgtgcgaga

>IGHV1/OR15-5*01

agaagcctggggcctcagtgaag

gtctcctgcaaggcttctggatacaccttcaccagctactgtatgcac

tgggtgcaccaggtccatgcacaagggcttgagtggatgggattggtgtgccctagt

gatggcagcacaagctatgcacagaagttccaggccagagtcaccataaccagg

gacacatccatgagcacagcctacatggagctaagcagtctgagatctgaggacacggcc

atgtattactgtgtgaga

>IGHV1/OR15-5*02

caggtacagctggtgcagtctggggctgaggtgaagaagcctggggcctcagtgaag

gtctcctgcaaggcttctggatacaccttcaccaactactgtatgcac

tgggtgcgccaggtccatgcacaagggcttgagtggatgggattggtgtgccctagt

gatggcagcacaagctatgcacaaaagttccaggccagagtcaccataaccagg

gacacatccatgagcacagcctacatggagctaagcagtctgagatctgaggacacggcc

atgtattactgtgtgaga

>IGHV1/OR15-9*01

caggtacagctgatgcagtctggggctgaggtgaagaagcctggggcctcagtgagg

atctcctgcaaggcttctggatacaccttcaccagctactgtatgcac

tgggtgtgccaggcccatgcacaagggcttgagtggatgggattggtgtgccctagt

gatggcagcacaagctatgcacagaagttccagggcagagtcaccataaccagg

gacacatccatgggcacagcctacatggagctaagcagcctgagatctgaggacacggcc

atgtattactgtgtgagaga

>IGHV1/OR21-1*01

caggtacagctggtgcagtctggggctgaggtgaagaagcctggggcctcagtgaag

gtctcctgcaaggcttctggatacaccatcaccagctactgtatgcac

tgggtgcaccaggtccatgcacaagggcttgagtggatgggattggtgtgccctagt

gatggcagcacaagctatgcacagaagttccaggccagagtcaccataaccagg

gacacatccatgagcacagcctacatggagctaagcagtctgagatctgaggacacggcc

atgtattactgtgtgagaga

>IGHV2-10*01

caggtcaccttgaaggagtctggtcctgcactggtgaaacccacacagaccctcatg

ctgacctgcaccttctctgggttctcactcagcacttctggaatgggtgtgggt

tagatctgtcagccctcagcaaaggccctggagtggcttgcacacatttattagaat

gataataaatactacagcccatctctgaagagtaggctcattatctccaag

gacacctccaagaatgaagtggttctaacagtgatcaacatggacattgtggacacagcc

acacattactgtgcaaggagac

>IGHV2-26*01

caggtcaccttgaaggagtctggtcctgtgctggtgaaacccacagagaccctcacg

ctgacctgcaccgtctctgggttctcactcagcaatgctagaatgggtgtgagc

tggatccgtcagcccccagggaaggccctggagtggcttgcacacattttttcgaat

gacgaaaaatcctacagcacatctctgaagagcaggctcaccatctccaag

gacacctccaaaagccaggtggtccttaccatgaccaacatggaccctgtggacacagcc

acatattactgtgcacggatac

>IGHV2-26*02

caggtcaccttgaaggagtctggtcctgtgctggtgaaacccacagagaccctcacg

ctgacctgcaccgtctctgggttctcactcagcaatgctagaatgggtgtgagc

tggatccgtcagcccccagggaaggccctggagtggcttgcacacattttttcgaat

gacgaaaaatcctacagcacatctctgaagagcaggctcaccatctccaag

gacacctccaaaagccaggtggtccttaccatgaccaatatggaccctgtggacacagcc

acatattactgtgcacggatac

>IGHV2-26*03

caggtcaccttgaaggagtctggtcctgtgctggtgaaacccacagagaccctcacg

ctgacctgcaccatctctgggttctcactcagcaatgctagaatgggtgtgagc

tggatccgtcagcccccagggaaggccctggagtggcttgcacacattttttcgaat

gacgaaaaatcctacagcacatctctgaagagcaggctcaccatctccaag

gacacctccaaaagccaggtggtccttaccatgaccaacatggaccctgtggacacagcc

acatattactgtgcacggatac

>IGHV2-5*01

cagatcaccttgaaggagtctggtcctacgctggtgaaacccacacagaccctcacg

ctgacctgcaccttctctgggttctcactcagcactagtggagtgggtgtgggc

tggatccgtcagcccccaggaaaggccctggagtggcttgcactcatttattggaat

gatgataagcgctacagcccatctctgaagagcaggctcaccatcaccaag

gacacctccaaaaaccaggtggtccttacaatgaccaacatggaccctgtggacacagcc

acatattactgtgcacacagac

>IGHV2-5*02

cagatcaccttgaaggagtctggtcctacgctggtgaaacccacacagaccctcacg

ctgacctgcaccttctctgggttctcactcagcactagtggagtgggtgtgggc

tggatccgtcagcccccaggaaaggccctggagtggcttgcactcatttattgggat

gatgataagcgctacagcccatctctgaagagcaggctcaccatcaccaag

gacacctccaaaaaccaggtggtccttacaatgaccaacatggaccctgtggacacagcc

acatattactgtgcacacagac

>IGHV2-5*03

gctggtgaaacccacacagaccctcacg

ctgacctgcaccttctctgggttctcactcagcactagtggagtgggtgtgggc

tggatccgtcagcccccaggaaaggccctggagtggcttgcactcatttattgggat

gatgataagcgctacagcccatctctgaagagcaggctcaccattaccaag

gacacctccaaaaaccaggt

>IGHV2-5*04

cagatcaccttgaaggagtctggtcctacgctggtgaaacccacacagaccctcacg

ctgacctgcaccttctctgggttctcactcagcactagtggagtgggtgtgggc

tggatccgtcagcccccaggaaaggccctggagtggcttgcactcatttattggaat

gatgataagcgctacagcccatctctgaagagcaggctcaccatcaccaag

gacacctccaaaaaccaggtggtccttacaatgaccaacatggaccctgtggacacaggc

acatattactgtgtac

>IGHV2-5*05

cagatcaccttgaaggagtctggtcctacgctggtgaaacccacacagaccctcacg

ctgacctgcaccttctctgggttctcactcagcactagtggagtgggtgtgggc

tggatccgtcagcccccaggaaaggccctggagtggcttgcactcatttattgggat

gatgataagcgctacggcccatctctgaagagcaggctcaccatcaccaag

gacacctccaaaaaccaggtggtccttacaatgaccaacatggaccctgtggacacagcc

acatattactgtgcacacagac

>IGHV2-5*06

cagatcaccttgaaggagtctggtcctacgctggtaaaacccacacagaccctcacg

ctgacctgcaccttctctgggttctcactcagcactagtggagtgggtgtgggc

tggatccgtcagcccccaggaaaggccctggagtggcttgcactcatttattgggat

gatgataagcgctacggcccatctctgaagagcaggctcaccatcaccaag

gacacctccaaaaaccaggtggtccttacaatgaccaacatggaccctgtggacacagcc

acatattactgtgcacacaga

>IGHV2-5*08

caggtcaccttgaaggagtctggtcctgcgctggtgaaacccacacagaccctcaca

ctgacctgcaccttctctgggttctcactcagcactagtggaatgcgtgtgagc

tggatccgtcagcccccaggaaaggccctggagtggcttgcactcatttattgggat

gatgataagcgctacagcccatctctgaagagcaggctcaccatcaccaag

gacacctccaaaaaccaggtggtccttacaatgaccaacatggaccctgtggacacagcc

acatattactgtgcacacagac

>IGHV2-5*09

caggtcaccttgaaggagtctggtcctacgctggtgaaacccacacagaccctcacg

ctgacctgcaccttctctgggttctcactcagcactagtggagtgggtgtgggc

tggatccgtcagcccccaggaaaggccctggagtggcttgcactcatttattgggat

gatgataagcgctacggcccatctctgaagagcaggctcaccatcaccaag

gacacctccaaaaaccaggtggtccttacaatgaccaacatggaccctgtggacacagcc

acatattactgtgcacacagac

>IGHV2-70*01

caggtcaccttgagggagtctggtcctgcgctggtgaaacccacacagaccctcaca

ctgacctgcaccttctctgggttctcactcagcactagtggaatgtgtgtgagc

tggatccgtcagcccccagggaaggccctggagtggcttgcactcattgattgggat

gatgataaatactacagcacatctctgaagaccaggctcaccatctccaag

gacacctccaaaaaccaggtggtccttacaatgaccaacatggaccctgtggacacagcc

acgtattactgtgcacggatac

>IGHV2-70*02

caggtcaccttgagggagtctggtcctgcgctggtgaaacccacacagaccctcaca

ctgacctgcaccttctctgggttctcactcagcactagtggaatgtgtgtgagc

tggatccgtcagcccccagggaaggccctggagtggcttgcactcattgattgggat

gatgataaatactacagcacatctctgaagaccaggctcaccatctccaag

gacacctccaaaaaccaggtggtccttacaatgaccaacatggaccctgtggacacggcc

gtgtattactg

>IGHV2-70*03

caggtcaccttgaaggagtctggtcctgcgctggtgaaacccacacagaccctcaca

ctgacctgcaccttctctgggttctcactcagcactagtggaatgcgtgtgagc

tggatccgtcagcccccagggaaggccctggagtggcttgcacgcattgattgggat

gatgataaattctacagcacatctctgaagaccaggctcaccatctccaag

gacacctccaaaaaccaggtggtccttacaatgaccaacatggaccctgtggacacggcc

gtgtattactg

>IGHV2-70*04

caggtcaccttgaaggagtctggtcctgcgctggtgaaacccacacagaccctcaca

ctgacctgcaccttctctgggttctcactcagcactagtggaatgcgtgtgagc

tggatccgtcagcccccagggaaggccctggagtggcttgcacgcattgattgggat

gatgataaattctacagcacatctctgaagaccaggctcaccatctccaag

gacacctccaaaaaccaggtggtccttacaatgaccaacatggaccctgtggacacagcc

acgtattactgtgcacggatac

>IGHV2-70*05

tgcgctggtgaaacccacacagaccctcaca

ctgacctgcaccttctctgggttctcactcagcactagtggaatgcgtgcgagc

tggatccgtcagcccccagggaaggccctggagtggcttgcacgcattgattgggat

gatgataaattctacagcacatctctgaagaccaggctcaccatctccaag

gacacctccaaaaaccaggtggtccttacaatgaccaacatgga

>IGHV2-70*06

caggtcaccttgaaggagtctggtcctgcgctggtgaaacccacacagaccctcaca

ctgacctgcaccttctctgggttctcactcagcactagtggaatgcgtgtgagc

tggatccgtcagcccccagggaaggccctggagtggcttgcacgcattgattgggat

gatgataaattctacagcacatccctgaagaccaggctcaccatctccaag

gacacctccaaaaaccaggtggtccttacaatgaccaacatggaccctgtggacacggcc

gtgtattactg

>IGHV2-70*07

caggtcaccttgagggagtctggtcctgcgctggtgaaacccacacagaccctcaca

ctgacctgcaccttctctgggttctcactcagcactagtggaatgtgtgtgagc

tggatccgtcagcccccggggaaggccctggagtggcttgcactcattgattgggat

gatgataaatactacagcacatctctgaagaccaggctcaccatctccaag

gacacctccaaaaaccaggtggtccttacaatgaccaacatggaccctgtggacacggcc

gtgtattactg

>IGHV2-70*08

caggtcaccttgagggagtctggtcctgcgctggtgaaacccacacagaccctcaca

ctgacctgcgccttctctgggttctcactcagcactagtggaatgtgtgtgagc

tggatccgtcagcccccagggaaggccctggagtggcttgcacgcattgattgggat

gatgataaatactacagcacatctctgaagaccaggctcaccatctccaag

gacacctccaaaaaccaggtggtccttacaatgaccaacatggaccctgtggacacggcc

gtgtattactg

>IGHV2-70*09

cagatcaccttgaaggagtctggtcctacgctggtgaaacccacacagaccctcacg

ctgacccgcaccttctctgggttctcactcagcactagtggaatgtgtgtgagc

tggatccgtcagcccccagggaaggccctggagtggcttgcactcattgattgggat

gatgataaatactacagcacatctctgaacaccaggctcaccatctccaag

gacacctccaaaaaccaggtggtccttacaatgaccaacatggaccctgtggacacaggc

acatattactgtgtacgg

>IGHV2-70*10

caggtcaccttgaaggagtctggtcctgcgctggtgaaacccacacagaccctcaca

ctgacctgcaccttctctgggttctcactcagcactagtggaatgcgtgtgagc

tggatccgtcagcccccagggaaggccctggagtggattgcacgcattgattgggat

gatgataaatactacagcacatctctgaagaccaggctcaccatctccaag

gacacctccaaaaaccaggtggtccttacaatgaccaacatggaccctgtggacacagcc

acgtattactgtgcacggatac

>IGHV2-70*11

cgggtcaccttgagggagtctggtcctgcgctggtgaaacccacacagaccctcaca

ctgacctgcaccttctctgggttctcactcagcactagtggaatgtgtgtgagc

tggatccgtcagcccccagggaaggccctggagtggcttgcacgcattgattgggat

gatgataaatactacagcacatctctgaagaccaggctcaccatctccaag

gacacctccaaaaaccaggtggtccttacaatgaccaacatggaccctgtggacacagcc

acgtattactgtgcacggatac

>IGHV2-70*12

cagatcaccttgaaggagtctggtcctacgctggtgaaacccacacagaccctcacg

ctgacctgcaccttctctgggttctcactcagcactagtggaatgtgtgtgagc

tggatccgtcagcccccagggaaggccctggagtggcttgcactcattgattgggat

gatgataaatactacagcacatctctgaagaccaggctcaccatctccaag

gacacctccaaaaaccaggtggtccttacaatgaccaacatggaccctgtggacacagcc

acatattactgtgcacacagac

>IGHV2-70*13

caggtcaccttgagggagtctggtcctgcgctggtgaaacccacacagaccctcaca

ctgacctgcaccttctctgggttctcactcagcactagtggaatgtgtgtgagc

tggatccgtcagcccccagggaaggccctggagtggcttgcactcattgattgggat

gatgataaatactacagcacatctctgaagaccaggctcaccatctccaag

gacacctccaaaaaccaggtggtccttacaatgaccaacatggaccctgtggacacagcc

acgtattattgtgcacggatac

>IGHV2-70*15

caggtcaccttgagggagtctggtcctgcgctggtgaaacccacacagaccctcaca

ctgacctgcaccttctctgggttctcactcagcactagtggaatgtgtgtgagc

tggatccgtcagcccccagggaaggccctggagtggcttgcacgcattgattgggat

gatgataaatactacagcacatctctgaagaccaggctcaccatctccaag

gacacctccaaaaaccaggtggtccttacaatgaccaacatggaccctgtggacacagcc

acgtattactgtgcacggatac

>IGHV2-70*16

caggtcaccttgaaggagtctggtcctgtgctggtgaaacccacacagaccctcaca

ctgacctgcaccttctctgggttctcactcagcactagtggaatgtgtgtgagc

tggatccgtcagcccccagggaaggccctggagtggcttgcacgcattgattgggat

gatgataaattctacagcacatctctgaagaccaggctcaccatctccaag

gacacctccaaaaaccaggtggtccttacaatgaccaacatggaccctgtggacacagcc

acgtattactgtgcacggatac

>IGHV2-70*17

caggtcaccttgagggagtctggtcctgcgctggtgaaacccacacagaccctcaca

ctgacctgcaccttctctgggttctcactcagcactagtggaatgtgtgtgagc

tggatccgtcagcccccagggaaggccctggagtggcttgcacgcattgattgggat

gatgataaattctacagcacatctctgaagaccaggctcaccatctccaag

gacacctccaaaaaccaggtggtccttacaatgaccaacatggaccctgtggacacagcc

acgtattactgtgcacggatac

>IGHV2-70*18

caggtcaccttgagggagtctggtcctgcgctggtgaaacccacacagaccctcacc

ctgacctgcaccttctctgggttctcactcagcactagtgaaatgtgtgtgagc

tgggtccgtcagcccccagggaaggccctggagtggcttgcactcattgattgggat

gatgataaatactacagcacatctctgaagaccaggctcaccatctccaag

gacacctccaaaaaccaggtggtccttacaatgaccaacatggaccctgtggacacagcc

acgtattactgtgcacggatac

>IGHV2-70*19

caggtcaccttgagggagtctggtcctgcgctggtgaaacccacacagaccctcaca

ctgacctgcaccttctctgggttctcactcagcactagtggaatgtgtgtgagc

tgggtccgtcagcccccagggaaggccctggagtggcttgcactcattgattgggat

gatgataaacactacagcacatctctgaagaccaggctcaccatctccaag

gacacctccaaaaaccaggtggtccttacaatgaccaacatggaccctgtggacacagcc

acgtattactgtgcacggatac

>IGHV2-70D*04

caggtcaccttgaaggagtctggtcctgcgctggtgaaacccacacagaccctcaca

ctgacctgcaccttctctgggttctcactcagcactagtggaatgcgtgtgagc

tggatccgtcagcccccagggaaggccctggagtggcttgcacgcattgattgggat

gatgataaattctacagcacatctctgaagaccaggctcaccatctccaag

gacacctccaaaaaccaggtggtccttacaatgaccaacatggaccctgtggacacagcc

acgtattactgtgcacggatac

>IGHV2-70D*14

caggtcaccttgaaggagtctggtcctgcgctggtgaaacccacacagaccctcaca

ctgacctgcaccttctctgggttctcactcagcactagtggaatgcgtgtgagc

tggatccgtcagcccccaggtaaggccctggagtggcttgcacgcattgattgggat

gatgataaattctacagcacatctctgaagaccaggctcaccatctccaag

gacacctccaaaaaccaggtggtccttacaatgaccaacatggaccctgtggacacagcc

acgtattactgtgcacggatac

>IGHV2/OR16-5*01

caggtcaccttgaaggagtctggtcctgcgctggtgaaacccacagagaccctcacg

ctgacctgcactctctctgggttctcactcagcacttctggaatgggtatgagc

tggatccgtcagcccccagggaaggccctggagtggcttgctcacatttttttgaat

gacaaaaaatcctacagcacgtctctgaagaacaggctcatcatctccaag

gacacctccaaaagccaggtggtccttaccatgaccaacatggaccctgtggacacagcc

acgtattactgtgcatggagag

>IGHV3-11*01

caggtgcagctggtggagtctgggggaggcttggtcaagcctggagggtccctgaga

ctctcctgtgcagcctctggattcaccttcagtgactactacatgagc

tggatccgccaggctccagggaaggggctggagtgggtttcatacattagtagtagt

ggtagtaccatatactacgcagactctgtgaagggccgattcaccatctccagg

gacaacgccaagaactcactgtatctgcaaatgaacagcctgagagccgaggacacggcc

gtgtattactgtgcgagaga

>IGHV3-11*03

caggtgcagctgttggagtctgggggaggcttggtcaagcctggagggtccctgaga

ctctcctgtgcagcctctggattcaccttcagtgactactacatgagc

tggatccgccaggctccagggaaggggctggagtgggtttcatacattagtagtagt

agtagttacacaaactacgcagactctgtgaagggccgattcaccatctccaga

gacaacgccaagaactcactgtatctgcaaatgaacagcctgagagccgaggacacggcc

gtgtattactgtgcgaga

>IGHV3-11*04

caggtgcagctggtggagtctgggggaggcttggtcaagcctggagggtccctgaga

ctctcctgtgcagcctctggattcaccttcagtgactactacatgagc

tggatccgccaggctccagggaaggggctggagtgggtttcatacattagtagtagt

ggtagtaccatatactacgcagactctgtgaagggccgattcaccatctccagg

gacaacgccaagaactcactgtatctgcaaatgaacagcctgagagccgaggacacggct

gtgtattactgtgcgagaga

>IGHV3-11*05

caggtgcagctggtggagtctgggggaggcttggtcaagcctggagggtccctgaga

ctctcctgtgcagcctctggattcaccttcagtgactactacatgagc

tggatccgccaggctccagggaaggggctggagtgggtttcatacattagtagtagt

agtagttacacaaactacgcagactctgtgaagggccgattcaccatctccaga

gacaacgccaagaactcactgtatctgcaaatgaacagcctgagagccgaggacacggcc

gtgtattactgtgcgagaga

>IGHV3-11*06

caggtgcagctggtggagtctgggggaggcttggtcaagcctggagggtccctgaga

ctctcctgtgcagcctctggattcaccttcagtgactactacatgagc

tggatccgccaggctccagggaaggggctggagtgggtttcatacattagtagtagt

agtagttacacaaactacgcagactctgtgaagggccgattcaccatctccaga

gacaacgccaagaactcactgtatctgcaaatgaacagcctgagagccgaggacacggct

gtgtattactgtgcgagaga

>IGHV3-13*01

gaggtgcagctggtggagtctgggggaggcttggtacagcctggggggtccctgaga

ctctcctgtgcagcctctggattcaccttcagtagctacgacatgcac

tgggtccgccaagctacaggaaaaggtctggagtgggtctcagctattggtactgct

ggtgacacatactatccaggctccgtgaagggccgattcaccatctccaga

gaaaatgccaagaactccttgtatcttcaaatgaacagcctgagagccggggacacggct

gtgtattactgtgcaagaga

>IGHV3-13*02

gaggtgcatctggtggagtctgggggaggcttggtacagcctgggggggccctgaga

ctctcctgtgcagcctctggattcaccttcagtaactacgacatgcac

tgggtccgccaagctacaggaaaaggtctggagtgggtctcagccaatggtactgct

ggtgacacatactatccaggctccgtgaaggggcgattcaccatctccaga

gaaaatgccaagaactccttgtatcttcaaatgaacagcctgagagccggggacacggct

gtgtattactgtgcaagaga

>IGHV3-13*03

gaggtgcagctggtggagtctgggggaggcttggtacagcctggggggtccctgaga

ctctcctgtgcagcctgtggattcaccttcagtagctacgacatgcac

tgggtccgccaagctacaggaaaaggtctggagtgggtctcagctattggtactgct

ggtgacacatactatccaggctccgtgaagggccaattcaccatctccaga

gaaaatgccaagaactccttgtatcttcaaatgaacagcctgagagccggggacacggct

gtgtattactgtgcaaga

>IGHV3-13*04

gaggtgcagctggtggagtctgggggaggcttggtacagcctggggggtccctgaga

ctctcctgtgcagcctctggattcaccttcagtagctacgacatgcac

tgggtccgccaagctacaggaaaaggtctggaatgggtctcagctattggtactgct

ggtgacacatactatccaggctccgtgaagggccgattcaccatctccaga

gaaaatgccaagaactccttgtatcttcaaatgaacagcctgagagccggggacacggct

gtgtattactgtgcaagaga

>IGHV3-13*05

gaggtgcagctggtggagtctgggggaggcttggtacagcctggggggtccctgaga

ctctcctgtgcagcctctggattcaccttcagtagctacgacatgcac

tgggtccgccaagctacaggaaaaggtctggagtgggtctcagctattggtactgct

ggtgacccatactatccaggctccgtgaagggccgattcaccatctccaga

gaaaatgccaagaactccttgtatcttcaaatgaacagcctgagagccggggacacggct

gtgtattactgtgcaagaga

>IGHV3-15*01

gaggtgcagctggtggagtctgggggaggcttggtaaagcctggggggtcccttaga

ctctcctgtgcagcctctggattcactttcagtaacgcctggatgagc

tgggtccgccaggctccagggaaggggctggagtgggttggccgtattaaaagcaaaact

gatggtgggacaacagactacgctgcacccgtgaaaggcagattcaccatctcaaga

gatgattcaaaaaacacgctgtatctgcaaatgaacagcctgaaaaccgaggacacagcc

gtgtattactgtaccacaga

>IGHV3-15*02

gaggtgcagctggtggagtctgggggagccttggtaaagcctggggggtcccttaga

ctctcctgtgcagcctctggattcactttcagtaacgcctggatgagc

tgggtccgccaggctccagggaaggggctggagtgggttggccgtattaaaagcaaaact

gatggtgggacaacagactacgctgcacccgtgaaaggcagattcaccatctcaaga

gatgattcaaaaaacacgctgtatctgcaaatgaacagcctgaaaaccgaggacacagcc

gtgtattactgtaccacaga

>IGHV3-15*03

gaggtgcagctggtggagtctgccggagccttggtacagcctggggggtcccttaga

ctctcctgtgcagcctctggattcacttgcagtaacgcctggatgagc

tgggtccgccaggctccagggaaggggctggagtgggttggccgtattaaaagcaaagct

aatggtgggacaacagactacgctgcacctgtgaaaggcagattcaccatctcaaga

gttgattcaaaaaacacgctgtatctgcaaatgaacagcctgaaaaccgaggacacagcc

gtgtattactgtaccacaga

>IGHV3-15*04

gaggtgcagctggtggagtctgggggaggcttggtaaagcctggggggtcccttaga

ctctcctgtgcagcctctggattcactttcagtaacgcctggatgagc

tgggtccgccaggctccagggaaggggctggagtgggttggccgtattgaaagcaaaact

gatggtgggacaacagactacgctgcacccgtgaaaggcagattcaccatctcaaga

gatgattcaaaaaacacgctgtatctgcaaatgaacagcctgaaaaccgaggacacagcc

gtgtattactgtaccacaga

>IGHV3-15*05

gaggtgcagctggtggagtctgggggaggcttggtaaagcctggggggtcccttaga

ctctcctgtgcagcctctggattcactttcagtaacgcctggatgagc

tgggtccgccaggctccagggaaggggctggagtgggttggccgtattaaaagcaaaact

gatggtgggacaacagactacgctgcacccgtgaaaggcagattcaccatctcaaga

gatgattcaaaaaacacgctgtatctgcaaatgaacagtctgaaaaccgaggacacagcc

gtgtattactgtaccacaga

>IGHV3-15*06

gaggtgcagctggtggagtctgggggaggcttggtaaagcctggggggtcccttaga

ctctcctgtgcagcctctggattcactttcagtaacgcctggatgagc

tgggtccgccaggctccagggaaggggctggagtgggtcggccgtattaaaagcaaaact

gatggtgggacaacaaactacgctgcacccgtgaaaggcagattcaccatctcaaga

gatgattcaaaaaacacgctgtatctgcaaatgaacagcctgaaaaccgaggacacagcc

gtgtattactgtaccacaga

>IGHV3-15*07

gaggtgcagctggtggagtctgggggaggcttggtaaagcctggggggtcccttaga

ctctcctgtgcagcctctggtttcactttcagtaacgcctggatgaac

tgggtccgccaggctccagggaaggggctggagtgggtcggccgtattaaaagcaaaact

gatggtgggacaacagactacgctgcacccgtgaaaggcagattcaccatctcaaga

gatgattcaaaaaacacgctgtatctgcaaatgaacagcctgaaaaccgaggacacagcc

gtgtattactgtaccacaga

>IGHV3-15*08

gaggtgcagctggtggagtctgcgggaggcttggtacagcctggggggtcccttaga

ctctcctgtgcagcctctggattcacttgcagtaacgcctggatgagc

tgggtccgccaggctccagggaaggggctggagtgggttggctgtattaaaagcaaagct

aatggtgggacaacagactacgctgcacctgtgaaaggcagattcaccatctcaaga

gatgattcaaaaaacacgctgtatctgcaaatgatcagcctgaaaaccgaggacacggcc

gtgtattactgtaccacagg

>IGHV3-16*01

gaggtacaactggtggagtctgggggaggcttggtacagcctggggggtccctgaga

ctctcctgtgcagcctctggattcaccttcagtaacagtgacatgaac

tgggcccgcaaggctccaggaaaggggctggagtgggtatcgggtgttagttggaat

ggcagtaggacgcactatgtggactccgtgaagcgccgattcatcatctccaga

gacaattccaggaactccctgtatctgcaaaagaacagacggagagccgaggacatggct

gtgtattactgtgtgagaaa

>IGHV3-16*02

gaggtgcagctggtggagtctgggggaggcttggtacagcctggggggtccctgaga

ctctcctgtgcagcctctggattcaccttcagtaacagtgacatgaac

tgggcccgcaaggctccaggaaaggggctggagtgggtatcgggtgttagttggaat

ggcagtaggacgcactatgtggactccgtgaagcgccgattcatcatctccaga

gacaattccaggaactccctgtatctgcaaaagaacagacggagagccgaggacatggct

gtgtattactgtgtgagaaa

>IGHV3-19*01

acagtgcagctggtggagtctgggggaggcttggtagagcctggggggtccctgaga

ctctcctgtgcagcctctggattcaccttcagtaacagtgacatgaac

tgggtccgccaggctccaggaaaggggctggagtgggtatcgggtgttagttggaat

ggcagtaggacgcactatgcagactctgtgaagggccgattcatcatctccaga

gacaattccaggaacttcctgtatcagcaaatgaacagcctgaggcccgaggacatggct

gtgtattactgtgtgagaaa

>IGHV3-20*01

gaggtgcagctggtggagtctgggggaggtgtggtacggcctggggggtccctgaga

ctctcctgtgcagcctctggattcacctttgatgattatggcatgagc

tgggtccgccaagctccagggaaggggctggagtgggtctctggtattaattggaat

ggtggtagcacaggttatgcagactctgtgaagggccgattcaccatctccaga

gacaacgccaagaactccctgtatctgcaaatgaacagtctgagagccgaggacacggcc

ttgtatcactgtgcgagaga

>IGHV3-20*02

gaggtgcagctggtggagtctgggggaggtgtggtacggcctggggggtccctgaga

ctctcctttgcagcctctggattcacctttgatgattatggcatgagc

tgggtccgccaagctccagggaaggggctggagtgggtctctggtattaattggaat

ggtggtagcacaggttatgcagactctgtgaagggccgattcaccatctccaga

gacaacgccaagaactccctgtatctgcaaatgaacagtctgagagccgaggacacggcc

ttgtatcactgtgcgagaga

>IGHV3-20*03

gaggtgcagctggtggagtctgggggaggtgtggtacggcctggggggtccctgaga

ctctcctttgcagcctctggattcacctttgatgattatggcatgagc

tgggtccgccaagctccagggaaggggctggagtgggtctctggtattaattggaat

ggtggtagcacaggttatgcagactctgtgaagggccgattcaccatctccaga

gacaacgccaagaactccctgtatctgcaaatgaacagtctgagagccgaggacacggcc

ttgtattactgtgcgagaga

>IGHV3-20*04

gaggtgcagctggtggagtctgggggaggtgtggtacggcctggggggtccctgaga

ctctcctgtgcagcctctggattcacctttgatgattatggcatgagc

tgggtccgccaagctccagggaaggggctggagtgggtctctggtattaattggaat

ggtggtagcacaggttatgcagactctgtgaagggccgattcaccatctccaga

gacaacgccaagaactccctgtatctgcaaatgaacagtctgagagccgaggacacggcc

ttgtattactgtgcgagaga

>IGHV3-21*01

gaggtgcagctggtggagtctgggggaggcctggtcaagcctggggggtccctgaga

ctctcctgtgcagcctctggattcaccttcagtagctatagcatgaac

tgggtccgccaggctccagggaaggggctggagtgggtctcatccattagtagtagt

agtagttacatatactacgcagactcagtgaagggccgattcaccatctccaga

gacaacgccaagaactcactgtatctgcaaatgaacagcctgagagccgaggacacggct

gtgtattactgtgcgagaga

>IGHV3-21*02

gaggtgcaactggtggagtctgggggaggcctggtcaagcctggggggtccctgaga

ctctcctgtgcagcctctggattcaccttcagtagctatagcatgaac

tgggtccgccaggctccagggaaggggctggagtgggtctcatccattagtagtagt

agtagttacatatactacgcagactcagtgaagggccgattcaccatctccaga

gacaacgccaagaactcactgtatctgcaaatgaacagcctgagagccgaggacacggct

gtgtattactgtgcgagaga

>IGHV3-21*03

gaggtgcagctggtggagtctgggggaggcctggtcaagcctggggggtccctgaga

ctctcctgtgcagcctctggattcaccttcagtagctatagcatgaac

tgggtccgccaggctccagggaaggggctggagtgggtctcatccattagtagtagt

agtagttacatatactacgcagactcagtgaagggccgattcaccatctccaga

gacaacgccaagaactcactgtatctgcaaatgaacagcctgagagccgaggacacagct

gtgtattactgtgcgagaga

>IGHV3-21*04

gaggtgcagctggtggagtctgggggaggcctggtcaagcctggggggtccctgaga

ctctcctgtgcagcctctggattcaccttcagtagctatagcatgaac

tgggtccgccaggctccagggaaggggctggagtgggtctcatccattagtagtagt

agtagttacatatactacgcagactcagtgaagggccgattcaccatctccaga

gacaacgccaagaactcactgtatctgcaaatgaacagcctgagagccgaggacacggcc

gtgtattactgtgcgagaga

>IGHV3-21*05

gaggtgcagctggtggagtctgggggaggcctggtcaagcctggggggtccctgaga

ctctcctgtgcagcctctggattcaccttcagtagctatagcatgaac

tgggtccgccaggctccagggaaggggctggagtgggtttcatacattagtagtagt

agtagttacatatactacgcagactcagtgaagggccgattcaccatctccaga

gacaacgccaagaactcactgtatctgcaaatgaacagcctgagagccgaggacacggct

gtgtattactgtgcgagaga

>IGHV3-22*01

gaggtgcatctggtggagtctgggggagccttggtacagcctggggggtccctgaga

ctctcctgtgcagcctctggattcaccttcagttactactacatgagc

ggggtccgccaggctcccgggaaggggctggaatgggtaggtttcattagaaacaaagct

aatggtgggacaacagaatagaccacgtctgtgaaaggcagattcacaatctcaaga

gatgattccaaaagcatcacctatctgcaaatgaagagcctgaaaaccgaggacacggcc

gtgtattactgttccagaga

>IGHV3-22*02

gaggtgcagctggtggagtctgggggaggcttggtacagcctggggggtccctgaga

ctctcctgtgcagcctctggattcaccttcagttactactacatgagc

ggggtccgccaggctcccgggaaggggctggaatgggtaggtttcattagaaacaaagct

aatggtgggacaacagaatagaccacgtctgtgaaaggcagattcacaatctcaaga

gatgattccaaaagcatcacctatctgcaaatgaagagcctgaaaaccgaggacacggcc

gtgtattactgttccagaga

>IGHV3-23*01

gaggtgcagctgttggagtctgggggaggcttggtacagcctggggggtccctgaga

ctctcctgtgcagcctctggattcacctttagcagctatgccatgagc

tgggtccgccaggctccagggaaggggctggagtgggtctcagctattagtggtagt

ggtggtagcacatactacgcagactccgtgaagggccggttcaccatctccaga

gacaattccaagaacacgctgtatctgcaaatgaacagcctgagagccgaggacacggcc

gtatattactgtgcgaaaga

>IGHV3-23*02

gaggtgcagctgttggagtctgggggaggcttggtacagcctggggggtccctgaga

ctctcctgtgcagcctctggattcacctttagcagctatgccatgagc

tgggtccgccaggctccagggaaggggctggagtgggtctcagctattagtggtagt

ggtggtagcacatactacggagactccgtgaagggccggttcaccatctcaaga

gacaattccaagaacacgctgtatctgcaaatgaacagcctgagagccgaggacacggcc

gtatattactgtgcgaaaga

>IGHV3-23*03

gaggtgcagctgttggagtctgggggaggcttggtacagcctggggggtccctgaga

ctctcctgtgcagcctctggattcacctttagcagctatgccatgagc

tgggtccgccaggctccagggaaggggctggagtgggtctcagttatttatagcggt

ggtagtagcacatactatgcagactccgtgaagggccggttcaccatctccaga

gataattccaagaacacgctgtatctgcaaatgaacagcctgagagccgaggacacggcc

gtatattactgtgcgaaaga

>IGHV3-23*04

gaggtgcagctggtggagtctgggggaggcttggtacagcctggggggtccctgaga

ctctcctgtgcagcctctggattcacctttagcagctatgccatgagc

tgggtccgccaggctccagggaaggggctggagtgggtctcagctattagtggtagt

ggtggtagcacatactacgcagactccgtgaagggccggttcaccatctccaga

gacaattccaagaacacgctgtatctgcaaatgaacagcctgagagccgaggacacggcc

gtatattactgtgcgaaaga

>IGHV3-23*05

gaggtgcagctgttggagtctgggggaggcttggtacagcctggggggtccctgaga

ctctcctgtgcagcctctggattcacctttagcagctatgccatgagc

tgggtccgccaggctccagggaaggggctggagtgggtctcagctatttatagcagt

ggtagtagcacatactatgcagactccgtgaagggccggttcaccatctccaga

gacaattccaagaacacgctgtatctgcaaatgaacagcctgagagccgaggacacggcc

gtatattactgtgcgaaa

>IGHV3-23D*01

gaggtgcagctgttggagtctgggggaggcttggtacagcctggggggtccctgaga

ctctcctgtgcagcctctggattcacctttagcagctatgccatgagc

tgggtccgccaggctccagggaaggggctggagtgggtctcagctattagtggtagt

ggtggtagcacatactacgcagactccgtgaagggccggttcaccatctccaga

gacaattccaagaacacgctgtatctgcaaatgaacagcctgagagccgaggacacggcc

gtatattactgtgcgaaaga

>IGHV3-25*01

gagatgcagctggtggagtctgggggaggcttgcaaaagcctgcgtggtccccgaga

ctctcctgtgcagcctctcaattcaccttcagtagctactacatgaac

tgtgtccgccaggctccagggaatgggctggagttggtttgacaagttaatcctaat

gggggtagcacatacctcatagactccggtaaggaccgattcaatacctccaga

gataacgccaagaacacacttcatctgcaaatgaacagcctgaaaaccgaggacacggcc

ctctattagtgtaccagaga

>IGHV3-25*02

gagatgcagctggtggagtctgggggaggcttggcaaagcctgcgtggtccccgaga

ctctcctgtgcagcctctcaattcaccttcagtagctactacatgaac

tgtgtccgccaggctccagggaatgggctggagttggtttgacaagttaatcctaat

gggggtagcacatacctcatagactccggtaaggaccgattcaatacctccaga

gataacgccaagaacacacttcatctgcaaatgaacagcctgaaaaccgaggacacggcc

ctctattagtgtaccagaga

>IGHV3-25*03

gagatgcagctggtggagtctgggggaggcttggcaaagcctgcgtggtccccgaga

ctctcctgtgcagcctctcaattcaccttcagtagctactacatgaac

tgtgtccgccaggctccagggaatgggctggagttggttggacaagttaatcctaat

gggggtagcacatacctcatagactccggtaaggaccgattcaatacctccaga

gataacgccaagaacacacttcatctgcaaatgaacagcctgaaaaccgaggacacggcc

ctgtattagtgtaccaga

>IGHV3-25*04

gagacgcagctggtggagtctgggggaggcttggcaaagcctgggcggtccccgaga

ctctcctgtgcagcctctcaattcaccttcagtagctactacatgaac

tgtgtccgccaggctccagggaatgggctggagttggttggacaagttaatcctaat

gggggtagcacatacctcatagactccggtaaggaccgattcaatacctccaga

gataacgccaagaacacacttcatctgcaaatgaacagcctgaaaaccgaggacacggcc

ctgtattactgtaccagaga

>IGHV3-25*05

gagatgcagctggtggagtctgggggaggcttggcaaagcctgcgtggtccccgaga

ctctcctgtgcagcctctcaattcaccttcagtagctactacatgaac

tgtgtccgccaggctccagggaatgggctggagttggttggacaagttaatcctaat

gggggtagcacatacctcatagactccggtaaggaccgattcaatacctccaga

gataacgccaagaacacacttcatctgcaaatgaacagcctgaaaaccgaggacacggcc

ctctattagtgtaccagaga

>IGHV3-29*01

gaggtggagctgatagagcccacagaggacctgagacaacctgggaagttcctgaga

ctctcctgtgtagcctctagattcgccttcagtagcttctgaatgagc

ccagttcaccagtctgcaggcaaggggctggagtgagtaatagatataaaagatgat

ggaagtcagatacaccatgcagactctgtgaagggcagattctccatctccaaa

gacaatgctaagaactctctgtatctgcaaatgaacagtcagagaactgaggacatggct

gtgtatggctgtacataaggtt

>IGHV3-30*01

caggtgcagctggtggagtctgggggaggcgtggtccagcctgggaggtccctgaga

ctctcctgtgcagcctctggattcaccttcagtagctatgctatgcac

tgggtccgccaggctccaggcaaggggctagagtgggtggcagttatatcatatgat

ggaagtaataaatactacgcagactccgtgaagggccgattcaccatctccaga

gacaattccaagaacacgctgtatctgcaaatgaacagcctgagagctgaggacacggct

gtgtattactgtgcgagaga

>IGHV3-30*02

caggtgcagctggtggagtctgggggaggcgtggtccagcctggggggtccctgaga

ctctcctgtgcagcgtctggattcaccttcagtagctatggcatgcac

tgggtccgccaggctccaggcaaggggctggagtgggtggcatttatacggtatgat

ggaagtaataaatactatgcagactccgtgaagggccgattcaccatctccaga

gacaattccaagaacacgctgtatctgcaaatgaacagcctgagagctgaggacacggct

gtgtattactgtgcgaaaga

>IGHV3-30*03

caggtgcagctggtggagtctgggggaggcgtggtccagcctgggaggtccctgaga

ctctcctgtgcagcctctggattcaccttcagtagctatggcatgcac

tgggtccgccaggctccaggcaaggggctggagtgggtggcagttatatcatatgat

ggaagtaataaatactatgcagactccgtgaagggccgattcaccatctccaga

gacaattccaagaacacgctgtatctgcaaatgaacagcctgagagctgaggacacggct

gtgtattactgtgcgagaga

>IGHV3-30*04

caggtgcagctggtggagtctgggggaggcgtggtccagcctgggaggtccctgaga

ctctcctgtgcagcctctggattcaccttcagtagctatgctatgcac

tgggtccgccaggctccaggcaaggggctggagtgggtggcagttatatcatatgat

ggaagtaataaatactacgcagactccgtgaagggccgattcaccatctccaga

gacaattccaagaacacgctgtatctgcaaatgaacagcctgagagctgaggacacggct

gtgtattactgtgcgagaga

>IGHV3-30*05

caggtgcagctggtggagtctgggggaggcgtggtccagcctgggaggtccctgaga

ctctcctgtgcagcctctggattcaccttcagtagctatggcatgcac

tgggtccgccaggctccaggcaaggggctagagtgggtggcagttatatcatatgat

ggaagtaataaatactacgcagactccgtgaagggccgattcaccatctccaga

gacaattccaagaacacgctgtatctgcaaatgaacagcctgagagctgagggcacggct

gtgtattactgtgcgagaga

>IGHV3-30*06

caggtgcagctggtggagtctgggggaggcgtggtccagcctgggaggtccctgaga

ctctcctgtgcagcgtctggattcaccttcagtagctatggcatgcac

tgggtccgccaggctccaggcaaggggctagagtgggtggcagttatatcatatgat

ggaagtaataaatactacgcagactccgtgaagggccgattcaccatctccaga

gacaattccaagaacacgctgtatctgcaaatgaacagcctgagagctgaggacacggct

gtgtattactgtgcgagaga

>IGHV3-30*07

caggtgcagctggtggagtctgggggaggcgtggtccagcctgggaggtccctgaga

ctctcctgtgcagcctctggattcaccttcagtagctatgctatgcac

tgggtccgccaggctccaggcaaggggctagagtgggtggcagttatatcatatgat

ggaagtaataaatactacgcagactccgtgaagggccgattcaccatctccaga

gacaattccaagaacacgctgtatctgcaaatgaacagcctgagagccgaggacacggct

gtgtattactgtgcgagaga

>IGHV3-30*08

caggtgcagctggtggactctgggggaggcgtggtccagcctgggaggtccctgaga

ctctcctgtgcagcctctgcattcaccttcagtagctatgctatgcac

tgggtccgccaggctccaggcaaggggctagagtgggtggcagttatatcatatgat

ggaagtaataaatactacgcagactccgtgaagggccgattcaccatctccaga

gacaattccaagaacacgctgtatctgcaaatgaacagcctgagagctgaggacacggct

gtgtattactgtgcgaga

>IGHV3-30*09

caggtgcagctggtggagtctgggggaggcgtggtccagcctgggaggtccctgaga

ctctcctgtgcagcctctggattcaccttcagtagctatgctatgcac

tgggtccgccaggctccaggcaaggggctggagtgggtggcagttatatcatatgat

ggaagtaataaatactacgcagactccgtgaagggccgattcgccatctccaga

gacaattccaagaacacgctgtatctgcaaatgaacagcctgagagctgaggacacggct

gtgtattactgtgcgagaga

>IGHV3-30*10

caggtgcagctggtggagtctgggggaggcgtggtccagcctgggaggtccctgaga

ctctcctgtgcagcctctggattcaccttcagtagctatgctatgcac

tgggtccgccaggctccaggcaaggggctagagtgggtggcagttatatcatatgat

ggaagtaataaatactacacagactccgtgaagggccgattcaccatctccaga

gacaattccaagaacacgctgtatctgcaaatgaacagcctgagagctgaggacacggct

gtgtattactgtgcgagaga

>IGHV3-30*11

caggtgcagctggtggagtctgggggaggcgtggtccagcctgggaggtccctgaga

ctctcctgtgcagcgtctggattcaccttcagtagctatgctatgcac

tgggtccgccaggctccaggcaaggggctagagtgggtggcagttatatcatatgat

ggaagtaataaatactacgcagactccgtgaagggccgattcaccatctccaga

gacaattccaagaacacgctgtatctgcaaatgaacagcctgagagctgaggacacggct

gtgtattactgtgcgagaga

>IGHV3-30*12

caggtgcagctggtggagtctggggggggcgtggtccagcctgggaggtccctgaga

ctctcctgtgcagcgtctggattcaccttcagtagctatggcatgcac

tgggtccgccaggctccaggcaaggggctagagtgggtggcagttatatcatatgat

ggaagtaataaatactacgcagactccgtgaagggccgattcaccatctccaga

gacaattccaagaacacgctgtatctgcaaatgaacagcctgagagccgaggacacggct

gtgtattactgtgcgagaga

>IGHV3-30*13

caggtgcagctggtggagtctgggggaggcgtggtccagcctgggaggtccctgaga

ctctcctgtgcagcctctggattcaccttcagtagctatggcatgcac

tgggtccgccaggctccaggcaaggggctagagtgggtggcagttatatcatatgat

ggaagtaataaatactacgcagactccgtgaagggccgattcaccatctccaga

gacaattccaagaacaggctgtatctgcaaatgaacagcctgagagctgaggacacggct

gtgtattactgtgcgagaga

>IGHV3-30*14

caggtgcagctggtggagtctgggggaggcgtggtccagcctgggaggtccctgaga

ctctcctgtgcagcctctggattcaccttcagtagctatgctatgcac

tgggtccgccaggctccaggcaaggggctggagtgggtggcagttatatcatatgat

ggaagtaataaatactacgcagactccgtgaagggccgattcaccatctccaga

gacaattccaagaacacgctgtatcttcaaatgaacagcctgagagctgaggacacggct

gtgtattactgtgcgagaga

>IGHV3-30*15

caggtgcagctggtggagtctgggggaggcgtggtccagcctgggaggtccctgaga

ctctcctgtgcagcctctggattcaccttcagtagctatgctatgcac

tgggtccgccaggctccaggcaaggggctagagtgggtggcagttatatcatatgat

ggaagtaataaatactacgcagactccgtgaagggccgattcaccatctccaga

gacaattccaagaacacgctgtatctgcaaatgagcagcctgagagctgaggacacggct

gtgtattactgtgcgagaga

>IGHV3-30*16

caggtgcagctggtggagtctgggggaggcgtggtccagcctgggaggtccctgaga

ctctcctgtgcagcctctggattcaccttcagtagctatgctatgcac

tgggtccgccaggccccaggcaaggggctagagtgggtggcagttatatcatatgat

ggaagtaataaatactacgcagactccgtgaagggccgattcaccatctccaga

gacaattccaagaacacgctgtatctgcaaatgaacagcctgagagctgaggacacggct

gtgtattactgtgcgagaga

>IGHV3-30*17

caggtgcagctggtggagtctgggggaggcgtggtccagcctgggaggtccctgaga

ctctcctgtgcagcctctggattcaccttcagtagctatgctatgcac

tgggtccgccaggctccgggcaaggggctagagtgggtggcagttatatcatatgat

ggaagtaataaatactacgcagactccgtgaagggccgattcaccatctccaga

gacaattccaagaacacgctgtatctgcaaatgaacagcctgagagctgaggacacggct

gtgtattactgtgcgagaga

>IGHV3-30*18

caggtgcagctggtggagtctgggggaggcgtggtccagcctgggaggtccctgaga

ctctcctgtgcagcctctggattcaccttcagtagctatggcatgcac

tgggtccgccaggctccaggcaaggggctggagtgggtggcagttatatcatatgat

ggaagtaataaatactatgcagactccgtgaagggccgattcaccatctccaga

gacaattccaagaacacgctgtatctgcaaatgaacagcctgagagctgaggacacggct

gtgtattactgtgcgaaaga

>IGHV3-30*19

caggtgcagctggtggagtctgggggaggcgtggtccagcctgggaggtccctgaga

ctctcctgtgcagcgtctggattcaccttcagtagctatggcatgcac

tgggtccgccaggctccaggcaaggggctggagtgggtggcagttatatcatatgat

ggaagtaataaatactacgcagactccgtgaagggccgattcaccatctccaga

gacaattccaagaacacgctgtatctgcaaatgaacagcctgagagctgaggacacggct

gtgtattactgtgcgagaga

>IGHV3-30-2*01

gaggtacagctcgtggagtccggagaggacccaagacaacctgggggatccctgaga

ctctcctgtgcagactctggattaaccttcagtagctactgaaggaac

tcggtttcccaggctccagggaaggggctggagtgagtagtagatatacagtgtgat

ggaagtcagatatgttatgcataatctttgaagagcaaattcaccatctccaaa

gaaaatgccaagaactcactgtatttgctaatgaacagtctgagagcagcgggcacagct

gtgtgttactgtatgtgaggca

>IGHV3-30-22*01

gaggtggagctgatagagtccatagaggacctgagacaacctgggaagttcctgaga

ctctcctgtgtagcctctagattcgccttcagtagcttctgaatgagc

cgagttcaccagtctccaggcaaggggctggagtgagtaatagatataaaagatgat

ggaagtcagatacaccatgcagactctgtgaagggcagattctccatctccaaa

gacaatgctaagaactctctgtatctgcaaatgaacagtcagagagctgaggacatggac

gtgtatggctgtacataaggtc

>IGHV3-30-3*01

caggtgcagctggtggagtctgggggaggcgtggtccagcctgggaggtccctgaga

ctctcctgtgcagcctctggattcaccttcagtagctatgctatgcac

tgggtccgccaggctccaggcaaggggctggagtgggtggcagttatatcatatgat

ggaagcaataaatactacgcagactccgtgaagggccgattcaccatctccaga

gacaattccaagaacacgctgtatctgcaaatgaacagcctgagagctgaggacacggct

gtgtattactgtgcgagaga

>IGHV3-30-3*02

caggtgcagctggtggagtctgggggaggcgtggtccagcctgggaggtccctgaga

ctctcctgtgcagcgtctggattcaccttcagtagctatgctatgcac

tgggtccgccaggctccaggcaaggggctggagtgggtggcagttatatcatatgat

ggaagcaataaatactacgcagactccgtgaagggccgattcaccatctccaga

gacaattccaagaacacgctgtatctgcaaatgaacagcctgagagctgaggacacggct

gtgtattactgtgcgaaaga

>IGHV3-30-3*03

caggtgcagctggtggagtctgggggaggcgtggtccagcctgggaggtccctgaga

ctctcctgtgcagcctctggattcaccttcagtagctatgctatgcac

tgggtccgccaggctccaggcaaggggctggagtgggtggcagttatatcatatgat

ggaagtaataaatactacgcagactccgtgaagggccgattcaccatctccaga

gacaattccaagaacacgctgtatctgcaaatgaacagcctgagagctgaggacacggct

gtgtattactgtgcgagaga

>IGHV3-30-33*01

gaggtacagctcgtggagtccggagaggacccaagacaacctgggggatccctgaga

ctctcctgtgcagactctggattaaccttcagtagctactgaaggagc

tcggtttcccaggctccagggaaggggctggagtgagtagtagatatacagtgtgat

ggaagtcagatatgttatgcataatctttgaagagcaaattcaccatctccaaa

gaaaatgccaagaactcactgtatttgctaatgaacagtctgagagcagagggcacagct

gtgtgttactgtatgtgagg

>IGHV3-30-42*01

gaggtggagctgatagagcccacagaggacctgagacaacctgggaagttcctgaga

ctctcctgtgtagcctctagattcgccttcagtagcttctgaatgagc

ccagttcaccagtctgcaggcaaggggctggagtgagtaatagatataaaagatgat

ggaagtcagatacaccatgcagactctgtgaagggcagattctccatctccaaa

gacaatgctaagaactctctgtatctgcaaatgaacagtcagagaactgaggacatggct

gtgtatggctgtacataaggtt

>IGHV3-30-5*01

caggtgcagctggtggagtctgggggaggcgtggtccagcctgggaggtccctgaga

ctctcctgtgcagcctctggattcaccttcagtagctatggcatgcac

tgggtccgccaggctccaggcaaggggctggagtgggtggcagttatatcatatgat

ggaagtaataaatactatgcagactccgtgaagggccgattcaccatctccaga

gacaattccaagaacacgctgtatctgcaaatgaacagcctgagagctgaggacacggct

gtgtattactgtgcgaaaga

>IGHV3-30-5*02

caggtgcagctggtggagtctgggggaggcgtggtccagcctggggggtccctgaga

ctctcctgtgcagcgtctggattcaccttcagtagctatggcatgcac

tgggtccgccaggctccaggcaaggggctggagtgggtggcatttatacggtatgat

ggaagtaataaatactatgcagactccgtgaagggccgattcaccatctccaga

gacaattccaagaacacgctgtatctgcaaatgaacagcctgagagctgaggacacggct

gtgtattactgtgcgaaaga

>IGHV3-30-52*01

gaggtacagctcgtggagtccggagaggacccaagacaacctgggggatccctgaga

ctctcctgtgcagactctggattaaccttcagtagctactgaaggaac

tcggtttcccaggctccagggaaggggctggagtgagtagtagatatacagtgtgat

ggaagtcagatatgttatgcataatctttgaagagcaaattcaccatctccaaa

gaaaatgccaagaactcactgtatttgctaatgaacagtctgagagcagcgggcacagct

gtgtgttactgtatgtgagg

>IGHV3-32*01

gaggtggagctgatagagtccatagaggacctgagacaacctgggaagttcctgaga

ctctcctgtgtagcctctagattcgccttcagtagcttctgaatgagc

cgagttcaccagtctccaggcaaggggctggagtgagtaatagatataaaagatgat

ggaagtcagatacaccatgcagactctgtgaagggcagattctccatctccaaa

gacaatgctaagaactctctgtatctgcaaatgaacactcagagagctgaggacgtggcc

gtgtatggctatacataaggtc

>IGHV3-33*01

caggtgcagctggtggagtctgggggaggcgtggtccagcctgggaggtccctgaga

ctctcctgtgcagcgtctggattcaccttcagtagctatggcatgcac

tgggtccgccaggctccaggcaaggggctggagtgggtggcagttatatggtatgat

ggaagtaataaatactatgcagactccgtgaagggccgattcaccatctccaga

gacaattccaagaacacgctgtatctgcaaatgaacagcctgagagccgaggacacggct

gtgtattactgtgcgagaga

>IGHV3-33*02

caggtacagctggtggagtctgggggaggcgtggtccagcctgggaggtccctgaga

ctctcctgtgcagcgtctggattcaccttcagtagctatggcatgcac

tgggtccgccaggctccaggcaaggggctggagtgggtggcagttatatggtatgat

ggaagtaataaatactatgcagactccgcgaagggccgattcaccatctccaga

gacaattccacgaacacgctgtttctgcaaatgaacagcctgagagccgaggacacggct

gtgtattactgtgcgagaga

>IGHV3-33*03

caggtgcagctggtggagtctgggggaggcgtggtccagcctgggaggtccctgaga

ctctcctgtgcagcgtctggattcaccttcagtagctatggcatgcac

tgggtccgccaggctccaggcaaggggctggagtgggtggcagttatatggtatgat

ggaagtaataaatactatgcagactccgtgaagggccgattcaccatctccaga

gacaactccaagaacacgctgtatctgcaaatgaacagcctgagagccgaggacacggct

gtgtattactgtgcgaaaga

>IGHV3-33*04

caggtgcagctggtggagtctgggggaggcgtggtccagcctgggaggtccctgaga

ctctcctgtgcagcgtctggattcaccttcagtagctatggcatgcac

tgggtccgccaggctccaggcaaggggctagagtgggtggcagttatatggtatgac

ggaagtaataaatactatgcagactccgtgaagggccgattcaccatctccaga

gacaattccaagaacacgctgtatctgcaaatgaacagcctgagagccgaggacacggct

gtgtattactgtgcgagaga

>IGHV3-33*05

caggtgcagctggtggagtctgggggaggcgtggtccagcctgggaggtccctgaga

ctctcctgtgcagcgtctggattcaccttcagtagctatggcatgcac

tgggtccgccaggctccaggcaaggggctggagtgggtggcagttatatcatatgat

ggaagtaataaatactatgcagactccgtgaagggccgattcaccatctccaga

gacaattccaagaacacgctgtatctgcaaatgaacagcctgagagccgaggacacggct

gtgtattactgtgcgagaga

>IGHV3-33*06

caggtgcagctggtggagtctgggggaggcgtggtccagcctgggaggtccctgaga

ctctcctgtgcagcgtctggattcaccttcagtagctatggcatgcac

tgggtccgccaggctccaggcaaggggctggagtgggtggcagttatatggtatgat

ggaagtaataaatactatgcagactccgtgaagggccgattcaccatctccaga

gacaattccaagaacacgctgtatctgcaaatgaacagcctgagagccgaggacacggct

gtgtattactgtgcgaaaga

>IGHV3-33*07

caggtgcagctggtggagtctgggggacgcgtggtccagcctgggaggtccctgaga

ctctcctgtgcagcgtctggattcaccttcagtaggtatggcatgtac

tgggtccgccaggctccaggcaaggggctggagtgggtggcagttatatggtatgat

ggaagtaataaatactatgcagactccgtgaagggccgattcaccatctccaga

gacaattccaagaacacgctgtatctgcaaatgaacagcctgagagccgaggacacggct

gtgtattactgtgcgagaga

>IGHV3-33-2*01

gaggtacagctcgtggagtccggagaggacccaagacaacctgggggatccttgaga

ctctcctgtgcagactctggattaaccttcagtagctactgaatgagc

tcggtttcccaggctccagggaaggggctggagtgagtagtagatatacagtgtgat

ggaagtcagatatgttatgcccaatctgtgaagagcaaattcaccatctccaaa

gaaaatgccaagaactcactgtatttgcaaatgaacagtctgagagcagagggcacagct

gtgtgttactgtatgtgaggca

>IGHV3-35*01

gaggtgcagctggtggagtctgggggaggcttggtacagcctgggggatccctgaga

ctctcctgtgcagcctctggattcaccttcagtaacagtgacatgaac

tgggtccatcaggctccaggaaaggggctggagtgggtatcgggtgttagttggaat

ggcagtaggacgcactatgcagactctgtgaagggccgattcatcatctccaga

gacaattccaggaacaccctgtatctgcaaacgaatagcctgagggccgaggacacggct

gtgtattactgtgtgagaaa

>IGHV3-38*01

gaggtgcagctggtggagtctgggggaggcttggtacagcctagggggtccctgaga

ctctcctgtgcagcctctggattcaccgtcagtagcaatgagatgagc

tggatccgccaggctccagggaaggggctggagtgggtctcatccattagtggt

ggtagcacatactacgcagactccaggaagggcagattcaccatctccaga

gacaattccaagaacacgctgtatcttcaaatgaacaacctgagagctgagggcacggcc

gcgtattactgtgccagatata

>IGHV3-38*02

gaggtgcagctggtggagtctgggggaggcttggtacagcctagggggtccctgaga

ctctcctgtgcagcctctggattcaccgtcagtagcaatgagatgagc

tggatccgccaggctccagggaaggggctggagtgggtctcatccattagtggt

ggtagcacatactacgcagactccaggaagggcagattcaccatctccaga

gacaattccaagaacacgctgtatcttcaaatgaacaacctgagagctgagggcacggcc

gtgtattactgtgccagatata

>IGHV3-38*03

gaggtgcagctggtggagtctgggggaggcttggtacagcctagggggtccctgaga

ctctcctgtgcagcctctggattcaccgtcagtagcaatgagatgagc

tggatccgccaggctccagggaagggtctggagtgggtctcatccattagtggt

ggtagcacatactacgcagactccaggaagggcagattcaccatctccaga

gacaattccaagaacacgctgtatcttcaaatgaacaacctgagagctgagggcacggcc

gtgtattactgtgccagatata

>IGHV3-38-3*01

gaggtgcagctggtggagtctcggggagtcttggtacagcctggggggtccctgaga

ctctcctgtgcagcctctggattcaccgtcagtagcaatgagatgagc

tgggtccgccaggctccagggaagggtctggagtgggtctcatccattagtggt

ggtagcacatactacgcagactccaggaagggcagattcaccatctccaga

gacaattccaagaacacgctgcatcttcaaatgaacagcctgagagctgaggacacggct

gtgtattactgtaagaaaga

>IGHV3-41*02

gaggtgcagctggtggagtctgggggaggcttggtccagcctggggggtccctgaga

ctctcctgtgcagcctcaggattctcctttagtagctatggcatgagc

tgggtccgccaggctccagggaaggggctggactgagtggcacatatctggaatgat

ggaagtcagaaatactatgcagactctgtgaagggccgattcacaatctccaga

gacaattctaagagcatgctctatctgcaaatggacagtctgaaagctaaggacacggcc

atgtattactgtaccaga

>IGHV3-43*01

gaagtgcagctggtggagtctgggggagtcgtggtacagcctggggggtccctgaga

ctctcctgtgcagcctctggattcacctttgatgattataccatgcac

tgggtccgtcaagctccggggaagggtctggagtgggtctctcttattagttgggat

ggtggtagcacatactatgcagactctgtgaagggccgattcaccatctccaga

gacaacagcaaaaactccctgtatctgcaaatgaacagtctgagaactgaggacaccgcc

ttgtattactgtgcaaaagata

>IGHV3-43*02

gaagtgcagctggtggagtctgggggaggcgtggtacagcctggggggtccctgaga

ctctcctgtgcagcctctggattcacctttgatgattatgccatgcac

tgggtccgtcaagctccagggaagggtctggagtgggtctctcttattagtggggat

ggtggtagcacatactatgcagactctgtgaagggccgattcaccatctccaga

gacaacagcaaaaactccctgtatctgcaaatgaacagtctgagaactgaggacaccgcc

ttgtattactgtgcaaaagata

>IGHV3-43D*03

gaagtgcagctggtggagtctgggggagtcgtggtacagcctggggggtccctgaga

ctctcctgtgcagcctctggattcacctttgatgattatgccatgcac

tgggtccgtcaagctccggggaagggtctggagtgggtctctcttattagttgggat

ggtggtagcacctactatgcagactctgtgaagggtcgattcaccatctccaga

gacaacagcaaaaactccctgtatctgcaaatgaacagtctgagagctgaggacaccgcc

ttgtattactgtgcaaaagata

>IGHV3-43D*04

gaagtgcagctggtggagtctgggggagtcgtggtacagcctggggggtccctgaga

ctctcctgtgcagcctctggattcacctttgatgattatgccatgcac

tgggtccgtcaagctccggggaagggtctggagtgggtctctcttattagttgggat

ggtggtagcacatactatgcagactctgtgaagggtcgattcaccatctccaga

gacaacagcaaaaactccctgtatctgcaaatgaacagtctgagagctgaggacaccgcc

ttgtattactgtgcaaaagata

>IGHV3-47*01

gaggatcagctggtggagtctgggggaggcttggtacagcctggggggtccctgcga

ccctcctgtgcagcctctggattcgccttcagtagctatgctctgcac

tgggttcgccgggctccagggaagggtctggagtgggtatcagctattggtactggt

ggtgatacatactatgcagactccgtgatgggccgattcaccatctccaga

gacaacgccaagaagtccttgtatcttcatatgaacagcctgatagctgaggacatggct

gtgtattattgtgcaaga

>IGHV3-47*02

gaggatcagctggtggagtctgggggaggcttggtacagcctggggggtccctgaga

ccctcctgtgcagcctctggattcgccttcagtagctatgttctgcac

tgggttcgccgggctccagggaagggtccggagtgggtatcagctattggtactggt

ggtgatacatactatgcagactccgtgatgggccgattcaccatctccaga

gacaacgccaagaagtccttgtatcttcaaatgaacagcctgatagctgaggacatggct

gtgtattattgtgcaagaga

>IGHV3-48*01

gaggtgcagctggtggagtctgggggaggcttggtacagcctggggggtccctgaga

ctctcctgtgcagcctctggattcaccttcagtagctatagcatgaac

tgggtccgccaggctccagggaaggggctggagtgggtttcatacattagtagtagt

agtagtaccatatactacgcagactctgtgaagggccgattcaccatctccaga

gacaatgccaagaactcactgtatctgcaaatgaacagcctgagagccgaggacacggct

gtgtattactgtgcgagaga

>IGHV3-48*02

gaggtgcagctggtggagtctgggggaggcttggtacagcctggggggtccctgaga

ctctcctgtgcagcctctggattcaccttcagtagctatagcatgaac

tgggtccgccaggctccagggaaggggctggagtgggtttcatacattagtagtagt

agtagtaccatatactacgcagactctgtgaagggccgattcaccatctccaga

gacaatgccaagaactcactgtatctgcaaatgaacagcctgagagacgaggacacggct

gtgtattactgtgcgagaga

>IGHV3-48*03

gaggtgcagctggtggagtctgggggaggcttggtacagcctggagggtccctgaga

ctctcctgtgcagcctctggattcaccttcagtagttatgaaatgaac

tgggtccgccaggctccagggaaggggctggagtgggtttcatacattagtagtagt

ggtagtaccatatactacgcagactctgtgaagggccgattcaccatctccaga

gacaacgccaagaactcactgtatctgcaaatgaacagcctgagagccgaggacacggct

gtttattactgtgcgagaga

>IGHV3-48*04

gaggtgcagctggtggagtctgggggaggcttggtacagcctggggggtccctgaga

ctctcctgtgcagcctctggattcaccttcagtagctatagcatgaac

tgggtccgccaggctccagggaaggggctggagtgggtttcatacattagtagtagt

agtagtaccatatactacgcagactctgtgaagggccgattcaccatctccaga

gacaacgccaagaactcactgtatctgcaaatgaacagcctgagagccgaggacacggct

gtgtattactgtgcgagaga

>IGHV3-49*01

gaggtgcagctggtggagtctgggggaggcttggtacagccagggcggtccctgaga

ctctcctgtacagcttctggattcacctttggtgattatgctatgagc

tggttccgccaggctccagggaaggggctggagtgggtaggtttcattagaagcaaagct

tatggtgggacaacagaatacaccgcgtctgtgaaaggcagattcaccatctcaaga

gatggttccaaaagcatcgcctatctgcaaatgaacagcctgaaaaccgaggacacagcc

gtgtattactgtactagaga

>IGHV3-49*02

gaggtgcagctggtggagtctgggggaggcttggtacagccagggccgtccctgaga

ctctcctgtacagcttctggattcacctttgggtattatcctatgagc

tgggtccgccaggctccagggaaggggctggagtgggtaggtttcattagaagcaaagct

tatggtgggacaacagaatacgccgcgtctgtgaaaggcagattcaccatctcaaga

gatgattccaaaagcatcgcctatctgcaaatgaacagcctgaaaaccgaggacacagcc

gtgtattactgtactagaga

>IGHV3-49*03

gaggtgcagctggtggagtctgggggaggcttggtacagccagggcggtccctgaga

ctctcctgtacagcttctggattcacctttggtgattatgctatgagc

tggttccgccaggctccagggaaggggctggagtgggtaggtttcattagaagcaaagct

tatggtgggacaacagaatacgccgcgtctgtgaaaggcagattcaccatctcaaga

gatgattccaaaagcatcgcctatctgcaaatgaacagcctgaaaaccgaggacacagcc

gtgtattactgtactagaga

>IGHV3-49*04

gaggtgcagctggtggagtctgggggaggcttggtacagccagggcggtccctgaga

ctctcctgtacagcttctggattcacctttggtgattatgctatgagc

tgggtccgccaggctccagggaaggggctggagtgggtaggtttcattagaagcaaagct

tatggtgggacaacagaatacgccgcgtctgtgaaaggcagattcaccatctcaaga

gatgattccaaaagcatcgcctatctgcaaatgaacagcctgaaaaccgaggacacagcc

gtgtattactgtactagaga

>IGHV3-49*05

gaggtgcagctggtggagtctgggggaggcttggtaaagccagggcggtccctgaga

ctctcctgtacagcttctggattcacctttggtgattatgctatgagc

tggttccgccaggctccagggaaggggctggagtgggtaggtttcattagaagcaaagct

tatggtgggacaacagaatacgccgcgtctgtgaaaggcagattcaccatctcaaga

gatgattccaaaagcatcgcctatctgcaaatgaacagcctgaaaaccgaggacacagcc

gtgtattactgtactagaga

>IGHV3-52*01

gaggtgcagctggtggagtctgggtgaggcttggtacagcctggagggtccctgaga

ctctcctgtgcagcctctggattcaccttcagtagctcctggatgcac

tgggtctgccaggctccggagaaggggctggagtgggtggccgacataaagtgtgac

ggaagtgagaaatactatgtagactctgtgaagggccgattgaccatctccaga

gacaatgccaagaactccctctatctgcaagtgaacagcctgagagctgaggacatgacc

gtgtattactgtgtgagagg

>IGHV3-52*02

gaggtgcagctggtggagtctgggtgaggcttggtacagcctggagggtccctgaga

ctctcctgtgcagcctctggattcaccttcagtagctcctggatgcac

tgggtctgccaggctccggagaaggggcaggagtgggtggccgacataaagtgtgac

ggaagtgagaaatactatgtagactctgtgaagggccgattgaccatctccaga

gacaatgccaagaactccctctatctgcaagtgaacagcctgagagctgaggacatgacc

gtgtattactgtgtgaga

>IGHV3-52*03

gaggtgcagctggtcgagtctgggtgaggcttggtacagcctggagggtccctgaga

ctctcctgtgcagcctctggattcaccttcagtagctcctggatgcac

tgggtctgccaggctccggagaaggggctggagtgggtggccgacataaagtgtgac

ggaagtgagaaatactatgtagactctgtgaagggccgattgaccatctccaga

gacaatgccaagaactccctctatctgcaagtgaacagcctgagagctgaggacatgacc

gtgtattactgtgtgaga

>IGHV3-53*01

gaggtgcagctggtggagtctggaggaggcttgatccagcctggggggtccctgaga

ctctcctgtgcagcctctgggttcaccgtcagtagcaactacatgagc

tgggtccgccaggctccagggaaggggctggagtgggtctcagttatttatagcggt

ggtagcacatactacgcagactccgtgaagggccgattcaccatctccaga

gacaattccaagaacacgctgtatcttcaaatgaacagcctgagagccgaggacacggcc

gtgtattactgtgcgagaga

>IGHV3-53*02

gaggtgcagctggtggagactggaggaggcttgatccagcctggggggtccctgaga

ctctcctgtgcagcctctgggttcaccgtcagtagcaactacatgagc

tgggtccgccaggctccagggaaggggctggagtgggtctcagttatttatagcggt

ggtagcacatactacgcagactccgtgaagggccgattcaccatctccaga

gacaattccaagaacacgctgtatcttcaaatgaacagcctgagagccgaggacacggcc

gtgtattactgtgcgagaga

>IGHV3-53*03

gaggtgcagctggtggagtctggaggaggcttgatccagcctggggggtccctgaga

ctctcctgtgcagcctctgggttcaccgtcagtagcaactacatgagc

tgggtccgccagcctccagggaaggggctggagtgggtctcagttatttatagcggt

ggtagcacatactacgcagactctgtgaagggccgattcaccatctccaga

gacaattccaagaacacgctgtatcttcaaatgaacagcctgagagccgaggacacggcc

gtgtattactgtgctaggga

>IGHV3-53*04

gaggtgcagctggtggagtctggaggaggcttggtccagcctggggggtccctgaga

ctctcctgtgcagcctctgggttcaccgtcagtagcaactacatgagc

tgggtccgccaggctccagggaaggggctggagtgggtctcagttatttatagcggt

ggtagcacatactacgcagactccgtgaagggccgattcaccatctccaga

cacaattccaagaacacgctgtatcttcaaatgaacagcctgagagctgaggacacggcc

gtgtattactgtgcgagaga

>IGHV3-53*05

gaggtgcagctggtggagactggaggaggcttgatccagcctggggggtccctgaga

ctctcctgtgcagcctctgggttcaccgtcagtagcaactacatgagc

tgggtccgccaggctccagggaaggggctggagtgggtctcagttatttatagcggt

ggtagcacatactacgcagactccgtgaagggccgattcaccatctccaga

gacaattccaagaacacgctgtatcttcaaatgaacagcctgagagctgaggacacggcc

gtgtattactgtgcgagaga

>IGHV3-54*01

gaggtacagctggtggagtctgaagaaaaccaaagacaacttgggggatccctgaga

ctctcctgtgcagactctggattaaccttcagtagctactgaatgagc

tcagattcccaagctccagggaaggggctggagtgagtagtagatatatagtaggat

agaagtcagctatgttatgcacaatctgtgaagagcagattcaccatctccaaa

gaaaatgccaagaactcactctgtttgcaaatgaacagtctgagagcagagggcacggcc

gtgtattactgtatgtgagt

>IGHV3-54*02

gaggtacagctggtggagtctgaagaaaaccaaagacaacttgggggatccctgaga

ctctcctgtgcagactctggattaaccttcagtagctactgaatgagc

tcagattcccaggctccagggaaggggctggagtgagtagtagatatatagtacgat

agaagtcagatatgttatgcacaatctgtgaagagcagattcaccatctccaaa

gaaaatgccaagaactcactccgtttgcaaatgaacagtctgagagcagagggcacggcc

gtgtattactgtatgtgagg

>IGHV3-54*04

gaggtacagctggtggagtctgaagaaaaccaaagacaacttgggggatccctgaga

ctctcctgtgcagactctggattaaccttcagtagctactgaatgagc

tcagattcccaggctccagggaaggggctggagtgagtagtagatatatagtaggat

agaagtcagctatgttatgcacaatctgtgaagagcagattcaccatctccaaa

gaaaatgccaagaactcactctgtttgcaaatgaacagtctgagagcagagggcacggcc

gtgtattactgtatgtgagt

>IGHV3-62*01

gaggtgcagctggtggagtctggggaaggcttggtccagcctggggggtccctgaga

ctctcctgtgcagcctctggattcaccttcagtagctctgctatgcac

tgggtccgccaggctccaagaaagggtttgtagtgggtctcagttattagtacaagt

ggtgataccgtactctacacagactctgtgaagggccgattcaccatctccaga

gacaatgcccagaattcactgtctctgcaaatgaacagcctgagagccgagggcacagtt

gtgtactactgtgtgaaaga

>IGHV3-62*03

gaggtgcagctggtggagtctggggaaggcttggtccagcctggggggtccctgaga

ctctcctgtgcagcctctggattcaccttcagtagctctgctatgcac

tgggtccgccaggctccaagaaagggtttgtagtgggtctcagttattagtacaagt

ggtgataccgtactctacacagactctgtgaagggccgattcaccatctccaga

gacaatgcccagaattcactgtatctgcaaatgaacagcctgagagccgacgacatggct

gtgtattactgtgtgaaaga

>IGHV3-62*04

gaggtgcagctggtgaagtctggaggaggcttggtacagcctggggggtccctgaga

ctctcctgtgcagcctctggattcaccttcagtagctctgctatgcac

tgggtccgccaggctccaagaaagggtttggagtgggtctcagttattagtacaagt

ggtgataccgtactctacacagactctgtgaagggccgattcaccatctccaga

gacaatgcccagaattcactgtctctgcaaatgaacagcctgagagccgaggacatggct

gtgtattactgtgtgaaaga

>IGHV3-63*01

gaggtggagctgatagagtccatagagggcctgagacaacttgggaagttcctgaga

ctctcctgtgtagcctctggattcaccttcagtagctactgaatgagc

tgggtcaatgagactctagggaaggggctggagggagtaatagatgtaaaatatgat

ggaagtcagatataccatgcagactctgtgaagggcagattcaccatctccaaa

gacaatgctaagaactcaccgtatctccaaacgaacagtctgagagctgaggacatgacc

atgcatggctgtacataaggtt

>IGHV3-63*02

gaggtggagctgatagagtccatagagggcctgagacaacttgggaagttcctgaga

ctctcctgtgtagcctctggattcaccttcagtagctactgaatgagc

tgggtcaatgagactctagggaaggggctggagggagtaatagatgtaaaatatgat

ggaagtcagatataccatgcagactctgtgaagggcagattcaccatctccaaa

gacaatgctaagaactcaccgtatctgcaaacgaacagtctgagagctgaggacatgacc

atgcatggctgtacataa

>IGHV3-64*01

gaggtgcagctggtggagtctgggggaggcttggtccagcctggggggtccctgaga

ctctcctgtgcagcctctggattcaccttcagtagctatgctatgcac

tgggtccgccaggctccagggaagggactggaatatgtttcagctattagtagtaat

gggggtagcacatattatgcaaactctgtgaagggcagattcaccatctccaga

gacaattccaagaacacgctgtatcttcaaatgggcagcctgagagctgaggacatggct

gtgtattactgtgcgagaga

>IGHV3-64*02

gaggtgcagctggtggagtctggggaaggcttggtccagcctggggggtccctgaga

ctctcctgtgcagcctctggattcaccttcagtagctatgctatgcac

tgggtccgccaggctccagggaagggactggaatatgtttcagctattagtagtaat

gggggtagcacatattatgcagactctgtgaagggcagattcaccatctccaga

gacaattccaagaacacgctgtatcttcaaatgggcagcctgagagctgaggacatggct

gtgtattactgtgcgagaga

>IGHV3-64*03

gaggtgcagctggtggagtctgggggaggcttggtccagcctggggggtccctgaga

ctctcctgttcagcctctggattcaccttcagtagctatgctatgcac

tgggtccgccaggctccagggaagggactggaatatgtttcagctattagtagtaat

gggggtagcacatactacgcagactcagtgaagggcagattcaccatctccaga

gacaattccaagaacacgctgtatgtccaaatgagcagtctgagagctgaggacacggct

gtgtattactgtgtgaaaga

>IGHV3-64*04

caggtgcagctggtggagtctgggggaggcttggtccagcctggggggtccctgaga

ctctcctgttcagcctctggattcaccttcagtagctatgctatgcac

tgggtccgccaggctccagggaagggactggaatatgtttcagctattagtagtaat

gggggtagcacatactacgcagactcagtgaagggcagattcaccatctccaga

gacaattccaagaacacgctgtatctgcaaatgaacagcctgagagctgaggacacggct

gtgtattactgtgcgagaga

>IGHV3-64*05

gaggtgcagctggtggagtctgggggaggcttggtccagcctggggggtccctgaga

ctctcctgttcagcctctggattcaccttcagtagctatgctatgcac

tgggtccgccaggctccagggaagggactggaatatgtttcagctattagtagtaat

gggggtagcacatactacgcagactcagtgaagggcagattcaccatctccaga

gacaattccaagaacacgctgtatgttcaaatgagcagtctgagagctgaggacacggct

gtgtattactgtgtgaaaga

>IGHV3-64*07

gaggtgcagctggtggagtctgggggaggcttggtccagcctggggggtccctgaga

ctctcctgtgcagcctctggattcaccttcagtagctatgctatgcac

tgggtccgccaggctccagggaagggactggaatatgtttcagctattagtagtaat

gggggtagcacatattatgcagactctgtgaagggcagattcaccatctccaga

gacaattccaagaacacgctgtatcttcaaatgggcagcctgagagctgaggacatggct

gtgtattactgtgcgagaga

>IGHV3-64D*06

gaggtgcagctggtggagtctgggggaggcttggtccagcctggggggtccctgaga

ctctcctgttcagcctctggattcaccttcagtagctatgctatgcac

tgggtccgccaggctccagggaagggactggaatatgtttcagctattagtagtaat

gggggtagcacatactacgcagactccgtgaagggcagattcaccatctccaga

gacaattccaagaacacgctgtatcttcaaatgagcagtctgagagctgaggacacggct

gtgtattactgtgtgaaaga

>IGHV3-66*01

gaggtgcagctggtggagtctgggggaggcttggtccagcctggggggtccctgaga

ctctcctgtgcagcctctggattcaccgtcagtagcaactacatgagc

tgggtccgccaggctccagggaaggggctggagtgggtctcagttatttatagcggt

ggtagcacatactacgcagactccgtgaagggcagattcaccatctccaga

gacaattccaagaacacgctgtatcttcaaatgaacagcctgagagccgaggacacggct

gtgtattactgtgcgagaga

>IGHV3-66*02

gaggtgcagctggtggagtctgggggaggcttggtccagcctggggggtccctgaga

ctctcctgtgcagcctctggattcaccgtcagtagcaactacatgagc

tgggtccgccaggctccagggaaggggctggagtgggtctcagttatttatagcggt

ggtagcacatactacgcagactccgtgaagggccgattcaccatctccaga

gacaattccaagaacacgctgtatcttcaaatgaacagcctgagagctgaggacacggct

gtgtattactgtgcgaga

>IGHV3-66*03

gaggtgcagctggtggagtctggaggaggcttgatccagcctggggggtccctgaga

ctctcctgtgcagcctctgggttcaccgtcagtagcaactacatgagc

tgggtccgccaggctccagggaaggggctggagtgggtctcagttatttatagctgt

ggtagcacatactacgcagactccgtgaagggccgattcaccatctccaga

gacaattccaagaacacgctgtatcttcaaatgaacagcctgagagctgaggacacggct

gtgtattactgtgcgagaga

>IGHV3-66*04

gaggtgcagctggtggagtctgggggaggcttggtccagcctggggggtccctgaga

ctctcctgtgcagcctctggattcaccgtcagtagcaactacatgagc

tgggtccgccaggctccagggaaggggctggagtgggtctcagttatttatagcggt

ggtagcacatactacgcagactccgtgaagggcagattcaccatctccaga

gacaattccaagaacacgctgtatcttcaaatgaacagcctgagagccgaggacacggct

gtgtattactgtgcgagaca

>IGHV3-69-1*01

gaggtgcagctggtggagtctgggggaggcttggtaaagcctggggggtccctgaga

ctctcctgtgcagcctctggattcaccttcagtgactactacatgaac

tgggtccgccaggctccagggaaggggctggagtgggtctcatccattagtagtagt

agtaccatatactacgcagactctgtgaagggccgattcaccatctccaga

gacaacgccaagaactcactgtatctgcaaatgaacagcctgagagccgaggacacggct

gtgtattactgtgcgagaga

>IGHV3-69-1*02

gaggtgcagctggtggagtctgggggaggcttggtaaagcctggggggtccctgaga

ctctcctgtgcagcctctggattcaccttcagtgactactacatgaac

tgggtccgccaggctccagggaaggggctggagtgggtctcatccattagtagtagt

agtaccatatactacgcagactctgtgaagggccgattcaccatctccaga

gacaacgccaagaactcactgtatctgcaaatgaacagcctgagagccgaggacacggct

gtttattactgtgcgagaga

>IGHV3-7*01

gaggtgcagctggtggagtctgggggaggcttggtccagcctggggggtccctgaga

ctctcctgtgcagcctctggattcacctttagtagctattggatgagc

tgggtccgccaggctccagggaaggggctggagtgggtggccaacataaagcaagat

ggaagtgagaaatactatgtggactctgtgaagggccgattcaccatctccaga

gacaacgccaagaactcactgtatctgcaaatgaacagcctgagagccgaggacacggct

gtgtattactgtgcgagaga

>IGHV3-7*02

gaggtgcagctggtggagtctgggggaggcttggtccagcctggggggtccctgaga

ctctcctgtgcagcctctggattcacctttagtagctattggatgagc

tgggtccgccaggctccagggaaagggctggagtgggtggccaacataaagcaagat

ggaagtgagaaatactatgtggactctgtgaagggccgattcaccatctccaga

gacaacgccaagaactcactgtatctgcaaatgaacagcctgagagccgaggacacggct

gtgtattactgtgcgaga

>IGHV3-7*03

gaggtgcagctggtggagtctgggggaggcttggtccagcctggggggtccctgaga

ctctcctgtgcagcctctggattcacctttagtagctattggatgagc

tgggtccgccaggctccagggaaggggctggagtgggtggccaacataaagcaagat

ggaagtgagaaatactatgtggactctgtgaagggccgattcaccatctccaga

gacaacgccaagaactcactgtatctgcaaatgaacagcctgagagccgaggacacggcc

gtgtattactgtgcgagaga

>IGHV3-7*04

gaggtgcagctggtggagtctgggggaggcttggtccagcctggggggtccctgaga

ctctcctgtgcagcctctggattcacctttagtagctattggatgagc

tgggtccgccaggctccagggaaagggctggagtgggtggccaacataaagcaagat

ggaagtgagaaatactatgtggactctgtgaagggccgattcaccatctccaga

gacaacgccaagaactcactgtatctgcaaatgaacagcctgagagccgaggacacggct

gtgtattactgtgcgaggga

>IGHV3-71*01

gaggtgcagctggtggagtccgggggaggcttggtccagcctggggggtccctgaga

ctctcctgtgcagcctctggattcaccttcagtgactactacatgagc

tgggtccgccaggctcccgggaaggggctggagtgggtaggtttcattagaaacaaagct

aatggtgggacaacagaatagaccacgtctgtgaaaggcagattcacaatctcaaga

gatgattccaaaagcatcacctatctgcaaatgaacagcctgagagccgaggacacggcc

gtgtattactgtgcgagaga

>IGHV3-71*02

gaggtgcagctggtggagtctgggggaggcttggtccagcctggggggtccctgaga

ctctcctgtgcagcctctggattcaccttcagtgactactacatgagc

tgggtccgccaggctcccgggaaggggctggagtgggtaggtttcattagaaacaaagct

aatggtgggacaacagaatagaccacgtctgtgaaaggcagattcacaatctcaaga

gatgattccaaaagcatcacctatctgcaaatgaacagcctgagagccgaggacatggct

gtgtattactgtgcgagaga

>IGHV3-71*03

gaggtgcagctggtggagtctgggggaggcttggtccagcctggggggtccctgaga

ctctcctgtgcagcctctggtttcaccttcagtgactactacatgagc

tgggtccgccaggctcccgggaaggggctggagtgggtaggtttcattagaaacaaagct

aatggtgggacaacagaatagaccacgtctgtgaaaggcagattcacaatctcaaga

gatgattccaaaagcatcacctatctgcaaatgaacagcctgagagccgaggacacggct

gtgtattactgtgcgagaga

>IGHV3-71*04

gaggtgcagctggtggagtctgggggaggcttggtccagcctggggggtccctgaga

ctctcctgtgcagcctctggattcaccttcagtgactactacatgagc

tgggtccgccaggctcccgggaaggggctggagtgggtaggtttcattagaaacaaagct

aatggtgggacaacagaatagaccacgtctgtgaaaggcagattcacaatctcaaga

gatgattccaaaagcatcacctatctgcaaatgaacagcctgagagccgaggacacggcc

gtgtattactgtgcgagaga

>IGHV3-72*01

gaggtgcagctggtggagtctgggggaggcttggtccagcctggagggtccctgaga

ctctcctgtgcagcctctggattcaccttcagtgaccactacatggac

tgggtccgccaggctccagggaaggggctggagtgggttggccgtactagaaacaaagct

aacagttacaccacagaatacgccgcgtctgtgaaaggcagattcaccatctcaaga

gatgattcaaagaactcactgtatctgcaaatgaacagcctgaaaaccgaggacacggcc

gtgtattactgtgctagaga

>IGHV3-72*02

accttcagtgaccactacatggac

tgggtccgccaggctccagggaaggggctggagtgggttggccgtactagaaacaaagct

aacagctacaccacagaatacgccgcgtctgtgaaaggcagattcaccatctcaaga

gatgattcaaagaactcactgtat

>IGHV3-73*01

gaggtgcagctggtggagtctgggggaggcttggtccagcctggggggtccctgaaa

ctctcctgtgcagcctctgggttcaccttcagtggctctgctatgcac

tgggtccgccaggcttccgggaaagggctggagtgggttggccgtattagaagcaaagct

aacagttacgcgacagcatatgctgcgtcggtgaaaggcaggttcaccatctccaga

gatgattcaaagaacacggcgtatctgcaaatgaacagcctgaaaaccgaggacacggcc

gtgtattactgtactagaca

>IGHV3-73*02

gaggtgcagctggtggagtccgggggaggcttggtccagcctggggggtccctgaaa

ctctcctgtgcagcctctgggttcaccttcagtggctctgctatgcac

tgggtccgccaggcttccgggaaagggctggagtgggttggccgtattagaagcaaagct

aacagttacgcgacagcatatgctgcgtcggtgaaaggcaggttcaccatctccaga

gatgattcaaagaacacggcgtatctgcaaatgaacagcctgaaaaccgaggacacggcc

gtgtattactgtactagaca

>IGHV3-74*01

gaggtgcagctggtggagtccgggggaggcttagttcagcctggggggtccctgaga

ctctcctgtgcagcctctggattcaccttcagtagctactggatgcac

tgggtccgccaagctccagggaaggggctggtgtgggtctcacgtattaatagtgat

gggagtagcacaagctacgcggactccgtgaagggccgattcaccatctccaga

gacaacgccaagaacacgctgtatctgcaaatgaacagtctgagagccgaggacacggct

gtgtattactgtgcaagaga

>IGHV3-74*02

gaggtgcagctggtggagtctgggggaggcttagttcagcctggggggtccctgaga

ctctcctgtgcagcctctggattcaccttcagtagctactggatgcac

tgggtccgccaagctccagggaaggggctggtgtgggtctcacgtattaatagtgat

gggagtagcacaagctacgcggactccgtgaagggccgattcaccatctccaga

gacaacgccaagaacacgctgtatctgcaaatgaacagtctgagagccgaggacacggct

gtgtattactgtgcaaga

>IGHV3-74*03

gaggtgcagctggtggagtccgggggaggcttagttcagcctggggggtccctgaga

ctctcctgtgcagcctctggattcaccttcagtagctactggatgcac

tgggtccgccaagctccagggaaggggctggtgtgggtctcacgtattaatagtgat

gggagtagcacaacgtacgcggactccgtgaagggccgattcaccatctccaga

gacaacgccaagaacacgctgtatctgcaaatgaacagtctgagagccgaggacacggct

gtgtattactgtgcaagaga

>IGHV3-9*01

gaagtgcagctggtggagtctgggggaggcttggtacagcctggcaggtccctgaga

ctctcctgtgcagcctctggattcacctttgatgattatgccatgcac

tgggtccggcaagctccagggaagggcctggagtgggtctcaggtattagttggaat

agtggtagcataggctatgcggactctgtgaagggccgattcaccatctccaga

gacaacgccaagaactccctgtatctgcaaatgaacagtctgagagctgaggacacggcc

ttgtattactgtgcaaaagata

>IGHV3-9*02

gaagtgcagctggtggagtctgggggaggcttggtacagcctggcaggtccctgaga

ctctcctgtgcagcctctggattcacctctgatgattatgccatgcac

tgggtccggcaagctccagggaagggcctggagtgggtctcaggtattagttggaat

agtggtagcataggctatgcggactctgtgaagggccgattcaccatctccaga

gacaacgccaagaactccctgtatctgcaaatgaacagtctgagagctgaggacacggcc

ttgtattactgtgcaaaagata

>IGHV3-9*03

gaagtgcagctggtggagtctgggggaggcttggtacagcctggcaggtccctgaga

ctctcctgtgcagcctctggattcacctttgatgattatgccatgcac

tgggtccggcaagctccagggaagggcctggagtgggtctcaggtattagttggaat

agtggtagcataggctatgcggactctgtgaagggccgattcaccatctccaga

gacaacgccaagaactccctgtatctgcaaatgaacagtctgagagctgaggacatggcc

ttgtattactgtgcaaaagata

>IGHV3-NL1*01

caggtgcagctggtggagtctgggggaggcgtggtccagcctggggggtccctgaga

ctctcctgtgcagcgtctggattcaccttcagtagctatggcatgcac

tgggtccgccaggctccaggcaaggggctggagtgggtctcagttatttatagcggt

ggtagtagcacatactatgcagactccgtgaagggccgattcaccatctccaga

gacaattccaagaacacgctgtatctgcaaatgaacagcctgagagctgaggacacggct

gtgtattactgtgcgaaaga

>IGHV3/OR15-7*01

gaggtgcagctggtggagtctgggggaggcttggtccagcctgggggttctctgaga

ctctcatgtgcagcctctggattcaccttcagtgaccactacatgagc

tgggtccgccaggctcaagggaaagggctagagttggtaggtttaataagaaacaaagct

aacagttacacgacagaatatgctgcgtctgtgaaaggcagacttaccatctcaaga

gaggattcaaagaacacgatgtatctgcaaatgagcaacctgaaaaccgaggacttggcc

gtgtattactgtgctaga

>IGHV3/OR15-7*02

gaggtgcagctgttggagtctgggggaggcttggtccagcctgggggttctctgaga

ctctcatgtgctgcctctggattcaccttcagtgaccactacatgagc

tgggtccgccaggctcaagggaaagggctagagttggtaggtttaataagaaacaaagct

aacagttacacgacagaatatgctgcgtctgtgaaaggcagacttaccatctcaaga

gaggattcaaagaacacgctgtatctgcaaatgagcagcctgaaaaccgaggacttggcc

gtgtattactgtgctaga

>IGHV3/OR15-7*03

gaggtgcagctggtggagtctgggggaggcttggtccagcctgggggttctctgaga

ctctcatgtgcagcctctggattcaccttcagtgaccactacatgagc

tgggtccgccaggctcaagggaaagggctagagttggtaggtttaataagaaacaaagct

aacagttacacgacagaatatgctgcgtctgtgaaaggcagacttaccatctcaaga

gaggattcaaagaacacgctgtatctgcaaatgagcagcctgaaaaccgaggacttggcc

gtgtattactgtgctaga

>IGHV3/OR15-7*05

gaggtgcagctggtggagtctgggggaggcttggtccagcctgggggttctctgaga

ctctcatgtgcagcctctggattcaccttcagtgaccactacatgagc

tgggtccgccaggctcaagggaaagggctagagttggtaggtttaataagaaacaaagct

aacagttacacgacagaatatgctgcgtctgtgaaaggcagacttaccatctcaaga

gaggattcaaagaacacgctgtatctgcaaatgagcaacctgaaaaccgaggacttggcc

gtgtattactgtgctagaga

>IGHV3/OR16-10*01

gaggttcagctggtgcagtctgggggaggcttggtacatcctggggggtccctgaga

ctctcctgtgcaggctctggattcaccttcagtagctatgctatgcac

tgggttcgccaggctccaggaaaaggtctggagtgggtatcagctattggtactggt

ggtggcacatactatgcagactccgtgaagggccgattcaccatctccaga

gacaatgccaagaactccttgtatcttcaaatgaacagcctgagagccgaggacatggct

gtgtattactgtgcaaga

>IGHV3/OR16-10*02

gaggttcagctggtgcagtctgggggaggcttggtacagcctggggggtccctgaga

ctctcctgtgcaggctctggattcaccttcagtagctatgctatgcac

tgggttcgccaggctccaggaaaaggtctggagtgggtatcagctattggtactggt

ggtggcacatactatgcagactccgtgaagggccgattcaccatctccaga

gacaatgccaagaactccttgtatcttcaaatgaacagcctgagagccgaggacatggct

gtgtattactgtgcaaga

>IGHV3/OR16-10*03

gaggtgcagctggtggagtctgggggaggcttggtacagcctggggggtccctgaga

ctctcctgtgcaggctctggattcaccttcagtagctatgctatgcac

tgggttcgccaggctccaggaaaaggtctggagtgggtatcagctattggtactggt

ggtggcacatactatgcagactccgtgaagggccgattcaccatctccaga

gacaatgccaagaactccttgtatcttcaaatgaacagcctgagagccgaggacatggct

gtgtattactgtgcaagaga

>IGHV3/OR16-12*01

gaggtgcagctggtagagtctgggagaggcttggcccagcctggggggtacctaaaa

ctctccggtgcagcctctggattcaccgtcggtagctggtacatgagc

tggatccaccaggctccagggaagggtctggagtgggtctcatacattagtagtagt

ggttgtagcacaaactacgcagactctgtgaagggcagattcaccatctccaca

gacaactcaaagaacacgctctacctgcaaatgaacagcctgagagtggaggacacggcc

gtgtattactgtgcaaga

>IGHV3/OR16-13*01

gaggtgcagctggtggagtctgggggaggcttagtacagcctggagggtccctgaga

ctctcctgtgcagcctctggattcaccttcagtagctactggatgcac

tgggtccgccaagctccagggaaggggctggtgtgggtctcacgtattaatagtgat

gggagtagcacaagctacgcagactccatgaagggccaattcaccatctccaga

gacaatgctaagaacacgctgtatctgcaaatgaacagtctgagagctgaggacatggct

gtgtattactgtactaga

>IGHV3/OR16-14*01

gaggtgcagctggaggagtctgggggaggcttagtacagcctggagggtccctgaga

ctctcctgtgcagcctctggattcaccttcagtagctactggatgcac

tgggtccgccaatctccagggaaggggctggtgtgagtctcacgtattaatagtgat

gggagtagcacaagctacgcagactccttgaagggccaattcaccatctccaga

gacaatgctaagaacacgctgtatctgcaaatgaacagtctgagagctgaggacatggct

gtgtattactgtactaga

>IGHV3/OR16-15*01

gaagtgcagctggtggagtctgggggaggcttggtccagcctggggggtccctgaga

ctctcctgtgcagcctctgtattcaccttcagtaacagtgacataaac

tgggtcctctaggctccaggaaaggggctggagtgggtctcgggtattagttggaat

ggcggtaagacgcactatgtggactccgtgaagggccaattttccatctccaga

gacaattccagcaagtccctgtatctgcaaaagaacagacagagagccaaggacatggcc

gtgtattactgtgtgagaaa

>IGHV3/OR16-15*02

gaggtgcagctggtggagtctgggggaggcttggtccagcctggggggtccctgaga

cactcctgtgcagcctctggattcaccttcagtaacagtgacatgaac

tgggtcctctaggctccaggaaaggggctggagtgggtctcgggtattagttggaat

ggcggtaagacgcactatgtggactccgtgaagggccaatttaccatctccaga

gacaattccagcaagtccctgtatctgcaaaagaacagacagagagccaaagacatggcc

gtgtattactgtgtgaga

>IGHV3/OR16-16*01

gaggtgcagctggtggagtctgggggaggcttggtccagcctggggggtccctgaga

cactcctgtgcagcctctggattcaccttcagtaacagtgacatgaac

tgggtcctctaggctccaggaaaggggctggagtgggtctcggatattagttggaat

ggcggtaagacgcactatgtggactccgtgaagggccaatttaccatctccaga

gacaattccagcaagtccctgtatctgcaaaagaacagacagagagccaaggacatggcc

gtgtattactgtgtgaga

>IGHV3/OR16-6*02

gaggtgcagctggtggagtctgcgggaggccttggtacagcctgggggtcccttaga

ctctcctgtgcagcctctggattcacttgcagtaacgcctggatgagc

tgggtccgccaggctccagggaaggggctggagtgggttggctgtattaaaagcaaagct

aatggtgggacaacagactacgctgcacctgtgaaaggcagattcaccatctcaaga

gatgattcaaaaaacacgctgtatctgcaaatgatcagcctgaaaaccgaggacacggcc

gtgtattactgtaccacagg

>IGHV3/OR16-8*01

gaggtgcagctggtggagtctgggggaggcttggtacagcctggggggtccctgaga

ctgtcctgtccagcctctggattcaccttcagtaaccactacatgagc

tgggtccgccaggctccagggaagggactggagtgggtttcatacattagtggtgat

agtggttacacaaactacgcagactctgtgaagggccgattcaccatctccagg

gacaacgccaataactcaccgtatctgcaaatgaacagcctgagagctgaggacacggct

gtgtattactgtgtgaaa

>IGHV3/OR16-8*02

gaggtgcagctggtggagtctgggggaggcttggtacagcctggggggtccctgaga

ctgtcctgtccagactctggattcaccttcagtaaccactacatgagc

tgggtccgccaggctccagggaagggactggagtggatttcatacattagtggtgat

agtggttacacaaactacgcagactctgtgaagggccgattcaccatctccagg

gacaacgccaataactcaccgtatctgcaaatgaacagcttgagagctgaggacacggct

gtgtattactgtgtgaaaca

>IGHV3/OR16-9*01

gaggtgcagctggtggagtctggaggaggcttggtacagcctggggggtccctgaga

ctctcctgtgcagcctctggattcaccttcagtaaccactacacgagc

tgggtccgccaggctccagggaagggactggagtgggtttcatacagtagtggtaat

agtggttacacaaactacgcagactctgtgaaaggccgattcaccatctccagg

gacaacgccaagaactcactgtatctgcaaatgaacagcctgagagccgaggacacggct

gtgtattactgtgtgaaa

>IGHV4-28*01

caggtgcagctgcaggagtcgggcccaggactggtgaagccttcggacaccctgtcc

ctcacctgcgctgtctctggttactccatcagcagtagtaactggtggggc

tggatccggcagcccccagggaagggactggagtggattgggtacatctattatagt

gggagcacctactacaacccgtccctcaagagtcgagtcaccatgtcagta

gacacgtccaagaaccagttctccctgaagctgagctctgtgaccgccgtggacacggcc

gtgtattactgtgcgagaaa

>IGHV4-28*02

caggtgcagctgcaggagtcgggcccaggactggtgaagccttcacagaccctgtcc

ctcacctgcgctgtctctggttactccatcagcagtagtaactggtggggc

tggatccggcagcccccagggaagggactggagtggattgggtacatctattatagt

gggagcatctactacaacccgtccctcaagagtcgagtcaccatgtcagta

gacacgtccaagaaccagttctccctgaagctgagctctgtgaccgccgtggacacggcc

gtgtattactgtgcgagaaa

>IGHV4-28*03

caggtgcagctgcaggagtcgggcccaggactggtgaagccttcggacaccctgtcc

ctcacctgcgctgtctctggttactccatcagcagtagtaactggtggggc

tggatccggcagcccccagggaagggactggagtggattgggtacatctattatagt

gggagcacctactacaacccgtccctcaagagtcgagtcaccatgtcagta

gacacgtccaagaaccagttctccctgaagctgagctctgtgaccgccgtggacacggcc

gtgtattactgtgcgagaga

>IGHV4-28*04

caggtgcagctgcaggagtcgggcccaggactggtgaagccttcggacaccctgtcc

ctcacctgcgctgtctctggttactccatcagcagtagtaactggtggggc

tggatccggcagcccccagggaagggactggagtggattgggtacatctattatagt

gggagcacctactacaacccgtccctcaagagtcgagtcaccatgtcagta

gacacgtccaagaaccagttctccctgaagctgagctctgtgaccgccgtggacaccggc

gtgtattactgtgcgaga

>IGHV4-28*05

caggtgcagctgcaggagtcgggcccaggactggtgaagccttcggacaccctgtcc

ctcacctgcgctgtctctggttactccatcagcagtagtaactggtggggc

tggatccggcagcccccagggaagggactggagtggattgggtacatctattatagt

gggagcatctactacaacccgtccctcaagagtcgagtcaccatgtcagta

gacacgtccaagaaccagttctccctgaagctgagctctgtgaccgccgtggacacggcc

gtgtattactgtgcgagaaa

>IGHV4-28*06

caggtgcagctacaggagtcgggcccaggactggtgaagccttcggacaccctgtcc

ctcacctgcgctgtctctggttactccatcagcagtagtaactggtggggc

tggatccggcagcccccagggaagggactggagtggattgggtacatctattatagt

gggagcaccaactacaacccgtccctcaagagtcgagtcaccatgtcagta

gacacgtccaagaaccagttctccctgaagctgagctctgtgaccgccttggacacggcc

gtgtattactgtgcgagaaa

>IGHV4-28*07

caggtacagctgcaggagtcgggcccaggactggtgaagccttcggacaccctgtcc

ctcacctgcgctgtctctggttactccatcagcagtagtaactggtggggc

tggatccggcagcccccagggaagggactggagtggattgggtacatctattatagt

gggagcacctactacaacccgtccctcaagagtcgagtcaccatgtcagta

gacacgtccaagaaccagttctccctgaagctgagctctgtgaccgccgtggacacggcc

gtgtattactgtgcgagaaa

>IGHV4-30-2*01

cagctgcagctgcaggagtccggctcaggactggtgaagccttcacagaccctgtcc

ctcacctgcgctgtctctggtggctccatcagcagtggtggttactcctggagc

tggatccggcagccaccagggaagggcctggagtggattgggtacatctatcatagt

gggagcacctactacaacccgtccctcaagagtcgagtcaccatatcagta

gacaggtccaagaaccagttctccctgaagctgagctctgtgaccgccgcggacacggcc

gtgtattactgtgccagaga

>IGHV4-30-2*02

cagctgcagctgcaggagtccggctcaggactggtgaagccttcacagaccctgtcc

ctcacctgcgctgtctctggtggctccatcagcagtggtggttactcctggagc

tggatccggcagccaccagggaagggcctggagtggattgggtacatctatcatagt

gggagcacctactacaacccgtccctcaagagtcgagtcaccatatcagta

gacaggtccaagaaccagttctccctgaagctgagctctgtgaccgctgcggacacggcc

gtgtattactgtgcg

>IGHV4-30-2*03

cagctgcagctgcaggagtccggctcaggactggtgaagccttcacagaccctgtcc

ctcacctgcgctgtctctggtggctccatcagcagtggtggttactcctggagc

tggatccggcagccaccagggaagggcctggagtggattgggagtatctattatagt

gggagcacctactacaacccgtccctcaagagtcgagtcaccatatccgta

gacacgtccaagaaccagttctccctgaagctgagctctgtgaccgctgcagacacggct

gtgtattactgtgcgagaca

>IGHV4-30-2*04

tctggtggctccatcagcagtggtggttactcctggagc

tggatccggcagccaccagggaagggcctggagtggattgggtacatctatcatagt

gggagcacctactacaacccgtccctcaagagtcgagtcaccatatcagta

gacacgtccaagaaccagttctccctgaagctgagctctgtgaccgccgcagacacggcc

gtgtattactgtgcgagaga

>IGHV4-30-2*05

cagctgcagctgcaggagtccggctcaggactggtgaagccttcacagaccctgtcc

ctcacctgcgctgtctctggtggctccatcagcagtggtggttactcctggagc

tggatccggcagccaccagggaagggcctggagtggattgggtacatctatcatagt

gggagcacctactacaacccgtccctcaagagtcgagttaccatatcagta

gacacgtccaagaaccagttctccctgaagctgagctctgtgactgccgcagacacggcc

gtgtattactgtgccagaga

>IGHV4-30-2*06

cagctgcagctgcaggagtccggctcaggactggtgaagccttcacagaccctgtcc

ctcacctgcgctgtctctggtggctccatcagcagtggtggttactcctggagc

tggatccggcagtcaccagggaagggcctggagtggattgggtacatctatcatagt

gggagcacctactacaacccgtccctcaagagtcgagtcaccatatcagta

gacaggtccaagaaccagttctccctgaagctgagctctgtgaccgccgcggacacggcc

gtgtattactgtgccagaga

>IGHV4-30-4*01

caggtgcagctgcaggagtcgggcccaggactggtgaagccttcacagaccctgtcc

ctcacctgcactgtctctggtggctccatcagcagtggtgattactactggagt

tggatccgccagcccccagggaagggcctggagtggattgggtacatctattacagt

gggagcacctactacaacccgtccctcaagagtcgagttaccatatcagta

gacacgtccaagaaccagttctccctgaagctgagctctgtgactgccgcagacacggcc

gtgtattactgtgccagaga

>IGHV4-30-4*02

caggtgcagctgcaggagtcgggcccaggactggtgaagccttcggacaccctgtcc

ctcacctgcactgtctctggtggctccatcagcagtggtgattactactggagt

tggatccgccagcccccagggaagggcctggagtggattgggtacatctattacagt

gggagcacctactacaacccgtccctcaagagtcgagttaccatatcagta

gacacgtccaagaaccagttctccctgaagctgagctctgtgactgcagcagacacggcc

gtgtattactgtgccagaga

>IGHV4-30-4*03

caggtgcagctgcaggagtcgggcccaggactggtgaagccttcacagaccctgtcc

ctcacctgcactgtctctggtggctccatcagcagtggtgattactactggagt

tggatccgccagcccccagggaagggcctggagtggattgggtacatctattacagt

gggagcacctactacaacccgtccctcaagagtcgagttaccatatcagta

gacacgtccaagaaccagttctccctgaagctgagctctgtgactgccgcggacacggcc

gtgtattactg

>IGHV4-30-4*04

caggtgcagctgcaggactcgggcccaggactggtgaagccttcacagaccctgtcc

ctcacctgcactgtctctggtggctccatcagcagtggtgattactactggagt

tggatccgccagcccccagggaagggcctggagtggattgggtacttctattacagt

gggagcacctactacaacccgtccctcaagagtcgagttaccatatcagta

gacacgtccaagaaccagttctccctgaagctgagctctgtgactgccgcagacacggcc

gtgtattactg

>IGHV4-30-4*05

ctctggtggctccatcagcagtggtgattactactggagt

tggatccgccagcncccagggaagggcctggagtggattgggtacatctattacagt

gggagcacctactacaacccgtccctcaagagtcgagtcaccatatcagta

gacacgtccaagaaccagttctccctgaagctgagctctgtgactgccgcagacacggcc

gtgtattactgtgccagaga

>IGHV4-30-4*06

tctggtggctccatcagcagtggtgattactactggagt

tggatccgccagcacccagggaagggcctggagtggattgggtacatctattacagt

gggagcacctactacaacccgtccctcaagagtcgagttaccatatcagta

gacacgtccaagaaccagttctccctgaagctgagctctgtgactgccgcagacacggcc

gtgtattactgtgccagaga

>IGHV4-30-4*07

caggtgcagctgcaggagtcgggcccaggactggtgaagccttcacagaccctgtcc

ctcacctgcgctgtctctggtggctccatcagcagtggtggttactcctggagc

tggatccggcagccaccagggaagggactggagtggattgggtatatctattacagt

gggagcacctactacaacccgtccctcaagagtcgagttaccatatcagta

gacacgtccaagaaccagttctccctgaagctgagctctgtgaccgccgcggacacggcc

gtgtattactgtgccagaga

>IGHV4-30-4*08

caggtgcagctgcaggagtcgggcccaggactggtgaagccttcacagaccctgtcc

ctcacctgcactgtctctggtggctccatcagcagtggtgattactactggagc

tggatccgccagcccccagggaagggcctggagtggattgggtacatctattacagt

gggagcacctactacaacccgtccctcaagagtcgagttaccatatcagta

gacacgtccaagaaccagttctccctgaagctgagctctgtgactgccgcagacacggcc

gtgtattactgtgccagag

>IGHV4-31*01

caggtgcagctgcaggagtcgggcccaggactggtgaagccttcacagaccctgtcc

ctcacctgcactgtctctggtggctccatcagcagtggtggttactactggagc

tggatccgccagcacccagggaagggcctggagtggattgggtacatctattacagt

gggagcacctactacaacccgtccctcaagagtctagttaccatatcagta

gacacgtctaagaaccagttctccctgaagctgagctctgtgactgccgcggacacggcc

gtgtattactgtgcgagaga

>IGHV4-31*02

caggtgcagctgcaggagtcgggcccaggactggtgaagccttcacagaccctgtcc

ctcacctgtactgtctctggtggctccatcagcagtggtggttactactggagc

tggatccgccagcacccagggaagggcctggagtggattgggtacatctattacagt

gggagcacctactacaacccgtccctcaagagtcgagttaccatatcagta

gacacgtctaagaaccagttctccctgaagctgagctctgtgactgccgcggacacggcc

gtgtattactgtgcgagaga

>IGHV4-31*03

caggtgcagctgcaggagtcgggcccaggactggtgaagccttcacagaccctgtcc

ctcacctgcactgtctctggtggctccatcagcagtggtggttactactggagc

tggatccgccagcacccagggaagggcctggagtggattgggtacatctattacagt

gggagcacctactacaacccgtccctcaagagtcgagttaccatatcagta

gacacgtctaagaaccagttctccctgaagctgagctctgtgactgccgcggacacggcc

gtgtattactgtgcgagaga

>IGHV4-31*04

caggtgcggctgcaggagtcgggcccaggactggtgaagccttcacagaccctgtcc

ctcacctgcactgtctctggtggctccatcagcagtggtggttactactggagc

tggatccgccagcacccagggaagggcctggagtggattgggtacatctattacagt

gggagcacctactacaacccgtccctcaagagtcgagttaccatatcagta

gacacgtctaagaaccagttctccctgaagctgagctctgtgactgccgcggacacggcc

gtgtattactgtgcg

>IGHV4-31*05

caggtgcagctgcaggagtcgggcccaggactggtgaagccttcacagaccctgtcc

ctcacctgcactgtctctggtggctccatcagcagtggtggttactactggagc

tggatccgccagcacccagggaagggcctggagtggattgggtacatctattacagt

gggagcacctactacaacccgtccctcaagagtcgagttaccatatcagta

gacacgtctaagaaccagttctccctgaagctgagctctgtgaccgcggacgcggcc

gtgtattactgtgcg

>IGHV4-31*06

caggtgcagctgcaggagtcgggcccaggactggtgaagccttcacagaccctgtcc

ctcacctgcactgtctctggtggctccatcagcagtggtagttactactggagc

tggatccgccagcacccagggaagggcctggagtggattgggtacatctattacagt

gggagcacctactacaacccgtccctcaagagtcgagttaccatatcagta

gacacgtctaagaaccagttctccctgaagctgagctctgtgactgccgcggacacggcc

gtgtattactg

>IGHV4-31*07

caggtgcagctgcaggagtcgggcccaggactggtgaagccttcacagaccctgtcc

ctcacctgcactgtctctggtggatccatcagcagtggtggttactactggagc

tggatccgccagcacccagggaagggcctggagtggattgggtacatctattacagt

gggagcacctactacaacccgtccctcaagagtcgagttaccatatcagta

gacacgtctaagaaccagttctccctgaagctgagctctgtgactgccgcggacacggcc

gtgtattactg

>IGHV4-31*08

caggtgcagctgcaggagtcgggcccaggactggtgaagccttcacagaccctgtcc

ctcacctgcactgtctctggtggctccatcagcagtggtggttactactggagc

tggatccgccagcacccagggaagggcctggagtggattgggtacatctattacagt

gggagcacctactacaacccgtccctcaagagtcgagttaccatatccgta

gacacgtccaagaaccagttctccctgaagctgagctctgtgactgccgcggacacggcc

gtgtattactg

>IGHV4-31*09

caggtgcagctgcaggagtcgggcccaggactggtgaagccttcacagaccctgtcc

ctcacctgcactgtctctggtggctccatcagcagtggtggttactactggagc

tggatccgccagcacccagggaagggcctggagtggattgggtacatctattacagt

gggagcacctactacaacccgtccctcaagagtcgagttaccatatcagta

gacaagtccaagaaccagttctccctgaagctgagctctgtgaccgccgcggacacggcc

gtgtattactg

>IGHV4-31*10

caggtgcagctgcaggagtcgggcccaggactgttgaagccttcacagaccctgtcc

ctcacctgcactgtctctggtggctccatcagcagtggtggttactactggagc

tggatccgccagcacccagggaagggcctggagtggattgggtgcatctattacagt

gggagcacctactacaacccgtccctcaagagtcgagttaccatatcagta

gacccgtccaagaaccagttctccctgaagccgagctctgtgactgccgcggacacggcc

gtggattactgtgcgagaga

>IGHV4-31*11

caggtgcagctgcaggagtcgggcccaggactggtgaagccttcacagaccctgtcc

ctcacctgcgctgtctctggtggctccatcagcagtggtggttactactggagc

tggatccgccagcacccagggaagggcctggagtggattgggtacatctattacagt

gggagcacctactacaacccgtccctcaagagtcgagttaccatatcagta

gacacgtctaagaaccagttctccctgaagctgagctctgtgactgccgcggacacggcc

gtgtattactgtgcgagaga

>IGHV4-34*01

caggtgcagctacagcagtggggcgcaggactgttgaagccttcggagaccctgtcc

ctcacctgcgctgtctatggtgggtccttcagtggttactactggagc

tggatccgccagcccccagggaaggggctggagtggattggggaaatcaatcatagt

ggaagcaccaactacaacccgtccctcaagagtcgagtcaccatatcagta

gacacgtccaagaaccagttctccctgaagctgagctctgtgaccgccgcggacacggct

gtgtattactgtgcgagagg

>IGHV4-34*02

caggtgcagctacaacagtggggcgcaggactgttgaagccttcggagaccctgtcc

ctcacctgcgctgtctatggtgggtccttcagtggttactactggagc

tggatccgccagcccccagggaaggggctggagtggattggggaaatcaatcatagt

ggaagcaccaactacaacccgtccctcaagagtcgagtcaccatatcagta

gacacgtccaagaaccagttctccctgaagctgagctctgtgaccgccgcggacacggct

gtgtattactgtgcgagagg

>IGHV4-34*03

caggtgcagctacagcagtggggcgcaggactgttgaagccttcggagaccctgtcc

ctcacctgcgctgtctatggtgggtccttcagtggttactactggagc

tggatccgccagcccccagggaaggggctggagtggattggggaaatcaatcatagt

ggaagcaccaactacaacccgtccctcaagagtcgagtcaccatatcagta

gacacgtccaagaaccagttctccctgaagctgagctctgtgaccgccgcggacacggcc

gtgtattactg

>IGHV4-34*04

caggtgcagctacagcagtggggcgcaggactgttgaagccttcggagaccctgtcc

ctcacctgcgctgtctatggtgggtccttcagtggttactactggagc

tggatccgccagcccccagggaaggggctggagtggattggggaaatcaatcatagt

ggaagcaccaacaacaacccgtccctcaagagtcgagccaccatatcagta

gacacgtccaagaaccagttctccctgaagctgagctctgtgaccgccgcggacacggct

gtgtattactgtgcgagagg

>IGHV4-34*05

caggtgcagctacagcagtggggcgcaggactgttgaagccttcggagaccctgtcc

ctcacctgcgctgtctatggtgggtccttcagtggttactactggtgc

tggatccgccagcccctagggaaggggctggagtggattggggaaatcaatcatagt

ggaagcaccaacaacaacccgtccctcaagagtcgagccaccatatcagta

gacacgtccaagaaccagttctccctgaagctgagctctgtgaccgccgcggacacggct

gtgtattactgtgcgagagg

>IGHV4-34*06

caggtgcagctacagcagtggggcgcaggactgttgaagccttcggagaccctgtcc

ctcacctgcgctgtctatggtgggtccttcagtggttactactggagc

tggatccgccagcccccagggaaggggctggagtggattggggaaatcaatcatagt

ggaagcaccaactacaacccgtccctcaagagtcgagtcaccatatcagta

gacacgtccaagaaccagttctccctgaagctgggctctgtgaccgccgcggacacggcc

gtgtattactg

>IGHV4-34*07

caggtgcagctacagcagtggggcgcaggactgttgaagccttcggagaccctgtcc

ctcacctgcgctgtctatggtgggtccttcagtggttactactggagc

tggatccgccagcccccagggaaggggctggagtggattggggaaatcaaccatagt

ggaagcaccaactacaacccgtccctcaagagtcgagtcaccatatcagta

gacacgtccaagaaccagttctccctgaagctgagctctgtgaccgccgcggacacggcc

gtgtattactg

>IGHV4-34*08

caggtgcagctacagcagtggggcgcaggactgttgaagccttcggagaccctgtcc

ctcacctgcgctgtctatggtgggaccttcagtggttactactggagc

tggatccgccagcccccagggaaggggctggagtggattggggaaatcaatcatagt

ggaagcaccaactacaacccgtccctcaagagtcgagtcaccatatcagta

gacacgtccaagaaccagttctccctgaagctgagctctgtgaccgccgcggacacggct

gtgtattactgtgcg

>IGHV4-34*09

caggtgcagctgcaggagtcgggcccaggactggtgaagccttcacagaccctgtcc

ctcacctgcgctgtctatggtgggtccttcagtggttactactggagc

tggatccgccagcccccagggaagggactggagtggattggggaaatcaatcatagt

ggaagcaccaactacaacccgtccctcaagagtcgagttaccatatcagta

gacacgtctaagaaccagttctccctgaagctgagctctgtgactgccgcggacacggcc

gtgtattactgtgcgagaga

>IGHV4-34*10

caggtgcagctgcaggagtcgggcccaggactggtgaagccttcggagaccctgtcc

ctcacctgcgctgtctatggtgggtccttcagtggttactactggagc

tggatccgccagcccccagggaagggactggagtggattggggaaatcaatcatagt

ggaagcaccaactacaacccgtccctcaagagtcgaatcaccatgtcagta

gacacgtccaagaaccagttctacctgaagctgagctctgtgaccgccgcggacacggcc

gtgtattactgtgcgagata

>IGHV4-34*11

caggtgcagctacagcagtggggcgcaggactgttgaagccttcggagaccctgtcc

ctcacctgcgctgtctatggtgggtccgtcagtggttactactggagc

tggatccggcagcccccagggaaggggctggagtggattgggtatatctattatagt

gggagcaccaacaacaacccctccctcaagagtcgagccaccatatcagta

gacacgtccaagaaccagttctccctgaacctgagctctgtgaccgccgcggacacggcc

gtgtattgctgtgcgagaga

>IGHV4-34*12

caggtgcagctacagcagtggggcgcaggactgttgaagccttcggagaccctgtcc

ctcacctgcgctgtctatggtgggtccttcagtggttactactggagc

tggatccgccagcccccagggaaggggctggagtggattggggaaatcattcatagt

ggaagcaccaactacaacccgtccctcaagagtcgagtcaccatatcagta

gacacgtccaagaaccagttctccctgaagctgagctctgtgaccgccgcggacacggct

gtgtattactgtgcgaga

>IGHV4-34*13

tatggtgggtccttcagtggttactactggagc

tggatccgccagcccccagggaaggggctggagtggattggggaaatcaatcatagt

ggaagcaccaactacaacccctccctcaagagtcgagtcaccatatcagta

gacacgtccaagaaccagttctccctgaagctgagctctgtgaccgccgcggacacggct

gtgtattactgtgcgagagg

>IGHV4-38-2*01

caggtgcagctgcaggagtcgggcccaggactggtgaagccttcggagaccctgtcc

ctcacctgcgctgtctctggttactccatcagcagtggttactactggggc

tggatccggcagcccccagggaaggggctggagtggattgggagtatctatcatagt

gggagcacctactacaacccgtccctcaagagtcgagtcaccatatcagta

gacacgtccaagaaccagttctccctgaagctgagctctgtgaccgccgcagacacggcc

gtgtattactgtgcgaga

>IGHV4-38-2*02

caggtgcagctgcaggagtcgggcccaggactggtgaagccttcggagaccctgtcc

ctcacctgcactgtctctggttactccatcagcagtggttactactggggc

tggatccggcagcccccagggaaggggctggagtggattgggagtatctatcatagt

gggagcacctactacaacccgtccctcaagagtcgagtcaccatatcagta

gacacgtccaagaaccagttctccctgaagctgagctctgtgaccgccgcagacacggcc

gtgtattactgtgcgagaga

>IGHV4-39*01

cagctgcagctgcaggagtcgggcccaggactggtgaagccttcggagaccctgtcc

ctcacctgcactgtctctggtggctccatcagcagtagtagttactactggggc

tggatccgccagcccccagggaaggggctggagtggattgggagtatctattatagt

gggagcacctactacaacccgtccctcaagagtcgagtcaccatatccgta

gacacgtccaagaaccagttctccctgaagctgagctctgtgaccgccgcagacacggct

gtgtattactgtgcgagaca

>IGHV4-39*02

cagctgcagctgcaggagtcgggcccaggactggtgaagccttcggagaccctgtcc

ctcacctgcactgtctctggtggctccatcagcagtagtagttactactggggc

tggatccgccagcccccagggaaggggctggagtggattgggagtatctattatagt

gggagcacctactacaacccgtccctcaagagtcgagtcaccatatccgta

gacacgtccaagaaccacttctccctgaagctgagctctgtgaccgccgcagacacggct

gtgtattactgtgcgagaga

>IGHV4-39*03

cagctgcagctgcaggagtcgggcccaggactggtgaagccttcggagaccctgtcc

ctcacctgcactgtctctggtggctccatcagcagtagtagttactactggggc

tggatccgccagcccccagggaaggggctggagtggattgggagtatctattatagt

gggagcacctactacaacccgtccctcaagagtcgagtcaccatatccgta

gacacgtccaagaaccagttctccctgaagctgagctctgtgaccgccgcagacacggcc

gtgtattactg

>IGHV4-39*04

gctccatcagcagtagtagttactactggggc

tggatccgccagcccccagggaaggggctggagtggattgggagtatctattatagt

gggagcacctactacaacccgtccctcaagagtcgagtcaccatatccgta

gacacgtccaagaaccagttctccctgaagctgagctctgtgaccgccgcggacac

>IGHV4-39*05

cagctgcagctgcaggagtcgggcccaggactggtgaagccttcggagaccccgtcc

ctcacctgcactgtctctggtggctccatcagcagtagtagttactactggggc

tggatccgccagcccccagggaaggggctggagtggattgggagtatctattatagt

gggagcacctactacaacccgtccctcaagagtcgagtcaccatatccgta

gacacgtccaagaaccagttctccctgaagctgagctctgtgaccgccgcagacacggct

gtgtattactgtgcg

>IGHV4-39*06

cggctgcagctgcaggagtcgggcccaggactggtgaagccttcggagaccctgtcc

ctcacctgcactgtctctggtggctccatcagcagtagtagttactactggggc

tggatccgccagcccccagggaaggggctggagtggattgggagtatctattatagt

gggagcacctactacaacccgtccctcaagagtcgagtcaccatatcagta

gacacgtccaagaaccagttccccctgaagctgagctctgtgaccgccgcggacacggcc

gtgtattactgtgcgagaga

>IGHV4-39*07

cagctgcagctgcaggagtcgggcccaggactggtgaagccttcggagaccctgtcc

ctcacctgcactgtctctggtggctccatcagcagtagtagttactactggggc

tggatccgccagcccccagggaaggggctggagtggattgggagtatctattatagt

gggagcacctactacaacccgtccctcaagagtcgagtcaccatatcagta

gacacgtccaagaaccagttctccctgaagctgagctctgtgaccgccgcggacacggcc

gtgtattactgtgcgagaga

>IGHV4-4*01

caggtgcagctgcaggagtcgggcccaggactggtgaagcctccggggaccctgtcc

ctcacctgcgctgtctctggtggctccatcagcagtagtaactggtggagt

tgggtccgccagcccccagggaaggggctggagtggattggggaaatctatcatagt

gggagcaccaactacaacccgtccctcaagagtcgagtcaccatatcagta

gacaagtccaagaaccagttctccctgaagctgagctctgtgaccgccgcggacacggcc

gtgtattgctgtgcgagaga

>IGHV4-4*02

caggtgcagctgcaggagtcgggcccaggactggtgaagccttcggggaccctgtcc

ctcacctgcgctgtctctggtggctccatcagcagtagtaactggtggagt

tgggtccgccagcccccagggaaggggctggagtggattggggaaatctatcatagt

gggagcaccaactacaacccgtccctcaagagtcgagtcaccatatcagta

gacaagtccaagaaccagttctccctgaagctgagctctgtgaccgccgcggacacggcc

gtgtattactgtgcgagaga

>IGHV4-4*03

caggtgcagctgcaggagtcgggcccaggactggtgaagcctccggggaccctgtcc

ctcacctgcgctgtctctggtggctccatcagcagtagtaactggtggagt

tgggtccgccagcccccagggaaggggctggagtggattggggaaatctatcatagt

gggagcaccaactacaacccgtccctcaagagtcgagtcaccatatcagta

gacaagtccaagaaccagttctccctgaagctgagctctgtgaccgccgcggacacggcc

gtgtattactgtgcgagag

>IGHV4-4*04

caggtgcagctgcaggagtcgggcccaggactggtgaagcctccggggaccctgtcc

ctcacctgcgctatctctggtggctccatcagcagtagtaactggtggagt

tgggtccgccagcccccagggaaggggctggagtggattggggaaatctatcatagt

gggagcaccaactacaacccgtccctcaagagtcgagtcaccatatcagta

gacaagtccaagaaccagttctccctgaagctgagctctgtgaccgccgcggacacggcc

gtgtattactg

>IGHV4-4*05

caggtgcagctgcaggagttgggcccaggactggtgaagcctccggggaccctgtcc

ctcacctgcgctgtctctggtggctccatcagcagtagtaactggtggagt

tgggtccgccagcccccagggaaggggctggagtggattggggaaatctatcatagt

gggagcaccaactacaacccgtccctcaagagtcgagtcaccatatcagta

gacaagtccaagaaccagttctccctgaagctgagctctgtgaccgccgcggacacggcc

gtgtattactg

>IGHV4-4*06

tctggtggctccatcagcagtagtaactggtggagt

tgggtccgccagcccccagggannnggctggagtggattggggaaatctatcatagt

gggagcaccaactacaacccgtccctcaagagtcgagtcaccatgtcagta

gacacgtccaagaaccagttctccctgaagctgagctctgtgaccgccgcggacacggcc

gtgtattactgtgcgagaga

>IGHV4-4*07

caggtgcagctgcaggagtcgggcccaggactggtgaagccttcggagaccctgtcc

ctcacctgcactgtctctggtggctccatcagtagttactactggagc

tggatccggcagcccgccgggaagggactggagtggattgggcgtatctataccagt

gggagcaccaactacaacccctccctcaagagtcgagtcaccatgtcagta

gacacgtccaagaaccagttctccctgaagctgagctctgtgaccgccgcggacacggcc

gtgtattactgtgcgagaga

>IGHV4-4*08

caggtgcagctgcaggagtcgggcccaggactggtgaagccttcggagaccctgtcc

ctcacctgcactgtctctggtggctccatcagtagttactactggagc

tggatccggcagcccccagggaagggactggagtggattgggtatatctataccagt

gggagcaccaactacaacccctccctcaagagtcgagtcaccatatccgta

gacacgtccaagaaccagttctccctgaagctgagctctgtgaccgccgcagacacggcc

gtgtattactgtgcgagaga

>IGHV4-4*09

caggtgcagctgcaggagtcgggcccaggactggtgaagccttcggagaccctgtcc

ctcacctgcactgtctctggtggctccatcagtagttactactggagc

tggatccggcagcccccagggaagggactggagtggattgggtacatctataccagt

gggagcaccaactacaacccctccctcaagagtcgagtcaccatatcagta

gacacgtccaagaaccagttctccctgaagctgagctctgtgaccgccgcagacacggcc

gtgtattactgtgcgaga

>IGHV4-55*01

caggtgcagctgcaggagtcgggcccaggactggtgaagccttcggagaccctgtcc

ctcatctgcgctgtctctggtgactccatcagcagtggtaactggtgaatc

tgggtccgccagcccccagggaaggggctggagtggattggggaaatccatcatagt

gggagcacctactacaacccgtccctcaagagtcgaatcaccatgtccgta

gacacgtccaagaaccagttctacctgaagctgagctctgtgaccgccgcggacacggcc

gtgtattactgtgcgagata

>IGHV4-55*02

caggtgcagctgcaggagtcgggcccaggactggtgaagccttcggagaccctgtcc

ctcatctgcgctgtctctggtgactccatcagcagtggtaactggtgaatc

tgggtccgccagcccccagggaaggggctggagtggattggggaaatccatcatagt

gggagcacctactacaacccgtccctcaagagtcgaatcaccatgtcagta

gacacgtccaagaaccagttctacctgaagctgagctctgtgaccgccgcggacacggcc

gtgtattactgtgcgagata

>IGHV4-55*03

caggtgcagctgcaggagtcgggcccaggactggtgaagccttcggagaccctgtcc

ctcatctgcgctgtctctggtgactccatcagcagtggtaactggtgaatc

tgggtccgccagcccccagggaaggggctggagtggattggggaaatccatcatagt

gggagcacctactacaacccgtccctcaagagtcgaatcaccatgtcagta

gacacgtccaagaaccagttctccctgaagctgagctctgtgaccgccgcggacacggcc

gtgtattactg

>IGHV4-55*04

caggtgcagctgcaggagtcgggcccaggactggtgaagctttcggagaccctgtcc

ctcatctgcgctgtctctggtgactccatcagcagtggtaactggtgaatc

tgggtccgccagcccccagggaaggggctggagtggattggggaaatccatcatagt

gggagcacctactacaacccgtccctcaagagtcgaatcaccatgtcagta

gacacgtccaagaaccagttctacctgaagctgagctctgtgaccgccgcggacacggcc

gtgtattactg

>IGHV4-55*05

caggtgcagctgcaggagtcgggcccaggactggtgaagctttcggagaccctgtcc

ctcatctgcgctgtctctggtgactccatcagcagtggtaactggtgaatc

tgggtccgccagcccccagggaaggggctggagtggattggggaaatccatcatagt

gggagcacctactacaacccgtccctcaagagtcgaatcaccatgtccgta

gacacgtccaagaaccagttctacctgaagctgagctctgtgaccgccgcggacacggcc

gtgtattactg

>IGHV4-55*06

caggtgcagctgcaggagtcgggcccaggactggtgaagccttcggagaccctgtcc

ctcatctgcgctgtctctggtgactccatcagcagtggtaactggtgaatc

tgggtccgccagcccccagggaaggggctggagtggattggggaaatccatcatagt

gggagcacctactacaacccgtccctcaagagtcgaatcaccatgtccgta

gacacgtccaagaagcagttctacctgaagctgagctctgtgaccgctgcggacacggcc

gtgtattactg

>IGHV4-55*07

caggtgcagctgcaggagtcgggcccaggactggtgaagccttcggagaccctgtcc

ctcatctgcgctgtctctggtgactccatcagcagtggtaactggtgaatc

tgggtccgccagcccccagggaaggggctggagtggattggggaaatccatcatagt

gggagcacctactacaacccgtccctcaagagtcgaatcaccatgtccgta

gacacgtccaggaaccagttctccctgaagctgagctctgtgaccgccgcagacacggcc

gtgtattactg

>IGHV4-55*08

caggtgcagctgcaggagtcgggcccaggactggtgaagccttcggagaccctgtcc

ctcatctgcgctgtctctggtgactccatcagcagtggtaactggtgaatc

tgggtccgccagcccccagggaaggggctggagtggattggggaaatccatcatagt

gggagcacctactacaacccgtccctcaagagtcgaatcaccatgtcagta

gacacgtccaagaaccagttctacctgaagctgagctctgtgaccgccgcggacacggcc

gtgtattactgtgcgagaga

>IGHV4-55*09

caggtgcagctgcaggagtcgggcccaggactggtgaagccttcggagaccctgtcc

ctcatctgcgctgtctctggtgactccatcagcagtggtaactggtgaatc

tgggtccgccagcccccagggaaggggctggagtggattggggaaatccatcatagt

gggagcacctactacaacccgtccctcaagagtcgaatcaccatgtccgta

gacacgtccaagaaccagttctccctgaagctgagctctgtgaccgccgtggacacggcc

gtgtattactgtgcgagaaa

>IGHV4-59*01

caggtgcagctgcaggagtcgggcccaggactggtgaagccttcggagaccctgtcc

ctcacctgcactgtctctggtggctccatcagtagttactactggagc

tggatccggcagcccccagggaagggactggagtggattgggtatatctattacagt

gggagcaccaactacaacccctccctcaagagtcgagtcaccatatcagta

gacacgtccaagaaccagttctccctgaagctgagctctgtgaccgctgcggacacggcc

gtgtattactgtgcgagaga

>IGHV4-59*02

caggtgcagctgcaggagtcgggcccaggactggtgaagccttcggagaccctgtcc

ctcacctgcactgtctctggtggctccgtcagtagttactactggagc

tggatccggcagcccccagggaagggactggagtggattgggtatatctattacagt

gggagcaccaactacaacccctccctcaagagtcgagtcaccatatcagta

gacacgtccaagaaccagttctccctgaagctgagctctgtgaccgctgcggacacggcc

gtgtattactgtgcgagaga

>IGHV4-59*03

caggtgcagctgcaggagtcgggcccaggactggtgaagccttcggagaccctgtcc

ctcacctgcactgtctctggtggctccatcagtagttactactggagc

tggatccggcagcccccagggaagggactggagtggattgggtatatctattacagt

gggagcaccaactacaacccctccctcaagagtcgagtcaccatatcagta

gacacgtccaagaaccaattctccctgaagctgagctctgtgaccgctgcggacacggcc

gtgtattactgtgcg

>IGHV4-59*04

caggtgcagctgcaggagtcgggcccaggactggtgaagccttcggagaccctgtcc

ctcacctgcactgtctctggtggctccatcagtagttactactggagc

tggatccggcagcccccagggaagggactggagtggattgggtatatctattatagt

gggagcacctactacaacccgtccctcaagagtcgagtcaccatgtcagta

gacacgtccaagaaccagttctccctgaagctgagctctgtgaccgccgcagacacggct

gtgtattactgtgcg

>IGHV4-59*05

caggtgcagctgcaggagtcgggcccaggactggtgaagccttcggagaccctgtcc

ctcacctgcactgtctctggtggctccatcagtagttactactggagc

tggatccggcagccgccggggaagggactggagtggattgggcgtatctattatagt

gggagcacctactacaacccgtccctcaagagtcgagtcaccatatccgta

gacacgtccaagaaccagttctccctgaagctgagctctgtgaccgccgcagacacggct

gtgtattactgtgcg

>IGHV4-59*06

caggtgcagctgcaggagtcgggcccaggactggtgaagccttcggagaccctgtcc

ctcacctgcactgtcactggtggctccatcagtagttactactggagc

tggatccggcagcccgctgggaagggcctggagtggattgggtacatctattacagt

gggagcacctactacaacccgtccctcaagagtcgagttaccatatcagta

gacacgtctaagaaccagttctccctgaagctgagctctgtgactgccgcggacacggcc

gtgtattactgtgcg

>IGHV4-59*07

caggtgcagctgcaggagtcgggcccaggactggtgaagccttcggacaccctgtcc

ctcacctgcactgtctctggtggctccatcagtagttactactggagc

tggatccggcagcccccagggaagggactggagtggattgggtatatctattacagt

gggagcaccaactacaacccctccctcaagagtcgagtcaccatatcagta

gacacgtccaagaaccagttctccctgaagctgagctctgtgaccgctgcggacacggcc

gtgtattactgtgcgaga

>IGHV4-59*08

caggtgcagctgcaggagtcgggcccaggactggtgaagccttcggagaccctgtcc

ctcacctgcactgtctctggtggctccatcagtagttactactggagc

tggatccggcagcccccagggaagggactggagtggattgggtatatctattacagt

gggagcaccaactacaacccctccctcaagagtcgagtcaccatatcagta

gacacgtccaagaaccagttctccctgaagctgagctctgtgaccgccgcagacacggcc

gtgtattactgtgcgagaca

>IGHV4-59*09

tctggtggctccatcagtagttactactggagc

tggatccggcagcccccaggnannngactggagtggattgggtatatctattacagt

gggagcaccaactacaacccctccctcaagagtcgagtcaccatatcagta

gacacgtccaagaaccagttctccctgaagctgagctctgtgaccgctgcggacacggcc

gtgtattactgtgcgagagg

>IGHV4-59*10

caggtgcagctacagcagtggggcgcaggactgttgaagccttcggagaccctgtcc

ctcacctgcgctgtctatggtggctccatcagtagttactactggagc

tggatccggcagcccgccgggaaggggctggagtggattgggcgtatctataccagt

gggagcaccaactacaacccctccctcaagagtcgagtcaccatgtcagta

gacacgtccaagaaccagttctccctgaagctgagctctgtgaccgccgcggacacggcc

gtgtattactgtgcgagata

>IGHV4-59*11

caggtgcagctgcaggagtcgggcccaggactggtgaagccttcggagaccctgtcc

ctcacctgcactgtctctggtggctccatcagtagtcactactggagc

tggatccggcagcccccagggaagggactggagtggattgggtatatctattacagt

gggagcaccaactacaacccctccctcaagagtcgagtcaccatatcagta

gacacgtccaagaaccagttctccctgaagctgagctctgtgaccgctgcggacacggcc

gtgtattactgtgcgagaga

>IGHV4-59*12

caggtgcagctgcaggagtcgggcccaggactggtgaagccttcggagaccctgtcc

ctcacctgcactgtctctggtggctccatcagtagttactactggagc

tggatccggcagcccccagggaagggactggagtggattgggtatatctattacagt

gggagcaccaactacaacccctccctcaagagtcgagtcaccatatcagta

gacacgtccaagaaccagttctccctgaagctgagctctgtgaccgccgcggacacggcc

gtgtattactgtgcgagaga

>IGHV4-59*13

caggtgcagctgcaggagtcgggcccaggactggtgaagccttcggagaccctgtcc

ctcacctgcactgtctctggtggctccatcagtagttactactggagc

tggatccggcagcccccggggaagggactggagtggattgggtatatctattacagt

gggagcaccaactacaacccctccctcaagagtcgagtcaccatatcagta

gacacgtccaagaaccagttctccctgaagctgagctctgtgaccgctgcggacacggcc

gtgtattactgtgcgagaga

>IGHV4-61*01

caggtgcagctgcaggagtcgggcccaggactggtgaagccttcggagaccctgtcc

ctcacctgcactgtctctggtggctccgtcagcagtggtagttactactggagc

tggatccggcagcccccagggaagggactggagtggattgggtatatctattacagt

gggagcaccaactacaacccctccctcaagagtcgagtcaccatatcagta

gacacgtccaagaaccagttctccctgaagctgagctctgtgaccgctgcggacacggcc

gtgtattactgtgcgagaga

>IGHV4-61*02

caggtgcagctgcaggagtcgggcccaggactggtgaagccttcacagaccctgtcc

ctcacctgcactgtctctggtggctccatcagcagtggtagttactactggagc

tggatccggcagcccgccgggaagggactggagtggattgggcgtatctataccagt

gggagcaccaactacaacccctccctcaagagtcgagtcaccatatcagta

gacacgtccaagaaccagttctccctgaagctgagctctgtgaccgccgcagacacggcc

gtgtattactgtgcgagaga

>IGHV4-61*03

caggtgcagctgcaggagtcgggcccaggactggtgaagccttcggagaccctgtcc

ctcacctgcactgtctctggtggctccgtcagcagtggtagttactactggagc

tggatccggcagcccccagggaagggactggagtggattgggtatatctattacagt

gggagcaccaactacaacccctccctcaagagtcgagtcaccatatcagta

gacacgtccaagaaccacttctccctgaagctgagctctgtgaccgctgcggacacggcc

gtgtattactgtgcgagaga

>IGHV4-61*04

caggtgcagctgcaggagtcgggcccaggactggtgaagccttcggagaccctgtcc

ctcacctgcactgtctctggtggctccgtcagcagtggtagttactactggagc

tggatccggcagcccccagggaagggactggagtggattggatatatctattacagt

gggagcaccaactacaacccctccctcaagagtcgagtcaccatatcagta

gacacgtccaagaaccagttctccctgaagctgagctctgtgaccgctgacacggcc

gtgtattactg

>IGHV4-61*05

cagctgcagctgcaggagtcgggcccaggactggtgaagccttcggagaccctgtcc

ctcacctgcactgtctctggtggctccatcagcagtagtagttactactggggc

tggatccggcagcccccagggaagggactggagtggattgggtatatctattacagt

gggagcaccaactacaacccctccctcaagagtcgagtcaccatatcagta

gacaagtccaagaaccagttctccctgaagctgagctctgtgaccgccgcggacacggcc

gtgtattactgtgcgaga

>IGHV4-61*06

tctggtggctccgtcagcagtggtagttactactggagc

tggatccggcagcccccagggaagggactggagtggattgggtatatctattacagt

gggagcaccaactacaacccctccctcaagagtcgagtcaccatatcagta

gacacgtccaagaaccagttctccctgaagctgagctctgtgaccgccgcggacacggcc

gtgtattactgtgccagaga

>IGHV4-61*07

tctggtggctccgtcagcagtggtagttactactggagc

tggatccggcagcccccagggaagggactggagtggattgggtatatctattacagt

gggagcaccaactacaacccctccctcaagagtcgagtcaccatatcagta

gacacgtccaagaaccagttctccctgaagctgagctctgtgaccgctgcggacacggcc

gtgtattactgtgcgagaca

>IGHV4-61*08

caggtgcagctgcaggagtcgggcccaggactggtgaagccttcggagaccctgtcc

ctcacctgcactgtctctggtggctccgtcagcagtggtggttactactggagc

tggatccggcagcccccagggaagggactggagtggattgggtatatctattacagt

gggagcaccaactacaacccctccctcaagagtcgagtcaccatatcagta

gacacgtccaagaaccagttctccctgaagctgagctctgtgaccgctgcggacacggcc

gtgtattactgtgcgagaga

>IGHV4-61*09

caggtgcagctgcaggagtcgggcccaggattggtgaagccttcacagaccctgtcc

ctcacctgcactgtctctggtggctccatcagcagtggtagttactactggagc

tggatccggcagcccgccgggaagggactggagtggattgggcatatctataccagt

gggagcaccaactacaacccctccctcaagagtcgagtcaccatatcagta

gacacgtccaagaaccagttctccctgaagctgagctctgtgaccgccgcagacacggcc

gtgtattactgtgcgagaga

>IGHV4-61*10

caggtgcagctgcaggagtcgggcccaggactggtgaagccttcggagaccctgtcc

ctcacctgcactgtctctggtggctccgtcagcagtggtagttactactggagc

tggatccggcagcccgccgggaagggactggagtggattgggtatatctattacagt

gggagcaccaactacaacccctccctcaagagtcgagtcaccatatcagta

gacacgtccaagaaccagttctccctgaagctgagctctgtgaccgctgcggacacggcc

gtgtattactgtgcgagag

>IGHV4/OR15-8*01

caggtgcagctgcaggagtcgggcccaggactggtgaagccttcggagaccctgtcc

ctcacctgcgttgtctctggtggctccatcagcagtagtaactggtggagc

tgggtccgccagcccccagggaaggggctggagtggattggggaaatctatcatagt

gggagccccaactacaacccgtccctcaagagtcgagtcaccatatcagta

gacaagtccaagaaccagttctccctgaagctgagctctgtgaccgccgcggacacggcc

gtgtattactgtgcgagaga

>IGHV4/OR15-8*02

caggtgcagctgcaggagtcgggcccaggactggtgaagccttcggagaccctgtcc

ctcacctgcgttgtctctggtggctccatcagcagtagtaactggtggagc

tgggtccgccagcccccagggaaggggctggagtggattggggaaatctatcatagt

gggaaccccaactacaacccgtccctcaagagtcgagtcaccatatcaata

gacaagtccaagaaccaattctccctgaagctgagctctgtgaccgccgcggacacggcc

gtgtattactgtgcgagaga

>IGHV4/OR15-8*03

caggtgcagctgcaggagtcgggcccaggactggtgaagccttcggagaccctgtcc

ctcacctgcgttgtctctggtggctccatcagcagtagtaactggtggagc

tgggtccgccagcccccagggaaggggctggagtggattggggaaatctatcatagt

gggagccccaactacaacccatccctcaagagtcgagtcaccatatcagta

gacaagtccaagaaccagttctccctgaagctgagctctgtgaccgccgcggacacggcc

gtgtattactgtgcgagaga

>IGHV5-10-1*01

gaagtgcagctggtgcagtctggagcagaggtgaaaaagcccggggagtctctgagg

atctcctgtaagggttctggatacagctttaccagctactggatcagc

tgggtgcgccagatgcccgggaaaggcctggagtggatggggaggattgatcctagt

gactcttataccaactacagcccgtccttccaaggccacgtcaccatctcagct

gacaagtccatcagcactgcctacctgcagtggagcagcctgaaggcctcggacaccgcc

atgtattactgtgcgaga

>IGHV5-10-1*02

gaagtgcagctggtgcagtctggagcagaggtgaaaaagcccggggagtctctgagg

atctcctgtaagggttctggatacagctttaccagctactggatcagc

tgggtgcgccagatgcccgggaaaggcttggagtggatggggaggattgatcctagt

gactcttataccaactacagcccgtccttccaaggccacgtcaccatctcagct

gacaagtccatcagcactgcctacctgcagtggagcagcctgaaggctcggacaccgcc

atgtattactgtgcgagaca

>IGHV5-10-1*03

gaagtgcagctggtgcagtccggagcagaggtgaaaaagcccggggagtctctgagg

atctcctgtaagggttctggatacagctttaccagctactggatcagc

tgggtgcgccagatgcccgggaaaggcctggagtggatggggaggattgatcctagt

gactcttataccaactacagcccgtccttccaaggccacgtcaccatctcagct

gacaagtccatcagcactgcctacctgcagtggagcagcctgaaggcctcggacaccgcc

atgtattactgtgcgaga

>IGHV5-10-1*04

gaagtgcagctggtgcagtctggagcagaggtgaaaaagcccggggagtctctgagg

atctcctgtaagggttctggatacagctttaccagctactggatcagc

tgggtgcgccagatgcccgggaaaggcctggagtggatggggaggattgatcctagt

gactcttataccaactacagcccgtccttccaaggccaggtcaccatctcagct

gacaagtccatcagcactgcctacctgcagtggagcagcctgaaggcctcggacaccgcc

atgtattactgtgcgaga

>IGHV5-51*01

gaggtgcagctggtgcagtctggagcagaggtgaaaaagcccggggagtctctgaag

atctcctgtaagggttctggatacagctttaccagctactggatcggc

tgggtgcgccagatgcccgggaaaggcctggagtggatggggatcatctatcctggt

gactctgataccagatacagcccgtccttccaaggccaggtcaccatctcagcc

gacaagtccatcagcaccgcctacctgcagtggagcagcctgaaggcctcggacaccgcc

atgtattactgtgcgagaca

>IGHV5-51*02

gaggtgcagctggtgcagtctggagcagaggtgaaaaagcccggggagtctctgaag

atctcctgtaagggttctggatacagctttaccagctactggaccggc

tgggtgcgccagatgcccgggaaaggcttggagtggatggggatcatctatcctggt

gactctgataccagatacagcccgtccttccaaggccaggtcaccatctcagcc

gacaagtccatcagcaccgcctacctgcagtggagcagcctgaaggcctcggacaccgcc

atgtattactgtgcgagaca

>IGHV5-51*03

gaggtgcagctggtgcagtctggagcagaggtgaaaaagccgggggagtctctgaag

atctcctgtaagggttctggatacagctttaccagctactggatcggc

tgggtgcgccagatgcccgggaaaggcctggagtggatggggatcatctatcctggt

gactctgataccagatacagcccgtccttccaaggccaggtcaccatctcagcc

gacaagtccatcagcaccgcctacctgcagtggagcagcctgaaggcctcggacaccgcc

atgtattactgtgcgaga

>IGHV5-51*04

gaggtgcagctggtgcagtctggagcagaggtgaaaaagccgggggagtctctgaag

atctcctgtaagggttctggatacagctttaccagctactggatcggc

tgggtgcgccagatgcccgggaaaggcctggagtggatggggatcatctatcctggt

gactctgataccagatacagcccgtccttccaaggccaggtcaccatctcagcc

gacaagcccatcagcaccgcctacctgcagtggagcagcctgaaggcctcggacaccgcc

atgtattactgtgcgaga

>IGHV5-51*05

aaaagcccggggagtctctgaag

atctcctgtaagggttctggatacagctttaccagctactggatcggc

tgggtgcgccagatgcccaggaaaggcctggagtggatggggatcatctatcctggt

gactctgataccagatacagcccgtccttccaaggccaggtcaccatctcagcc

gacaagtccatcagcaccgcctacctgcagtggagcagcctgaaggcctcggacaccgcc

atg

>IGHV5-51*06

gaggtgcagctggtgcagtctggagcagaggtgaaaaagccgggggagtctctgaag

atctcctgtaagggttctggatacagctttaccagctactggatcggc

tgggtgcgccagatgcccgggaaaggcctggagtggatggggatcatctatcctggt

gactctgataccagatacagcccgtccttccaaggccaggttaccatctcagcc

gacaagtccatcagcaccgcctacctgcagtggagcagcctgaaggcctcggacaccgcc

atgtattactgtgcgaga

>IGHV5-51*07

gaggtgcagctggtgcagtctggagcagaggtgaaaaagcccggggagtctctgaag

atctcctgtaagggttctggatacagctttaccagctactggatcggc

tgggtgcaccagatgcccgggaaaggcctggagtggatggggatcatctatcctggt

gactctgataccagatacagcccgtccttccaaggccaggtcaccatctcagcc

gacaagtccatcagcaccgcctacctgcagtggagcagcctgaaggcctcggacaccgcc

atgtattactgtgcgagaca

>IGHV5-78*01

gaggtgcagctgttgcagtctgcagcagaggtgaaaagacccggggagtctctgagg

atctcctgtaagacttctggatacagctttaccagctactggatccac

tgggtgcgccagatgcccgggaaagaactggagtggatggggagcatctatcctggg

aactctgataccagatacagcccatccttccaaggccacgtcaccatctcagcc

gacagctccagcagcaccgcctacctgcagtggagcagcctgaaggcctcggacgccgcc

atgtattattgtgtgaga

>IGHV6-1*01

caggtacagctgcagcagtcaggtccaggactggtgaagccctcgcagaccctctca

ctcacctgtgccatctccggggacagtgtctctagcaacagtgctgcttggaac

tggatcaggcagtccccatcgagaggccttgagtggctgggaaggacatactacaggtcc

aagtggtataatgattatgcagtatctgtgaaaagtcgaataaccatcaaccca

gacacatccaagaaccagttctccctgcagctgaactctgtgactcccgaggacacggct

gtgtattactgtgcaagaga

>IGHV6-1*02

caggtacagctgcagcagtcaggtccgggactggtgaagccctcgcagaccctctca

ctcacctgtgccatctccggggacagtgtctctagcaacagtgctgcttggaac

tggatcaggcagtccccatcgagaggccttgagtggctgggaaggacatactacaggtcc

aagtggtataatgattatgcagtatctgtgaaaagtcgaataaccatcaaccca

gacacatccaagaaccagttctccctgcagctgaactctgtgactcccgaggacacggct

gtgtattactgtgcaagaga

>IGHV6-1*03

caggtacagctgcagcagtcaggtccaggactggtgaagccctcgcagaccctctca

ctcacctgtgccatctccggggacagtgtctctagcaacagtgctgcttggaac

tggatcaggcagtccccatcgagaggccttgagtggctgggaaggacatactacaggtcc

aagtggtataatgattatgcagtatctgtgaaaagttgaataaccatcaaccca

gacacatccaagaaccagttctccctgcagctgaactctgtgactcccgaggacacggct

gtgtattactgtgcaagaga

>IGHV7-34-1*01

ctgcagctggtgcagtctgggcctgaggtgaagaagcctggggcctcagtgaag

gtctcctataagtcttctggttacaccttcaccatctatggtatgaat

tgggtatgatagacccctggacagggctttgagtggatgtgatggatcatcacctac

actgggaacccaacgtatacccacggcttcacaggatggtttgtcttctccatg

gacacgtctgtcagcacggcgtgtcttcagatcagcagcctaaaggctgaggacacggcc

gagtattactgtgcgaagta

>IGHV7-34-1*02

ctgcagctggtgcagtctgggcctgaggtgaagaagcctggggcctcagtgaag

gtctcctataagtcttctggttacaccttcaccatctatggtatgaat

tgggtatgatagacccctggacagggctttgagtggatgtgatggatcatcacctac

aatgggaacccaacgtatacccacggcttcacaggatggtttgtcttctccatg

gacacgtctgtcagcacggcgtgtcttcagatcagcagcctaaaggctgaggacacggcc

gagtattactgtgcgaagta

>IGHV7-34-1*03

ctgcagctggtgcagtctgggcctgaggtgaagaagcgtggggcctcagtgaag

gtctcctataagtcttctggttacaccttcaccatctatggtatgaat

tgggtatgatagacccctggacagggctttgagtggatgtgatggatcatcacctac

actgggaacccaacgtatacccacggcttcacaggatggtttgtcttctccatg

gacacgtctgtcagcacggcgtgtcttcagatcagcagcctaaaggctgaggacacggcc

gagtattactgtgcgaagta

>IGHV7-4-1*01

caggtgcagctggtgcaatctgggtctgagttgaagaagcctggggcctcagtgaag

gtttcctgcaaggcttctggatacaccttcactagctatgctatgaat

tgggtgcgacaggcccctggacaagggcttgagtggatgggatggatcaacaccaac

actgggaacccaacgtatgcccagggcttcacaggacggtttgtcttctccttg

gacacctctgtcagcacggcatatctgcagatctgcagcctaaaggctgaggacactgcc

gtgtattactgtgcgagaga

>IGHV7-4-1*02

caggtgcagctggtgcaatctgggtctgagttgaagaagcctggggcctcagtgaag

gtttcctgcaaggcttctggatacaccttcactagctatgctatgaat

tgggtgcgacaggcccctggacaagggcttgagtggatgggatggatcaacaccaac

actgggaacccaacgtatgcccagggcttcacaggacggtttgtcttctccttg

gacacctctgtcagcacggcatatctgcagatcagcagcctaaaggctgaggacactgcc

gtgtattactgtgcgagaga

>IGHV7-4-1*03

caggtgcagctggtgcaatctgggtctgagttgaagaagcctggggcctcagtgaag

gtttcctgcaaggcttctggatacaccttcactagctatgctatgaat

tgggtgcgacaggcccctggacaagggcttgagtggatgggatggatcaacaccaac

actgggaacccaacgtatgcccagggcttcacaggacggtttgtcttctccttg

gacacctctgtcagcacggcatatctgcagatcagcacgctaaaggctgaggacactg

>IGHV7-4-1*04

caggtgcagctggtgcaatctgggtctgagttgaagaagcctggggcctcagtgaag

gtttcctgcaaggcttctggatacaccttcactagctatgctatgaat

tgggtgcgacaggcccctggacaagggcttgagtggatgggatggatcaacaccaac

actgggaacccaacgtatgcccagggcttcacaggacggtttgtcttctccttg

gacacctctgtcagcatggcatatctgcagatcagcagcctaaaggctgaggacactgcc

gtgtattactgtgcgagaga

>IGHV7-4-1*05

caggtgcagctggtgcaatctgggtctgagttgaagaagcctggggcctcagtgaag

gtttcctgcaaggcttctggatacaccttcactagctatgctatgaat

tgggtgcgacaggcccctggacaagggcttgagtggatgggatggatcaacaccaac

actgggaacccaacgtatgcccagggcttcacaggacggtttgtcttctccttg

gacacctctgtcagcatggcatatctgcagatcagcagcctaaaggctgaggacactgcc

gtgtgttactgtgcgagaga

>IGHV7-40*03

ttttcaatagaaaagtcaaataatctaagtgtcaatcagtggatgattagataaaat

atgatatatgtaaatcatggaatactatgcagccagtatggtatgaat

tcagtgtgaccagcccctggacaagggcttgagtggatgggatggatcatcacctac

actgggaacccaacatataccaacggcttcacaggacggtttctattctccatg

gacacctctgtcagcatggcgtatctgcagatcagcagcctaaaggctgaggacacggcc

gtgtatgactgtatgagaga

>IGHV7-81*01

caggtgcagctggtgcagtctggccatgaggtgaagcagcctggggcctcagtgaag

gtctcctgcaaggcttctggttacagtttcaccacctatggtatgaat

tgggtgccacaggcccctggacaagggcttgagtggatgggatggttcaacacctac

actgggaacccaacatatgcccagggcttcacaggacggtttgtcttctccatg

gacacctctgccagcacagcatacctgcagatcagcagcctaaaggctgaggacatggcc

atgtattactgtgcgagata

>IGHV8-51-1*01

gaggcccagcttacagagtctgggggagacttggtacactgagaggggcccctgagg

ctctcctgtgcagcctcttggttcaccttcagtatctatgagattcac

tgggtttgccaggcctcagggaaggggctggaatgggttgcagttatatggcgtagt

gaaagtcatcaatacaatgcagactatgttaggggcagactcaccacttccaga

gacaacaccaagtacatgctgtacatgcaaatgaacagcctgagaacccagaacatggca

gcatttaactgtgcaggaaa

>IGHV8-51-1*02

gaggcccagcttacagagtctgggggagacttggtacacttagaggggcccctgagg

ctctcctgtgcagcctcttggttcaccttcagtatctatgagattcac

tgggtttgccaggcctcagggaaggggctggaatgggttgcagttatatggcgtggt

gaaagtcatcaatacaatgcagactatgttaggggcagactcaccacttccaga

gacaacaccaagtacatgctgtacatgcaaatgatcagcctgagaacccagaacatggca

gcatttaactgtgcaggaaa

>IGHV8-51-1*03

gaggcccagcttacagagtctgggggagacttggtacactgagaggggcccctgagg

ctctcctgtgcagcctcttggttcaccttcagtatctatgagattcac

tgggtttgccaggcctcagggaaggggctggaatgggttgcagttatatggcgtggt

gaaagtcatcaatacaatgcagactatgttaggggcagactcaccacttccaga

gacaacaccaagtacatgctgtacatgcaaatgaacagcctgagaacccagaacatggca

gcatttaactgtgcaggaaa

>IGHD1-1*01

ggtacaactggaacgac

>IGHD1-14*01

ggtataaccggaaccac

>IGHD1-20*01

ggtataactggaacgac

>IGHD1-26*01

ggtatagtgggagctactac

>IGHD1-7*01

ggtataactggaactac

>IGHD1/OR15-1a*01

ggtataactggaacaac

>IGHD1/OR15-1b*01

ggtataactggaacaac

>IGHD2-15*01

aggatattgtagtggtggtagctgctactcc

>IGHD2-2*01

aggatattgtagtagtaccagctgctatgcc

>IGHD2-2*02

aggatattgtagtagtaccagctgctatacc

>IGHD2-2*03

tggatattgtagtagtaccagctgctatgcc

>IGHD2-21*01

agcatattgtggtggtgattgctattcc

>IGHD2-21*02

agcatattgtggtggtgactgctattcc

>IGHD2-8*01

aggatattgtactaatggtgtatgctatacc

>IGHD2-8*02

aggatattgtactggtggtgtatgctatacc

>IGHD2/OR15-2a*01

agaatattgtaatagtactactttctatgcc

>IGHD2/OR15-2b*01

agaatattgtaatagtactactttctatgcc

>IGHD3-10*01

gtattactatggttcggggagttattataac

>IGHD3-10*02

gtattactatgttcggggagttattataac

>IGHD3-16*01

gtattatgattacgtttgggggagttatgcttatacc

>IGHD3-16*02

gtattatgattacgtttgggggagttatcgttatacc

>IGHD3-22*01

gtattactatgatagtagtggttattactac

>IGHD3-3*01

gtattacgatttttggagtggttattatacc

>IGHD3-3*02

gtattagcatttttggagtggttattatacc

>IGHD3-9*01

gtattacgatattttgactggttattataac

>IGHD3/OR15-3a*01

gtattatgatttttggactggttattatacc

>IGHD3/OR15-3b*01

gtattatgatttttggactggttattatacc

>IGHD4-11*01

tgactacagtaactac

>IGHD4-17*01

tgactacggtgactac

>IGHD4-23*01

tgactacggtggtaactcc

>IGHD4-4*01

tgactacagtaactac

>IGHD4/OR15-4a*01

tgactatggtgctaactac

>IGHD4/OR15-4b*01

tgactatggtgctaactac

>IGHD5-12*01

gtggatatagtggctacgattac

>IGHD5-18*01

gtggatacagctatggttac

>IGHD5-24*01

gtagagatggctacaattac

>IGHD5-5*01

gtggatacagctatggttac

>IGHD5/OR15-5a*01

gtggatatagtgtctacgattac

>IGHD5/OR15-5b*01

gtggatatagtgtctacgattac

>IGHD6-13*01

gggtatagcagcagctggtac

>IGHD6-19*01

gggtatagcagtggctggtac

>IGHD6-25*01

gggtatagcagcggctac

>IGHD6-6*01

gagtatagcagctcgtcc

>IGHD7-27*01

ctaactgggga

>IGHD1-1*01

ggtacaactggaacgac

>IGHD1-14*01

ggtataaccggaaccac

>IGHD1-20*01

ggtataactggaacgac

>IGHD1-26*01

ggtatagtgggagctactac

>IGHD1-7*01

ggtataactggaactac

>IGHD1/OR15-1a*01

ggtataactggaacaac

>IGHD1/OR15-1b*01

ggtataactggaacaac

>IGHD2-15*01

aggatattgtagtggtggtagctgctactcc

>IGHD2-2*01

aggatattgtagtagtaccagctgctatgcc

>IGHD2-2*02

aggatattgtagtagtaccagctgctatacc

>IGHD2-2*03

tggatattgtagtagtaccagctgctatgcc

>IGHD2-21*01

agcatattgtggtggtgattgctattcc

>IGHD2-21*02

agcatattgtggtggtgactgctattcc

>IGHD2-8*01

aggatattgtactaatggtgtatgctatacc

>IGHD2-8*02

aggatattgtactggtggtgtatgctatacc

>IGHD2/OR15-2a*01

agaatattgtaatagtactactttctatgcc

>IGHD2/OR15-2b*01

agaatattgtaatagtactactttctatgcc

>IGHD3-10*01

gtattactatggttcggggagttattataac

>IGHD3-10*02

gtattactatgttcggggagttattataac

>IGHD3-16*01

gtattatgattacgtttgggggagttatgcttatacc

>IGHD3-16*02

gtattatgattacgtttgggggagttatcgttatacc

>IGHD3-22*01

gtattactatgatagtagtggttattactac

>IGHD3-3*01

gtattacgatttttggagtggttattatacc

>IGHD3-3*02

gtattagcatttttggagtggttattatacc

>IGHD3-9*01

gtattacgatattttgactggttattataac

>IGHD3/OR15-3a*01

gtattatgatttttggactggttattatacc

>IGHD3/OR15-3b*01

gtattatgatttttggactggttattatacc

>IGHD4-11*01

tgactacagtaactac

>IGHD4-17*01

tgactacggtgactac

>IGHD4-23*01

tgactacggtggtaactcc

>IGHD4-4*01

tgactacagtaactac

>IGHD4/OR15-4a*01

tgactatggtgctaactac

>IGHD4/OR15-4b*01

tgactatggtgctaactac

>IGHD5-12*01

gtggatatagtggctacgattac

>IGHD5-18*01

gtggatacagctatggttac

>IGHD5-24*01

gtagagatggctacaattac

>IGHD5-5*01

gtggatacagctatggttac

>IGHD5/OR15-5a*01

gtggatatagtgtctacgattac

>IGHD5/OR15-5b*01

gtggatatagtgtctacgattac

>IGHD6-13*01

gggtatagcagcagctggtac

>IGHD6-19*01

gggtatagcagtggctggtac

>IGHD6-25*01

gggtatagcagcggctac

>IGHD6-6*01

gagtatagcagctcgtcc

>IGHD7-27*01

ctaactgggga

>IGHJ1*01

gctgaatacttccagcactggggccagggcaccctggtcaccgtctcctcag

>IGHJ2*01

ctactggtacttcgatctctggggccgtggcaccctggtcactgtctcctcag

>IGHJ3*01

tgatgcttttgatgtctggggccaagggacaatggtcaccgtctcttcag

>IGHJ3*02

tgatgcttttgatatctggggccaagggacaatggtcaccgtctcttcag

>IGHJ4*01

actactttgactactggggccaaggaaccctggtcaccgtctcctcag

>IGHJ4*02

actactttgactactggggccagggaaccctggtcaccgtctcctcag

>IGHJ4*03

gctactttgactactggggccaagggaccctggtcaccgtctcctcag

>IGHJ5*01

acaactggttcgactcctggggccaaggaaccctggtcaccgtctcctcag

>IGHJ5*02

acaactggttcgacccctggggccagggaaccctggtcaccgtctcctcag

>IGHJ6*01

attactactactactacggtatggacgtctgggggcaagggaccacggtcaccgtctcctcag

>IGHJ6*02

attactactactactacggtatggacgtctggggccaagggaccacggtcaccgtctcctca

>IGHJ6*03

attactactactactactacatggacgtctggggcaaagggaccacggtcaccgtctcctca

>IGHJ6*04

attactactactactacggtatggacgtctggggcaaagggaccacggtcaccgtctcctcag
