## Supplementary data 2 for "Computational inference, validation, and analysis of 5’UTR-leader sequences of alleles of immunoglobulin heavy chain variable genes"

**Supplementary Data 2.** Database of upstream regions of IGHV genes, as FASTA entries, inferred from transcriptome data. Added to the list are upstream regions of two poorly expressed genes that were not inferred in the present study but defined in a separate study (23).

>IGHV1-18*01-A|2

ACCCAAAAACCACACCCCTCCTTGGGAGAATCCCCTAGATCACAGCTCCTCACCATGGACTGGACCTGGAGCATCCTTTTCTTGCTGGCAGCAGCAACAGGTGCCCACTCC

>IGHV1-18*01-B|94

ACCCAAAAACCACACCCCTCCTTGGGAGAATCCCCTAGATCACAGCTCCTCACCATGGACTGGACCTGGAGCATCCTTTTCTTGGTGGCAGCAGCAACAGGTGCCCACTCC

>IGHV1-18*04-A|31

ACCCAAAAACCACACCCCTCCTTGGGAGAATCCCCTAGATCACAGCTCCTCACCATGGACTGGACCTGGAGCATCCTTTTCTTGGTGGCAGCAGCAACAGGTGCCCACTCC

>IGHV1-24*01-A|2

ATCACACAACAGCCACATCCCTCCCCTACAGAAGCCCCCAGAGCACAGCACCTCACCATGGACTGCACCTGGAGGATCCTCTTCTTGGTGGCAGCAGCTACAGGCACCCACGCC

>IGHV1-24*01-B|30

ATCACACAACAGCCACATCCCTCCCCTACAGAAGCCCCCAGAGTGCAGCACCTCACCATGGACTGCACCTGGAGGATCCTCTTCTTGGTGGCAGCAGCTACAGGCACCCACGCC

>IGHV1-24*01-C|91

ATCACACAACAGCCACATCCCTCCCCTACAGAAGCCCCCAGAGCGCAGCACCTCACCATGGACTGCACCTGGAGGATCCTCTTCTTGGTGGCAGCAGCTACAGGCACCCACGCC

>IGHV1-2*02-A|68

AGCATCACCCAGCAACCACATCTGTCCTCTAGAGAATCCCCTGAGAGCTCCGTTCCTCACCATGGACTGGACCTGGAGGATCCTCTTCTTGGTGGCAGCAGCCACAGGAGCCCACTCC

>IGHV1-2*04-A|64

AGCATCACCCAGCAACCACATCTGTCCTCTAGAGAATCCCCTGAGAGCTCCGTTCCTCACCATGGACTGGACCTGGAGGATCCTCTTCTTGGTGGCAGCAGCCACAGGAGCCCACTCC

>IGHV1-2*05-A|1

AGCATCACCCAGCAACCACATCTGTCCTCTAGAGAATCCCCTGAGAGCTCCGTTCCTCACCATGGACTGGACCTGGAGGATCCTCTTCTTGGTGGCAGCAGCCACAGGAGCCCACTCC

>IGHV1-2*06-A|19

AGCATCACCCAGCAACCACATCTGTCCTCTAGAGAATCCCCTGAGAGCTCCGTTCCTCACCATGGACTGGACCTGGAGGATCCTCTTCTTGGTGGCAGCAGCCACAGGAGCCCACTCC

>IGHV1-3*01-A|1

AACCACATCCCTCCTCAGAAGCCCCCAGAGCACAACGCCTCACCATGGACTGGACCTGGAGGATCCTCTTTTTGGTGGCAGCAGCCACAGGTGTCCACTCC

>IGHV1-3*01-B|9

AACCACATCCCTCCTCAGAAGCCCCCAGAGCACAACTCCTCACCATGGACTGGACCTGGAGGATCCTCTTTTTGGTGGCAGCAGCCACAGGTGCCCACTCC

>IGHV1-3*01-C|26

AACCACATCCCTCCTCAGAAGCCCCCAGAGCACAACTCCTTACCATGGACTGGACCTGGAGGATCCTCTTTTTGGTGGCAGCAGCCACAGGTGCCCACTCC

>IGHV1-3*01-D|57

AACCACATCCCTCCTCAGAAGCCCCCAGAGCACAACGCCTCACCATGGACTGGACCTGGAGGATCCTCTTTTTGGTGGCAGCAGCCACAGGTGCCCACTCC

>IGHV1-3*01_S6816-A|2

AACCACATCCCTCCTCAGAAGCCCCCAGAGCACAACGCCTCACCATGGACTGGACCTGGAGGATCCTCTTTTTGGTGGCAGCAGCCACAGGTGCCCACTCC

>IGHV1-46*01-A|90

ATCATCCAACAACCACATCCCTTCTCTACAGAAGCCTCTGAGAGGAAAGTTCTTCACCATGGACTGGACCTGGAGGGTCTTCTGCTTGCTGGCTGTAGCTCCAGGTGCTCACTCC

>IGHV1-46*03-A|49

ATCATCCAACAACCACATCCCTTCTCTACAGAAGCCTCTGAGAGGAAAGTTCTTCACCATGGACTGGACCTGGAGGGTCTTCTGCTTGCTGGCTGTAGCTCCAGGTGCTCACTCC

>IGHV1-46*04-A|2

ATCATCCAACAACCACATCCCTTCTCTACAGAAGCCTCTGAGAGGAAAGTTCTTCACCATGGACTGGACCTGGAGGGTCTTCTGCTTGCTGGCTGTAGCTCCAGGTGCTCACTCC

>IGHV1-58*01-A|61

ATCATCCAGAAACCACATCCCTCCGCTAGAGAAGCCCCTGACGGCACAGTTCCTCACTATGGACTGGATTTGGAGGATCCTCTTCTTGGTGGGAGCAGCGACAGGTGCCCACTCC

>IGHV1-58*02-A|23

ATCATCCAGAAACCACATCCCTCCGCTAGAGAAGCCCCTGACGGCACAGTTCCTCACTATGGACTGGATTTGGAGGATCCTCTTCTTGGTGGGAGCAGCGACAGGTGCCCACTCC

>IGHV1-58*02-B|16

ATCATCCAGAAACCACATCCCTCCGCTAGAGAAGCCCCTGACGGCACAGTTCCTCACTATGGACTGGATTTGGAGGGTCCTCTTCTTGGTGGGAGCAGCGACAGGTGCCCACTCC

>IGHV1-69-2*01-A|1

ATCACACAACAGCCACATCCCTCCCCTACAGAAGCCCCCAGAGCACAGCACCTCACCATGGACTGCACCTGGAGGATCCTCCTCTTGGTGGCAGCAGCTACAGGCACCCACGCC

>IGHV1-69-2*01-B|29

ATCACACAACAGCCACATCCCTCCCCTACAGAAGCCCCCAGAGAGCAGCACCTCACCATGGACTGCACCTGGAGGATCCTCCTCTTGGTGGCAGCAGCTACAGGCACCCACGCC

>IGHV1-69*01-A|76

ATCACATAACAACCACATTCCTCCTCTAAAGAAGCCCCTGGGAGCACAGCTCATCACCATGGACTGGACCTGGAGGTTCCTCTTTGTGGTGGCAGCAGCTACAGGTGTCCAGTCC

>IGHV1-69*01-B|1

ATCACATAACAACCACATTCCTCCTCTGAAGAAGCCCCTGGGAGCACAGCTCATCACCATGGACTGGACCTGGAGGTTCCTCTTTGTGGTGGCAGCAGCTACAGGTGTCCAGTCC

>IGHV1-69*01_S8909-A|1

ATCACATAACAACCACATTCCTCCTCTAAAGAAGCCCCTGGGAGCACAGCTCATCACCATGGACTGGACCTGGAGGTTCCTCTTTGTGGTGGCAGCAGCTACAGGTGTCCAGTCC

>IGHV1-69*02-A|45

TCACATAACAACCAGATTCCTCCTCTAAAGAAGCCCCTGGGAGCACAGCTCATCACCATGGACTGGACCTGGAGGTTCCTCTTTGTGGTGGCAGCAGCTACAGGTGTCCAGTCC

>IGHV1-69*04-A|24

ATCACATAACAACCAGATTCCTCCTCTAAAGAAGCCCCTGGGAGCACAGCTCATCACCATGGACTGGACCTGGAGGTTCCTCTTTGTGGTGGCAGCAGCTACAGGTGTCCAGTCC

>IGHV1-69*04-B|2

ATCACATAACAACCACATTCCTCCTCTGAAGAAGCCCCTGGGAGCACAGCTCATCACCATGGACTGGACCTGGAGGTTCCTCTTTGTGGTGGCAGCAGCTACAGGTGTCCAGTCC

>IGHV1-69*04_S3852-A|2

ATCACATAACAACCAGATTCCTCCTCTAAAGAAGCCCCTGGGAGCACAGCTCATCACCATGGACTGGACCTGGAGGTTCCTCTTTGTGGTGGCAGCAGCTACAGGTGTCCAGTCC

>IGHV1-69*06-A|40

ATCACATAACAACCACATTCCTCCTCTGAAGAAGCCCCTGGGAGCACAGCTCATCACCATGGACTGGACCTGGAGGTTCCTCTTTGTGGTGGCAGCAGCTACAGGTGTCCAGTCC

>IGHV1-69*06-B|2

ATCACATAACAACCACATTCCTCCTCTAAAGAAGCCCCTGGGAGCACAGCTCATCACCATGGACTGGACCTGGAGGTTCCTCTTTGTGGTGGCAGCAGCTACAGGTGTCCAGTCC

>IGHV1-69*06_S0471-A|1

ATCACATAACAACCACATTCCTCCTCTGAAGAAGCCCCTGGGAGCACAGCTCATCACCATGGACTGGACCTGGAGGTTCCTCTTTGTGGTGGCAGCAGCTACAGGTGTCCAGTCC

>IGHV1-69*09-A|2

ACATAACAACCACATTCCTCCTCTAAAGAAGCCCCTGGGAGCACAGCTCATCACCATGGACTGGACCTGGAGGTTCCTCTTTGTGGTGGCAGCAGCTACAGGTGTCCAGTCC

>IGHV1-69*09-B|1

ATCACATAACAACCAGATTCCTCCTCTAAAGAAGCCCCTGGGAGCACAGCTCATCACCATGGACTGGACCTGGAGGTTCCTCTTTGTGGTGGCAGCAGCTACAGGTGTCCAGTCC

>IGHV1-69*09-C|5

ATCACATAACAACCACATTCCTCCTCTGAAGAAGCCCCTGGGAGCACAGCTCATCACCATGGACTGGACCTGGAGGTTCCTCTTTGTGGTGGCAGCAGCTACAGGTGTCCAGTCC

>IGHV1-69*10-A|2

ATCACATAACAACCAGATTCCTCCTCTAAAGAAGCCCCTGGGAGCACAGCTCATCACCATGGACTGGACCTGGAGGTTCCTCTTTGTGGTGGCAGCAGCTACAGGTGTCCAGTCC

>IGHV1-69*12-A|3

ACATAACAACCAGATTCCTCCTCTAAAGAAGCCCCTGGGAGCACAGCTCATCACCATGGACTGGACCTGGAGGTTCCTCTTTGTGGTGGCAGCAGCTACAGGTGTCCAGTCC

>IGHV1-69*17-A|2

ATCACATAACAACCACATTCCTCCTCTGAAGAAGCCCCTGGGAGCACAGCTCATCACCATGGACTGGACCTGGAGGTTCCTCTTTGTGGTGGCAGCAGCTACAGGTGTCCAGTCC

>IGHV1-8*01-A|77

ACTCAACAACCACATCTGTCCTCTAGAGAAAACCCTGTGAGCACAGCTCCTCACCATGGACTGGACCTGGAGGATCCTCTTCTTGGTGGCAGCAGCTACAAGTGCCCACTCC

>IGHV1-8*03-A|2

ACTCAACAACCACATCTGTCCTCTAGAGAAAACCCTGTGAGCACAGCTCCTCACCATGGACTGGACCTGGAGGATCCTCTTCTTGGTGGCAGCAGCTACAAGTGCCCACTCC

>IGHV2-26*01-A|98

ACTCCTGTGCCCCACCATGGACACACTTTGCTACACACTCCTGCTGCTGACCACCCCTTCCTGGGTCTTGTCC

>IGHV2-26*02_S3803-A|1

ACTCCTGTGCCCCACCATGGACACGCTTTGCTACACACTCCTGCTGCTGACCACCCCTTCCTGGGTCTTGTCC

>IGHV2-5*01-A|57

ACTCCTGTGCCCCACCATGGACACACTTTGCTCCACGCTCCTGCTGCTGACCATCCCTTCATGGGTCTTGTCC

>IGHV2-5*02-A|82

ACTCCTGTGCCCCACCATGGACACACTTTGCTCCACGCTCCTGCTGCTGACCATCCCTTCATGGGTCTTGTCC

>IGHV2-70*01-A|63

AATCCTGCTCTCCACCATGGACATACTTTGTTCCACGCTCCTGCTACTGACTGTCCCGTCCTGGGTCTTATCC

>IGHV2-70*04-A|26

AATCCTGCTCTCCACCATGGACATACTTTGTTCCACGCTCCTGCTACTGACTGTCCCGTCCTGGGTCTTATCC

>IGHV2-70*04_S5392-A|1

AATCCTGCTCTCCACCATGGACATACTTTGTTCCACGCTCCTGCTACTGACTGTCCCGTCCTGGGTCTTATCC

>IGHV2-70*15-A|6

AATCCTGCTCTCCACCATGGACATACTTTGTTCCACGCTCCTGCTACTGACTGTCCCGTCCTGGGTCTTATCC

>IGHV2-70*15-B|36

AATCCTGCTCCCCACCATGGACATACTTTGTTCCACGCTCCTGCTACTGACTGTCCCGTCCTGGGTCTTATCC

>IGHV3-11*01-A|76

AGCTCTGGGAGAAGAGCCCCAGCCCCAGAATTCCCAGGAGTTTCCATTCGGTGATCAGCACTGAACACAGAGGACTCACCATGGAGTTTGGGCTGAGCTGGGTTTTCCTTGTTGCTATTATAAAAGGTGTCCAGTGT

>IGHV3-11*04-A|2

AGCTCTGGGAGAAGAGCCCCAGCCCCAGAATTCCCAGGAGTTTCCATTCGGTGATCAGCACTGAACACAGAGGACTCACCATGGAGTTTGGGCTGAGCTGGGTTTTCCTTGTTGCTATTATAAAAGGTGTCCAGTGT

>IGHV3-11*05-A|22

AGCTCTGGGAGAAGAGCCCCAGCCCCAGAATTCCCAGGAGTTTCCATTCGGTGATCAGCACTGAACACAGAGGACTCACCATGGAGTTTGGGCTGAGCTGGGTTTTCCTTGTTGCTATTATAAAAGGTGTCCAGTGT

>IGHV3-11*06-A|47

AGCTCTGGGAGAAGAGCCCCAGCCCCAGAATTCCCAGGAGTTTCCATTCGGTGATCAGCACTGAACACAGAGGACTCACCATGGAGTTTGGGCTGAGCTGGGTTTTCCTTGTTGCTATTATAAAAGGTGTCCAGTGT

>IGHV3-13*01-A|61

AGCTCTGGGAGTGGAGCCCCAGCCTTGGGATTCCCAAGTGTTTGTATTCAGTGATCAGGACTGAACACACAGGACTCACCATGGAGTTGGGGCTGAGCTGGGTTTTCCTTGTTGCTATATTAGAAGGTGTCCAGTGT

>IGHV3-13*01_S3164-A|8

CTCTGGGAGTGGAGCCCCAGCCTTGGGATTCCCAAGTGTTTGTATTCAGTGATCAGGACTGAACACACAGGACTCACCATGGAGTTGGGGCTGAGCTGGGTTTTCCTTGTTGCTATATTAGAAGGTGTCCAGTGT

>IGHV3-13*04-A|19

AGCTCTGGGAGTGGAGCCCCAGCCTTGGGATTCCCAAGTGTTTGTATTCAGTGATCAGGACTGAACACACAGGACTCACCATGGAGTTGGGGCTGAGCTGGGTTTTCCTTGTTGCTATATTAGAAGGTGTCCAGTGT

>IGHV3-13*05-A|22

AGCTCTGGGAGTGGAGCCCCAGCCTTGGGATTCCCAAGTGTTTGTATTCAGTGATCAGGACTGAACACACAGGACTCACCATGGAGTTGGGGCTGAGCTGGGTTTTCCTTGTTGCTATATTAGAAGGTGTCCAGTGT

>IGHV3-15*01-A|96

AGCTCTGGGAGAGGAGCCCCAGCCTTGGGATTCCCAAGTGTTTTCATTCAGTGATCAGGACTGAACACAGAGGACTCACCATGGAGTTTGGGCTGAGCTGGATTTTCCTTGCTGCTATTTTAAAAGGTGTCCAGTGT

>IGHV3-15*07-A|25

AGCTCTGGGAGAGGAGCCCCAGCCTTGGGATTCCCAAGTGTTTTCATTCAGTGATCAGGACTGAACACAGAGGACTCACCATGGAGTTTGGGCTGAGCTGGATTTTCCTTGCTGCTATTTTAAAAGGTGTCCAGTGT

>IGHV3-20*01-A|53

AGCTCTGGGAGAGGAGCCCCAGCCCTGAGATTCCCACGTGTTTCCATTCAGTGATCAGCACTGAACACAGAGGACTCGCCATGGAGTTTGGGCTGAGCTGGGTTTTCCTTGTTGCTATTTTAAAAGGTGTCCAGTGT

>IGHV3-20*04-A|30

AGCTCTGGGAGAGGAGCCCCAGCCCTGAGATTCCCACGTGTTTCCATTCAGTGATCAGCACTGAACACAGAGGACTCGCCATGGAGTTTGGGCTGAGCTGGGTTTTCCTTGTTGCTATTTTAAAAGGTGTCCAGTGT

>IGHV3-21*01-A|95

AGCTCTGAGAGAGGAGCCTTAGCCCTGGATTCCAAGGCCTATCCACTTGGTGATCAGCACTGAGCACCGAGGATTCACCATGGAACTGGGGCTCCGCTGGGTTTTCCTTGTTGCTATTTTAGAAGGTGTCCAGTGT

>IGHV3-21*01-B|15

AGCTCTGAGAGAGGAGCCTTAGCCCTGGATTCCAAGGCCTATCCACTTGGTGATCAGCACGGAGCACCGAGGATTCACCATGGAACTGGGGCTCCGCTGGGTTTTCCTTGTTGCTATTTTAGAAGGTGTCCAGTGT

>IGHV3-21*01_S4935-A|1

AGCTCTGAGAGAGGAGCCTTAGCCCTGGATTCCAAGGCCTATCCACTTGGTGATCAGCACTGAGCACCGAGGATTCACCATGGAACTGGGGCTCCGCTGGGTTTTCCTTGTTGCTATTTTAGAAGGTGTCCAGTGT

>IGHV3-21*01_S5913-A|1

AGCTCTGAGAGAGGAGCCTTAGCCCTGGATTCCAAGGCCTATCCACTTGGTGATCAGCACTGAGCACCGAGGATTCACCATGGAACTGGGGCTCCGCTGGGTTTTCCTTGTTGCTATTTTAGAAGGTGTCCAGTGT

>IGHV3-23*01-A|97

AGCTCTGAGAGAGGAGCCCAGCCCTGGGATTTTCAGGTGTTTTCATTTGGTGATCAGGACTGAACAGAGAGAACTCACCATGGAGTTTGGGCTGAGCTGGCTTTTTCTTGTGGCTATTTTAAAAGGTGTCCAGTGT

>IGHV3-23*04-A|20

AGCTCTGAGAGAGGAGCCCAGCCCTGGGATTTTCAGGTGTTTTCATTTGGTGATCAGGACTGAACAGAGAGAACTCACCATGGAGTTTGGGCTGAGCTGGCTTTTTCTTGTGGCTATTTTAAAAGGTGTCCAGTGT

>IGHV3-30-3*01-A|69

AGCTCTGGGAGACGAGCCCAGCACTGGAAGTCGCCGGTGTTTCCATTCGGTGATCATCACTGAACACAGAGGACTCACCATGGAGTTTGGGCTGAGCTGGGTTTTCCTCGTTGCTCTTTTAAGAGGTGTCCAGTGT

>IGHV3-30*01-A|11

AGCTCTGGGAGAGGAGCCCAGCACTGGAAGTCGCCGGTGTTTCCATTCGGTGATCAGCACTGAACACAGAGGACTCACCATGGAGTTTGGGCTGAGCTGGGTTTTCCTCGTTGCTCTTTTAAGAGGTGTCCAGTGT

>IGHV3-30*02-A|3

AGCTCTGGGAGAGGAGCCCAGCACTAGAAGTCGGCGGTGTTTCCATTCGGTGATCATCACTGAACACAGAGGACTCACCATGGAGTTTGGGCTGAGCTGGGTTTTCCTCGTTGCTCTTTTAAGAGGTGTCCAGTGT

>IGHV3-30*02-B|6

AGCTCTGGGAGAGGAGCCCAGCACTAGAAGTCGGCGGTGTTTCCATTCGGTGATCAGCACTGAACACAGAGGACTCACCATGGAGTTTGGGCTGAGCTGGGTTTTCCTCGTTGCTCTTTTAAGAGGTGTCCAGTGT

>IGHV3-30*02_S4989-A|1

AGCTCTGGGAGAGGAGCCCAGCACTAGAAGTCGGCGGTGTTTCCATTCGGTGATCAGCACTGAACACAGAGGACTCACCATGGAGTTTGGGCTGAGCTGGGTTTTCCTCGTTGCTCTTTTAAGAGGTGTCCAGTGT

>IGHV3-30*03-A|5

AGCTCTGGGAGAGGAGCCCAGCACTAGAAGTCGGCGGTGTTTCCATTCGGTGATCAGCACTGAACACAGAGGACTCACCATGGAGTTTGGGCTGAGCTGGGTTTTCCTCGTTGCTCTTTTAAGAGGTGTCCAGTGT

>IGHV3-30*04-A|6

AGCTCTGGGAGACGAGCCCAGCACTGGAAGTCGCCGGTGTTTCCATTCGGTGATCATCACTGAACACAGAGGACTCACCATGGAGTTTGGGCTGAGCTGGGTTTTCCTCGTTGCTCTTTTAAGAGGTGTCCAGTGT

>IGHV3-30*04-B|12

AGCTCTGGGAGACGAGCCCAGCACTGGAAGTCGCCGGTGTTTCCATTCGGTGATCAGCACTGAACACAGAGGACTCACCATGGAGTTTGGGCTGAGCTGGGTTTTCCTCGTTGCTCTTTTAAGAGGTGTCCAGTGT

>IGHV3-30*04_S7005-A|1

AGCTCTGGGAGAGGAGCCCAGCACTAGAAGTCGGCGGTGTTTCCATTCGGTGATCAGCACTGAACACAGAGGACTCACCATGGAGTTTGGGCTGAGCTGGGTTTTCCTCGTTGCTCTTTTAAGAGGTGTCCAGTGT

>IGHV3-30*18-A|92

AGCTCTGGGAGAGGAGCCCAGCACTAGAAGTCGGCGGTGTTTCCATTCGGTGATCAGCACTGAACACAGAGGACTCACCATGGAGTTTGGGCTGAGCTGGGTTTTCCTCGTTGCTCTTTTAAGAGGTGTCCAGTGT

>IGHV3-30*19_S5956-A|1

AGCTCTGGGAGACGAGCCCAGCACTGGAAGTCGCCGGTGTTTCCATTCGGTGATCATCACTGAACACAGAGGACTCACCATGGAGTTTGGGCTGAGCTGGGTTTTCCTCGTTGCTCTTTTAAGAGGTGTCCAGTGT

>IGHV3-33*01-A|96

AGCTCTGGGAGAGGAGCCCAGCACTAGAAGTCGGCGGTGTTTCCATTCGGTGATCAGCACTGAACACAGAGGACTCACCATGGAGTTTGGGCTGAGCTGGGTTTTCCTCGTTGCTCTTTTAAGAGGTGTCCAGTGT

>IGHV3-33*01_S3418-A|1

AGCTCTGGGAGAGGAGCCCAGCACTAGAAGTCGGCGGTGTTTCCATTCGGTGATCAGCACTGAACACAGAGGACTCACCATGGAGTTTGGGCTGAGCTGGGTTTTCCTCGTTGCTCTTTTAAGAGGTGTCCAGTGT

>IGHV3-43D*03-A|15

AGCTCTGGGAAAGGAGCCCCAGCCCTGAGATTCCCAGGTGTTTCCATTCGGTGATCAGCACTGAACACAGAACTCACCATGGAGTTTGGACTGAGCTGGGTTTTCCTTGTTGCTATTTTAAAAGGTGTCCAGTGT

>IGHV3-43D*04-A|13

AGCTCTGGGAAAGGAGCCCCAGCCCTGAGATTCCCAGGTGTTTCCATTCGGTGATCAGCACTGAACACAGAACTCACCATGGAGTTTGGACTGAGCTGGGTTTTCCTTGTTGCTATTTTAAAAGGTGTCCAGTGT

>IGHV3-43D*04_S5432-A|1

AGCTCTGGGAAAGGAGCCCCAGCCCTGAGATTCCCAGGTGTTTCCATTCGGTGATCAGCACTGAACACAGAACTCACCATGGAGTTTGGACTGAGCTGGGTTTTCCTTGTTGCTATTTTAAAAGGTGTCCAGTGT

>IGHV3-43*01-A|73

AGCTCTGGGAGAGGAGCCCCAGCCCTGAGATTCCCAGGTGTTTCCATTCGGTGATCAGCACTGAACACAGAGAACGCACCATGGAGTTTGGACTGAGCTGGGTTTTCCTTGTTGCTATTTTAAAAGGTGTCCAGTGT

>IGHV3-43*02-A|11

AGCTCTGGGAGAGGAGCCCCAGCCCTGAGATTCCCAGGTGTTTCCATTCGGTGATCAGCACTGAACACAGAGAACTCACCATGGAGTTTGGACTGAGCTGGGTTTTCCTTGTTGCTATTTTAAAAGGTGTCCAGTGT

>IGHV3-48*01-A|50

AGAGAGGTGCCTTAGCCCTGGATTCCAAGGCATTTCCACTTGGTGATCAGCACTGAACACAGAGGACTCACCATGGAGTTGGGGCTGTGCTGGGTTTTCCTTGTTGCTATTTTAGAAGGTGTCCAGTGT

>IGHV3-48*02-A|65

AGAGAGGTGCCTTAGCCCTGGATTCCAAGGCATTTCCACTTGGTGATCAGCACTGAACACAGAGGACTCACCATGGAGTTGGGGCTGTGCTGGGTTTTCCTTGTTGCTATTTTAGAAGGTGTCCAGTGT

>IGHV3-48*03-A|34

TCTCAGAGAGGTGCCTTAGCCCTGGATTCCAAGGCATTTCCACTTGGTGATCAGCACTGAACACAGAGGACTCACCATGGAGTTGGGGCTGTGCTGGGTTTTCCTTGTTGCTATTTTAGAAGGTGTCCAGTGT

>IGHV3-48*04-A|10

AGAGAGGTGCCTTAGCCCTGGATTCCAAGGCATTTCCACTTGGTGATCAGCACTGAACACAGAGGACTCACCATGGAGTTGGGGCTGTGCTGGGTTTTCCTTGTTGCTATTTTAGAAGGTGTCCAGTGT

>IGHV3-49*03-A|59

CTCTGGGAGAGGAGCCCCAGCCGTGAGATTCCCAGGAGTTTCCACTTGGTGATCAGCACTGAACACAGACCACCAACCATGGAGTTTGGGCTTAGCTGGGTTTTCCTTGTTGCTATTTTAAAAGGTGTCCAATGT

>IGHV3-49*04-A|37

AGCTCTGGGAGAGGAGCCCCAGCCGTGAGATTCCCAGGAGTTTCCACTTGGTGATCAGCACTGAACACAGACCACCAACCATGGAGTTTGGGCTTAGCTGGGTTTTCCTTGTTGCTATTTTAAAAGGTGTCCAATGT

>IGHV3-49*05-A|60

CTCTGGGAGAGGAGCCCCAGCCGTGAGATTCCCAGGAGTTTCCACTTGGTGATCAGCACTGAACACAGACCACCAACCATGGAGTTTGGGCTTAGCTGGGTTTTCCTTGTTGCTATTTTAAAAGGTGTCCAATGT

>IGHV3-53*01-A|68

AGAGGAGCCCAGCACTGGGATTCCGAGGTGTTTCCATTCGGTGATCAGCACTGAACACAGAGGACTCACCATGGAGTTTTGGCTGAGCTGGGTTTTCCTTGTTGCTATTTTAAAAGGTGTCCAGTGT

>IGHV3-53*02-A|49

GAGGAGCCCAGCACTGGGATTCCGAGGTGTTTCCATTCGGTGATCAGCACTGAACACAGAGGACTCACCATGGAGTTTTGGCTGAGCTGGGTTTTCCTTGTTGCTATTTCAAAAGGTGTCCAGTGT

>IGHV3-53*02_S9017-A|1

AGAGGAGCCCAGCACTGGGATTCCGAGGTGTTTCCATTCGGTGATCAGCACTGAACACAGAGGACTCACCATGGAGTTTTGGCTGAGCTGGGTTTTCCTTGTTGCTATTTCAAAAGGTGTCCAGTGT

>IGHV3-53*04-A|29

AGAGGAGCCCAGCACTGGGATTCCGAGGTGTTTCCATTCGGTGATCAGCACTGAACACAGAGGACTCACCATGGAGTTTTGGCTGAGCTGGGTTTTCCTTGTTGCTATTTTAAAAGGTGTCCAGTGT

>IGHV3-64D*06-A|52

AGCTCTGGGAGAGGAGCCCCAGGCCCGGGATTCCCAGGTGTTTCCATTCAGTGATCAGCACTGAAGACAGAAGACTCATCATGGAGTTCTGGCTGAGCTGGGTTCTCCTTGTTGCCATTTTAAAAGATGTCCAGTGT

>IGHV3-64D*06_S5429-A|12

AGCTCTGGGAGAGGAGCCCCAGGCCCGGGATTCCCAGGTGTTTCCATTCAGTGATCAGCACTGAAGACAGAAGACTCATCATGGAGTTCTGGCTGAGCTGGGTTCTCCTTGTTGCCGTTTTAAAAGATGTCCAGTGT

>IGHV3-64*01-A|46

GCTCTGGGAGAGGAGCCCCCGCCCTGGGATTCCCAGGTGTTTTCATTTGGTGATCAGCACTGAACACAGAAGAGTCATGACGGAGTTTGGGCTGAGCTGGGTTTTCCTTGTTGCTATTTTTAAAGGTGTCCAGTGT

>IGHV3-64*05_S2482-A|10

AGCTCTGGGAGAGGAGCCCCAGGCCCGGGATTCCCAGGTGTTTCCATTCAGTGATCAGCACTGAAGACAGAAGACTCATCATGGAGTTCTGGCTGAGCTGGGTTCTCCTTGTTGCCATTTTAAAAGATGTCCAGTGT

>IGHV3-66*01-A|42

AGCTCTGGGAGAGGAGCCCAGCACTGGGATTCCGAGGTGTTTCCATTCAGTGATCTGCACTGAACACAGAGGACTCGCCATGGAGTTTGGGCTGAGCTGGGTTTTCCTTGTTGCTATTTTAAAAGGTGTCCAGTGT

>IGHV3-66*02-A|29

AGCTCTGGGAGAGGAGCCCAGCACTGGGATTCCGAGGTGTTTCCATTCAGTGATCTGCACTGAACACAGAGGACTCGCCATGGAGTTTGGGCTGAGCTGGGTTTTCCTTGTTGCTATTTTAAAAGGTGTCCAGTGT

>IGHV3-66*02_S8911-A|1

AGCTCTGGGAGAGGAGCCCAGCACTGGGATTCCGAGGTGTTTCCATTCAGTGATCTGCACTGAACACAGAGGACTCGCCATGGAGTTTGGGCTGAGCTGGGTTTTCCTTGTTGCTATTTTAAAAGGTGTCCAGTGT

>IGHV3-72*01-A|47

AGAGCGGAGCCCCAGCCCCAGAATTCCCAGGTGTTTTCATTTGGTGATCAGCACTGAACACAGAGGACTCACCATGGAGTTTGGGCTGAGCTGGGTTTTCCTTGTTGTTATTTTACAAGGTGTCCAGTGT

>IGHV3-73*01-A|51

AGCTCTGGGAGAGGAGCTCCAGCCTTGGGATTCCCAGCTGTCTCCACTCGGTGATCGGCACTGAATACAGGAGACTCACCATGGAGTTTGGGCTGAGCTGGGTTTTCCTTGTTGCTATTTTAAAAGGTGTCCAGTGT

>IGHV3-73*02-A|64

AGCTCTGGGAGAGGAGCTCCAGCCTTGGGATTCCCAGCTGTCTCCACTCGGTGATCGGCACTGAATACAGGAGACTCACCATGGAGTTTGGGCTGAGCTGGGTTTTCCTTGTTGCTATTTTAAAAGGTGTCCAGTGT

>IGHV3-74*01-A|98

AGCCCCAGCCCTGGGATTCCCAGCTGTTTCTGCTTGCTGATCAGGACTGCACACAGAGAACTCACCATGGAGTTTGGGCTGAGCTGGGTTTTCCTTGTTGCTATTTTAAAAGGTGTCCAGTGT

>IGHV3-7*01-A|76

TCTCAGAGAGGAGCCTTAGCCCTGGACTCCAAGGCCTTTCCACTTGGTGATCAGCACTGAGCACAGAGGACTCACCATGGAATTGGGGCTGAGCTGGGTTTTCCTTGTTGCTATTTTAGAAGGTGTCCAGTGT

>IGHV3-7*01-B|2

AGAGAGGAGCCTTAGCCCTGGACTCCAAGGCCTTTCCACTTGGTGATCAGCACTGAGCACAGAGGACTCACCATGGAGTTGGGGCTGAGCTGGGTTTTCCTTGTTGCTATTTTAGAAGGTGTCCAGTGT

>IGHV3-7*03-A|44

CTCAGAGAGGAGCCTTAGCCCTGGACTCCAAGGCCTTTCCACTTGGTGATCAGCACTGAGCACAGAGGACTCACCATGGAGTTGGGGCTGAGCTGGGTTTTCCTTGTTGCTATTTTAGAAGGTGTCCAGTGT

>IGHV3-7*03-B|1

CTCAGAGAGGAGCCTTAGCCCTGGACTCCAAGGCCTTTCCACTTGGTGATCAGCACTGAGCACAGAGGACTCACCATGGAATTGGGGCTGAGCTGGGTTTTCCTTGTTGCTATTTTAGAAGGTGTCCAGTGT

>IGHV3-7*03_S9833-A|11

AGAGAGGAGCCTTAGCCCTGGACTCCAAGGCCTTTCCACTTGGTGATCAGCACTGAGCACAGAGGACTCACCATGGAGTTGGGGCTGAGCTGGGTTTTCCTTGTTGCTATTTTAGAAGGTGTCCAGTGT

>IGHV3-7*04-A|21

AGCCTTAGCCCTGGACTCCAAGGCCTTTCCACTTGGTGATCAGCACTGAGCACAGAGGACTCACCATGGAGTTGGGGCTGAGCTGGGTTTTCCTTGTTGCTATTTTAGAAGGTGTCCAGTGT

>IGHV3-9*01-A|77

AGCTCTGGGAGAGGAGCCCCAGCCCTGAGATTCCCAGGTGTTTCCATTCAGTGATCAGCACTGAACACAGAGGACTCACCATGGAGTTGGGACTGAGCTGGATTTTCCTTTTGGCTATTTTAAAAGGTGTCCAGTGT

>IGHV3-9*03-A|2

AGCTCTGGGAGAGGAGCCCCAGCCCTGAGATTCCCAGGTGTTTCCATTCAGTGATCAGCACTGAACACAGAGGACTCACCATGGAGTTGGGACTGAGCTGGATTTTCCTTTTGGCTATTTTAAAAGGTGTCCAGTGT

>IGHV4-30-2*01-A|3

ACTTTCTGAGAGTCCTGGACCTCTGTGCAAGAACATGAAACACCTGTGGTTCTTCCTCCTGCTGGTGGCAGCTCCCAGATGGGTCCTGTCC

>IGHV4-30-2*01-B|1

ACTTTCTGAGAGTCCTGGACCTCCTGCACAAGAACATGAAACACCTGTGGTTCTTCCTCCTGCTGGTGGCAGCTCCCAGATGGGTCCTGTCC

>IGHV4-30-2*01-C|69

ACTTTCTGAGAGTCCTGGACCTCCTGTGCAAGAACATGAAACACCTGTGGTTCTTCCTCCTGCTGGTGGCAGCTCCCAGATGGGTCCTGTCC

>IGHV4-30-2*01_S6723-A|1

ACTTTCTGAGAGTCCTGGACCTCCTGTGCAAGAACATGAAACACCTGTGGTTCTTCCTCCTGCTGGTGGCAGCTCCCAGATGGGTCCTGTCC

>IGHV4-30-4*01-A|62

ATACTTTCTGAGAGTCCTGGACCTCCTGTGCAAGAACATGAAACACCTGTGGTTCTTCCTCCTGCTGGTGGCAGCTCCCAGATGGGTCCTGTCC

>IGHV4-30-4*01_S5061-A|2

ATACTTTCTGAGAGTCCTGGACCTCCTGTGCAAGAACATGAAACACCTGTGGTTCTTCCTCCTGCTGGTGGCAGCTCCCAGATGGGTCCTGTCC

>IGHV4-30-4*07-A|2

ATACTTTCTGAGAGTCCTGGACCTCTGTGCAAGAACATGAAACACCTGTGGTTCTTCCTCCTGCTGGTGGCAGCTCCCAGATGGGTCCTGTCC

>IGHV4-30-4*07-B|1

ATACTTTCTGAGAGTCCTGGACCTCCTGTGCAAGAACATGAAACACCTGTGGTTCTTCCTCCTGCTGGTGGCAGCTCCCAGATGGGTCCTGTCC

>IGHV4-30-4*08-A|1

ACTTTCTGAGAGTCCTGGACCTCCTGTGCAAGAACATGAAACACCTGTGGTTCTTCCTCCTGCTGGTGGCAGCTCCCAGATGGGTCCTGTCC

>IGHV4-31*01-A|9

ATACTTTCTGAGAGTCCTGGACCTCCTGTGCAAGAACATGAAACACCTGTGGTTCTTCCTCCTGCTGGTGGCAGCTCCCAGATGGGTCCTGTCC

>IGHV4-31*03-A|89

ATACTTTCTGAGAGTCCTGGACCTCCTGTGCAAGAACATGAAACACCTGTGGTTCTTCCTCCTGCTGGTGGCAGCTCCCAGATGGGTCCTGTCC

>IGHV4-34*01-A|98

AGTGCTTTCTGAGAGTCATGGACCTCCTGCACAAGAACATGAAACACCTGTGGTTCTTCCTCCTCCTGGTGGCAGCTCCCAGATGGGTCCTGTCC

>IGHV4-38-2*01-A|23

CTTTCTGAGAGTCATGGACCTCCTGTGCAAGAACATGAAGCACCTGTGGTTTTTCCTCCTGCTGGTGGCAGCTCCCAGATGGGTCCTGTCC

>IGHV4-38-2*02-A|19

CTTTCTGAGAGTCATGGACCTCCTGTGCAAGAACATGAAGCACCTGTGGTTTTTCCTCCTGCTGGTGGCAGCTCCCAGATGGGTCCTGTCC

>IGHV4-39*01-A|2

CTTTCTGAGAGTCATGGATCTCATGTGCAAGAAAATGAAGCACCTGTGGTTCTTCCTCCTGGTGGTGGCGGCTCCCAGATGGGTCCTGTCC

>IGHV4-39*01-B|92

CTTTCTGAGAGTCATGGATCTCATGTGCAAGAAAATGAAGCACCTGTGGTTCTTCCTCCTGCTGGTGGCGGCTCCCAGATGGGTCCTGTCC

>IGHV4-39*01_S7498-A|1

CTTTCTGAGAGTCATGGATCTCATGTGCAAGAAAATGAAGCACCTGTGGTTCTTCCTCCTGCTGGTGGCGGCTCCCAGATGGGTCCTGTCC

>IGHV4-39*07-A|13

CTTTCTGAGAGTCATGGACCTCCTGTGCAAGAACATGAAGCACCTGTGGTTCTTCCTCCTGCTGGTGGCGGCTCCCAGATGGGTCCTGTCC

>IGHV4-39*07_S0654-A|2

CTTTCTGAGAGTCATGGACCTCCTGTGCAAGAACATGAAGCACCTGTGGTTCTTCCTCCTGCTGGTGGCGGCTCCCAGATGGGTCCTGTCC

>IGHV4-4*02-A|7

TACTTTCTGAGACTCATGGGCCTCCTGCACAAGAACATGAAACACCTGTGGTTCTTCCTCCTCCTGGTGGCAGCTCCCAGATGGGTCCTGTCT

>IGHV4-4*02-B|1

ACTTTCTGAGAGTCATGGACCTCCTGCACAAGAACATGAAACACCTGTGGTTCTTCCTCCTCCTGGTGGCAGCTCCCAGATGGGTCCTGTCT

>IGHV4-4*02-C|57

TACTTTCTGAGACTCATGGACCTCCTGCACAAGAACATGAAACACCTGTGGTTCTTCCTCCTCCTGGTGGCAGCTCCCAGATGGGTCCTGTCT

>IGHV4-4*02-D|22

ACTTTCTGAGAGTCCTGGACCTCCTGCACAAGAACATGAAACACCTGTGGTTCTTCCTCCTCCTGGTGGCAGCTCCCAGATGGGTCCTGTCT

>IGHV4-4*02-E|3

ATACTTTCTGAGACTCATGGACCTCCTGTGCAAGAACATGAAACACCTGTGGTTCTTCCTCCTCCTGGTGGCAGCTCCCAGATGGGTCCTGTCT

>IGHV4-4*02-F|6

ATACTTTCTGAGACTCATGGACCTCCTGCACAAGAACATGAAACACCTGTGGTTCTTCCTCCTCCTGGTGGCAGCTCCCAGATGGGTCCTGTCC

>IGHV4-4*02_S2599-A|1

ACTTTCTGAGAGTCCTGGACCTCCTGCACAAGAACATGAAACACCTGTGGTTCTTCCTCCTCCTGGTGGCAGCTCCCAGATGGGTCCTGTCT

>IGHV4-4*07-A|2

ACTTTCTGAGAGTCCTGGACCTCCTGCACAAGAACATGAAACACCTGTGGTTCTTCCTCCTCCTGGTGGCAGCTCCCAGATGGGTCCTGTCC

>IGHV4-4*07-B|1

ACTTTCTGAGAGTCCTGGACCTCCTGCACAAGAACATGAAACACCTGTGGTTCTTCCTCCTGCTGGTGGCAGCTCCCAGATGGGTCCTGTCC

>IGHV4-4*07-C|1

ACTTTCTGAGAGTCCTGGACCTCCTGTGCAAGAACATGAAACACCTGTGGTTCTTCCTCCTGCTGGTGGCAGCTCCCAGATGGGTCCTGTCC

>IGHV4-4*07-D|8

ACTTTCTGAGACTCATGGACCTCCTGCACAAGAACATGAAACACCTGTGGTTCTTCCTCCTCCTGGTGGCAGCTCCCAGATGGGTCCTGTCC

>IGHV4-4*07-E|55

ACTTTCTGAGACTCATGGACCTCCTGCACAAGAACATGAAACACCTGTGGTTCTTCCTCCTGCTGGTGGCAGCTCCCAGATGGGTCCTGTCC

>IGHV4-4*07-F|1

ATACTTTCTGAGACTCATGGACCTCCTGTGCAAGAACATGAAACACCTGTGGTTCTTCCTCCTCCTGGTGGCAGCTCCCAGATGGGTCCTGTCC

>IGHV4-59*01-A|1

ACTTTCTGAGAGTCCTGGACCTCCTGTGCAAGAACATGAAACATCTGTGGTTCTTCCTTCTCCTGGTGGCGGCTCCCAGATGGGTCCTGTCC

>IGHV4-59*01-B|98

ACTTTCTGAGAGTCCTGGACCTCCTGTGCAAGAACATGAAACATCTGTGGTTCTTCCTTCTCCTGGTGGCAGCTCCCAGATGGGTCCTGTCC

>IGHV4-59*08-A|42

ACTTTCTGAGAGTCCTGGACCTCCTGTGCAAGAACATGAAACATCTGTGGTTCTTCCTTCTCCTGGTGGCAGCTCCCAGATGGGTCCTGTCC

>IGHV4-59*12-A|1

ATACTTTCTGAGACTCATGGACCTCCTGCACAAGAACATGAAACACCTGTGGTTCTTCCTCCTCCTGGTGGCAGCTCCCAGATGGGTCCTGTCC

>IGHV4-61*01-A|63

ACTTTCTGAGAGTCCTGGACCTCCTGTGCAAGAACATGAAACACCTGTGGTTCTTCCTCCTCCTGGTGGCAGCTCCCAGATGGGTCCTGTCC

>IGHV4-61*01_S9413-A|1

ACTTTCTGAGAGTCCTGGACCTCCTGTGCAAGAACATGAAACACCTGTGGTTCTTCCTCCTCCTGGTGGCAGCTCCCAGATGGGTCCTGTCC

>IGHV4-61*02-A|30

ACTTTCTGAGAGTCCTGGACCTCCTGTGCAAGAACATGAAACATCTGTGGTTCTTCCTCCTCCTGGTGGCAGCTCCCAGATGGGTCCTGTCC

>IGHV4-61*02_S0442-A|6

ACTTTCTGAGAGTCCTGGACCTCCTGTGCAAGAACATGAAACATCTGTGGTTCTTCCTCCTCCTGGTGGCAGCTCCCAGATGGGTCCTGTCC

>IGHV5-10-1*01-A|22

AGTCTCCTTCACCACCCAGCTGGGATCTCAGGGCTTCCTTTTCTGTCCTCCTCCAGGATGGGGTCAACCGCCATCCTTGGCCTCCTCCTGGCTGTTCTCCAAGGAGTCTGTGCC

>IGHV5-10-1*03-A|43

AGTCTCCTTCACCACCCAGCTGGGATCTCAGGGCTTCCTTTTCTGTCCTCCTCCAGGATGGGGTCAACCGCCATCCTCGCCCTCCTCCTGGCTGTTCTCCAAGGAGTCTGTGCC

>IGHV5-10-1*03-B|14

AGTCTCCTTCACCACCCAGCTGGGATCTCAGGGCTTCCTTTTCTGTCCTCCTCCAGGATGGGGTCAACCGCCATCCTTGGCCTCCTCCTGGCTGTTCTCCAAGGAGTCTGTGCC

>IGHV5-51*01-A|87

AGTCTCCCTCACTGCCCAGCTGGGATCTCAGGGCTTCATTTTCTGTCCTCCACCATCATGGGGTCAACCGCCATCCTCGCCCTCCTCCTGGCTGTTCTCCAAGGAGTCTGTGCC

>IGHV5-51*01-B|12

AGTCTCCCTCACTGCCCAGCTGGGATCTCAGGGCTTCATTTTCTGTCCTCCACCATCATGGGGTCAACCGCCATCCTTGCCCTCCTCCTGGCTGTTCTCCAAGGAGTCTGTGCC

>IGHV5-51*03-A|40

AGTCTCCCTCACTGCCCAGCTGGGATCTCAGGGCTTCATTTTCTGTCCTCCACCATCATGGGGTCAACCGCCATCCTCGCCCTCCTCCTGGCTGTTCTCCAAGGAGTCTGTGCC

>IGHV5-51*03-B|1

AGTCTCCCTCACCGCCCAGCTGGGATCTCAGGGCTTCATTTTCTGTCCTCCACCATCATGGGGTCAACCGCCATCCTCGCCCTCCTCCTGGCTGTTCTCCAAGGAGTCTGTGCC

>IGHV6-1*01-A|95

AGAGCCTGCTGAATTCTGGCTGACCAGGGCAGTCACCAGAGCTCCAGACAATGTCTGTCTCCTTCCTCATCTTCCTGCCCGTGCTGGGCCTCCCATGGGGTGTCCTGTCA

>IGHV7-4-1*01-A|1

ACCCAACAACCACACCCCTCCTAAGAAGAAGCCCCTAGACCACAGCTCCACACCATGGACTGGACCTGGAGGATCCTCTTCTTGGTGGCAGCAGCAACAGGTGCCCACTCC

>IGHV7-4-1*02-A|26

ACCCAACAACCACACCCCTCCTAAGAAGAAGCCCCTAGACCACAGCTCCACACCATGGACTGGACCTGGAGGATCCTCTTCTTGGTGGCAGCAGCAACAGGTGCCCACTCC
