## Supplementary Figure 1 for "Computational inference, validation, and analysis of 5’UTR-leader sequences of alleles of immunoglobulin heavy chain variable genes"

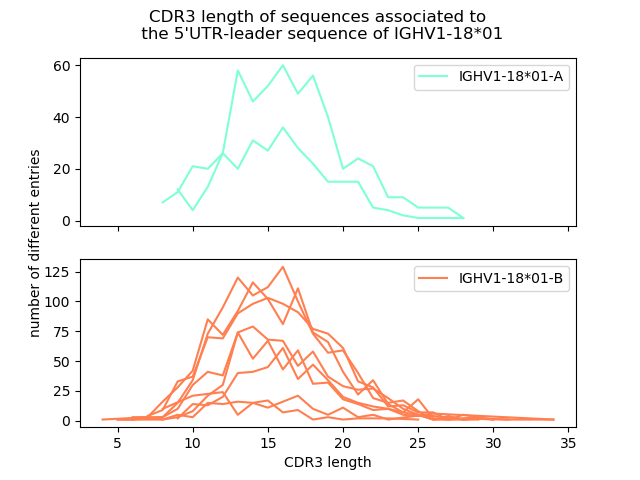

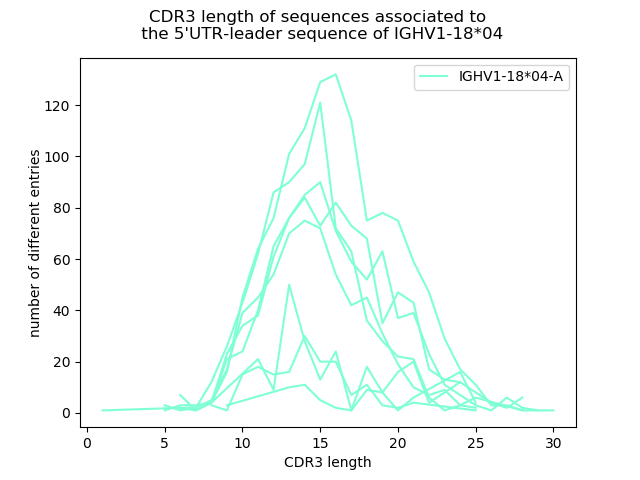

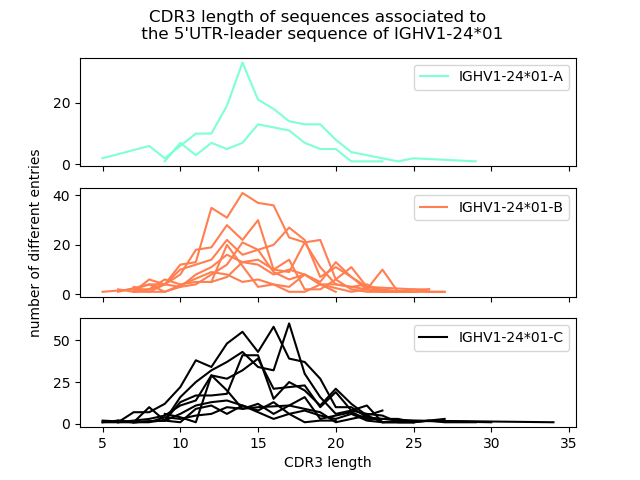

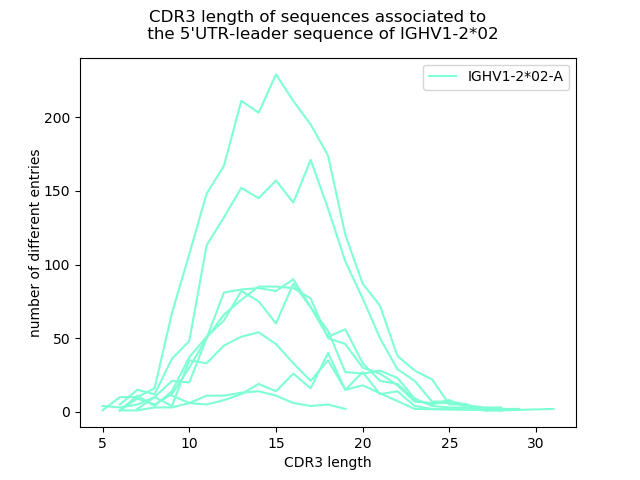

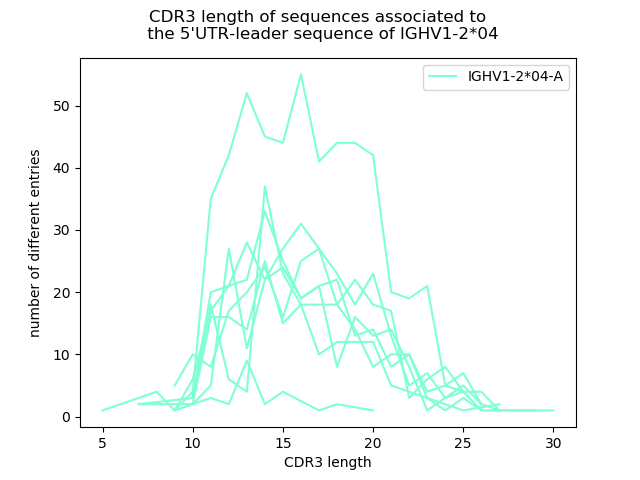

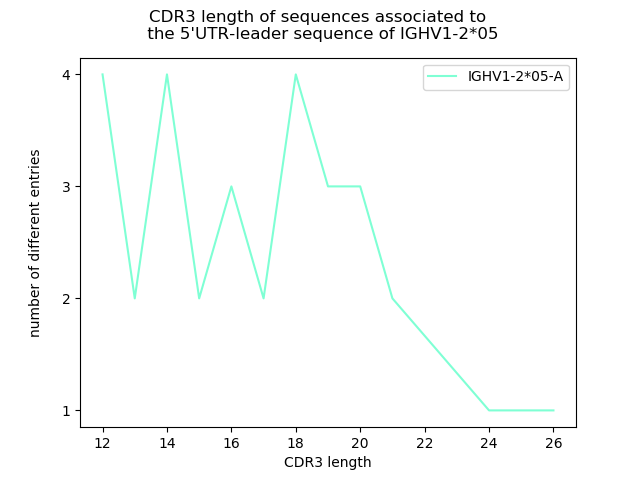

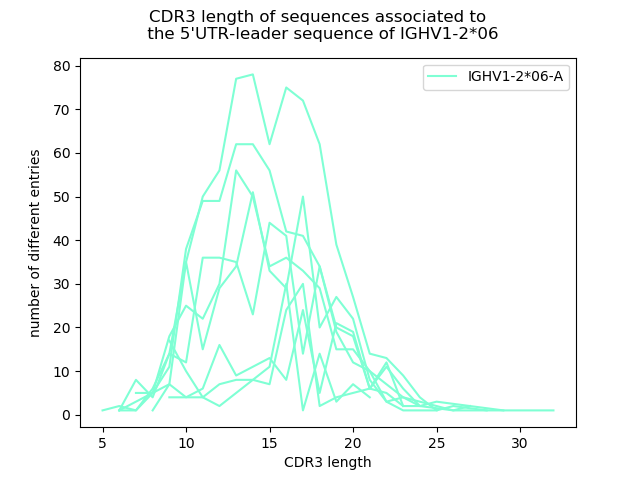

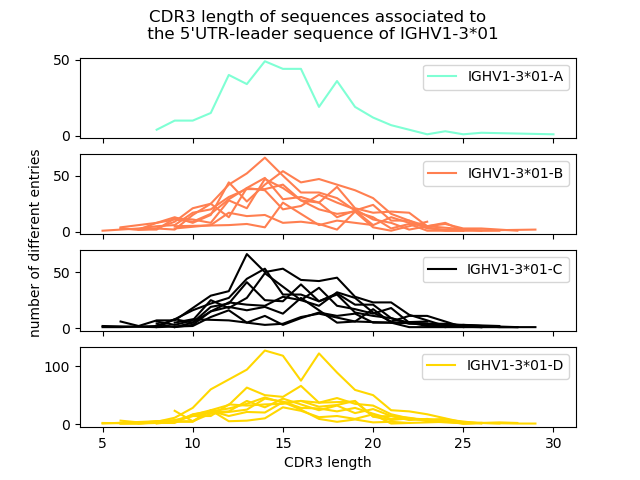

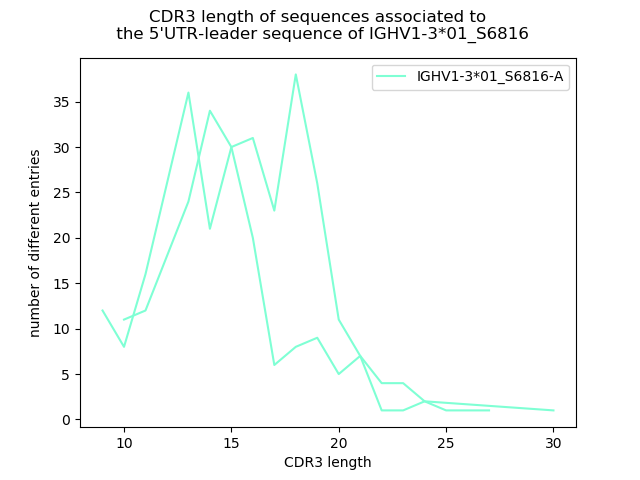

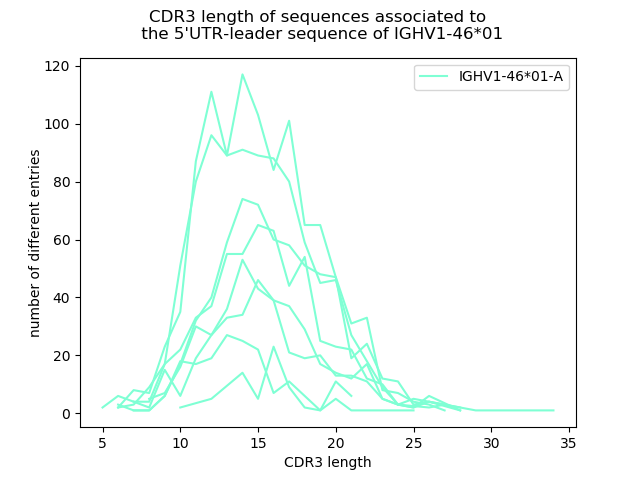

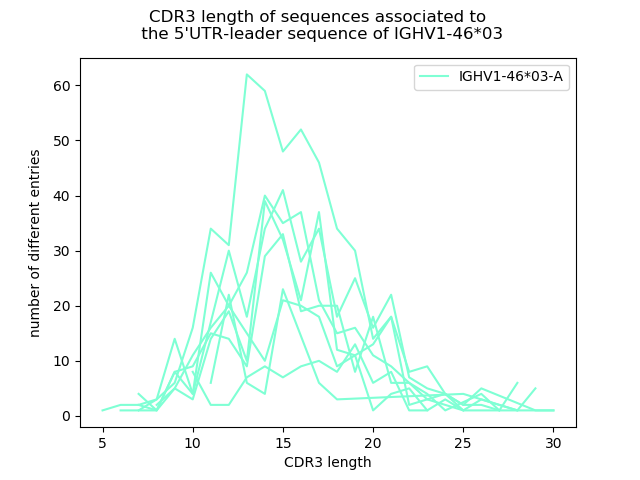

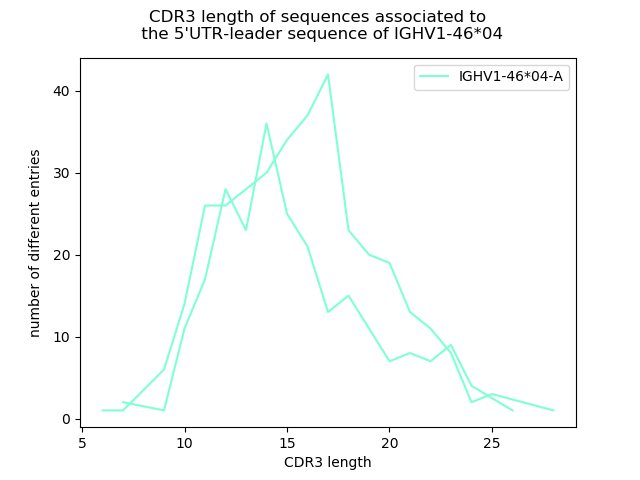

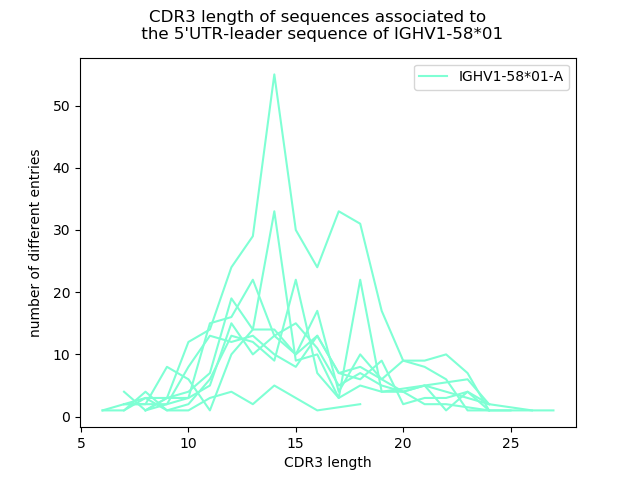

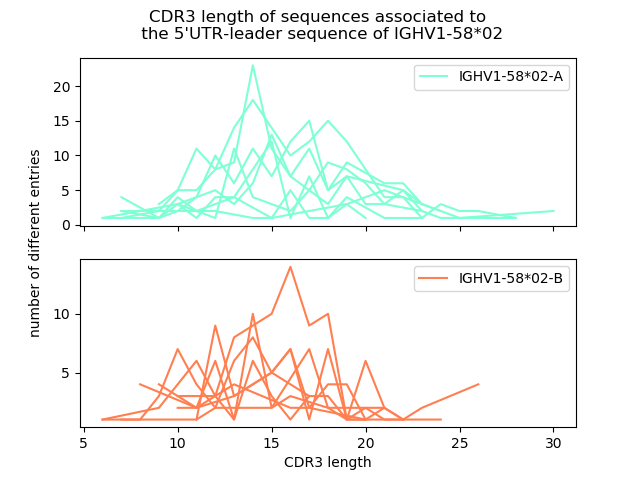

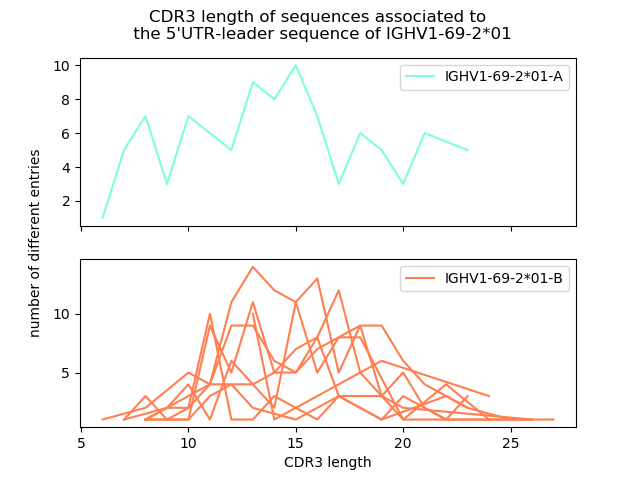

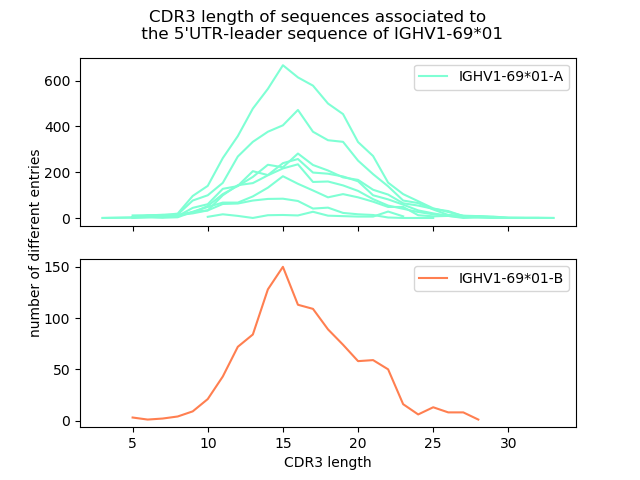

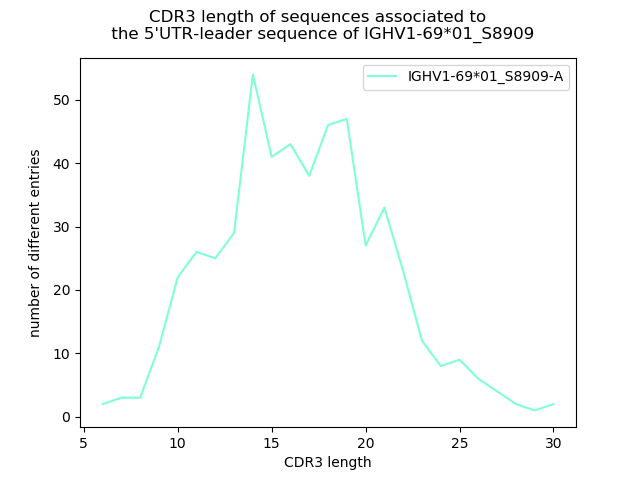

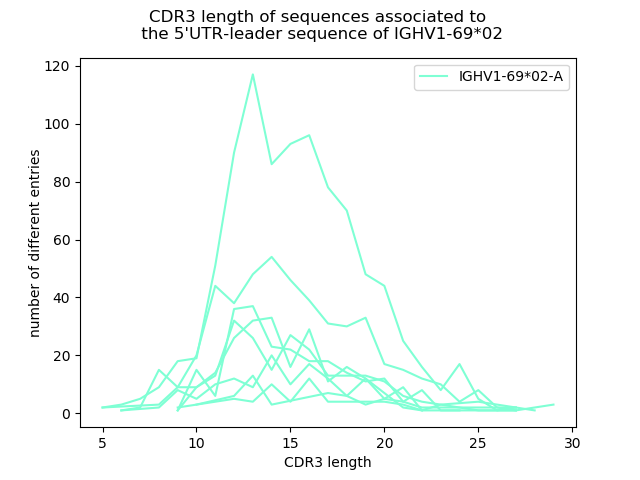

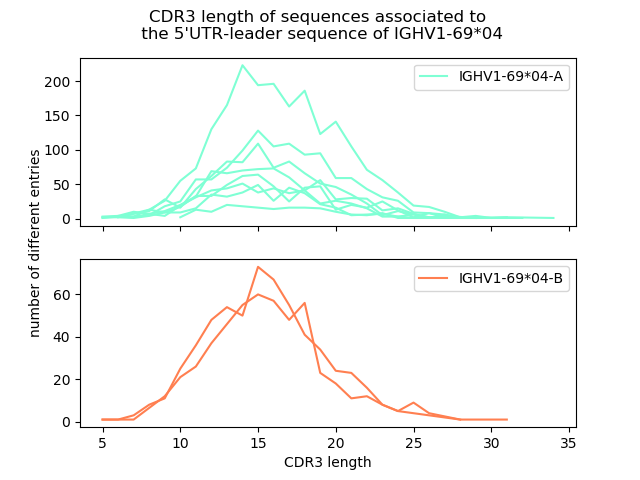

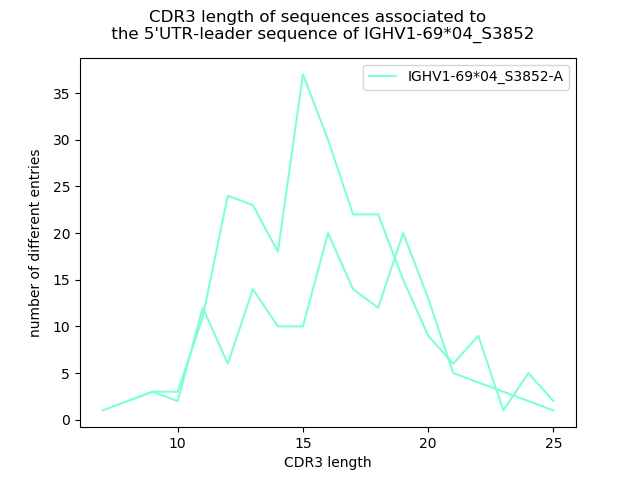

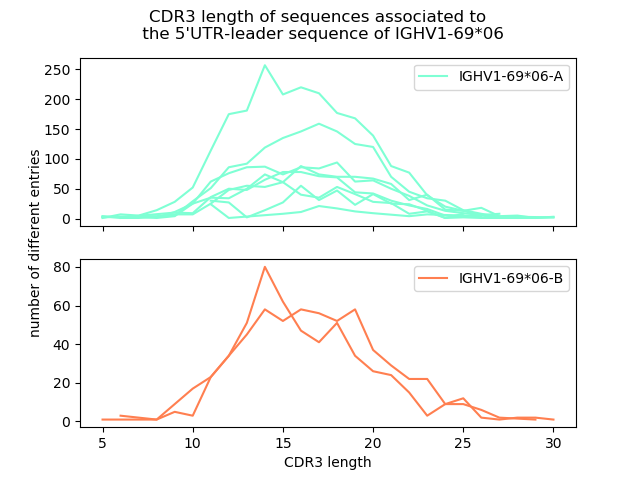

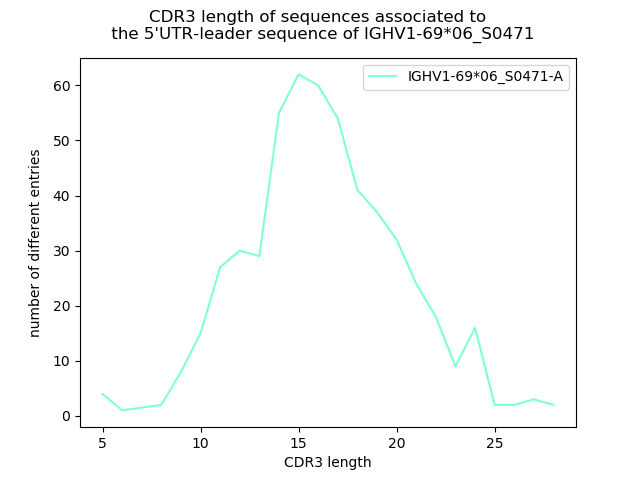

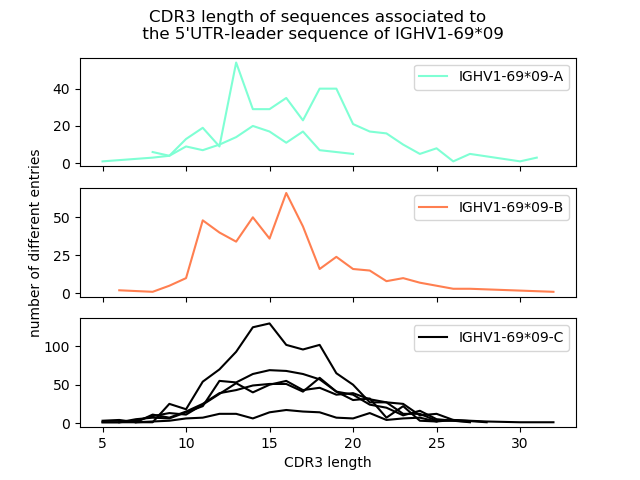

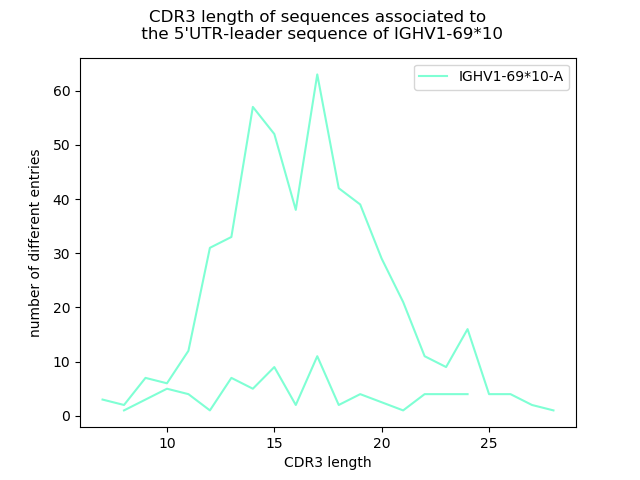

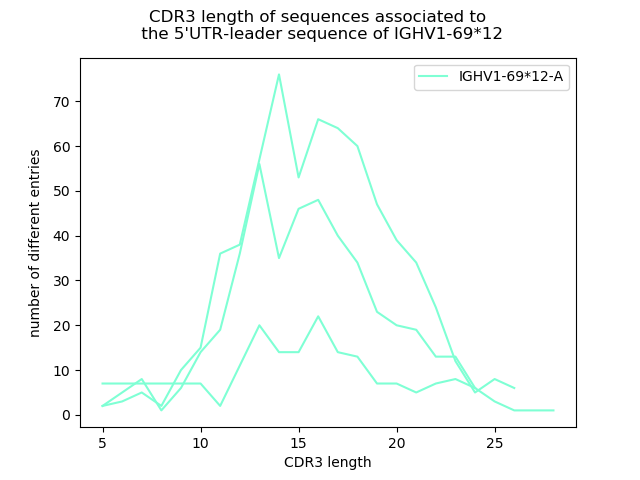

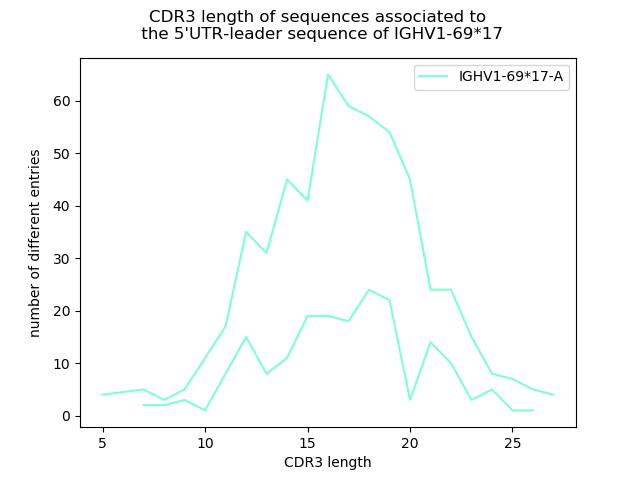

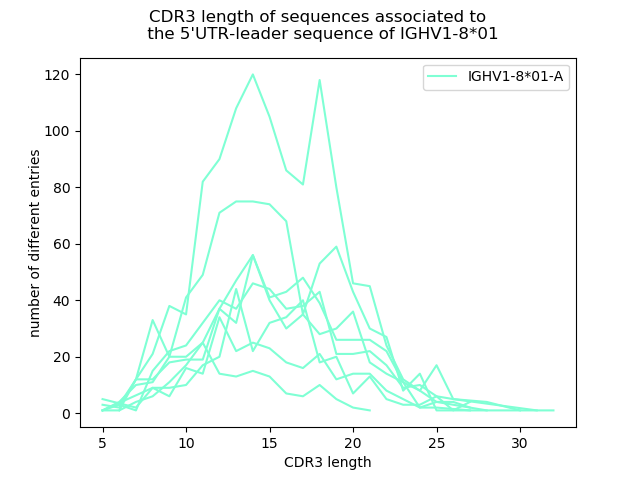

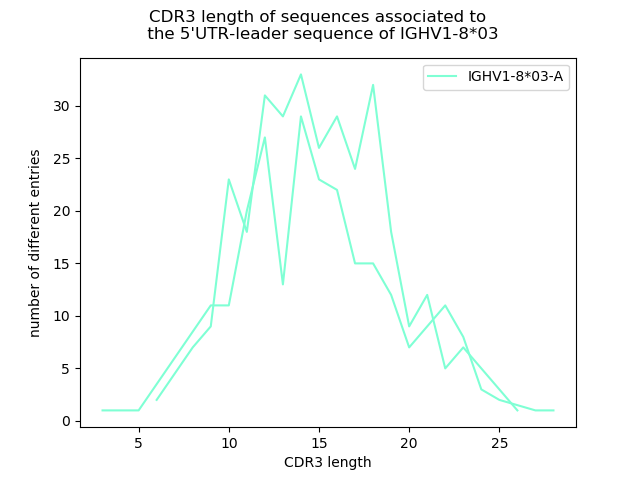

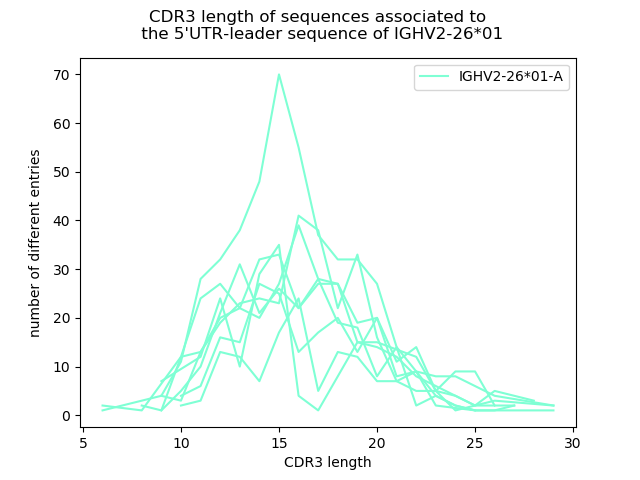

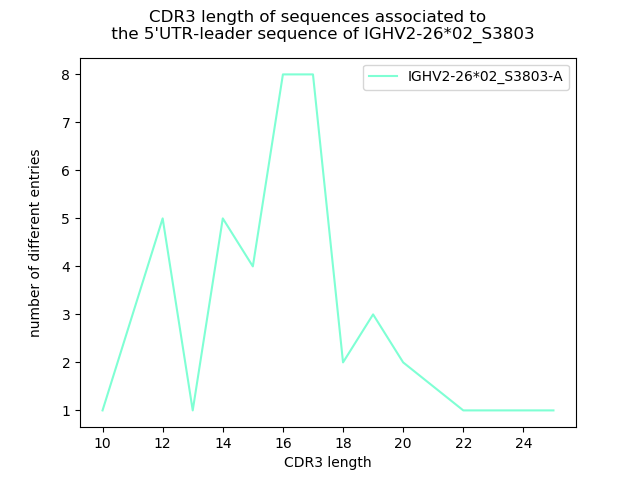

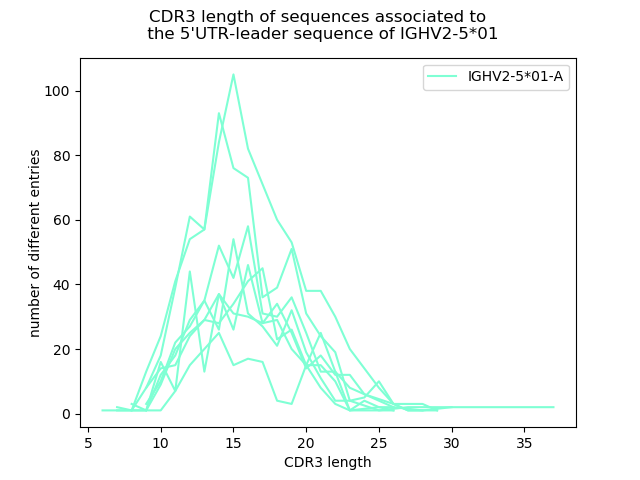

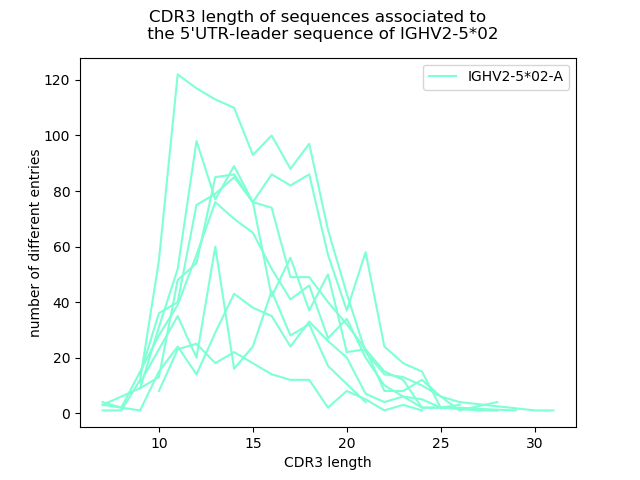

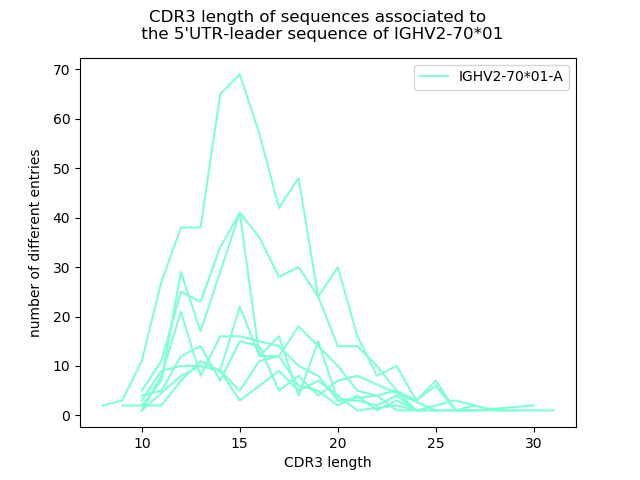

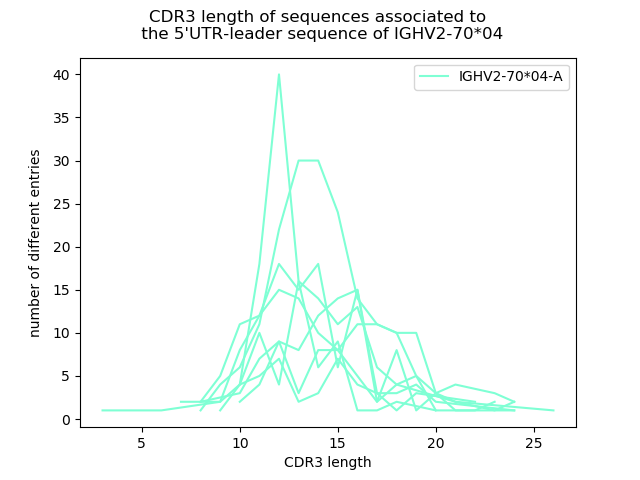

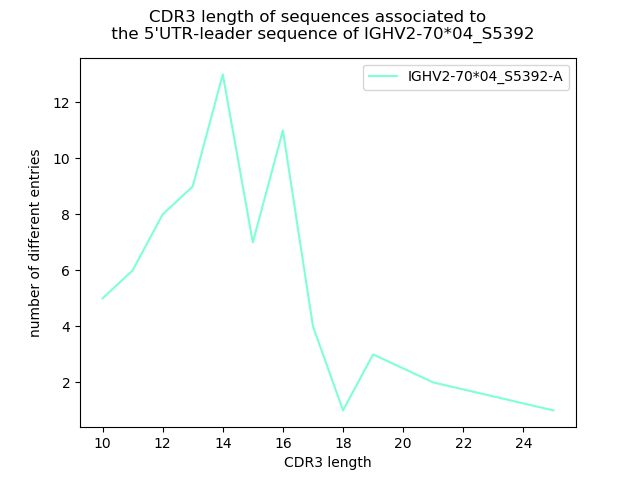

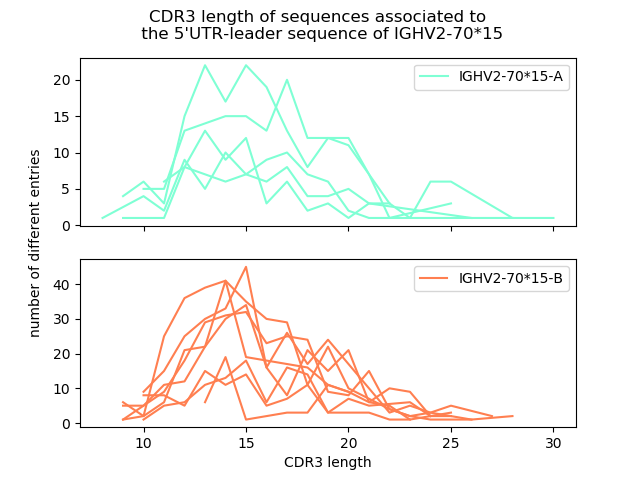

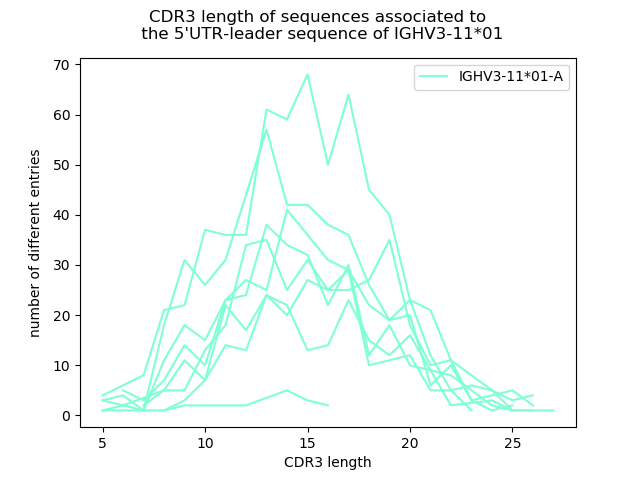

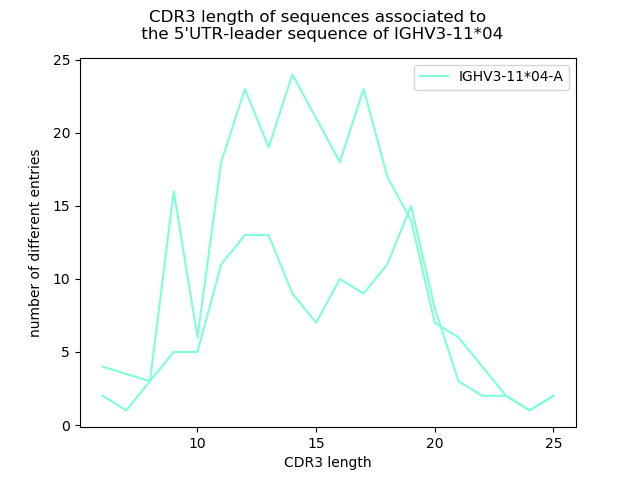

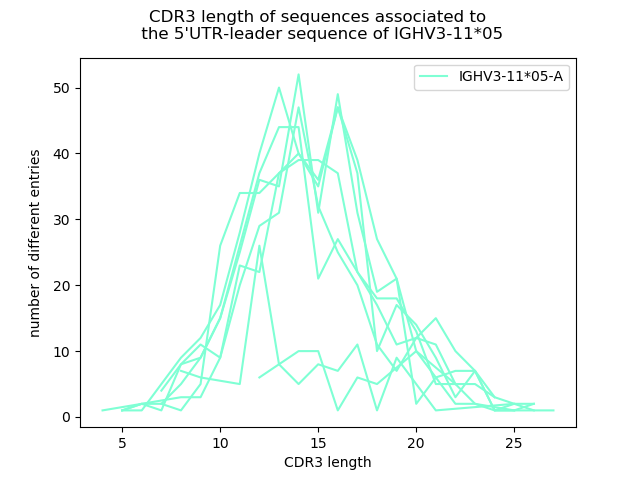

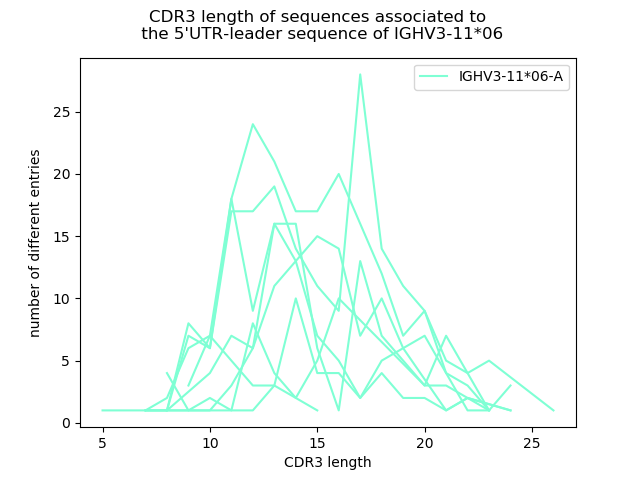

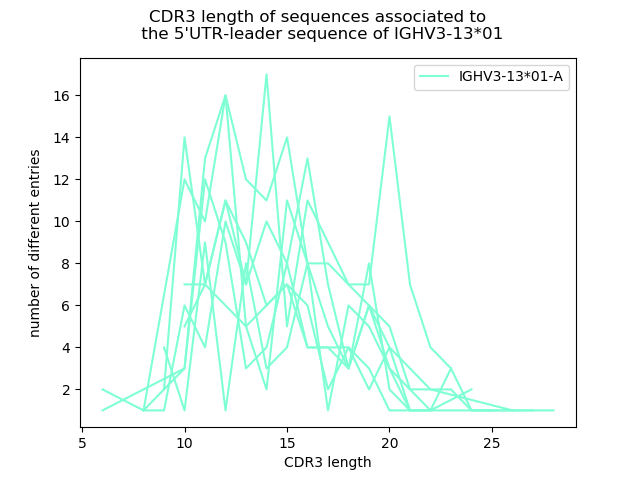

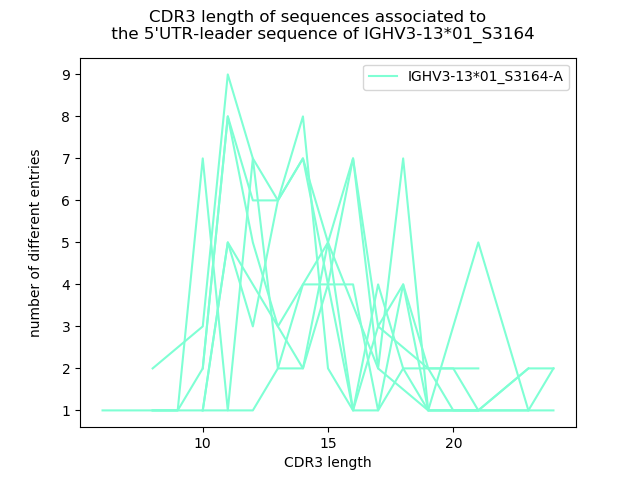

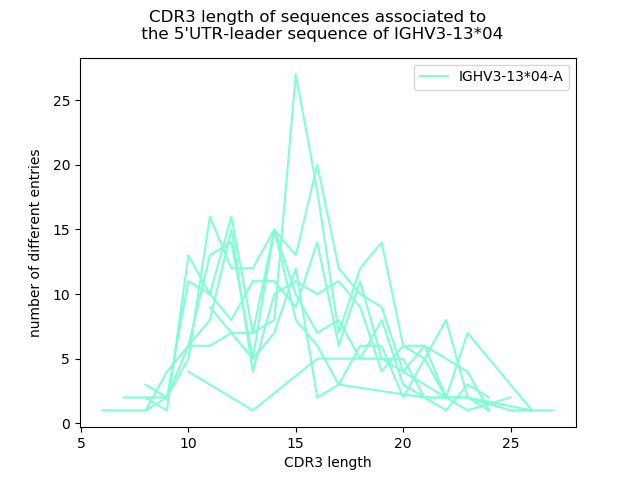

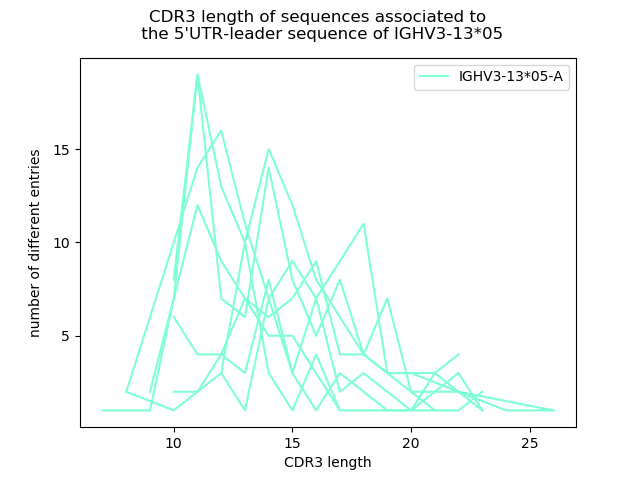

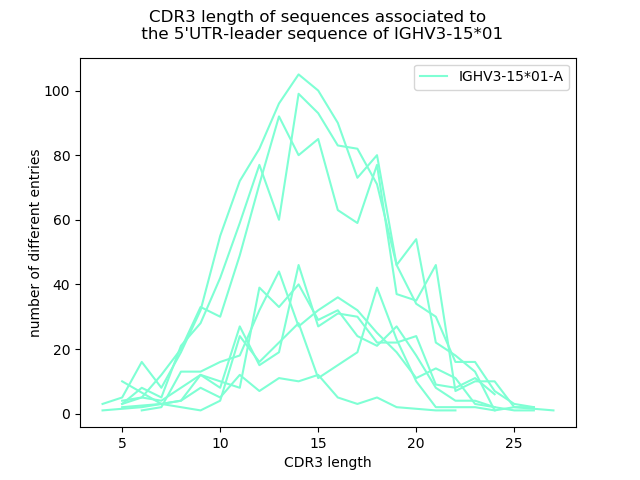

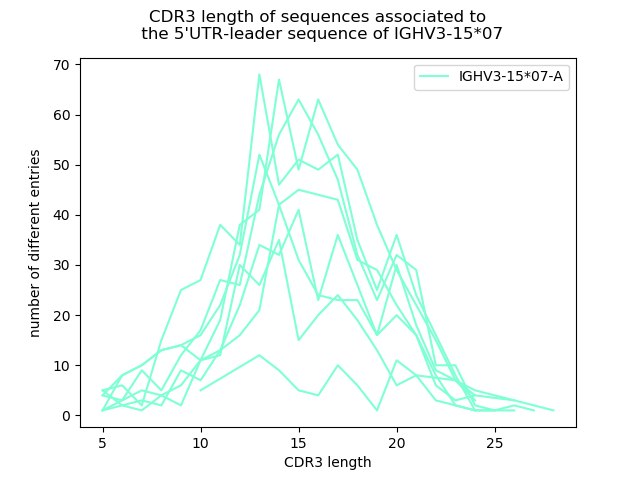

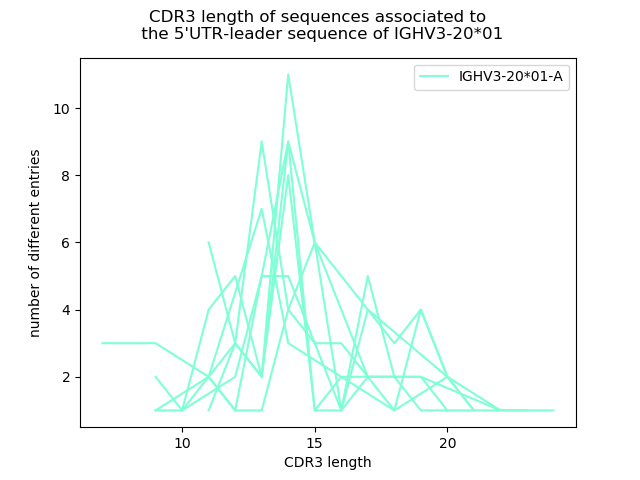

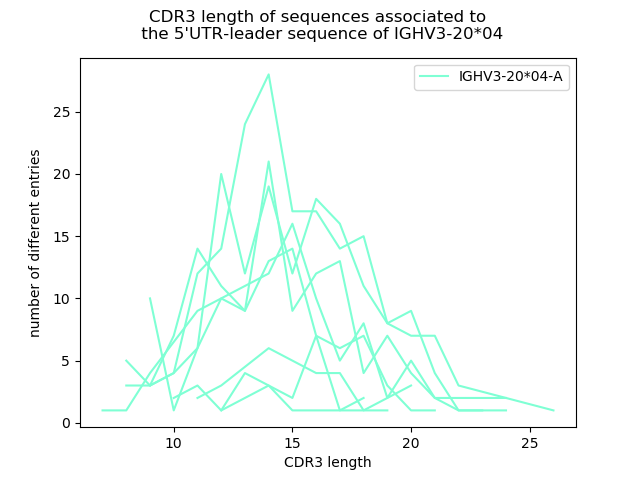

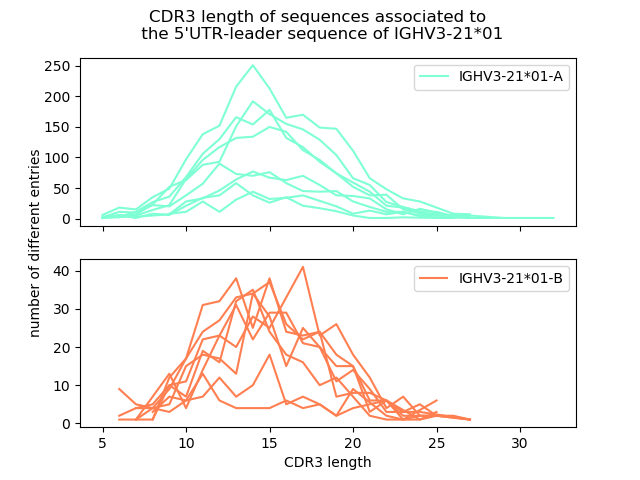

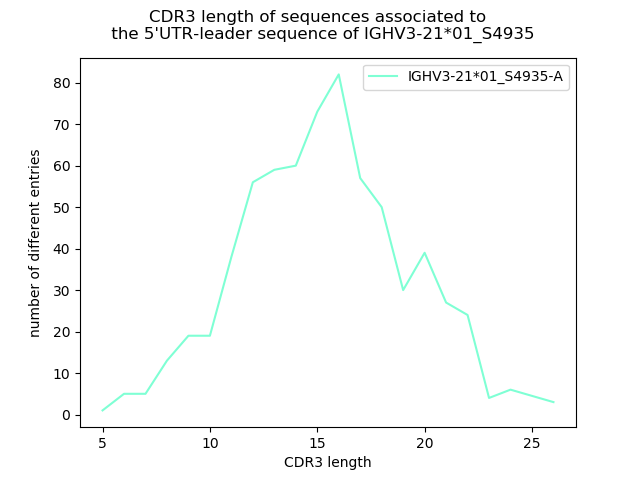

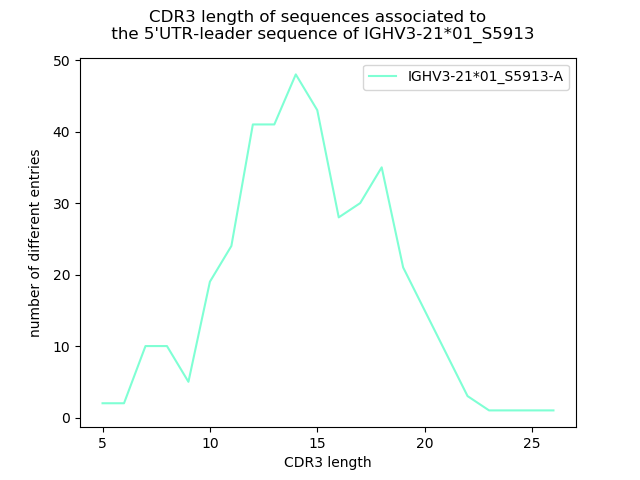

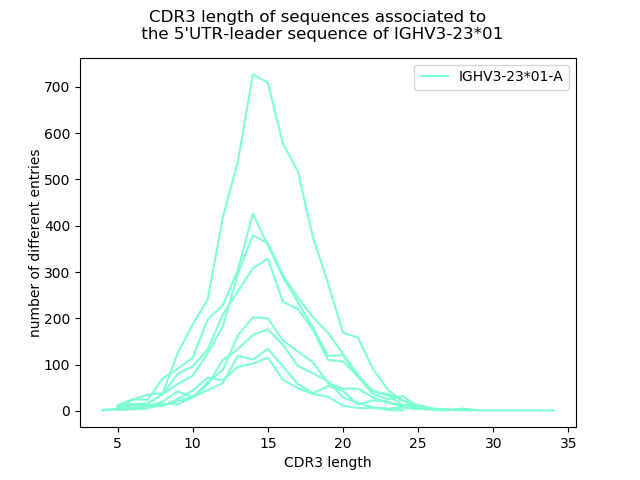

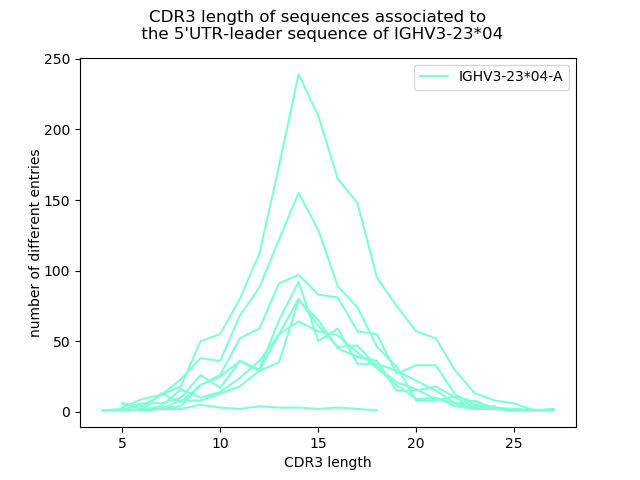

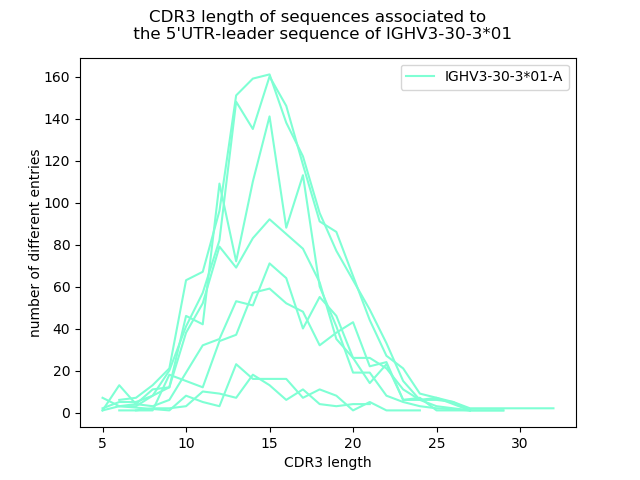

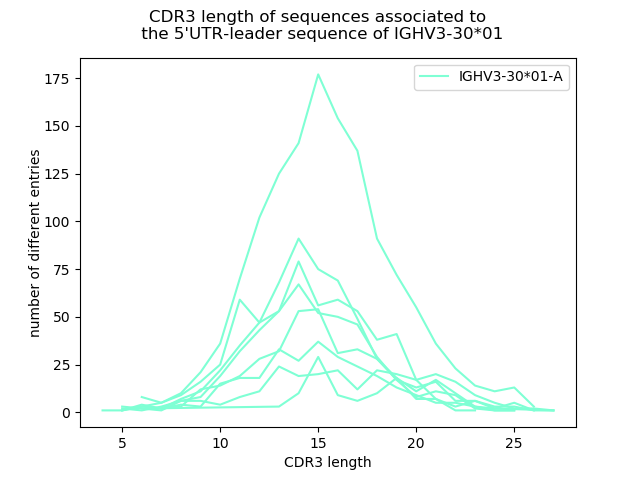

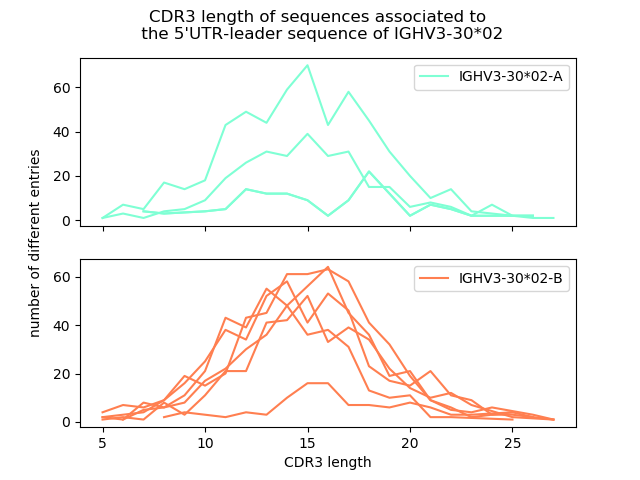

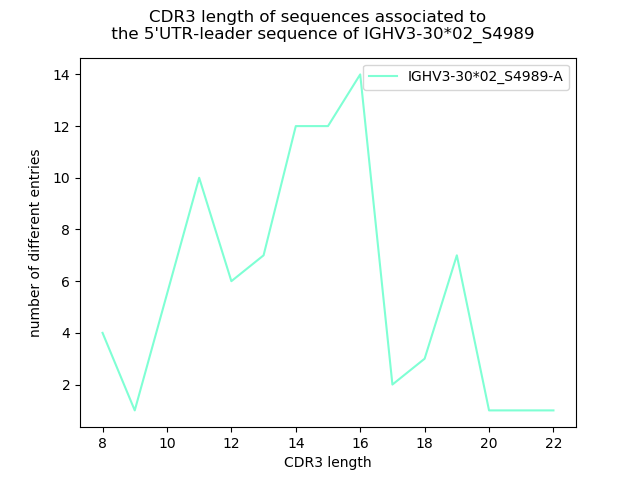

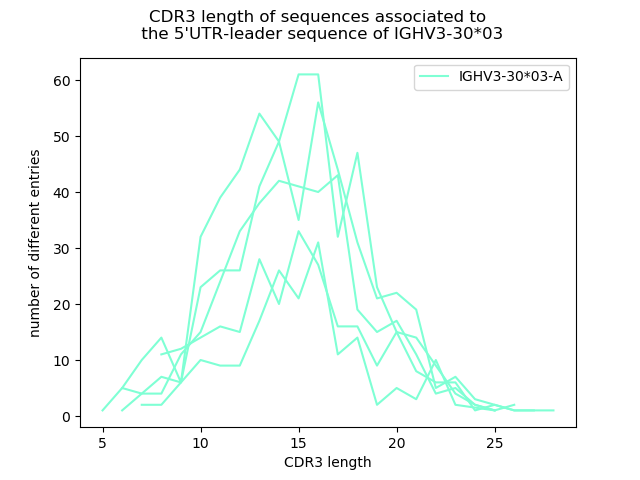

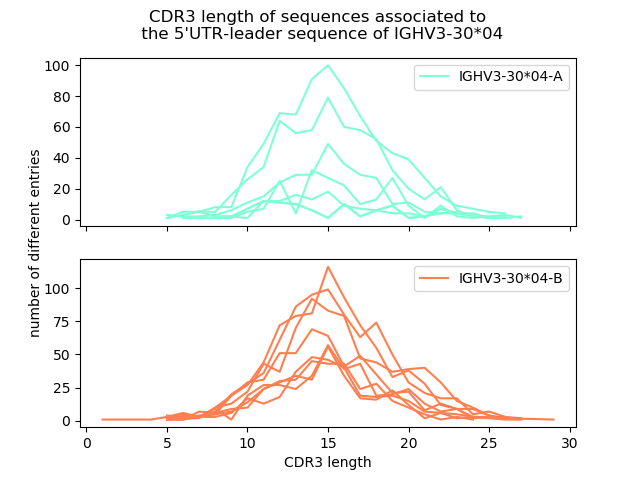

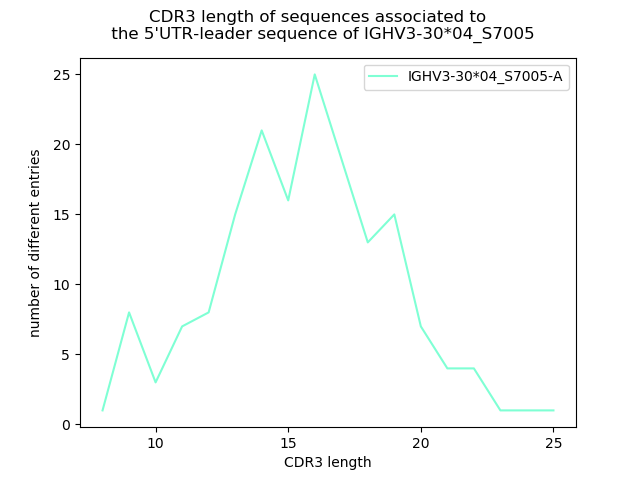

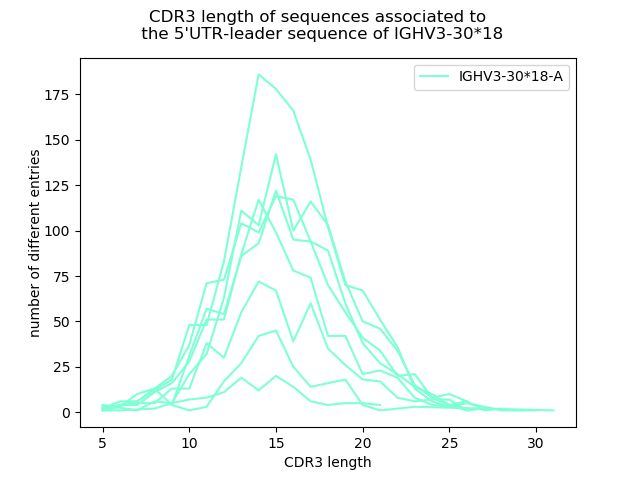

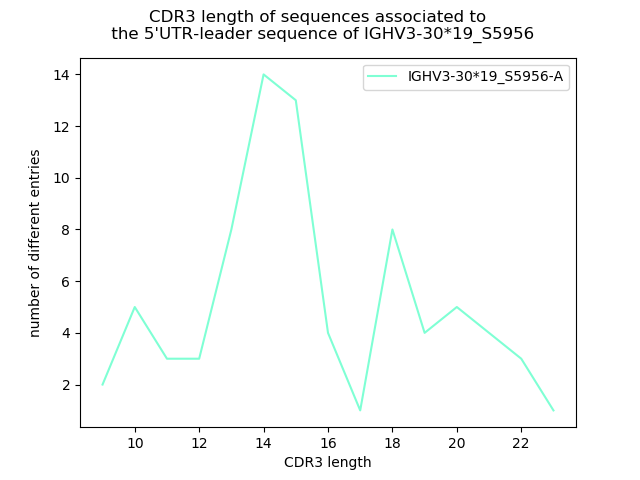

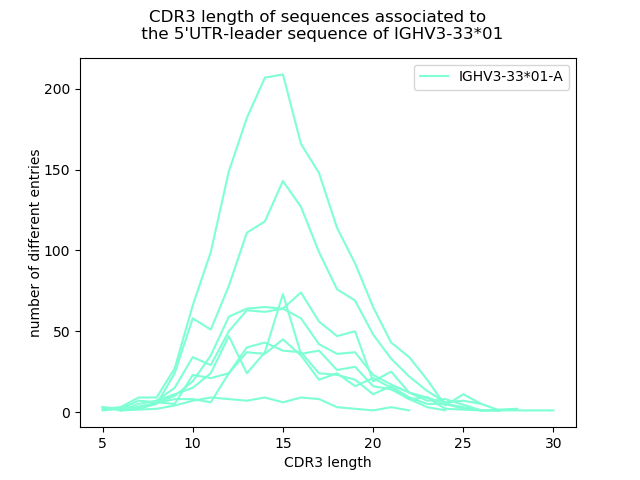

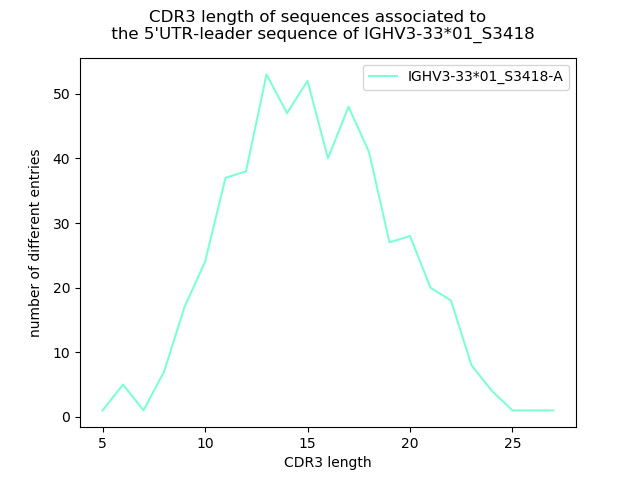

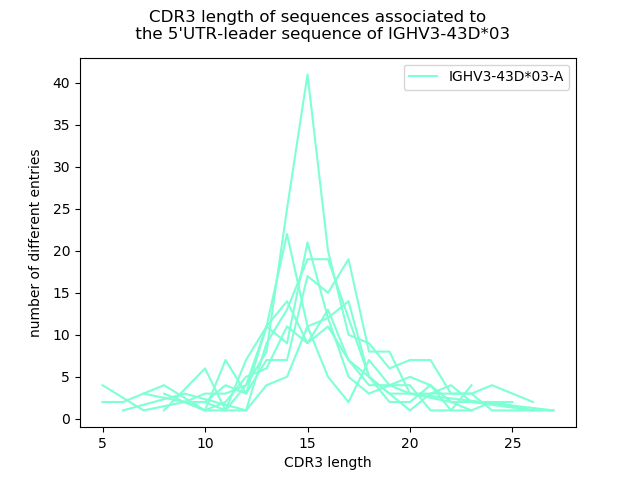

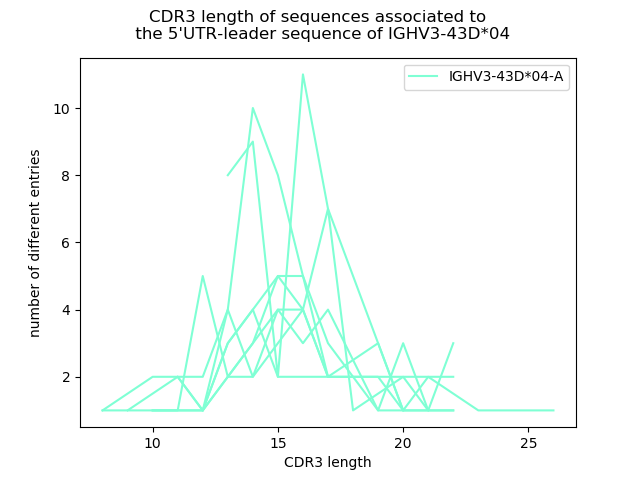

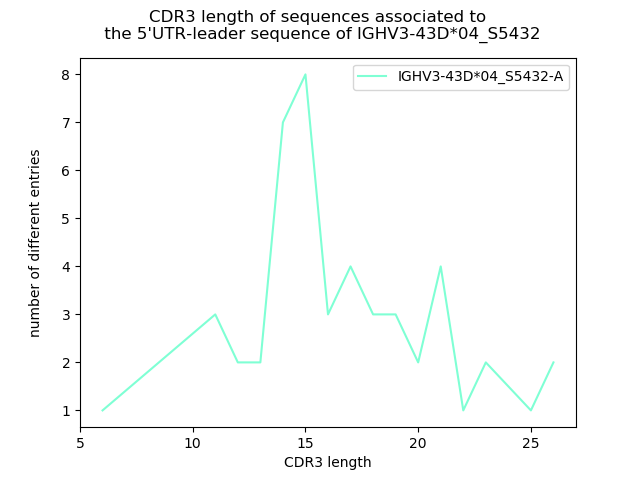

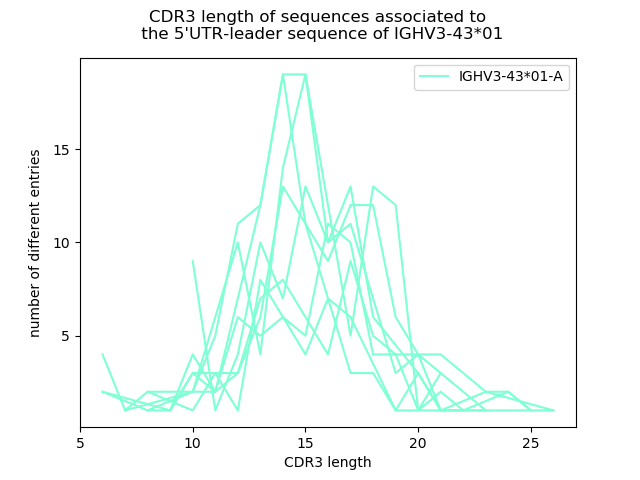

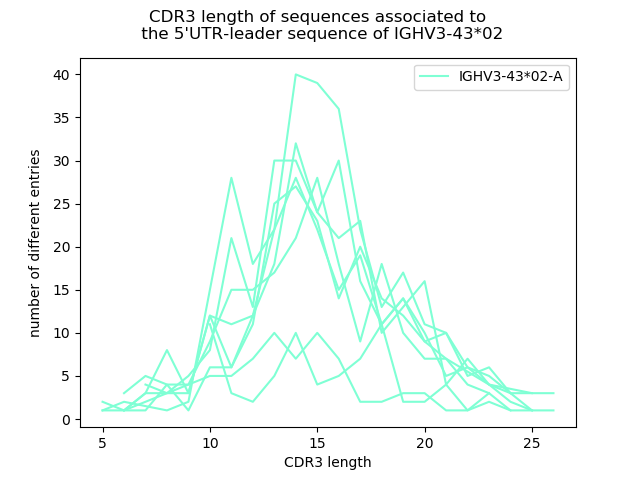

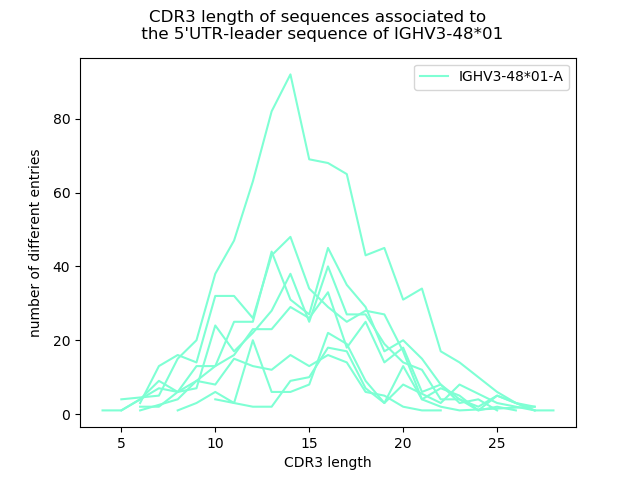

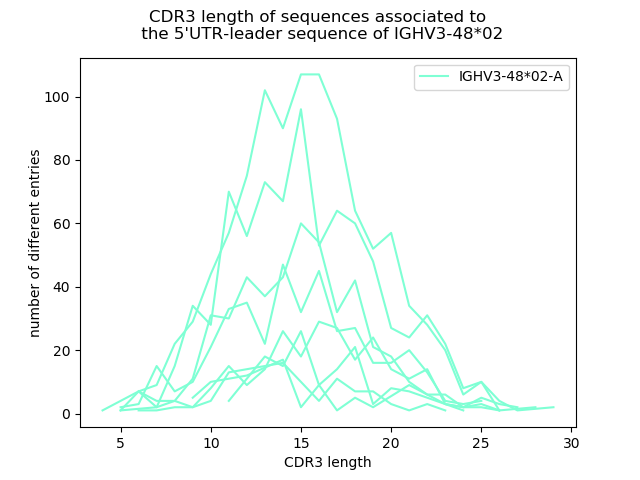

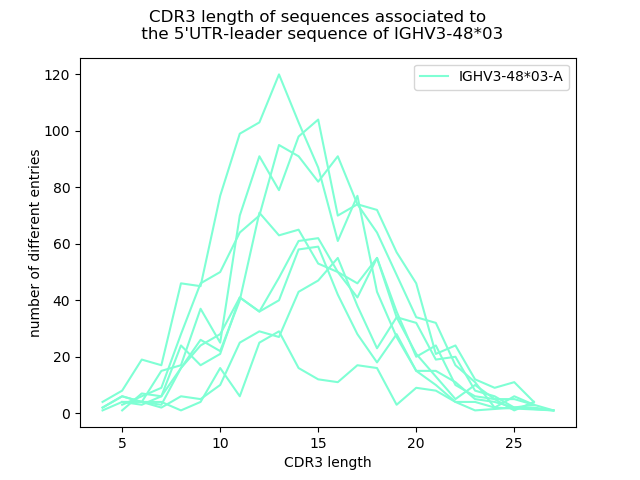

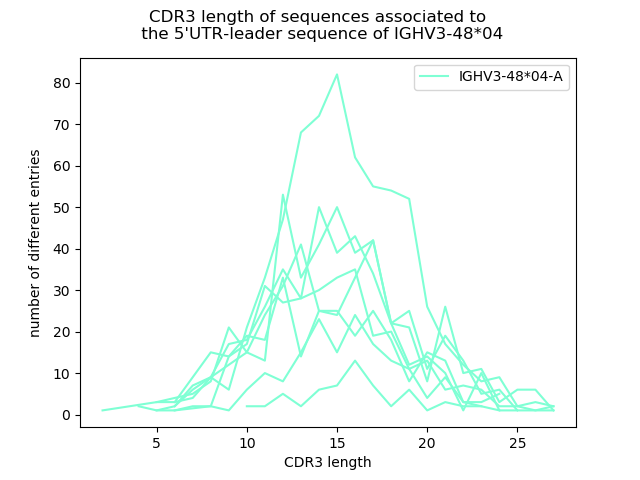

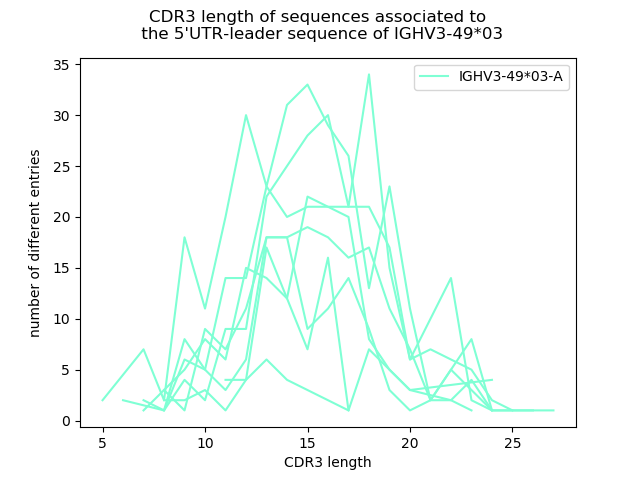

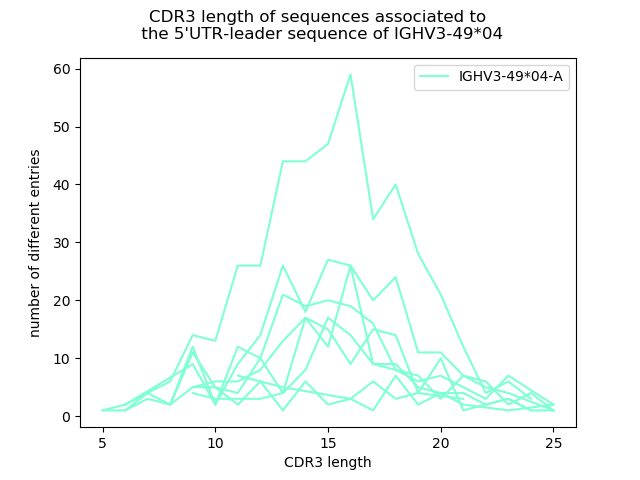

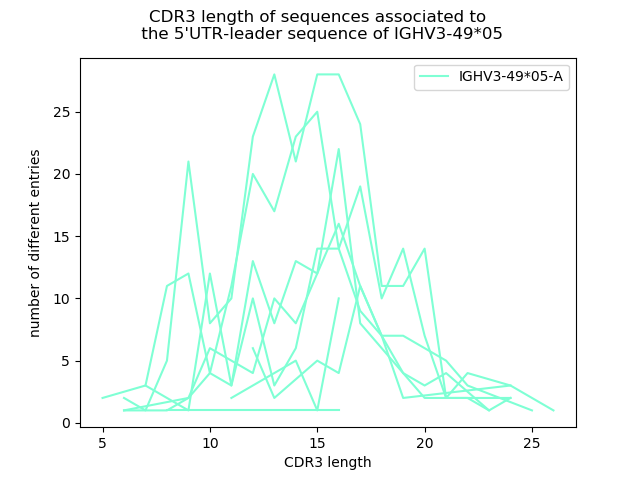

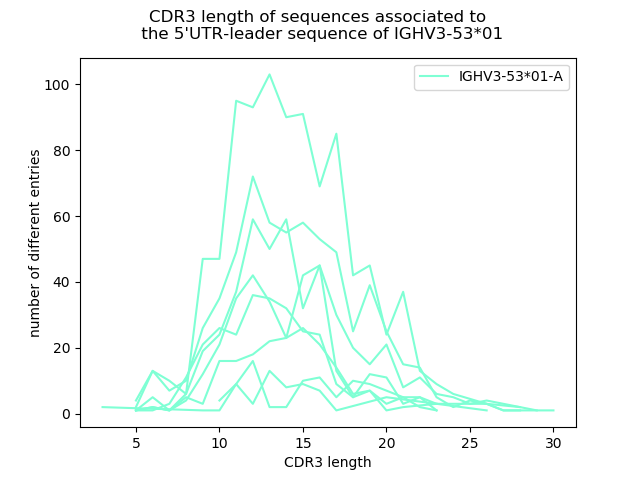

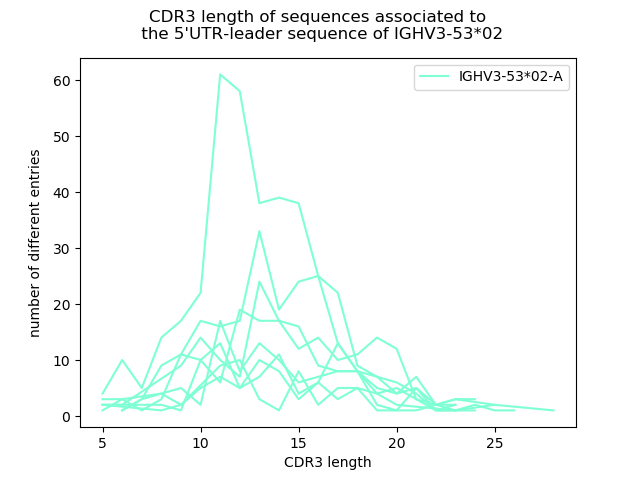

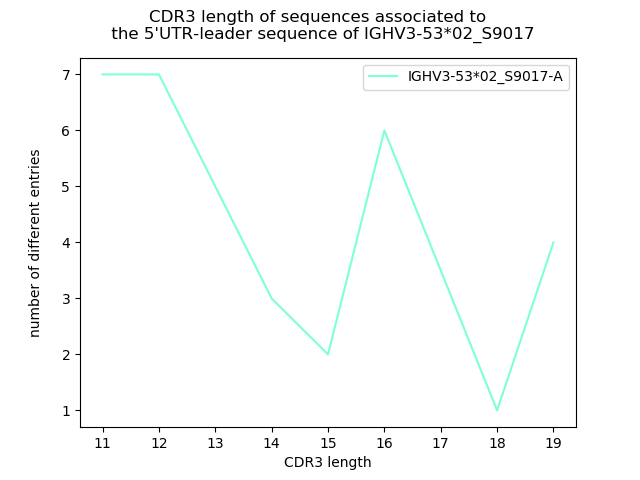

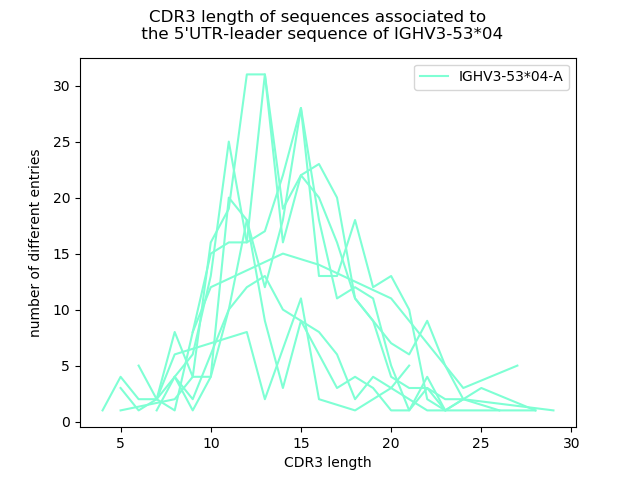

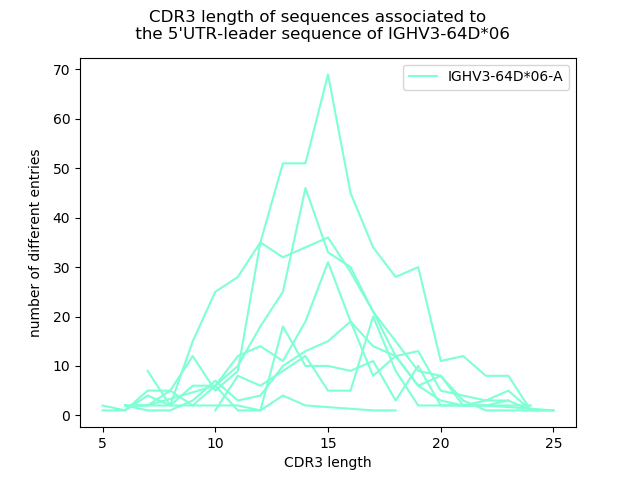

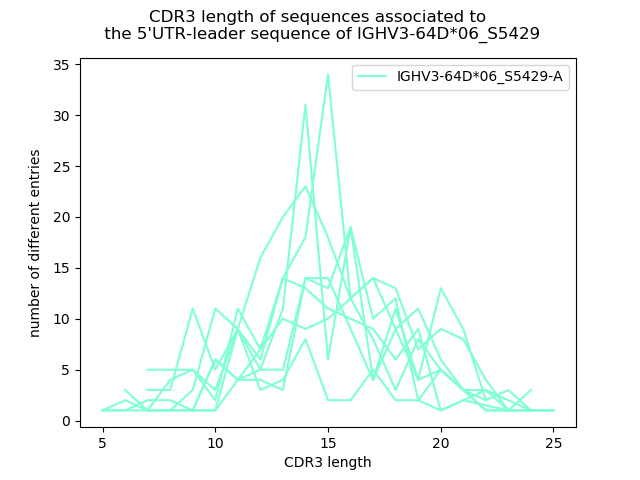

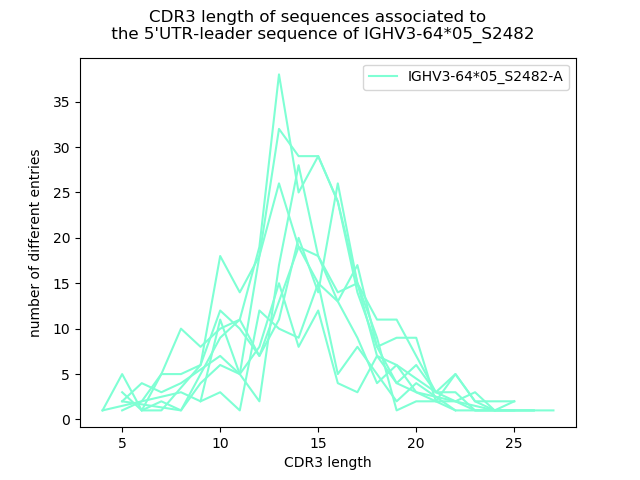

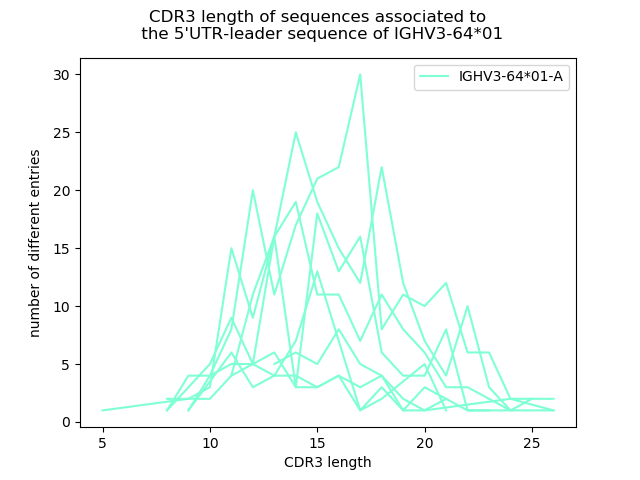

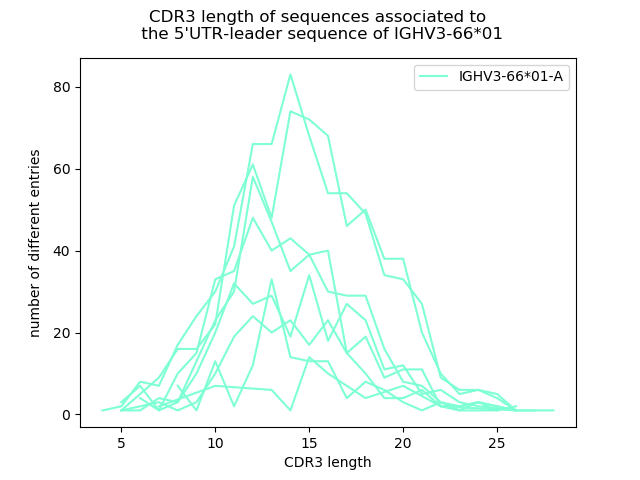

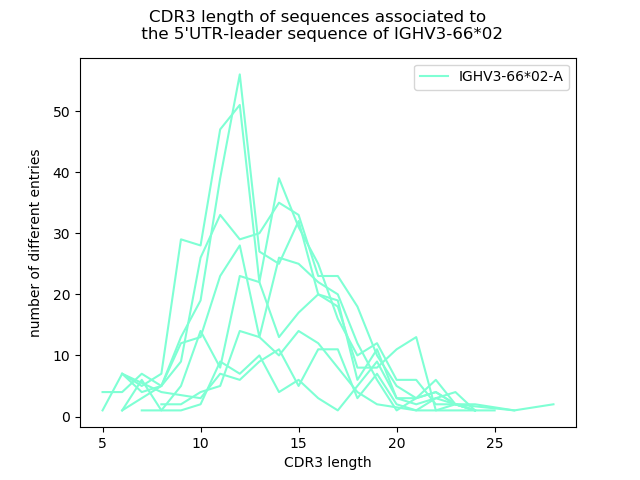

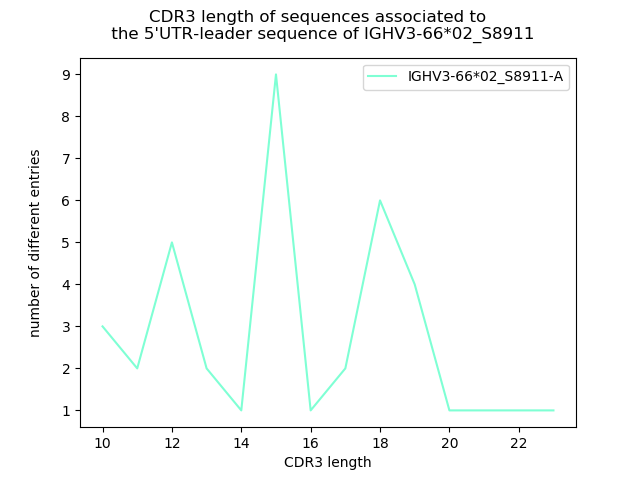

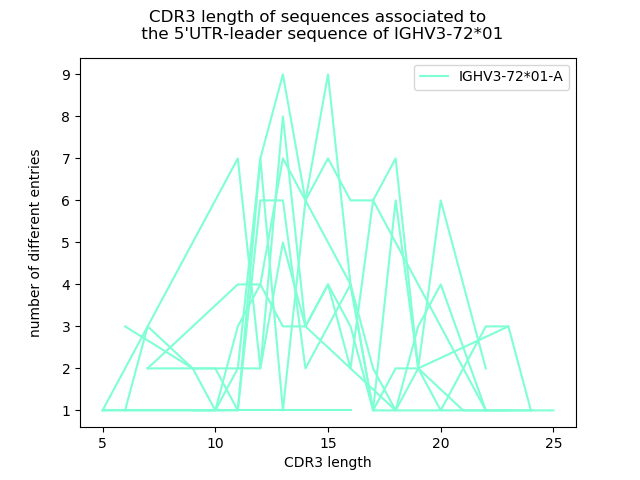

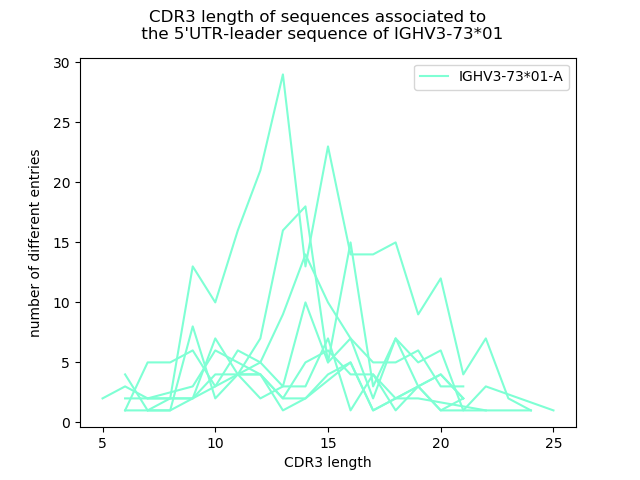

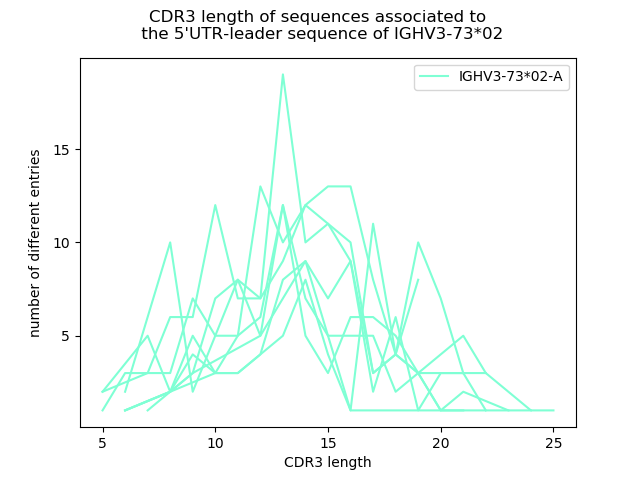

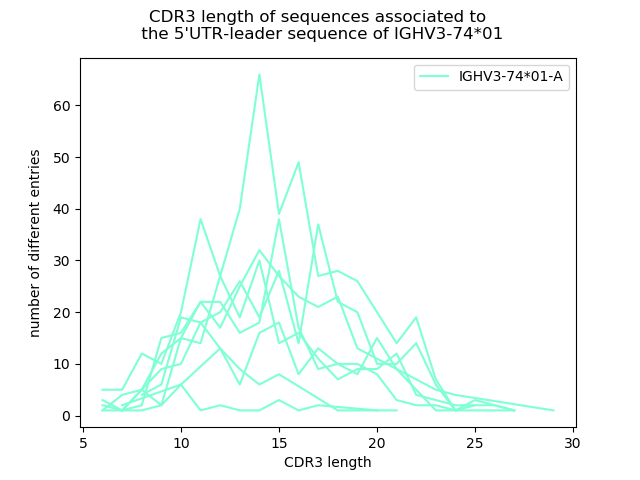

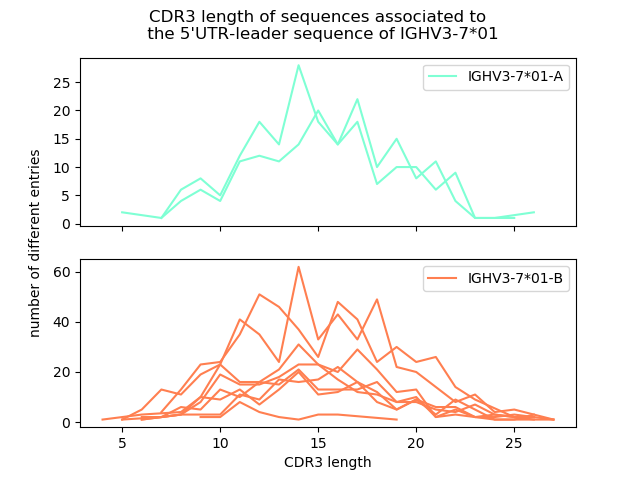

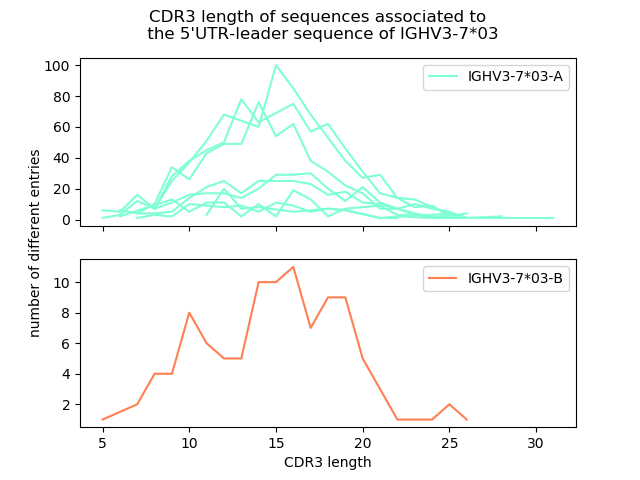

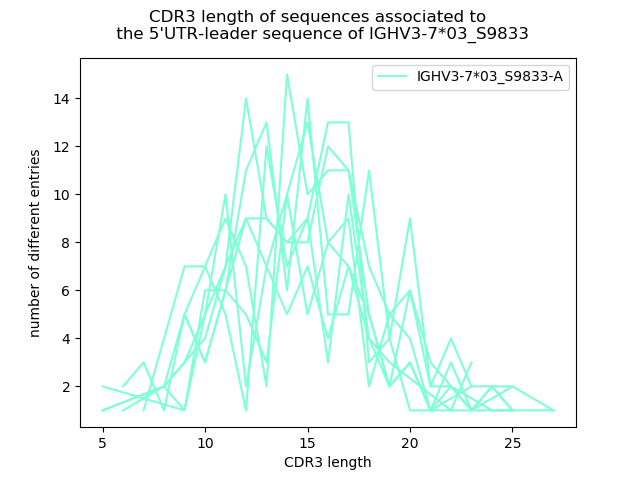

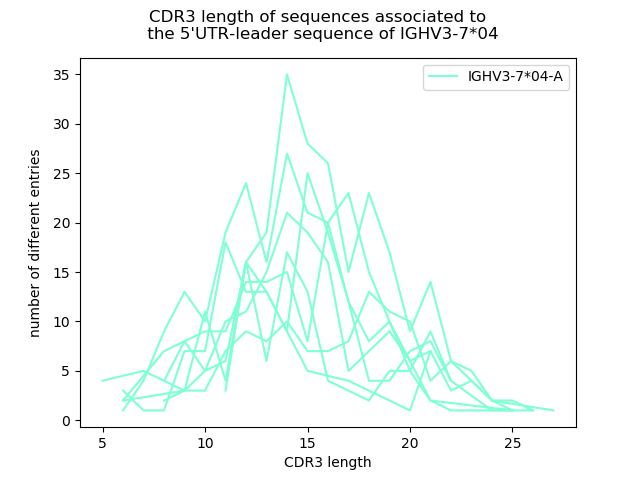

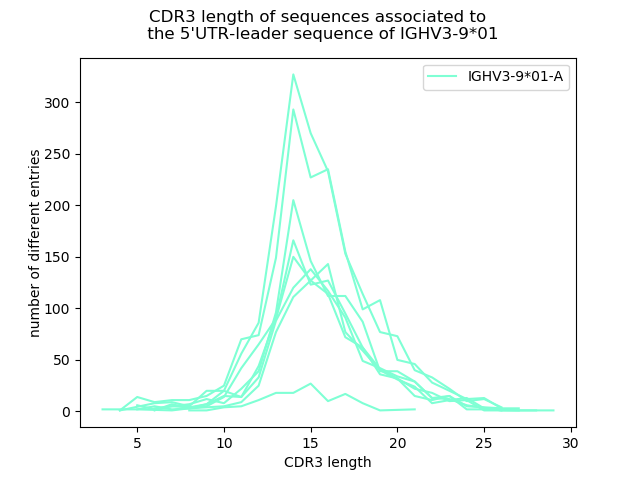

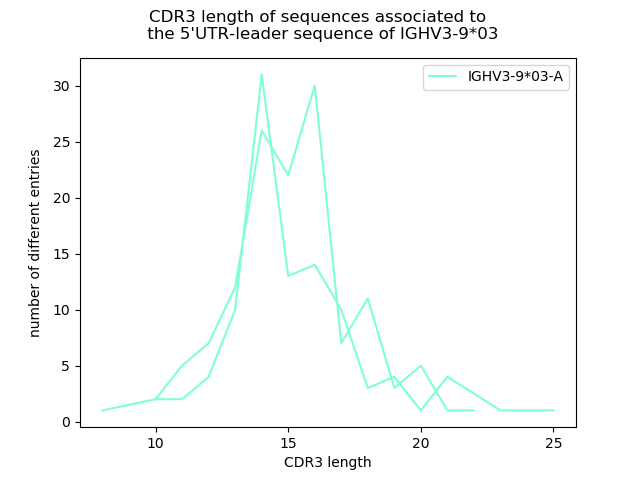

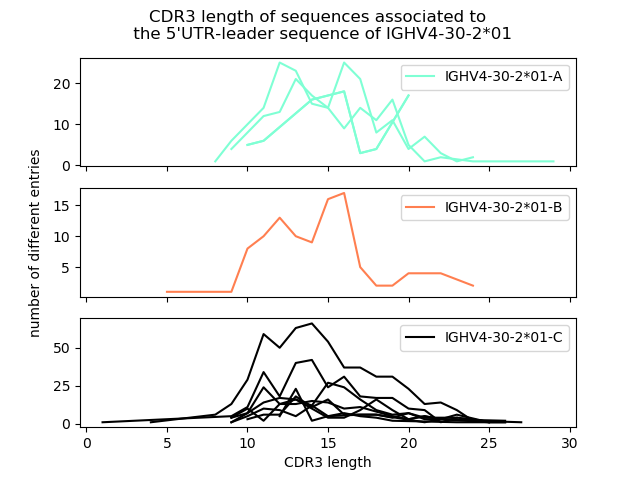

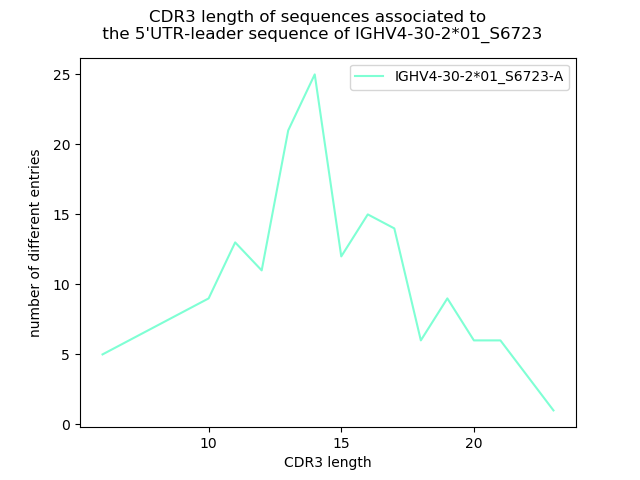

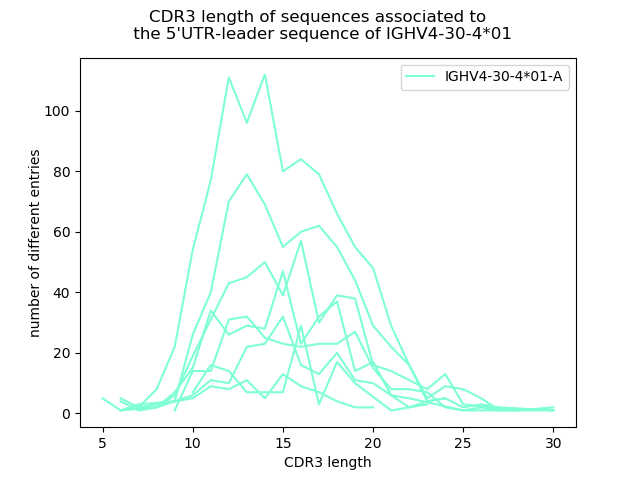

**Supplementary Figure 1.** Distribution patterns of CDR3 length encoded by transcripts associated to 5’UTR-leader sequences of different IGHV gene alleles. For each 5’UTR-leader sequence of a specific allele, the number of filtered reads in each length of CDR3 was counted to create the plots. Every line in the plots represents the CDR3 length distribution of one subject (at maximum 8 subjects were included in each plot).
