## Supplementary Table 3 for "Computational inference, validation, and analysis of 5’UTR-leader sequences of alleles of immunoglobulin heavy chain variable genes"

**Supplementary Table 3.** Alleles inferred as novel in relation to the database used in the study. Some genes have since been entered into the IMGT database, represents 3’-extensions of previously existing partial alleles, or have been provisionally approved by the Inferred Allele Review Committee (IARC). Evidence for novel alleles that have not previously been assessed is shown. The evidence include CDR3 length plots of unmutated reads (also available in Supplementary Figure 1), ORGDBstats plot obtained from VDJBase (30) (made available under a CC0 (Creative Commons 0) license), error plots visualizing the V_error distribution as determined by IgDiscover (9), haplotype data, and inferred gene sequences.

| Name of inferred allele | IMGT name | Allele name and mutation | Identified in (18) | Provisionally approved by IARC | Evidence of validity of previously unassessed alleles | | | | | | |
| --- | --- | --- | --- | --- | --- | --- | --- | --- | --- | --- | --- |
| IGHV1-3*01_S6816 | IGHV1-3*05 |  | yes |  |  | | | | | | |
| IGHV1-69*01_S8909 | IGHV1-69*19 |  | yes |  |  | | | | | | |
| IGHV1-69*04_S3852 |  | IGHV1-69*04 C184T | yes | yes° |  | | | | | | |
| IGHV1-69*06_S0471 |  | IGHV1-69*06 G240A | yes | yes° |  | | | | | | |
| IGHV1-69*08_S0647 | IGHV1-69*02 (3’-extension^&^) |  |  |  |  | | | | | | |
| IGHV2-26*02_S3803 | IGHV2-26*04 | IGHV2-26*02 C316T |  |  |  | | | | | | |
| IGHV2-70*04_S5392 |  | IGHV2-70*04 A14G or IGHV2-70*04D A14G, or IGHV2-70*17 T112C |  |  | A. CDR3 length  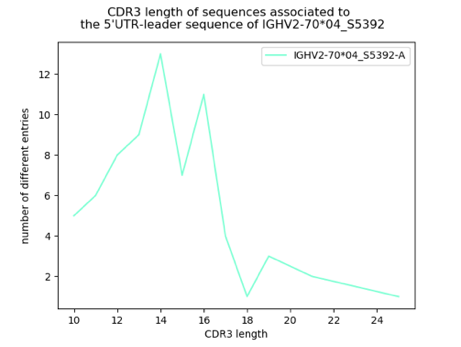 | | | B. Error plot  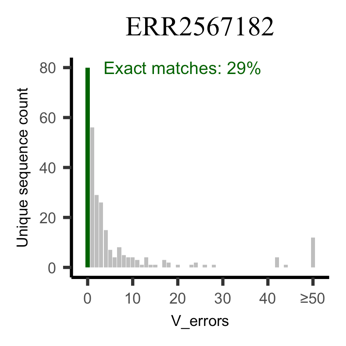  Exact matches, all genes^∞^: 23% | | | |
|  |  |  |  |  | C. Sequence of upstream region (excluding leader sequence intron) and the sequence encoding the final protein product.  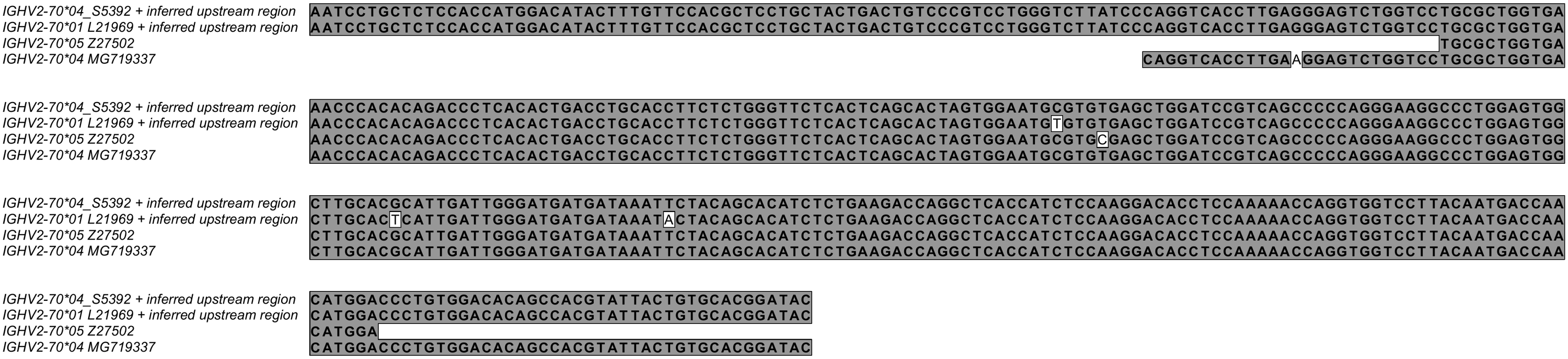 | | | | | | |
|  |  |  |  |  | Inferred by IgDiscover in ERR2567182 but it was not seen in VDJbase. Reads associated with this inferred germline gene allele are diverse in CDR3 length (A) and has a high level of unmutated reads (B), similar to the total frequency of exact matches in the same subject. Known alleles of IGHV2-70 carry either an A or a G at base position 14. No OGRDBStats is available for the allele in VDJbase. This genotype also carries IGHV2-70*01, an allele that features three differences from IGHV2-70*04_S5392. The inferred upstream regions of these two alleles are identical (C). The IMGT database also reports an incomplete allele (IGHV2-70*05) that in its defined sequence (that does not cover base 14) differs only in one position (base C116T) from IGHV2-70*04_S5392. Due to its incompleteness, IGHV2-70*05 does not perform well during the sequence alignment procedure that is part of the inference process. | | | | | | |
| IGHV3-13*01_S3164 |  | IGHV3-13*01 G290A T300A |  |  | A. CDR3 length  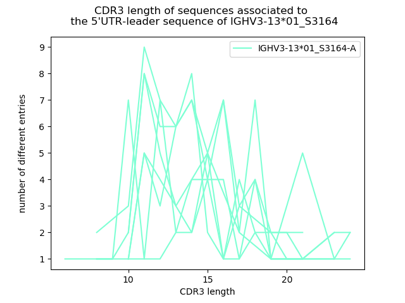 | | | | | | |
|  |  |  |  |  | 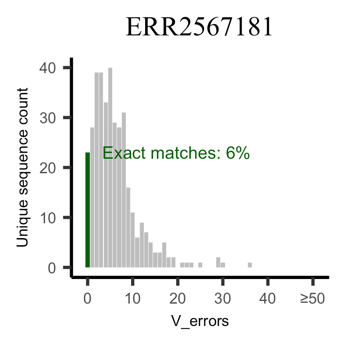  Exact matches, all genes^∞^:  6% | B. Error plots  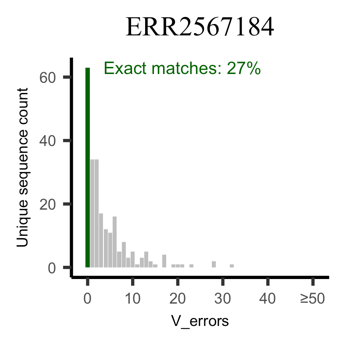  Exact matches, all genes^∞^: 25% | | | | 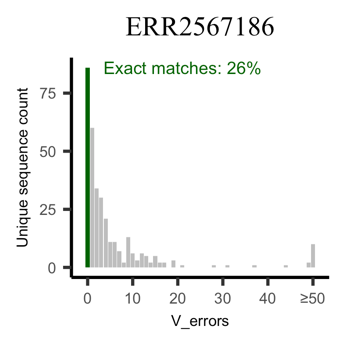  Exact matches, all genes^∞^: 28% | |
|  |  |  |  |  | 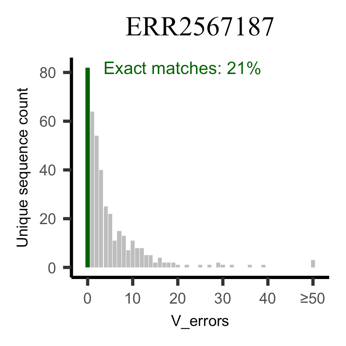  Exact matches, all genes^∞^: 23% | 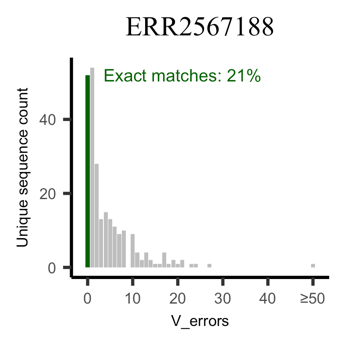  Exact matches, all genes^∞^: 23% | | | | 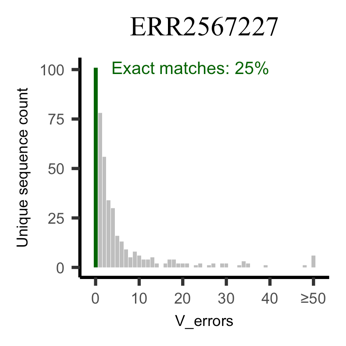 Exact matches, all genes^∞^: 26% | |
|  |  |  |  |  | 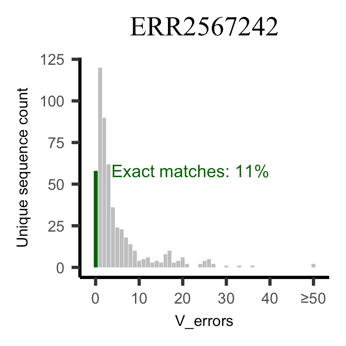  Exact matches, all genes^∞^:  12% | | | 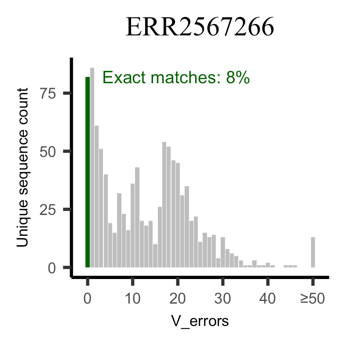  Exact matches, all genes^∞^:  17% | | | |
|  |  |  |  |  | C. ORGDBstats plot | | | | | | |
|  |  |  |  |  | ERR2567187  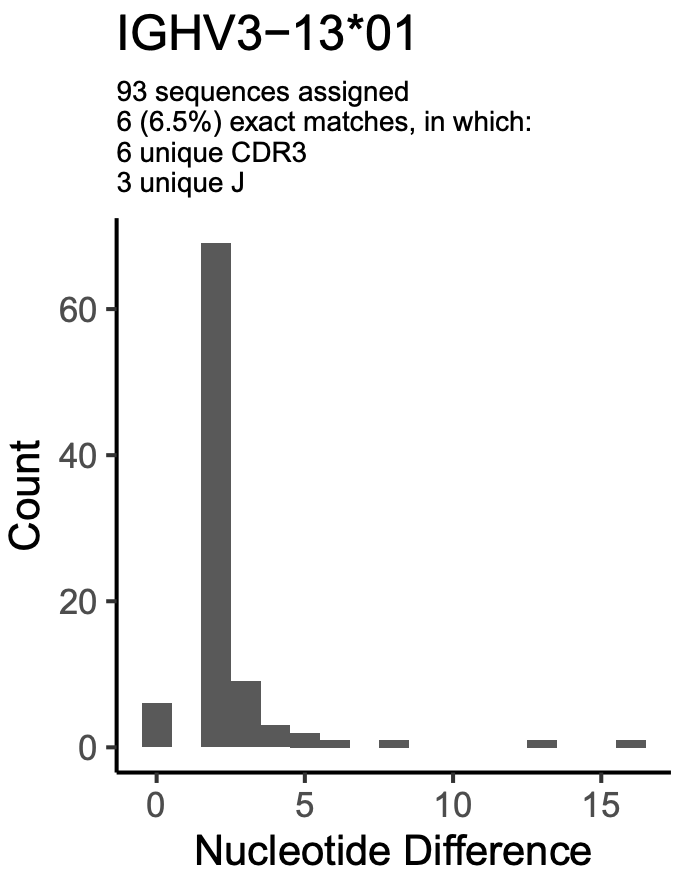  No other allele of IGHV3-13 is found in this subject. The gene is deleted from the other haplotype. | | ERR2567242  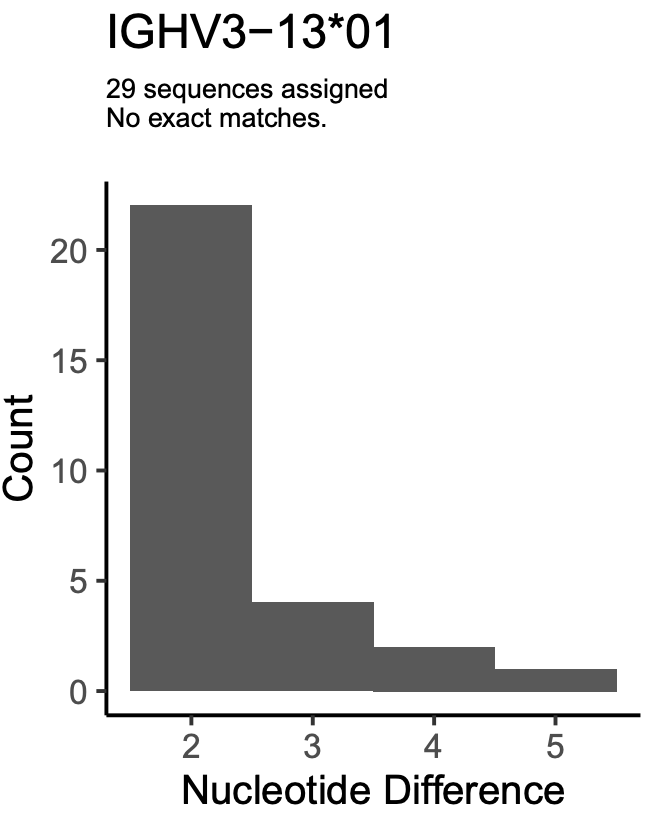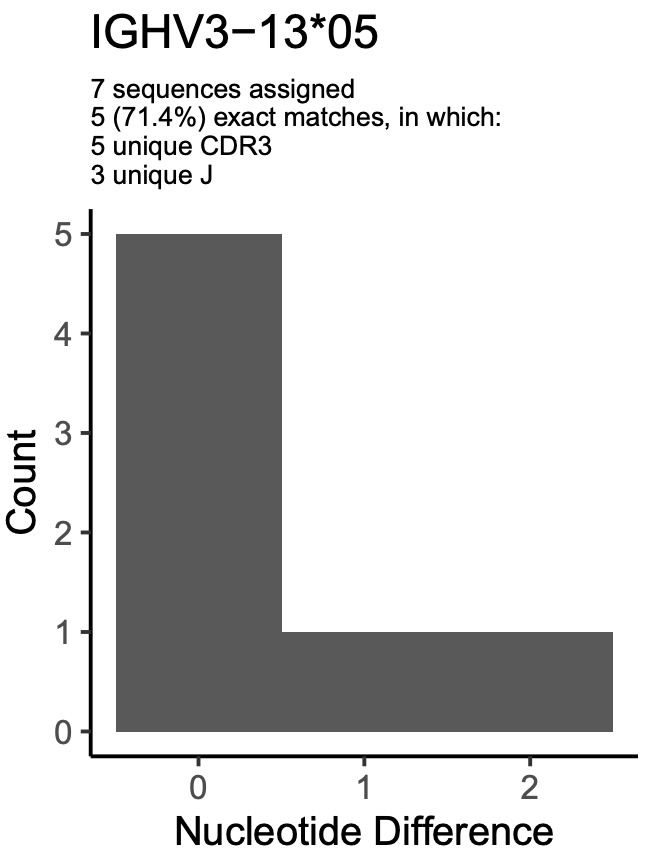 | | ERR2567266  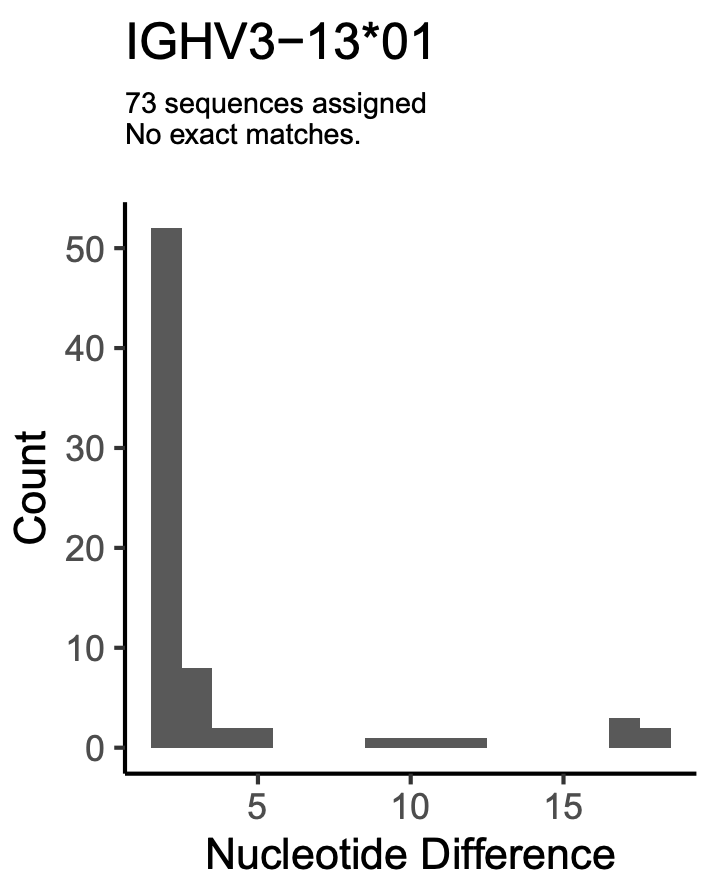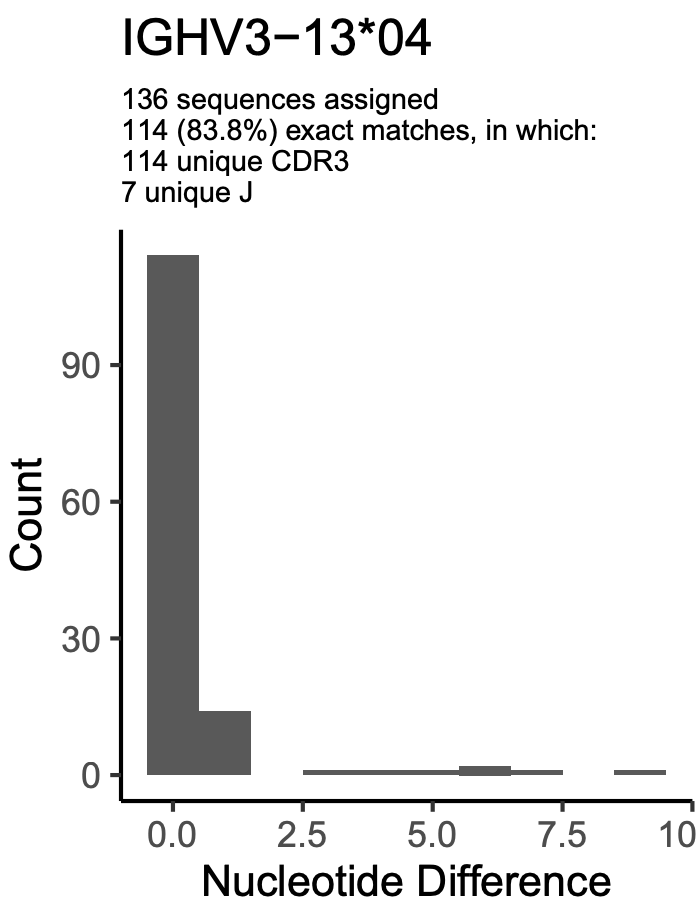 | | |
|  |  |  |  |  | D. Haplotyping data Data \| Germline gene/allele \| Haplotype \| \|; \| \| IGHJ6*02 \| IGHJ6*03 \| IGHJ6*04; ERR2567240 \| IGHV3-13*01 \| 17 \| 0 \|; \| IGHV3-13*01_S3164 \| 0 \| 16 \|; ERR2567242 \| IGHV3-13*01_S3164 \| \| 0 \| 43; \| IGHV3-13*05 \| \| 3 \| 0; ERR2567266 \| IGHV3-13*01_S3164 \| 25 \| 0 \|; \| IGHV3-13*04 \| 2 \| 25 \| | | | | | | |
|  |  |  |  |  | Inferred by IgDiscover in 8 samples. Reads associated with this inferred germline gene allele are diverse in CDR3 length (A). V_error profiles show high diversity between different subjects (B). 5 of 8 individuals, however, have an exact match rate of >20%, and the level of unmutated reads are similar to the total frequency of exact matches in the same subject, for all individuals but ERR2567266. VDJbase identify IGHV3-13*01 and not the novel variant, but the OGRDBStats reports identifies at least two mutations in such reads (while sequences of another allele of the same gene are mostly unmutated) (C). Reads haplotype as expected for alleles that are located on different chromosomes in ERR2567240, ERR2567242, and ERR2567266 (D). | | | | | | |
| IGHV3-21*01_S4935 |  | IGHV3-21*01 A184G T190A A191C | yes | yes§ |  | | | | | | |
| IGHV3-21*01_S5913 | IGHV3-21*06 |  |  |  |  | | | | | | |
| IGHV3-30*02_S4989 |  | IGHV3-30*02 G49A |  |  | A. CDR3 length  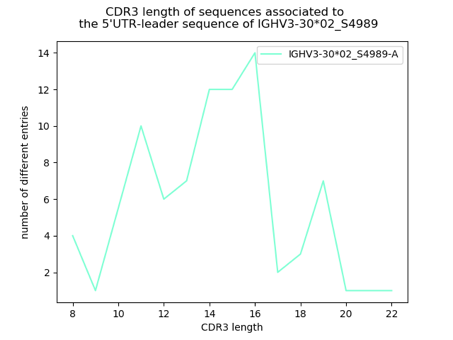 | | B. Error plot  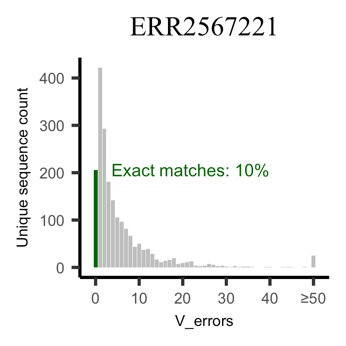  Exact matches, all genes^∞^: 10% | | | | C. ORGDBStats plot  P11_I25  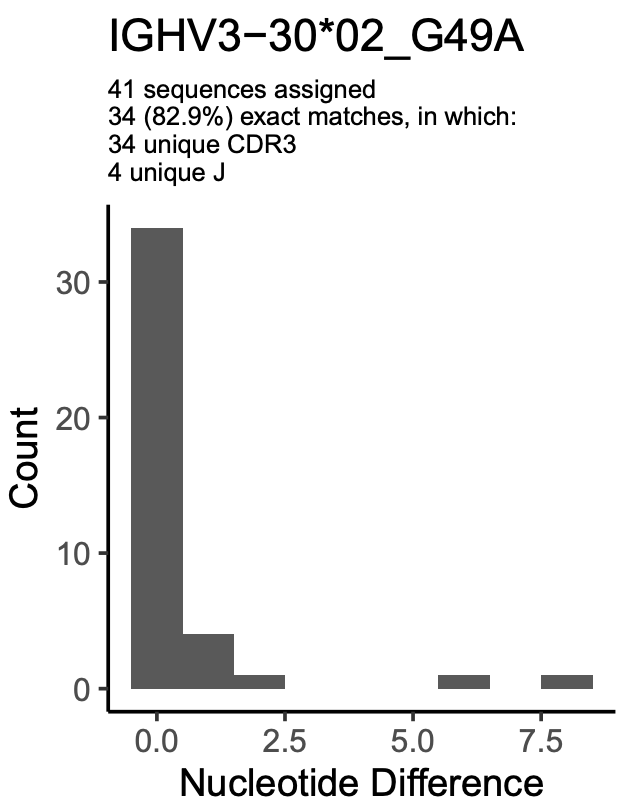 |
|  |  |  |  |  | D. Haplotyping data Data \| Germline gene/allele \| Haplotype \|; \| \| IGHJ6*02 \| IGHJ6*03; ERR2567221 \| IGHV3-30*02_S4989 \| 50 \| 0; \| IGHV3-30*04 \| 46 \| 0; \| IGHV3-30*18 \| 35 \| 37; \| IGHV3-30-3*01 \| 4 \| 8; \| IGHV3-33*01 \| 0 \| 12 | | | | | | |
|  |  |  |  |  | E. Sequence of upstream region (excluding leader sequence intron) and the sequence encoding the final protein product.  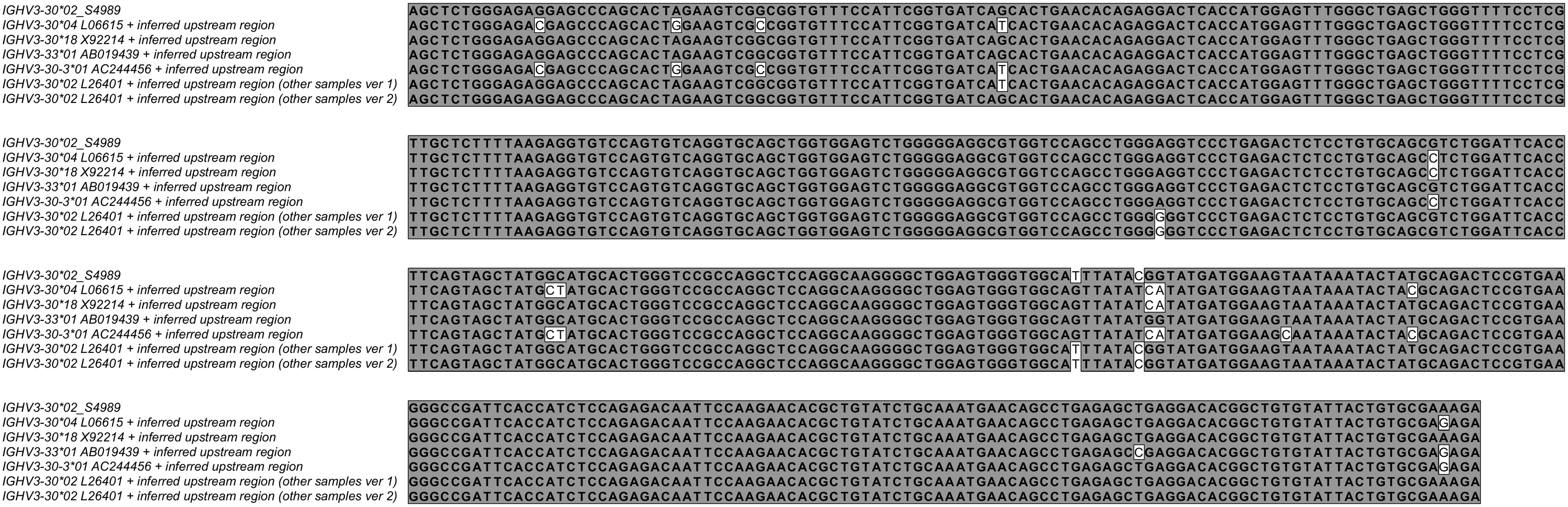 | | | | | | |
|  |  |  |  |  | Inferred by IgDiscover in ERR2567221. Reads associated with this inferred germline gene allele are diverse in CDR3 length (A). 10% of the assigned sequences in this subject are unmutated, similar to the total frequency of exact matches in the same subject (B). The allele has not been identified in the P1 study in VDJbase but it has been identified in a sample of study P11 in VDJbase (C). The subject has a complex haplotype (also including IGHV3-30*04, IGHV3-30*18, IGHV3-30-3*01 and IGHV3-33*01, but not IGHV3-30*02) in this complex part of the IGHV locus (D). The allele’s inferred upstream region is identical to those of IGHV3-30*18 and IGHV3-33*01 that also exist in the haplotype, but these alleles differ by 5 and 4 bases, respectively, in the coding region from the sequence of IGHV3-30*02_S4989 (E). The inferred allele cannot be a result of a single cross-over event during the PCR-process between products of other alleles of these genes. | | | | | | |
| IGHV3-30*04_S7005 |  | IGHV3-30*04 C201T G317A |  |  | A. CDR3 length  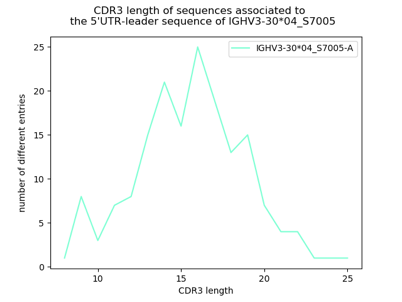 | | | B. Error plot  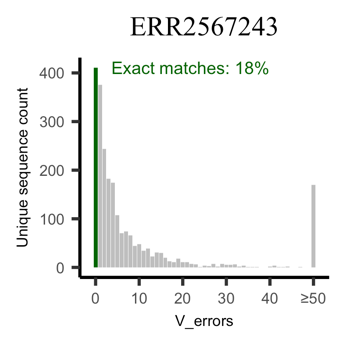  Exact matches, all genes^∞^: 24% | | | |
|  |  |  |  |  | C. Haplotyping data Data \| Germline gene/allele \| Haplotype \|; \| \| IGHJ6*02 \| IGHJ6*03; ERR2567243 \| IGHV3-30*04_S7005 \| 0 \| 63; \| IGHV3-30*18 \| 253 \| 128; \| IGHV3-30-3*01 \| 195 \| 111 | | | | | | |
|  |  |  |  |  | D. 5’UTR-leader sequence-IGHV  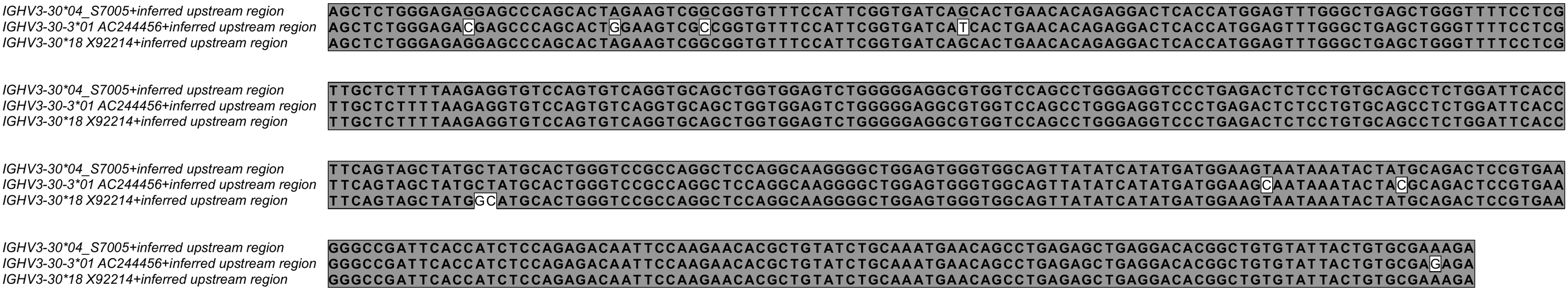 | | | | | | |
|  |  |  |  |  | Inferred by IgDiscover in ERR2567243. Reads associated with this inferred germline gene allele are diverse in CDR3 length (A) and has a high level of unmutated reads (B), however somewhat lower than the frequency of exact matches in the same subject. The subject has a complex haplotype (C) in this complex part of the IGHV locus. Two copies of IGHV3-30*18, and IGHV3-30-3*01, each, and one copy of IGHV3-30*04_S7005, but no copy of IGHV3-33 is identified in this subject. The upstream region of IGHV3-30*04_S7005 is identical to that of IGHV3-30*18 but not to that of IGHV3-30-3*01 (D). The inferred allele cannot be a result of a single cross-over event during the PCR-process between products of other alleles of these genes. Haplotyping shows that IGHV3-30*04_S7005 only exists on one of the haplotypes where IGHV3-30*18 and IGHV3-30-3*01 are found. | | | | | | |
| IGHV3-30*19_S5956 |  | IGHV3-30*19 T189C | yes | yes** | A. CDR3 length  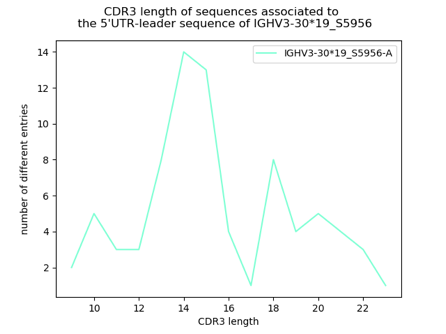 | | B. Error plot  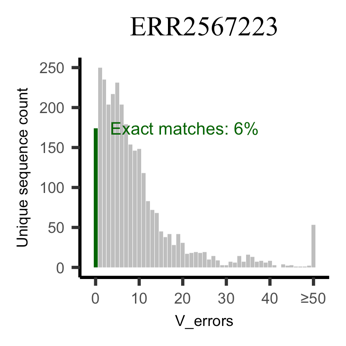  Exact matches, all genes^∞^: 8% | | | | C. ORGDBStats plot  ERR2567223  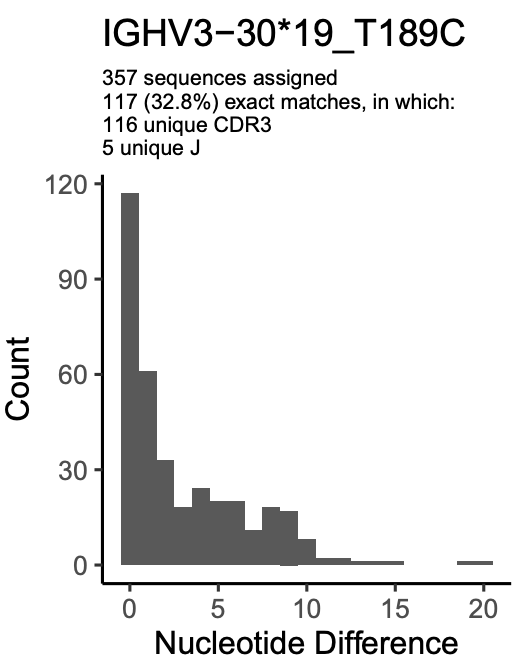 |
|  |  |  |  |  | D. Haplotyping data Data \| Germline gene/allele \| Haplotype \|; \| \| IGHJ6*02 \| IGHJ6*03; ERR2567223 \| IGHV3-30*18 \| 25 \| 4; \| IGHV3-30*19_S5956 \| 31 \| 0; \| IGHV3-30-3*01 \| 0 \| 10; \| IGHV3-33*01 \| 15 \| 13 | | | | | | |
|  |  |  |  |  | Inferred by IgDiscover in ERR2567223. Reads associated with this inferred germline gene allele are diverse in CDR3 length (A), and has a profile where sequences with errors ranging from 0 to 10 in the V region are found in similar levels (B). However, the level of unmutated reads is similar to the total frequency of exact matches in the same subject. This allele is also reported in VDJbase, to a large extent in unmutated form (C). This allele is part of the complex set of genes IGHV3-30, IGHV3-30-3, IGHV3-30-5, and IGHV3-33. Haplotyping data demonstrate that it resides on the same haplotype as IGHV3-30*18 and IGHV3-33*01, while the other haplotype carries IGHV3-30*18, IGHV3-30-3*01, and IGHV3-33*01. Its 5’UTR-leader sequence is identical to that of IGHV3-30-3*01, but these alleles segregate onto different haplotypes. It is thus possible that both IGHV3-30-3*01 and IGHV3-30*19_S5956 reside in gene IGHV3-30-3. | | | | | | |
| IGHV3-33*01_S3418 |  | IGHV3-33*01 G75C | yes | yes† |  | | | | | | |
| IGHV3-43D* 04_S5432 |  | IGHV3-43D*04 G4A |  |  | A. CDR3 length  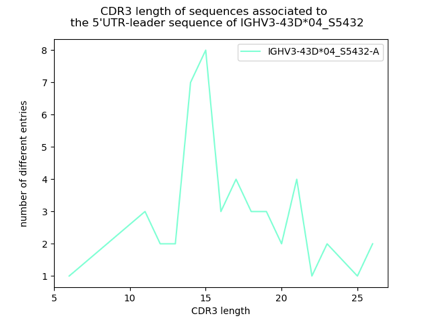 | | B. Error plot  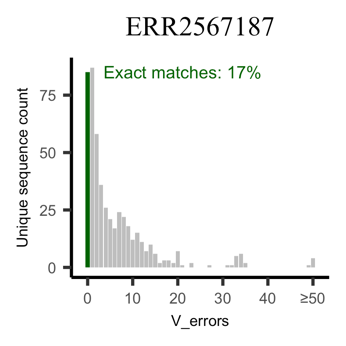  Exact matches, all genes^∞^: 23% | | | | C. ORGDBstats plot ERR2567187  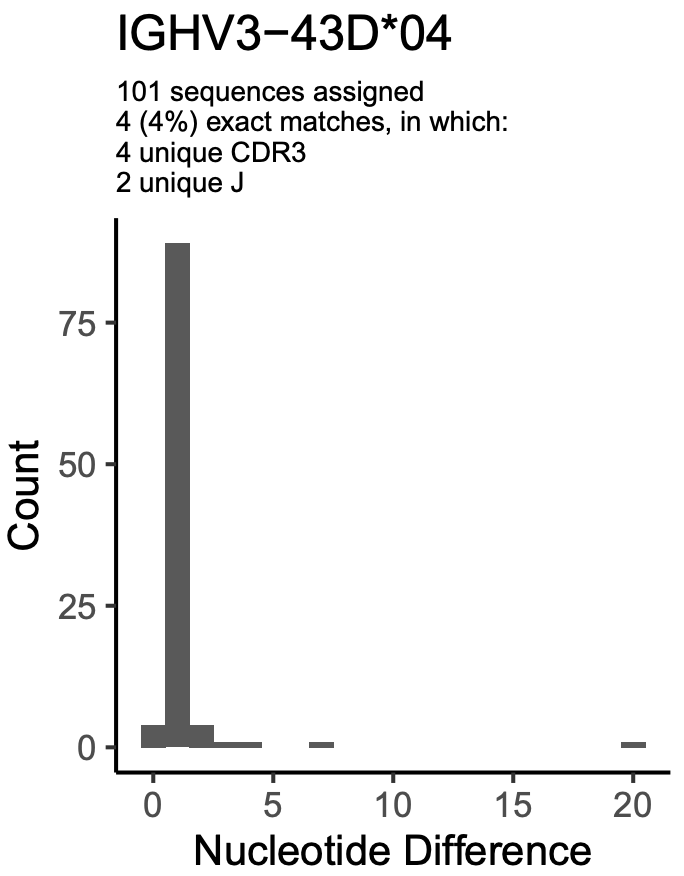 |
|  |  |  |  |  | D. Haplotyping data Data \| Germline gene/allele \| Haplotype \|; \| \| IGHJ6*02 \| IGHJ6*03; ERR2567187 \| IGHV3-43*01 \| 0 \| 20; \| IGHV3-43D*04_S5432 \| 49 \| 0 | | | | | | |
|  |  |  |  |  | Inferred by IgDiscover in ERR2567187. Reads associated with this inferred germline gene allele are diverse in CDR3 length (A) and has a high level of unmutated reads (B), however somewhat lower than the frequency of exact matches in the same subject. VDJbase identifies IGHV3-43D*04 but these sequences in contrast to sequences representing other alleles, almost all carry at least 1 mutation (C). The allele exists on the same haplotype as IGHV4-38-2 (D). The upstream region of this allele is identical to that of other alleles of IGHV3-43D but not to those of alleles of IGHV3-43. Altogether this suggests that IGHV3-43D*04_S5432 is an allele of IGHV3-43D and not of IGHV3-43. No other allele of IGH3-43D is found in the genotype. | | | | | | |
| IGHV3-53*02_S9017 |  | IGHV3-53*02 C259T |  |  | A. CDR3 length  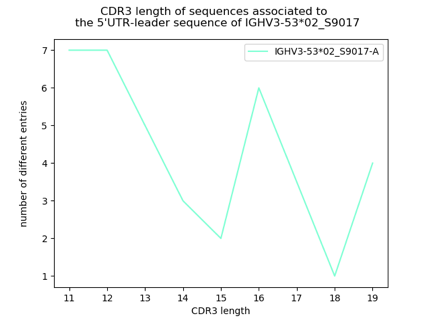 | | B. Error plot  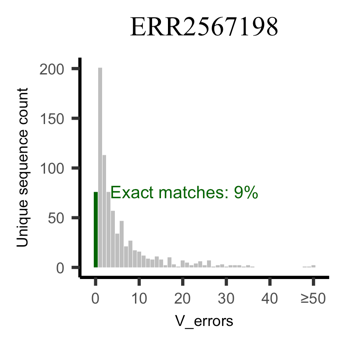  Exact matches, all genes^∞^: 10% | | | | C. OGRDBStats plot  ERR2567198  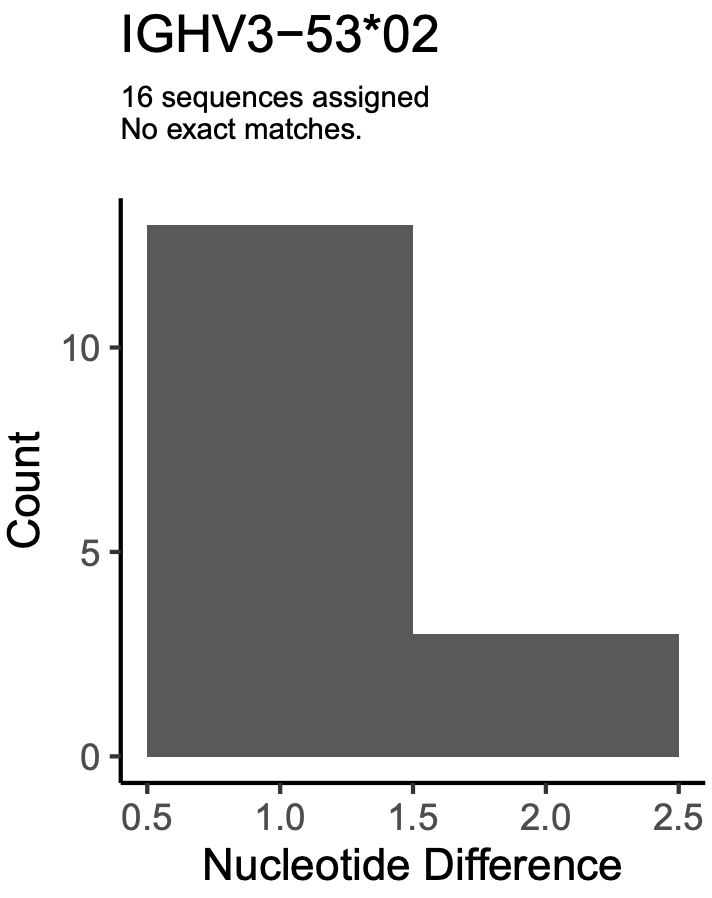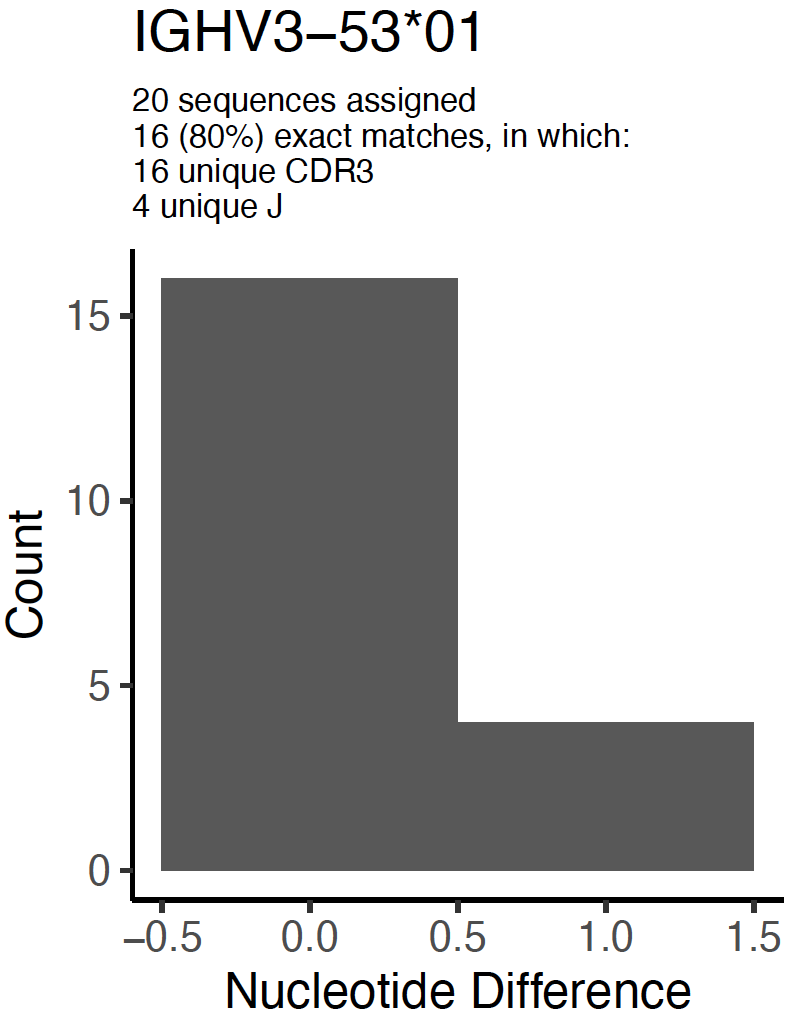 |
|  |  |  |  |  | Inferred by IgDiscover in ERR2567198. Reads associated with this inferred germline gene allele are diverse in CDR3 length (A), but most commonly carries 1 error in the V region (B). Unmutated reads, however, are present in a level similar to the total frequency of exact matches in the same subject. VDJbase identifies IGHV3-53*02 in this genotype, but these sequences, in contrast to sequences representing IGHV3-53*01 of the same subject, carry at least 1 mutation (C). The upstream region of IGHV3-53*02_S9017 differ in one base from the upstream region of IGHV3-53*01 that also exists in the genotype. A C259T variant does not exist in other known alleles of IGHV3-53 or alleles of the closely related gene IGHV3-66.  Population data: The T variant of SNP rs371918802 is very rare (<0.1%) in human populations (Ensembl release 103) | | | | | | |
| IGHV3-64*05_S2482 | IGHV3-64D*09 |  | yes |  |  | | | | | | |
| IGHV3-64D* 06_S5429 | IGHV3-64D*08 |  | yes |  |  | | | | | | |
| IGHV3-66*02_S8911 |  | IGHV3-66*02 G303A |  |  | A. CDR3 length  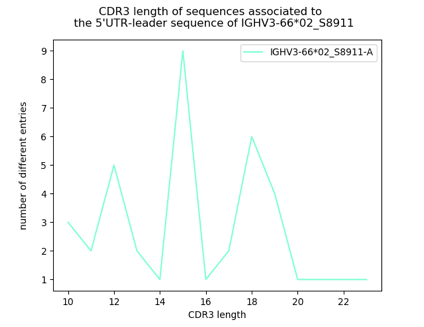 | | B. Error plot  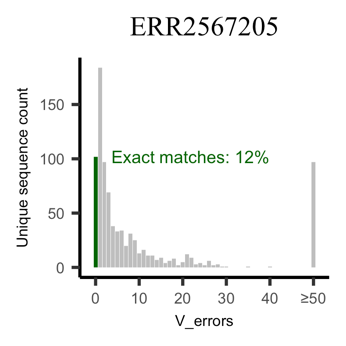  Exact matches, all genes^∞^: 12% | | | | C. OGRDBStats plot  ERR2567205  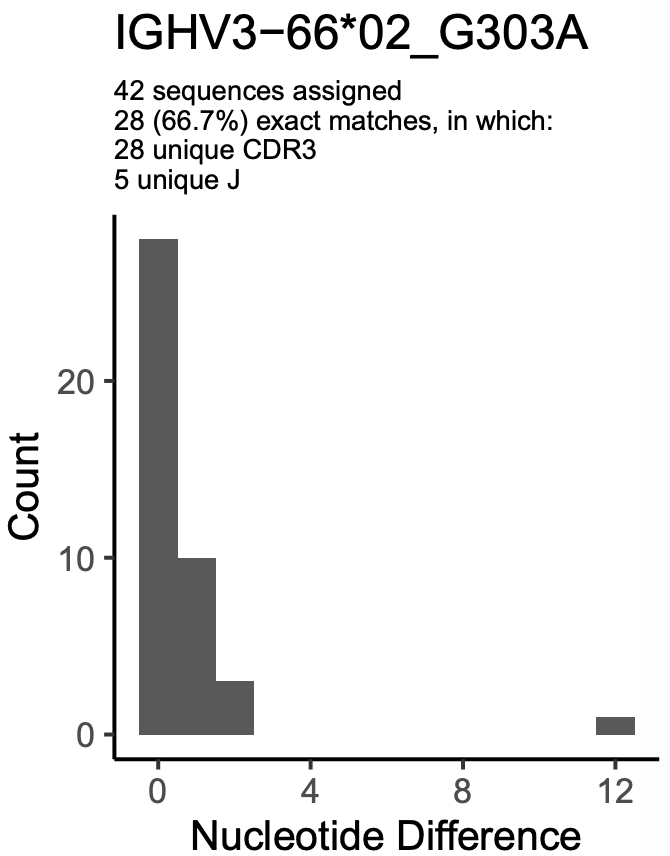 |
|  |  |  |  |  | Inferred by IgDiscover in ERR2567205. Reads associated with this inferred germline gene allele are diverse in CDR3 length (A), but most commonly carries 1 error in the V region (B). Unmutated reads, however, are present in a level similar to the total frequency of exact matches in the same subject. This allele is also reported in VDJbase, primarily in unmutated form (C). A G303A variant does not exist in other known alleles of IGHV3-66 or alleles of the closely related gene IGHV3-53.  Population data: The A variant of SNP rs201182063 is found at 0.3% in European populations (Ensembl release 103) | | | | | | |
| IGHV3-7*03_S9833 | IGHV3-7*05 |  | yes |  |  | | | | | | |
| IGHV4-30-2*01_S6723 |  | IGHV4-30-2*01 G70A | yes |  | A. CDR3 length  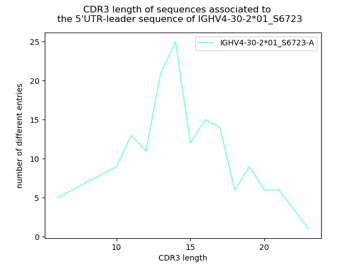 | | B. Error plot  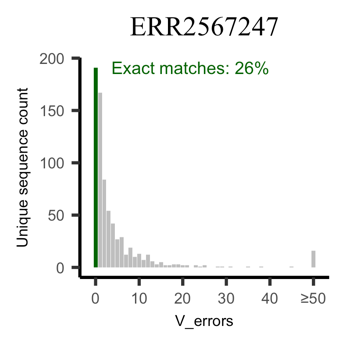  Exact matches, all genes^∞^: 23% | | | | C. OGRDBStats plot  ERR2567247  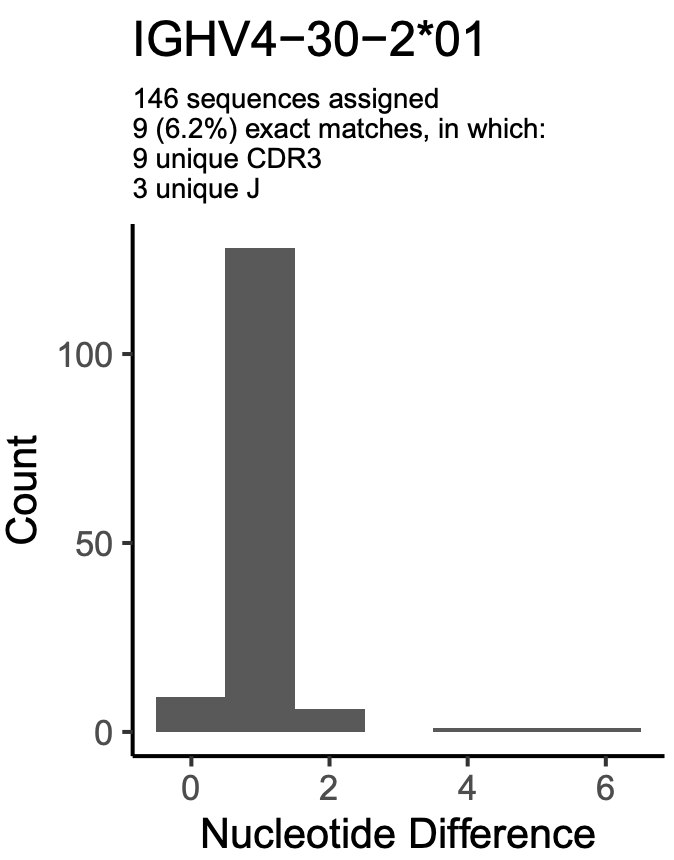 |
|  |  |  |  |  | Inferred by IgDiscover in ERR2567247 and in the study by Mikocziova et al. (2), but it is not identified in the current version of VDJbase (March, 2021). Reads associated with this inferred germline gene allele are diverse in CDR3 length (A) and has a high level of unmutated reads (B), similar to the total frequency of exact matches in the same subject. VDJbase identifies IGHV4-30-2*01 in this genotype but these sequences, in contrast to sequences representing other alleles, almost all carry at least 1 mutation as defined by its OGRDBStats plot (C). There is no other allele of IGHV4-30-2 in the genotype. No other allele of IGHV4-30-2 carry the G70A variant seen in this novel allele. | | | | | | |
| IGHV4-30-4*01_S5061 |  | IGHV4-30-4*01 A70G A107G | yes | yes† |  | | | | | | |
| IGHV4-38-2*02_S0038 | IGHV4-38-2*01 (3’-extension^&^) |  |  |  |  | | | | | | |
| IGHV4-39*01_S7498 |  | IGHV4-39*01 C66G | yes | yes† |  | | | | | | |
| IGHV4-39*07_S0654 |  | IGHV4-39*07 C288A | yes | yes* |  | | | | | | |
| IGHV4-4*02_S2599 |  | IGHV4-4*02 A106G | yes | yes** | A. CDR3 length  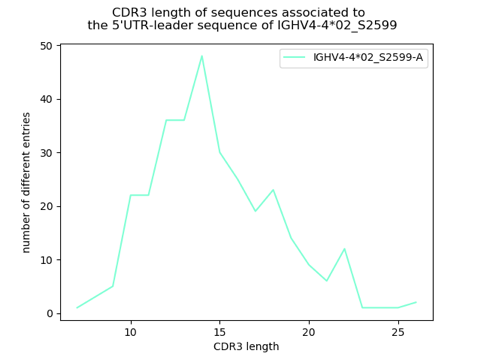 | | B. Error plot  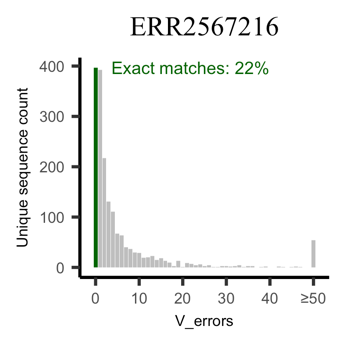  Exact matches, all genes^∞^: 21% | | | | C. OGRDBStats plot ERR2567216  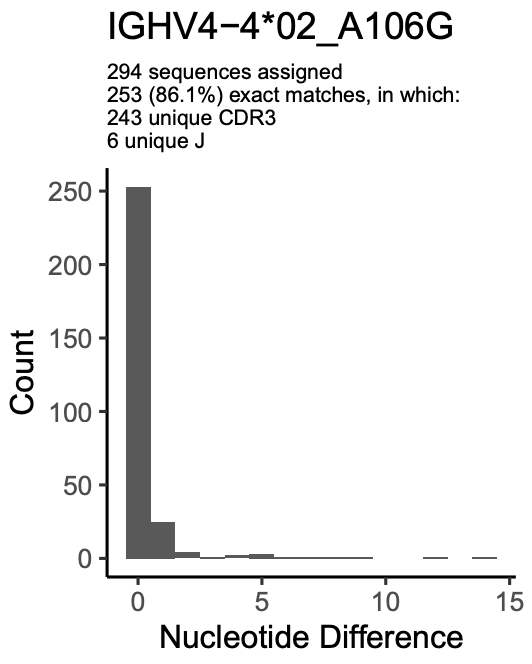 |
|  |  |  |  |  | Inferred by IgDiscover in ERR2567216, and in the study by Mikocziova et al. (2), and it is identified in the current version of VDJbase (May, 2021). Reads associated with this inferred germline gene allele are diverse in CDR3 length (A) and has a high level of unmutated reads (B), similar to the total frequency of exact matches in the same subject. VDJbase shows this inferred allele to be unmutated as defined by its OGRDBStats plot (C). The genotype of this subject carries the highly dissimilar allele IGHV4-4*07. Among alleles of genes similar to IGHV4-4, only allele IGHV4-61*08 carries the A106G modification, and this allele is not present in this genotype. | | | | | | |
| IGHV4-61*01_S9413 |  | IGHV4-61*01 A41G | yes | yes** | A. CDR3 length  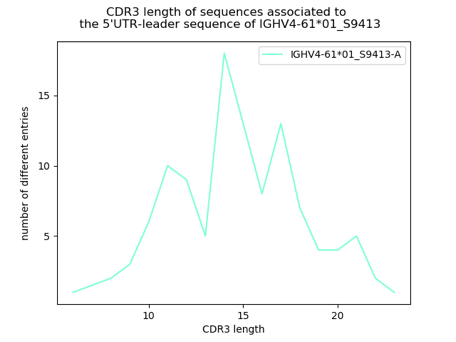 | | B. Error plot  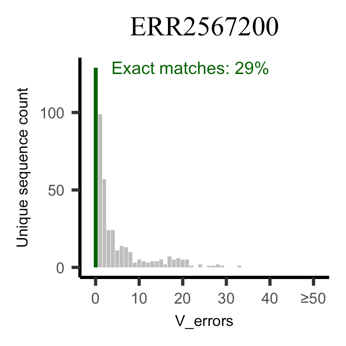  Exact matches, all genes^∞^: 28% | | | | C. OGRDBStats plot  ERR2567200  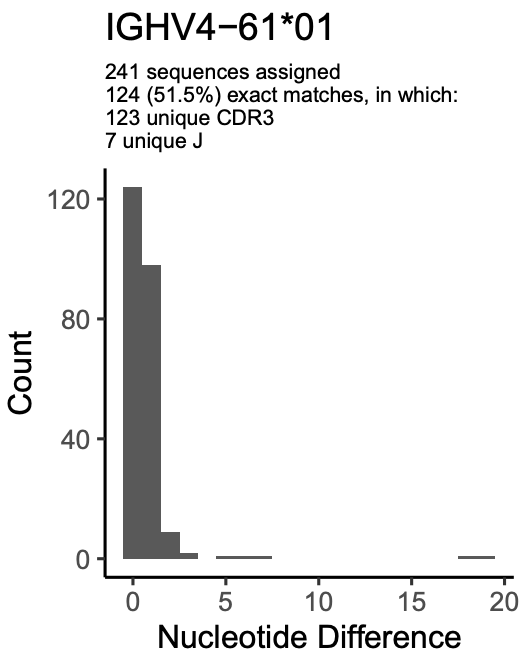 |
|  |  |  |  |  | E. Haplotyping data Data \| Germline gene/allele \| Haplotype \|; \| \| IGHJ6*02 \| IGHJ6*03; ERR2567200 \| IGHV4-61*01 \| 0 \| 21; \| IGHV4-61*01_S9413 \| 16 \| 0 | | | | | | |
|  |  |  |  |  | Inferred by IgDiscover in ERR2567200, and in the study by Mikocziova et al. (2), but it is not identified in the current version of VDJbase (May, 2021). Reads associated with this inferred germline gene allele are diverse in CDR3 length (A) and has a high level of unmutated reads (B), similar to the total frequency of exact matches in the same subject. VDJbase, that only inferred IGHV4-61*01, shows the presence of a large number of reads with one difference from the inferred allele, suggesting that a different allele might also be present in the genotype (C). Haplotyping suggests that the allele is present on one of two haplotypes that also carry IGHV4-61*01 (D). | | | | | | |
| IGHV4-61*02_S0442 |  | IGHV4-61*02 A234G | yes | yes* |  | | | | | | |
| IGHV5-51*01_S7082 | IGHV5-51*03 (3’-extension^&^) |  |  |  |  | | | | | | |

^&^ The IMGT database sequence is not of full length. Inference identified bases beyond the last base of the database sequence

^∞^ The frequency of unique sequences (all IGHV genes considered) having no errors in the V region, as determined by IgDiscover.

° Provisionally approved by IARC: https://www.antibodysociety.org/wordpress/wp-content/uploads/2020/12/Meeting-60-1_9_20-minutes.pdf

§ Provisionally approved by IARC: https://www.antibodysociety.org/wordpress/wp-content/uploads/2020/12/Meeting-61-13_10_20-minutes.pdf

† Provisionally approved by IARC: https://www.antibodysociety.org/wordpress/wp-content/uploads/2020/12/Meeting-62-3_11_20-minutes-1.pdf

* Provisionally approved by IARC: https://www.antibodysociety.org/wordpress/wp-content/uploads/2020/12/Meeting-63-24_11_20-minutes.pdf

** Provisionally approved by IARC at a lower level of confidence: https://www.antibodysociety.org/wordpress/wp-content/uploads/2020/12/Meeting-61-13_10_20-minutes.pdf and https://www.antibodysociety.org/wordpress/wp-content/uploads/2020/12/Meeting-63-24_11_20-minutes.pdf
