## Supplementary Figures 2-10, Supplementary Tables 1-2, 4-6 for "Computational inference, validation, and analysis of 5’UTR-leader sequences of alleles of immunoglobulin heavy chain variable genes"

**Content:**

Supplementary Tables 1-2, 4-6

Supplementary Figures 2-10

Supplementary Data 1 & 2, Supplementary Table 3, and Supplementary Figure 1 are provided as separate files.

**Supplementary Table 1.** Inference and population data of all IGHV 5’UTR-leader sequences identified in this study.

| Gene | Allele and 5’UTR-leader sequence | Diversified positions | Frequency | SNPs with MAF>1% in 1000 Genomes. Overall MAF, and MAF for highest (H),  lowest (L) and European (EUR) population) |
| --- | --- | --- | --- | --- |
| IGHV1-18 | 01-A | -27C | 2 | None |
|  | 01-B | -27G | 94 |  |
|  | 04-A | -27G | 31 |  |
| IGHV1-24 | 01-A | -70A, -71C | 2 | rs8013154, -33T/C: 2% (H: 7%, L: 0%, EUR: 0%) rs4511432, -71C/T: 11% (H: 31%, L: 1%, EUR: 13%) |
|  | 01-B | -70G, -71T | 30 |  |
|  | 01-C | -70G, -71C | 91 |  |
| IGHV1-2 | 02-A |  | 68 | None |
|  | 04-A |  | 64 |  |
|  | 05-A |  | 1 |  |
|  | 06-A |  | 19 |  |
| IGHV1-3 | 01-A | -8T, -61C, -65G | 1 | rs1143496, -8C/T: 42% (H: 63%, L: 26%, EUR: 41%) rs34315295, -61C/T: 27% (H: 43%, L: 15%, EUR: 15%) rs56372013, -65T/G: 28% (H: 43%, L: 8%, EUR: 43%) |
|  | 01-B | -8C, -61C, -65T | 9 |  |
|  | 01-C | -8C, -61T, -65T | 26 |  |
|  | 01-D | -8C, -61C, -65G | 57 |  |
|  | 01_S6816-A | -8C, -61C, -65G | 2 |  |
| IGHV1-46 | 01-A |  | 90 | None |
|  | 03-A |  | 49 |  |
|  | 04-A |  | 2 |  |
| IGHV1-58 | 01-A | -39A | 61 | rs148981028, -24G/A: 1% (H: 5%, L: 0%, EUR: 0%) rs1858692, -39A/G: 17% (H: 20%, L: 12%, EUR: 16%) |
|  | 02-A | -39A | 23 |  |
|  | 02-B | -39G | 16 |  |
| IGHV1-69-2 | 01-A | -70A, -71C | 1 | None^&^ |
|  | 01-B | -70G, -71A | 29 |  |
| IGHV1-69 | 01-A | -88A, -100C | 76 | rs4448834, -88A/G: 16% (H: 28%, L: 0%, EUR: 22%) rs10220412, -100G/C: 45% (H: 59%, L: 32%, EUR: 59%) rs59577815, -105A/C: 2% (H: 9%, L: 0%, EUR: 0%) |
|  | 01-B | -88G, -100C | 1 |  |
|  | 01_S8909-A | -88A, -100C | 1 |  |
|  | 02-A | -88A, -100G | 45 |  |
|  | 04-A | -88A, -100G | 24 |  |
|  | 04-B | -88G, -100C | 2 |  |
|  | 04_S3852-A | -88A, -100G | 2 |  |
|  | 06-A | -88G, -100C | 40 |  |
|  | 06-B | -88A, -100C | 2 |  |
|  | 06_S0471-A | -88G, -100C | 1 |  |
|  | 09-A | -88A, -100C | 2 |  |
|  | 09-B | -88A, -100G | 1 |  |
|  | 09-C | -88G, -100C | 5 |  |
|  | 10-A | -88A, -100G | 2 |  |
|  | 12-A | -88A, -100G | 3 |  |
|  | 17-A | -88G, -100C | 2 |  |
| IGHV1-8 | 01-A |  | 77 | None |
|  | 03-A |  | 2 |  |
| IGHV2-26 | 01-A | -49A | 98 | rs2073678, -49A/G: 13% (H: 25%, L: 3%, EUR: 3%) |
|  | 02_S3803-A | -49G | 1 |  |
| IGHV2-5 | 01-A |  | 57 | None |
|  | 02-A |  | 82 |  |
| IGHV2-70 | 01-A | -63T | 63 | rs61734101, -38C/T: 11% (H: 27%, L: 3%, EUR: 3%) rs10144801, -63C/T: 47% (H: 68%, L: 23%, EUR: 68%) |
|  | 04-A | -63T | 26 |  |
|  | 04_S5392-A | -63T | 1 |  |
|  | 15-A | -63T | 6 |  |
|  | 15-B | -63C | 36 |  |
| IGHV3-11 | 01-A |  | 76 | rs74207678, -39T/C: 1% (H: 7%, L: 0%, EUR: 0%) |
|  | 04-A |  | 2 |  |
|  | 05-A |  | 22 |  |
|  | 06-A |  | 47 |  |
| IGHV3-13 | 01-A |  | 61 | None |
|  | 01_S3164-A |  | 8 |  |
|  | 04-A |  | 19 |  |
|  | 05-A |  | 22 |  |
| IGHV3-15 | 01-A |  | 96 | rs10141897, -125G/A: 9% (H: 31%, L: 0%, EUR: 1%) |
|  | 07-A |  | 25 |  |
| IGHV3-20 | 01-A |  | 53 | rs12590474, -101C/T: 2% (H: 8%, L: 0%, EUR: 0%) |
|  | 04-A |  | 30 |  |
| IGHV3-21 | 01-A | -76T | 95 | rs73377566, -76T/G: 8% (H: 16%, L: 0%, EUR: 8%) |
|  | 01-B | -76G | 15 |  |
|  | 01_S4935-A | -76T | 1 |  |
|  | 01_S5913-A | -76T | 1 |  |
| IGHV3-23 | 01-A |  | 97 | rs112014369, -59C/T: 1% (H: 4%, L: 0%, EUR: 0%) rs12436700, -61C/T: 5% (H: 10%, L: 0%, EUR: 0%) rs12433853, -65A/G: 6% (H: 14%, L: 0%, EUR: 0%) |
|  | 04-A |  | 20 |  |
| IGHV3-30-3 | 01-A |  | 69 | No data° |
| IGHV3-30 | 01-A | -80G, -103C, -111G, -124G | 11 | rs371246072, -40C/G:2% (H: 9%, L: 0%, EUR: 0%) rs8021362, -103C/G: 48% (H: 62%, L: 36%, EUR: 62%) |
|  | 02-A | -80T, -103G, -111A, -124G | 3 |  |
|  | 02-B | -80G, -103G, -111A, -124G | 6 |  |
|  | 02_S4989-A | -80G, -103G, -111A, -124G | 1 |  |
|  | 03-A | -80G, -103G, -111A, -124G | 5 |  |
|  | 04-A | -80T, -103C, -111G, -124C | 6 |  |
|  | 04-B | -80G, -103C, -111G, -124C | 12 |  |
|  | 04_S7005-A | -80G, -103G, -111A, -124G | 1 |  |
|  | 18-A | -80G, -103G, -111A, -124G | 92 |  |
|  | 19_S5956-A | -80T, -103C, -111G, -124C | 1 |  |
| IGHV3-33 | 01-A |  | 96 | rs146691759, -80G/T: 12% (H: 37%, L: 0%, EUR: 3%) |
|  | 01_S3418-A |  | 1 |  |
| IGHV3-43D | 03-A |  | 15 | No data° |
|  | 04-A |  | 13 |  |
|  | 04_S5432-A |  | 1 |  |
| IGHV3-43 | 01-A | -62G | 73 | rs61999678, -62G/T: 16% (H: 36%, L: 4%, EUR: 6%) |
|  | 02-A | -62T | 11 |  |
| IGHV3-48 | 01-A |  | 50 | None |
|  | 02-A |  | 65 |  |
|  | 03-A |  | 34 |  |
|  | 04-A |  | 10 |  |
| IGHV3-49 | 03-A |  | 59 | None |
|  | 04-A |  | 37 |  |
|  | 05-A |  | 60 |  |
| IGHV3-53 | 01-A | -17T | 68 | rs2731152, -17T/C: 25% (H: 43%, L: 0%, EUR: 35%) |
|  | 02-A | -17C | 49 |  |
|  | 02_S9017-A | -17C | 1 |  |
|  | 04-A | -17T | 29 |  |
| IGHV3-64D | 06-A | -21A | 52 | No data° |
|  | 06_S5429-A | -21G | 12 |  |
|  | IGHV3-64*05_S2482-A^§^ | -21A | 10 |  |
| IGHV3-64 | 01-A |  | 46 | rs2073670, -56C/T: 42% (H: 61%, L: 24%, EUR: 61%) |
| IGHV3-66 | 01-A |  | 42 | None |
|  | 02-A |  | 29 |  |
|  | 02_S8911-A |  | 1 |  |
| IGHV3-72 | 01-A |  | 47 | None |
| IGHV3-73 | 01-A |  | 51 | rs61752554, -37G/C: 6% (H: 20%, L: 0%, EUR: 0%) |
|  | 02-A |  | 64 |  |
| IGHV3-74 | 01-A |  | 98 | rs140191839, -70C/G: 3% (H: 10%, L: 0%, EUR: 0%) |
| IGHV3-7 | 01-A | -52A | 76 | rs2073679, -52A/G: 46% (H: 58%, L: 31%, EUR: 43%) |
|  | 01-B | -52G | 2 |  |
|  | 03-A | -52G | 44 |  |
|  | 03-B | -52A | 1 |  |
|  | 03_S9833-A | -52G | 11 |  |
|  | 04-A | -52G | 21 |  |
| IGHV3-9 | 01-A |  | 77 | rs587598935, -60A/G: 8% (H: 13%, L: 2%, EUR: 13%) rs377293660, -88A/G: 1% (H: 3%, L: 1%, EUR: 1%) rs587754866, -101G/C: 20% (H: 30%, L: 10%, EUR: 26%) rs587664283, -127G/A:1% (H: 2%, L: 0%, EUR: 1%) |
|  | 03-A |  | 2 |  |
| IGHV4-30-2 | 01-A | -65G, -66T, -69Del | 3 | No data° |
|  | 01-B | -65A, -66C, -69C | 1 |  |
|  | 01-C | -65G, -66T, -69C | 69 |  |
|  | 01_S6723-A | -65G, -66T, -69C | 1 |  |
| IGHV4-30-4 | 01-A | -69C | 62 | No data° |
|  | 01_S5061-A | -69C | 2 |  |
|  | 07-A | -69Del | 2 |  |
|  | 07-B | -69C | 1 |  |
|  | 08-A | -69C | 1 |  |
| IGHV4-31 | 01-A |  | 9 | rs1393987777, -31G/C: 8% (H: 25%, L: 1%, EUR: 2%) rs371293120, -49C/T: 8% (H: 27%, L: 1%, EUR: 2%) rs112628447, -69C/Del: 1% (H: 3%, L: 0%, EUR: 0%) |
|  | 03-A |  | 89 |  |
| IGHV4-34 | 01-A |  | 98 | None |
| IGHV4-38-2 | 01-A |  | 23 | No data° |
|  | 02-A |  | 19 |  |
| IGHV4-39 | 01-A | -30G, -58A, -69A, -73T | 2 | rs4774113, -58A/C: 22% (H: 33%, L: 9%, EUR: 9%) rs4774114, -69A/C: 22% (H: 33%, L: 9%, EUR: 9%) rs4774115, -73T/C: 21% (H: 32%, L: 8%, EUR: 8%) |
|  | 01-B | -30C, -58A, -69A, -73T | 92 |  |
|  | 01_S7498-A | -30C, -58A, -69A, -73T | 1 |  |
|  | 07-A | -30C, -58C, -69C, -73C | 13 |  |
|  | 07_S0654-A | -30C, -58C, -69C, -73C | 2 |  |
| IGHV4-4 | 02-A | -1T, -31C, -65A, -66C, -74G, -78A, -81C | 7 | rs78225485, -1T/C: 49% (H: 65%, L: 37%, EUR: 51%) rs1141386. -31C/G: 18% (H: 32%, L: 3%, EUR: 32%) rs199897176, -52A/G: 2% (H: 6%, L: 0%, EUR: 0%) rs200645461, -65A/G: 5% (H: 13%, L: 2%, EUR: 2%) rs201659551, -66C/T: 5% (H: 13%, L: 2%, EUR: 2%) rs77360520, -74A/G: 18% (H: 40%, L: 5%, EUR: 5%) rs56086041, -78A/C: 31% (H: 61%, L: 13%, EUR: 17%) rs55998743, -81C/G: 34% (H: 65%, L: 14%, EUR: 18%) |
|  | 02-B | -1T, -31C, -65A, -66C, -74A, -78A, -81G | 1 |  |
|  | 02-C | -1T, -31C, -65A, -66C, -74A, -78A, -81C | 57 |  |
|  | 02-D | -1T, -31C, -65A, -66C, -74A, -78C, -81G | 22 |  |
|  | 02-E | -1T, -31C, -65G, -66T, -74A, -78A, -81C | 3 |  |
|  | 02-F | -1C, -31C, -65A, -66C, -74A, -78A, -81C | 6 |  |
|  | 02_S2599-A | -1T, -31C, -65A, -66C, -74A, -78C, -81G | 1 |  |
|  | 07-A | -1C, -31C, -65A, -66C, -74A, -78C, -81G | 2 |  |
|  | 07-B | -1C, -31G, -65A, -66C, -74A, -78C, -81G | 1 |  |
|  | 07-C | -1C, -31G, -65G, -66T, -74A, -78C, -81G | 1 |  |
|  | 07-D | -1C, -31C, -65A, -66C, -74A, -78A, -81C | 8 |  |
|  | 07-E | -1C, -31G, -65A, -66C, -74A, -78A, -81C | 55 |  |
|  | 07-F | -1C, -31C, -65G, -66T, -74A, -78A, -81C | 1 |  |
| IGHV4-59 | 01-A | -22G, -34T, -49T, -65G, -66T, -78C, -81G | 1 | rs56139643, -34T/C: 9% (H: 19%, L: 5%, EUR: 7%) |
|  | 01-B | -22A, -34T, -49T, -65G, -66T, -78C, -81G | 98 |  |
|  | 08-A | -22A, -34T, -49T, -65G, -66T, -78C, -81G | 42 |  |
|  | 12-A | -22A, -34C, -49C, -65A, -66C, -78A, -81C | 1 |  |
| IGHV4-61 | 01-A | -49C | 63 | rs201399631, -1C/T: 11% (H: 21%, L: 3%, EUR: 3%) rs2516897, -49C/T: 26% (H: 54%, L: 15%, EUR: 15%) rs200506032, -65G/A: 11% (H: 22%, L: 3%, EUR: 3%) rs201574166, -66T/C: 11% (H: 22%, L: 3%, EUR: 3%) |
|  | 01_S9413-A | -49C | 1 |  |
|  | 02-A | -49T | 30 |  |
|  | 02_S0442-A | -49T | 6 |  |
| IGHV5-10-1 | 01-A | -35G, -37T | 22 | None^&^ |
|  | 03-A | -35C, -37C | 43 |  |
|  | 03-B | -35G, -37T | 14 |  |
| IGHV5-51 | 01-A | -37C, -102T | 87 | rs116300992, -18C/A: 1% (H: 4%, L: 0%, EUR: 0%) rs61732461, -37C/T: 2% (H: 5%, L: 0%, EUR: 5%) rs77785266, -102T/C: 6% (H: 15%, L: 2%, EUR: 2%) |
|  | 01-B | -37T, -102T | 12 |  |
|  | 03-A | -37C, -102T | 40 |  |
|  | 03-B | -37C, -102C | 1 |  |
| IGHV6-1 | 01-A |  | 95 | None |
| IGHV7-4-1 | 01-A |  | 1 | None^&^ |
|  | 02-A |  | 26 |  |

^&^ The gene is not featured in GRCh37, and was thus analyzed using Ensembl (28) with GRCh38 as the reference genome.

° The gene is not featured in GRCh37 or GRCh38, and could thus not be analyzed.

^§^ Have now been included in IMGT database as IGHV3-64D*09 (Supplementary Table 3).

**Supplementary Table 2.** Haplotyping of distribution of reads associated to particular upstream regions of genes of the IGHV locus.

| Subject | IGHV gene and upstream region | IGHJ6 read distribution | |
| --- | --- | --- | --- |
|  |  | IGHJ6*02 | IGHJ6*03 |
| ERR2567187 | IGHV1-3*01-D | 72 | 0 |
|  | IGHV1-3*01-C | 0 | 76 |
| ERR2567204 | IGHV1-3*01-B | 160 | 0 |
|  | IGHV1-3*01-C | 0 | 55 |
| ERR2567243 | IGHV1-3*01-C | 48 | 0 |
|  | IGHV1-3*01-D | 1 | 47 |
| ERR2567249 | IGHV1-3*01-C | 1 | 44 |
|  | IGHV1-3*01-D | 54 | 0 |
| ERR2567266 | IGHV1-3*01-D | 78 | 0 |
|  | IGHV1-3*01-B | 0 | 50 |
| ERR2567271 | IGHV1-3*01-C | 28 | 0 |
|  | IGHV1-3*01-D | 0 | 23 |
| ERR2567189 | IGHV1-24*01-C | 31 | 0 |
|  | IGHV1-24*01-B | 0 | 9 |
| ERR2567206 | IGHV1-24*01-C | 0 | 12 |
|  | IGHV1-24*01-A | 15 | 0 |
| ERR2567226 | IGHV1-24*01-C | 0 | 8 |
|  | IGHV1-24*01-B | 9 | 0 |
| ERR2567230 | IGHV1-24*01-C | 50 | 0 |
|  | IGHV1-24*01-B | 0 | 8 |
| ERR2567246 | IGHV1-24*01-C | 49 | 0 |
|  | IGHV1-24*01-B | 0 | 5 |
| ERR2567263 | IGHV1-24*01-C | 54 | 0 |
|  | IGHV1-24*01-B | 1 | 29 |
| ERR2567271 | IGHV1-24*01-C | 12 | 0 |
|  | IGHV1-24*01-B | 0 | 4 |
| ERR2567277 | IGHV1-24*01-B | 0 | 19 |
|  | IGHV1-24*01-C | 18 | 0 |
| ERR2567206 | IGHV1-58*02-A | 20 | 0 |
|  | IGHV1-58*02-B | 0 | 3 |
| ERR2567243 | IGHV1-58*02-A | 11 | 0 |
|  | IGHV1-58*02-B | 0 | 2 |
| ERR2567187 | IGHV1-58*02-A | 0 | 14 |
|  | IGHV1-58*02-B | 9 | 0 |
| ERR2567264 | IGHV1-69*06-B | 2 | 231 |
|  | IGHV1-69*06-A | 1 | 204 |
|  | IGHV1-69*02-A | 95 | 0 |
| ERR2567220 | IGHV2-70*15-A | 24 | 1 |
|  | IGHV2-70*15-B | 0 | 18 |
| ERR2567192 | IGHV3-21*01-A | 0 | 15 |
|  | IGHV3-21*01-B | 19 | 0 |
| ERR2567266 | IGHV3-21*01-A | 0 | 53 |
|  | IGHV3-21*01-B | 64 | 1 |
| ERR2567266 | IGHV4-4*02-C | 58 | 0 |
|  | IGHV4-4*07-A | 1 | 107 |
| ERR2567189 | IGHV4-4*02-F | 0 | 24 |
|  | IGHV4-4*07-E | 37 | 0 |
| ERR2567200 | IGHV4-4*02-C | 0 | 46 |
|  | IGHV4-4*07-B | 48 | 0 |
| ERR2567230 | IGHV4-4*02-A | 0 | 38 |
|  | IGHV4-4*07-D | 72 | 0 |
| ERR2567192 | IGHV4-4*02-C | 16 | 0 |
|  | IGHV4-4*02-A | 0 | 17 |
| ERR2567204 | IGHV4-4*02-C | 74 | 0 |
|  | IGHV4-4*02-D | 0 | 75 |
| ERR2567246 | IGHV4-4*02-F | 0 | 36 |
|  | IGHV4-4*02-C | 65 | 0 |
| ERR2567254 | IGHV4-4*02-C | 55 | 0 |
|  | IGHV4-4*02-F | 0 | 42 |
| ERR2567261 | IGHV4-4*02-C | 99 | 0 |
|  | IGHV4-4*02-F | 0 | 78 |
| ERR2567271 | IGHV4-4*02-D | 51 | 0 |
|  | IGHV4-4*02-F | 0 | 5 |
| ERR2567274 | IGHV4-4*02-E | 24 | 0 |
|  | IGHV4-4*02-C | 0 | 21 |
| ERR2567187 | IGHV4-4*02-C | 65 | 0 |
|  | IGHV4-4*01^§^ | - | - |
| ERR2567201 | IGHV4-4*07-E | 27 | 0 |
|  | IGHV4-4*07-D | 0 | 35 |
| ERR2567263 | IGHV4-4*07-F | 0 | 94 |
|  | IGHV4-4*07-C | 84 | 1 |
| ERR2567215 | IGHV4-30-2*01-C | 0 | 39 |
|  | IGHV4-30-2*01-A | 31 | 0 |
| ERR2567254 | IGHV4-39*01-A | 0 | 190 |
|  | IGHV4-39*01-B | 118 | 0 |
| ERR2567220 | IGHV5-51*01-B | 25 | 0 |
|  | IGHV5-51*01-A | 0 | 36 |
| ERR2567231 | IGHV5-51*01-B | 0 | 35 |
|  | IGHV5-51*01-A | 48 | 0 |
| ERR2567264 | IGHV5-51*01-A | 2 | 52 |
|  | IGHV5-51*01-B | 41 | 0 |
| ERR2567266 | IGHV5-51*01-A | 72 | 0 |
|  | IGHV5-51*01-B | 0 | 35 |

^§^ IGHV4-4*01 was not identified in the present IgDiscover analysis, but in a recent study emoplying IMGT/HighV-quest analysis (23).

**Supplementary Table 4.** 5’-end of the 5’UTR-leader sequence of sets of genes inferred in the present study, 5’-end of the 5’UTR-leader sequence as previously defined as having variants with a 5’-terminal G, and population data (Ensembl release 103) (28) corresponding to the 5’-end of the 5’-UTR, and the extent of support for determination of diverse 5’-ends of 5’UTR-leader sequence. Only data related to genes represented in the Ensembl database are shown.

| Gene/allele^§^ | 5’end of inferred 5’UTR (this study)^†^ | 5’end of inferred 5’UTR by Mikocziova et al. (18)^†^ | Dominant base of SNP corresponding to the 5’-base inferred by Mikocziova et al. (18); (Ensembl; irrespective of allele) | Support in population data for different 5’-bases inferred by Mikocziova et al. (18) | |
| --- | --- | --- | --- | --- | --- |
| IGHV1-3*01 | 5’-A_-101_ACC | 5’-C_-102_AACC and  5’-G_-102_AACC | C_-102_ (Highest population MAF<0.01%) | 5’-C_-102:_ yes | 5’-G_-102:_ no |
| IGHV1-46*01 | 5’-A_-115_TCA | 5’-C_-116_ATCA and  5’-G_-116_ATCA | C_-116_ (Highest population MAF<0.01%) | 5’-C_-116:_ yes | 5’-G_-116:_ no |
| IGHV1-46*03 | 5’-A_-115_TCA | 5’-C_-116_ATCA and  5’-G_-116_ATCA |  |  |  |
| IGHV2-5*01 | 5’-A_-73_CT | 5’-T_-75_GACT and  5’-G_-75_GACT | T_-75_ (Highest population MAF<0.01%) | 5’-T_-75_: yes | 5’-G_-75_: no |
| IGHV2-5*02 | 5’-A_-73_CT | 5’-T_-75_GACT and  5’-G_-75_GACT |  |  |  |
| IGHV2-70*01 | 5’-A_-73_AT | 5’-T_-75_GAAT and  5’-G_-75_GAAT | T_-75_ (Highest population MAF<0.01%) | 5’-T_-75_: yes | 5’-G_-75_: no |
| IGHV2-70*04 | 5’-A_-73_AT | 5’-T_-75_GAAT and  5’-G_-75_GAAT |  |  |  |
| IGHV2-70*15 | 5’-A_-73_AT | 5’-G_-75_GAAT |  |  |  |
| IGHV3-11*01 | 5’-A_-137_GTC | 5’-C_-138_AGCT and  5’-G_-138_AGCT | C_-138_ (Highest population MAF<0.01%) | 5’-C_-138_: yes | 5’-G_-138_: no |
| IGHV3-11*05 | 5’-A_-137_GTC | 5’-G_-138_AGCT |  |  |  |
| IGHV3-11*06 | 5’-A_-137_GTC | 5’-C_-138_AGCT and  5’-G_-138_AGCT |  |  |  |
| IGHV3-20*01 | 5’-A_-137_GCT | 5’-G_-138_AGCT | C_-138_ (rs560966965; Highest population MAF (T): 4%) | 5’-C_-138_: yes | 5’-G_-138_: no |
| IGHV3-20*01 C307T (*04) | 5’-A_-137_GCT | 5’-G_-138_AGCT |  |  |  |
| IGHV3-43*01 | 5’-A_-137_GCTCT* | 5’-G_-136_CTCT | G (Highest population MAF<0.01%) |  | 5’-G_-136_: yes |
| IGHV3-43*02 | 5’-A_-137_GCTCT* | 5’-G_136_CTCT |  |  |  |
| IGHV3-53*01 | A_-127_GA | 5’-G_-131_GGGAGA | T (Highest population MAF<0.01%) | 5’-T_-131_: yes | 5’-G_-131_: no |
| IGHV3-53*02 | G_-126_A | 5’-G_-131_GGGAGA and  5’-T_-131_GGGAGA |  |  |  |
| IGHV3-53*04 | A_-127_GA | 5’-G_-131_GGGAGA |  |  |  |
| IGHV3-64*01 | G_-136_CT | 5’-C_-138_AGCT and  5’-G_-138_AGCT | C (Highest population MAF<0.01%) | 5’-C_-138_: yes | 5’-G_-138_: no |
| IGHV3-66*01 | 5’-A_-136_GCT | 5’-C_-137_AGCT and 5  ’-G_-137_AGCT | C (Highest population MAF<0.01%) | 5’-C_-137_: yes | 5’-G_-137_: no |
| IGHV3-66*02 | 5’-A_-136_GCT | 5’-G_-137_AGCT |  |  |  |
| IGHV3-72*01 | 5’-A_-130_GAGC | 5’-G_-138_AGCTCTGAGAGC | C_-138_ (Highest population MAF<0.01%) | 5’-C_-138_: yes | 5’-G_-138_: no |
| IGHV3-73*01 | 5’-A_-137_GCTC | 5’-C_-138_AGCT and  5’-G_-138_AGCT | C_-138_ (Highest population MAF<0.01%) | 5’-C_-138_: yes | 5’-G_-138_: no |
| IGHV3-73*02 | 5’-A_-137_GCTC | 5’-C_-138_AGCT and  5’-G_-138_AGCT |  |  |  |
| IGHV4-4*02 | 5’-A_-94_TACTT or 5’-T_-93_ACTT or 5’-A_-92_CTT** | 5’-T_-93_ACTT and  5’-G_-93_ACTT | T (rs531656003; highest population MAF (A): 1%) | 5’-T_-93_: yes | 5’-G_-93_: no |
| IGHV4-4*07 | 5’-A_-94_TACTT or 5’-A_-92_CTT** | 5’-T_-93_ACTT and  5’-G_-93_ACTT |  |  |  |
| IGHV4-31*03 | 5’-A_-94_TACTT | 5’-T_-93_ACTT and  5’-G_-93_ACTT | T_-93_ (Highest population MAF<0.01%) | 5’-T_-93_: yes | 5’-G_-93_: no |
| IGHV4-34*01 | 5’-A_-95_GTGCTTT*** | 5’-G_-92_CTTT | G_-92_ |  | 5’-G_-92_: yes |
| IGHV4-59*01 | 5’-A_-92_CTT | 5’-T_-93_ACTT and  5’-G_-93_ACTT | T_-93_ (Highest population MAF<0.01%) | 5’-T_-93_: yes | 5’-G_-93_: no |
| IGHV4-59*08 | 5’-A_-92_CTT | 5’-T_-93_ACTT and  5’-G_-93_ACTT |  |  |  |
| IGHV4-61*01 | 5’-A_-92_CTT | 5’-T_-93_ACTT and  5’-G_-93_ACTT | T_-93_ (Highest population MAF<0.01%) | 5’-T_-93_: yes | 5’-G_-93_: no |
| IGHV4-61*02 | 5’-A_-92_CTT | 5’-T_-93_ACTT and  5’-G_-93_ACTT |  |  |  |
| IGHV6-1*01 | 5’-A_-110_GAG | 5’-C_-111_AGAG and  5’-G_-111_AGAG | C_-111_ (Highest population MAF<0.01%) | 5’-C_-111_: yes | 5’-G_-111_: no |

^§^ Genes/alleles that feature a 5’G in (some of) its inferred 5’UTR-leader sequences and that are featured with population data in the Ensemble database.

^†^ The 5’-most base is numbered as the number of bases upstream of the first base encoding the H chain variable domain

* Additional 5’-A inferred in this study is supported by genomic data

** Different lengths determined depending on which 5’UTR-leader was inferred

*** Additional 5’-AGT inferred in this study is supported by genomic data

**Supplementary Table 5.** Lack of evidence for inference of base -93 in upstream regions of IGHV4 alleles. This base is close to the limit of called bases in reads representing upstream regions of genes belonging to this subgroup. The ratio should be close to 100% or 0% if evidence of G or T, respectively, at position -93 is present among the transcripts. Haplotype distribution of base -93, as defined by association to different alleles of IGHJ6, was calculated from the reads precisely matching the inferred 5’UTR-leader sequence from base -92 to -1, and unequivocally assigned by the IMGT/HighV-Quest tool (17) to the allele in question. The call of base -93 of alleles of IGHV1-46, a gene whose readable upstream region commonly extends beyond base -93, is shown for comparison, a call that with high confidence identify T as the base found in position -93.

| Dataset | Gene | Allele | 5’UTR-leader | IGHV6*02  base -93: G / (T+G) (%) | IGHV6*03  base -93: G / (T+G) (%) |
| --- | --- | --- | --- | --- | --- |
| ERR2567187 | IGHV4-30-2 | *01 | C | 54 | 51 |
|  | IGHV4-30-4 | *01 | A | 24 | 27 |
|  | IGHV4-31 | *03 | A | 39 | 32 |
|  | IGHV4-34 | *01 | A | 33 | 34 |
|  | IGHV4-38-2 | *01 | A | 67 |  |
|  | IGHV4-39 | *01 | B |  | 58 |
|  | IGHV4-4 | *01 |  |  | ^†^ |
|  |  | *02 | C | 43 |  |
|  | IGHV4-59 | *01 | B | 46 | 53 |
|  |  | *08 | A |  | 45 |
|  | IGHV4-61 | *02 | A | 78 |  |
|  | IGHV1-46 | *01 | A | 1 | 1 |
| ERR2567189 | IGHV4-30-2 | *01 | C | 52 |  |
|  | IGHV4-30-4 | *01 | A | 43 |  |
|  | IGHV4-31 | *03 | A | 33 |  |
|  | IGHV4-34 | *01 | A | 28 | 35 |
|  | IGHV4-38-2 | *02 | A | 60 |  |
|  | IGHV4-39 | *01 | B | 60 | 54 |
|  | IGHV4-4 | *02 | F |  | 37 |
|  |  | *07 | E | 48 |  |
|  | IGHV4-59 | *01 | B | 47 | 43 |
|  |  | *08 | A | 50 |  |
|  | IGHV4-61 | *01^∞^ | A |  | 43 |
|  | IGHV1-46 | *01 | A |  | 3 |
|  |  | *03 | A | 0 |  |
| ERR2567192 | IGHV4-30-2 | *01 | C |  | 69 |
|  | IGHV4-30-4 | *01^∞^ | A |  | 47 |
|  | IGHV4-31 | *03 | A | 41 | 0 |
|  | IGHV4-34 | *01 | A | 37 | 26 |
|  | IGHV4-39 | *01 | B | 50 | 73 |
|  | IGHV4-4 | *02 | C | 18 |  |
|  |  |  | A |  | 14 |
|  | IGHV4-59 | *01 | B | 25 | 46 |
|  |  | *08 | A | 13 | 30 |
|  | IGHV1-46 | *01 | A | 8 | 0 |
| ERR2567199 | IGHV4-31 | *03 | A |  | 41 |
|  | IGHV4-34 | *01 | A | 36 | 32 |
|  | IGHV4-38-2 | *02 | A | 70 |  |
|  | IGHV4-39 | *01 | B |  | 59 |
|  |  | *07 | A | 63 |  |
|  | IGHV4-4 | *02 | C |  | 54 |
|  |  | *07 | E | 36 |  |
|  | IGHV4-59 | *01 | B | 43 | 50 |
|  | IGHV4-61 | *01^∞^ | A | 58 | 40 |
|  | IGHV1-46 | *01 | A | 2 | 0 |
| ERR2567200 | IGHV4-30-2 | *01 | C | 49 |  |
|  | IGHV4-31 | *01^∞^ | A |  | 34 |
|  |  | *03 | A | 36 |  |
|  | IGHV4-34 | *01 | A | 30 | 34 |
|  | IGHV4-38-2 | *01 | A |  | 68 |
|  | IGHV4-39 | *01 | B | 57 |  |
|  | IGHV4-4 | *02 | C |  | 34 |
|  |  | *07 | B | 59 |  |
|  | IGHV4-59 | *01 | B | 46 | 52 |
|  | IGHV4-61 | *01^∞^ | A |  | 35 |
|  |  | *01__S9413 (A41G) | A | 43 |  |
|  | IGHV1-46 | *01 | A |  | 0 |
|  |  | *03 | A | 2 |  |
| ERR2567201 | IGHV4-30-2 | *01 | C | 67 | 41 |
|  | IGHV4-30-4 | *01 | A | 38 | 27 |
|  | IGHV4-31 | *03 | A | 57 | 37 |
|  | IGHV4-34 | *01 | A | 37 | 33 |
|  | IGHV4-39 | *01 | B | 56 | 52 |
|  | IGHV4-4 | *07 | E | 27 |  |
|  |  |  | D |  | 36 |
|  | IGHV4-59 | *01 | B | 52 | 49 |
|  | IGHV4-61 | *01^∞^ | A |  | 47 |
|  |  | *02 | A | 31 |  |
|  | IGHV1-46 | *01 | A |  | 3 |
|  |  | *03 | A | 1 |  |
| ERR2567204 | IGHV4-30-2 | *01 | C | 55 |  |
|  | IGHV4-31 | *03 | A |  | 37 |
|  | IGHV4-34 | *01 | A | 35 | 38 |
|  | IGHV4-39 | *01 | B | 64 | 58 |
|  | IGHV4-4 | *02 | C | 46 |  |
|  |  |  | D |  | 53 |
|  | IGHV4-59 | *01 | B | 54 | 46 |
|  | IGHV4-61 | *01^∞^ | A | 37 |  |
|  |  | *02 | A |  | 68 |
|  | IGHV1-46 | *01 | A | 1 |  |
|  |  | *03 | A |  | 4 |
| ERR2567206 | IGHV4-30-2 | *01 | C |  | 50 |
|  | IGHV4-34 | *01 | A | 31 | 27 |
|  | IGHV4-39 | *01 | B |  | 52 |
|  |  | *07_S0654 (C288A) | A | 62 |  |
|  | IGHV4-4 | *01 |  | ^†^ |  |
|  |  | *02 | F |  | 55 |
|  | IGHV4-59 | *01 | B | 50 | 45 |
|  |  | *08 | A | 44 |  |
|  | IGHV4-61 | *02 | A |  | 67 |
|  | IGHV1-46 | *01 | A | 6 |  |
|  |  | *03 | A |  | 4 |

^†^ IGHV4-4*01, a very poorly expressed allele not directly studied in the present investigation, is present in this haplotype (23).

^∞^ IGHV4-31*01 and IGHV4-61*01 are poorly expressed in comparison to IGHV4-31*03 and IGHV4-61*02, respectively, and may not be inferred depending on inference software settings.

**Supplementary Table 6.** Inference and population data suggesting that upstream regions of primary IMGT entries, in addition to IGHV2-5*01 (Supplementary Figure 5), may be in error.

| Germline gene | Inferred upstream sequences and sequences defined by primary IMGT entries* | Population data (Ensembl Genome Browser) |
| --- | --- | --- |
| IGHV1-18*01 | 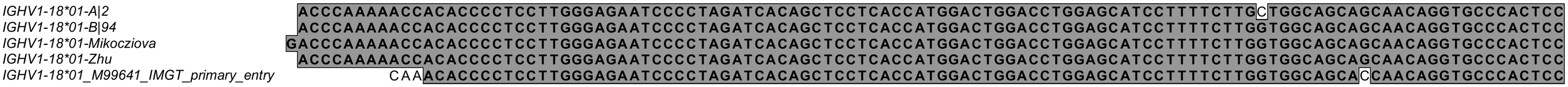 | Expected base at -18 is G. Highest population MAF for this base: <0.01% |
| IGHV1-8*01 | 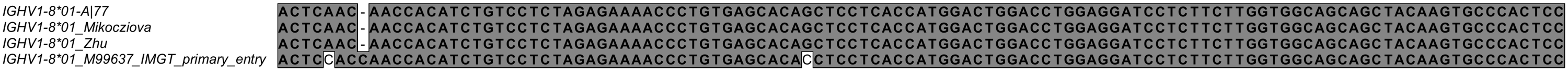 | Expected base at -67 is G. Highest population MAF for this base: <0.01% |
| IGHV3-11*01 | 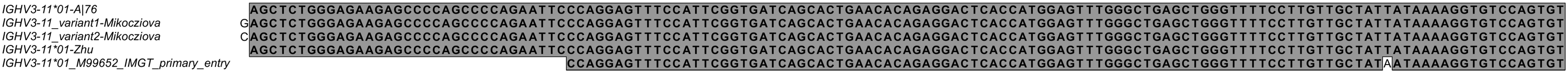 | Expected base at -19 is T. Highest population MAF for this base: <0.01% |
| IGHV3-23*01 | 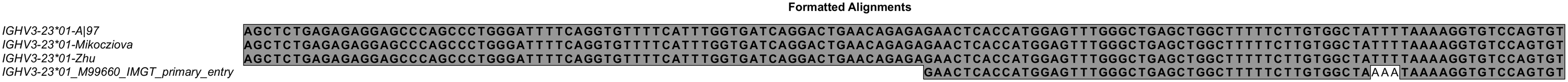 | Expected bases at -20 – -18 are TTT. Highest population MAF for these bases: <0.01% |
| IGHV3-64*01 | 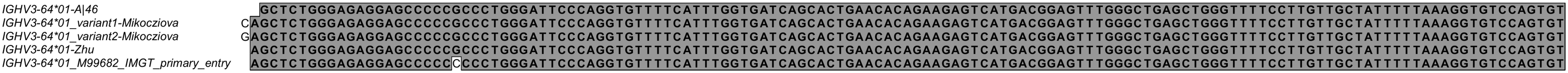 | Expected base at -116 is G. Highest population MAF for this base: <0.01% |
| IGHV3-72*01 | 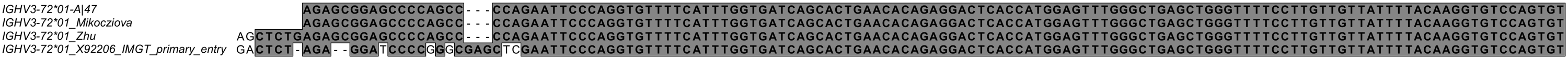 | Expected base for instance at bases -111 – -112 are AC. Highest population MAF for these bases: <0.01% |
| IGHV4-4*02 | 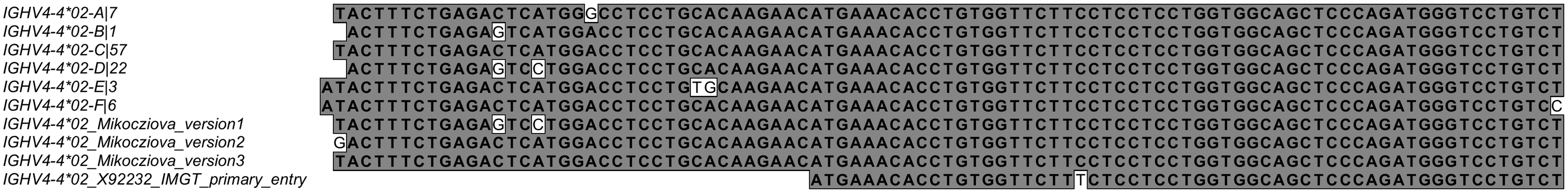 | Expected base at -37 is C. Highest population MAF for this base: <0.01% |
| IGHV5-51*01 | 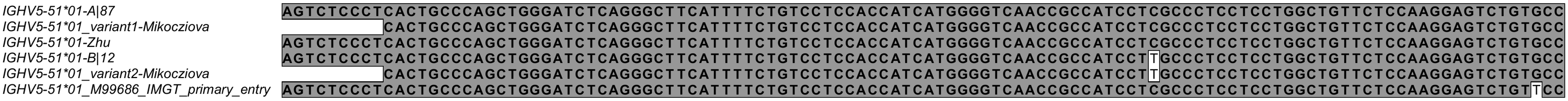 | Expected base at -3 is G. Highest population MAF for this base: <0.01% |

* Inferred upstream regions were defined in this study, by Mikocziova et al. (18) and Zhu et al. (14). In the latter case only the most frequent variants were included and major variants of IGHV4-4*02 upstream regions defined by Zhu et al are shown in Supplementary Figure 3..

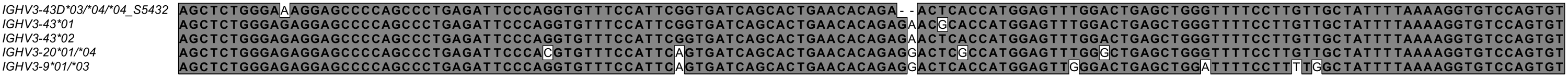

**Supplementary Figure 2.** Upstream region sequences of IGHV3-43D, IGHV3-43 and IGHV3-9, which in parts are very similar.

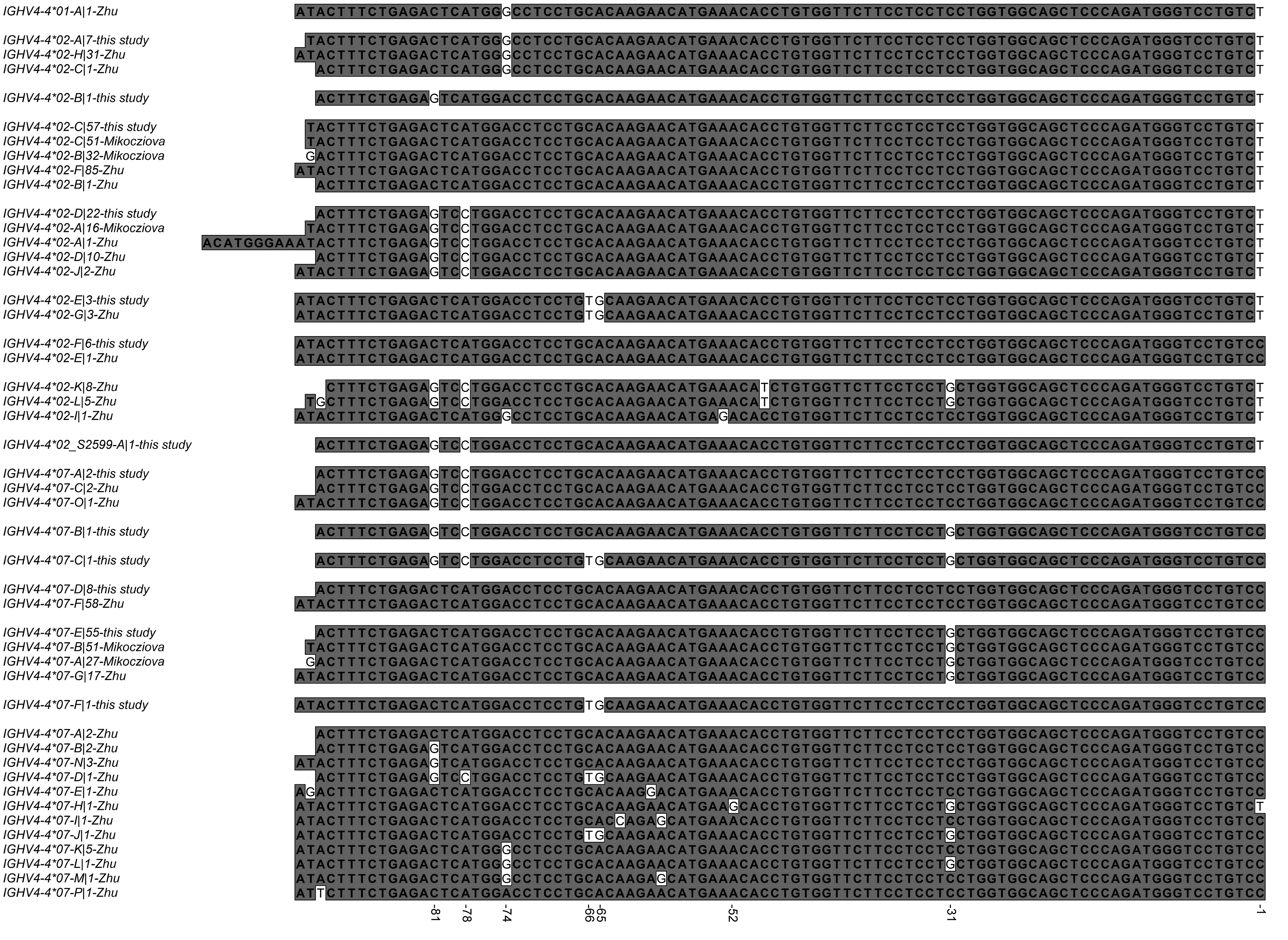

**Supplementary Figure 3.** The alignment of inferred upstream regions of alleles IGHV4-4*01, IGHV4-4*02 and IGHV4-4*07 as described in this study, Mikocziova et al. (18), and Zhu et al. (14). The sequences from Mikocziova et al.’s study and this study were inferred from exactly the same data set as used in this study. Names of upstream regions indicate “the IGHV allele to which the sequence was found to be associated – a letter separating different sequences associated to this allele number of instances in which it was found – study of identification”.

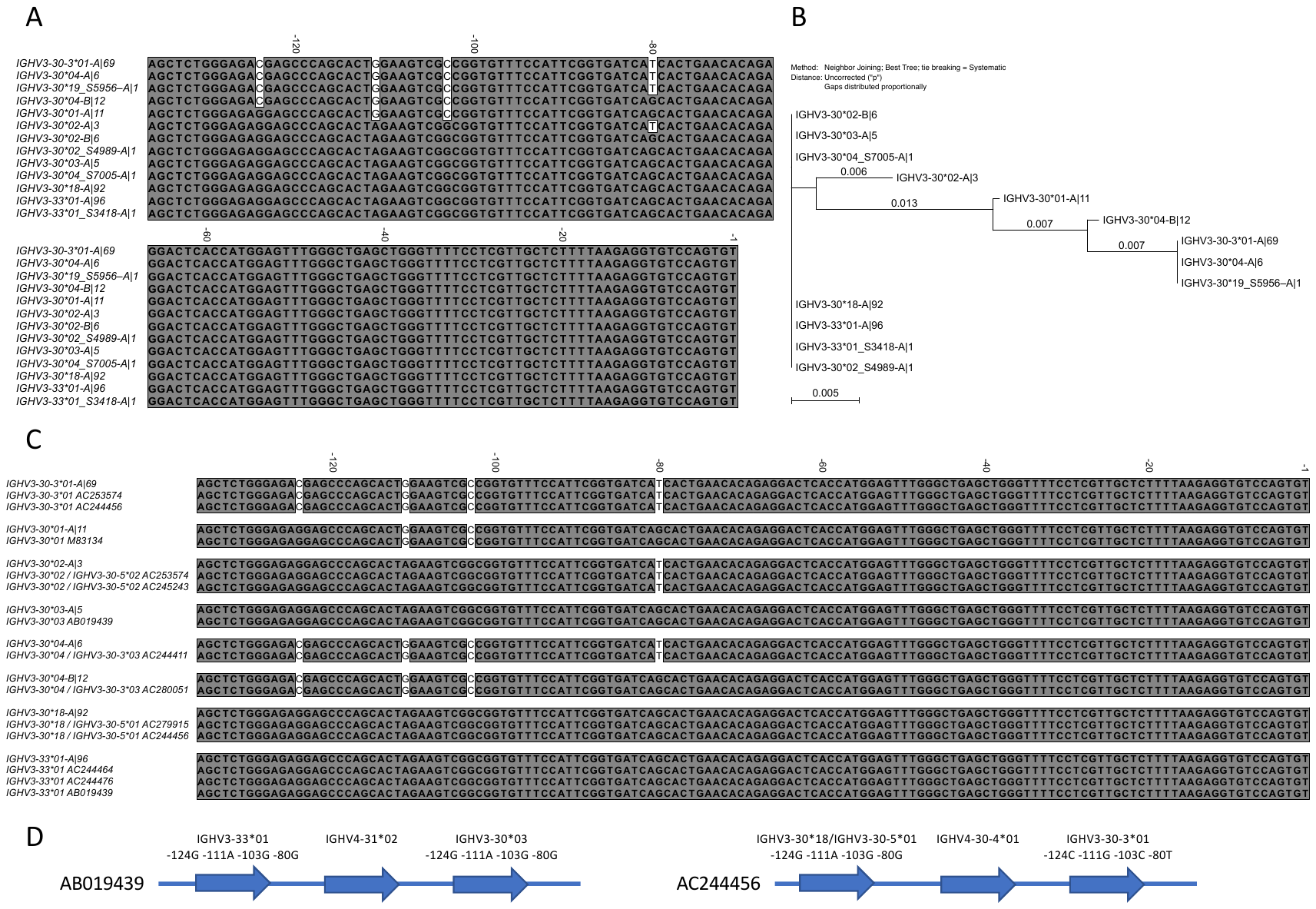

**Supplementary Figure 4.** Sequences (**A**) and Neighbour Joining Tree (**B**) of 5’UTR-leader sequences of alleles of IGHV3-30, IGHV3-30-3, and IGHV3-33. Examples of 5’UTR-leader sequences identified by inference have also been seen in genomic sequence data (genomic sequences are shown after removal of the leader intron; accession numbers are shown for selected cases) (**C**). Examples of part of the IGHV locus that also include alleles of IGHV3-30, IGHV3-30-3, IGHV3-30-5, and IGHV3-33 are illustrated (**D**).

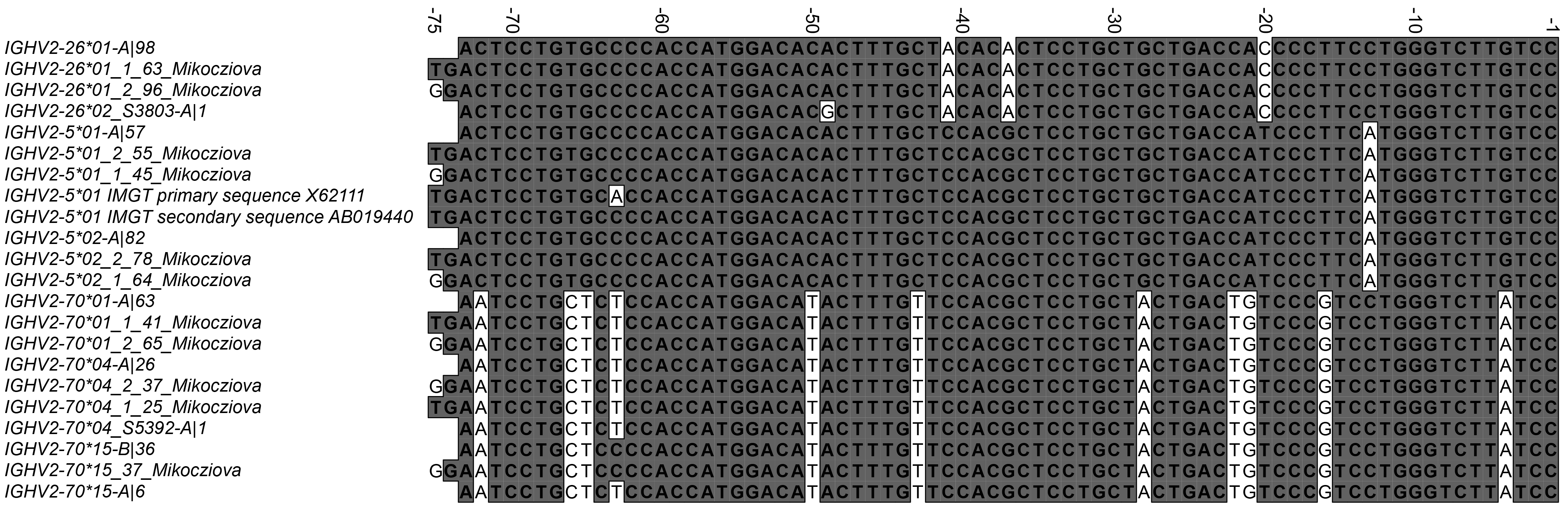

**Supplementary Figure 5.** Sequences of inferred upstream regions of genes/alleles of the IGHV2 subgroup (this study and Mikocziova et al [18]), and of the upstream regions of IGHV2-5*01 represented in two entries of the IMGT database, one of which features a C->A variant in base -63. This variant is not seen in human populations (highest population MAF<0.01%). The variant (T->G) previously inferred at high frequency in position -75 (18) is not reproduced in population studies as this base is represented by T at a frequency of 99.99% in IGHV2-26, IGHV2-5, and IGHV2-70 in studied human populations (http://www.ensembl.org) (28).

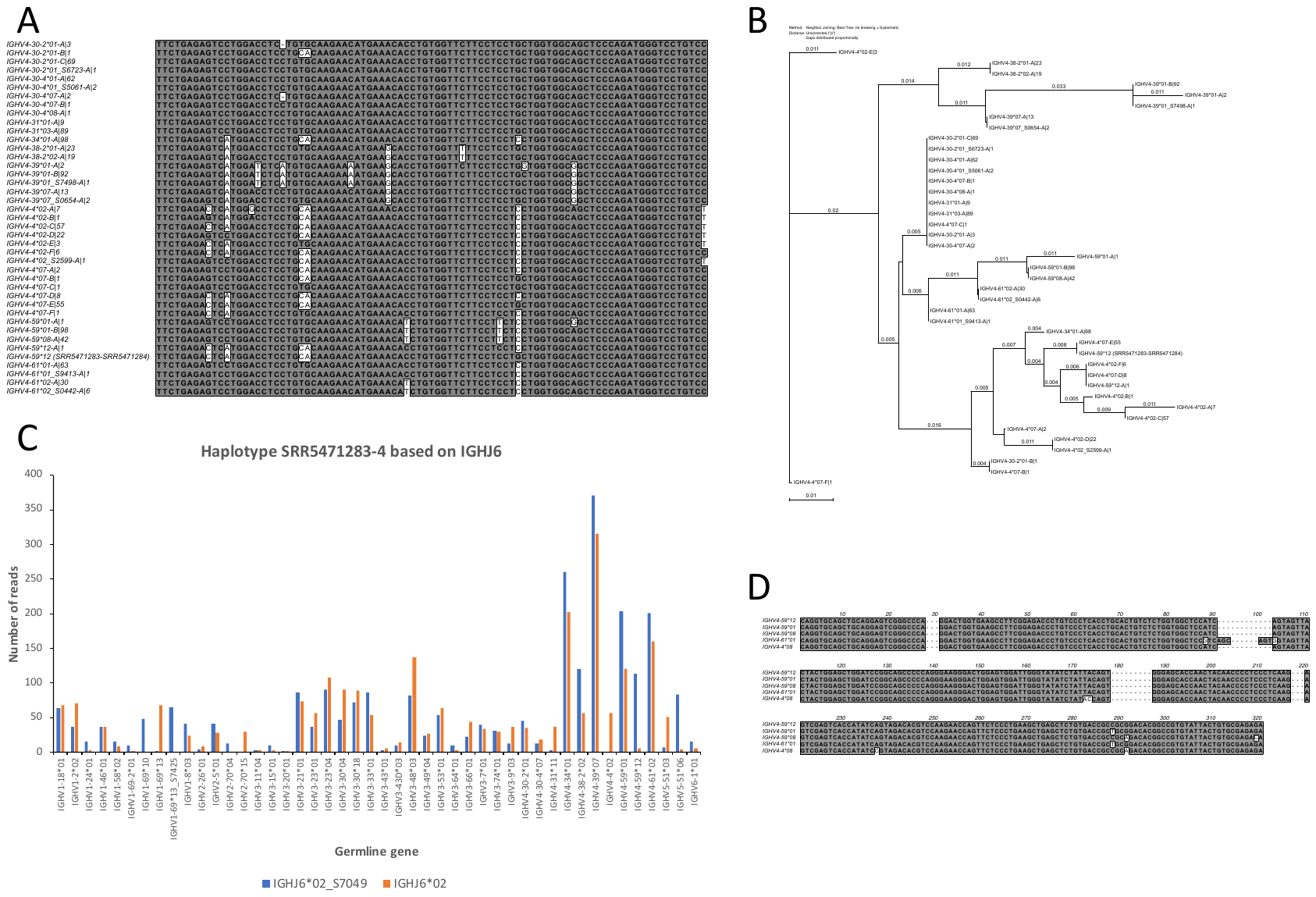

**Supplementary Figure 6.** Sequences (**A**) and neighbour Joining Tree (**B**) of upstream regions of genes belonging to subgroup 4, and haplotype analysis (**C**) of the individual (donor LP08248) where IGHV4-59*12 originally was identified (24). Upstream regions of IGHV4-59*12 are more closely related to upstream regions of most alleles of IGHV4-4 than to other upstream regions of alleles of IGHV4-59. Similarity of some alleles of IGHV4-4, IGHV4-59, and IGHV4-61 is shown (**D**).

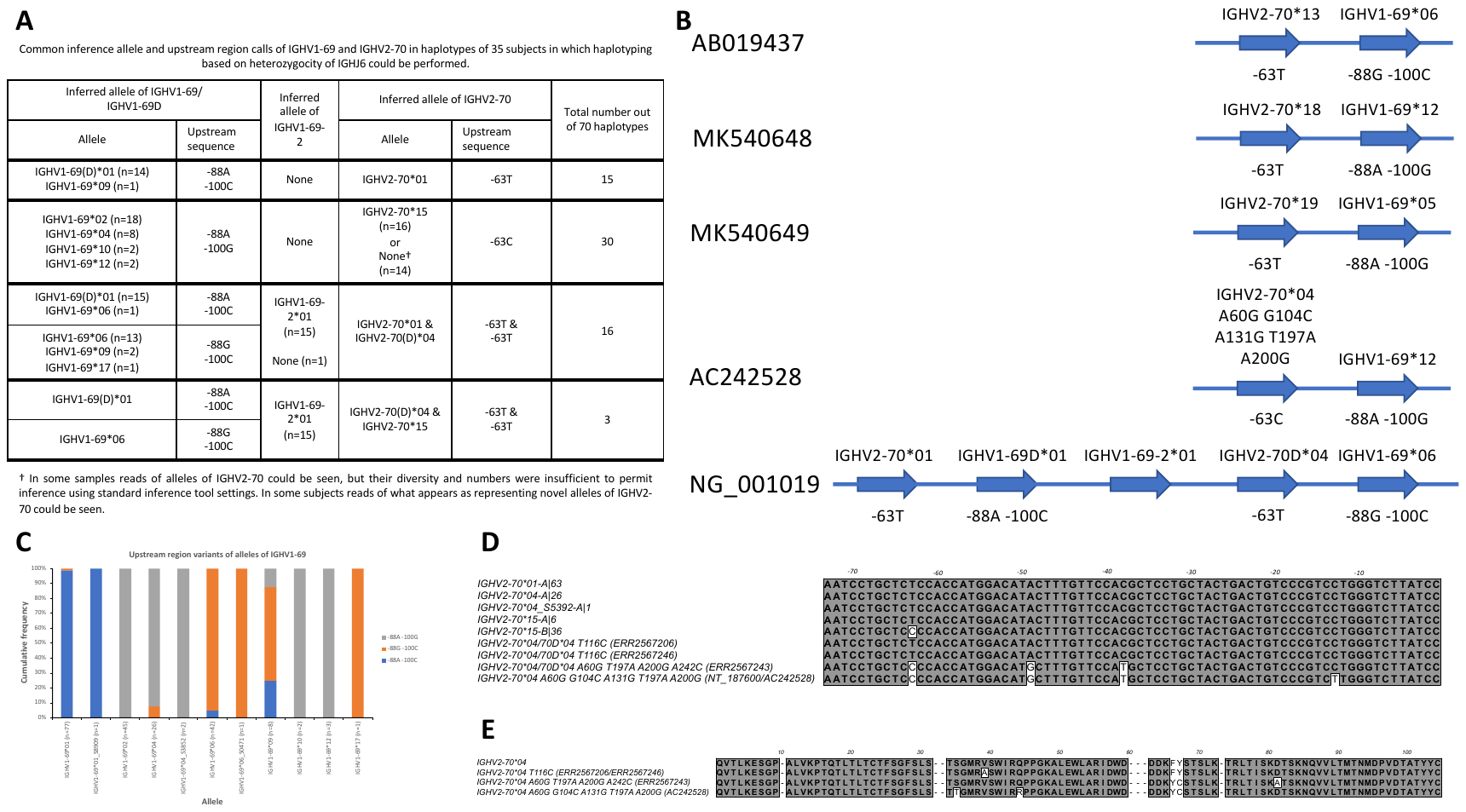

**Supplementary Figure 7.** Common variants of the IGHV1-69 and IGHV2-70 region of human haplotypes, including duplicated variants of this region, as represented by alleles of the genes and variants of their upstream regions identified by inference technology (**A**). Examples of germline this part of the IGHV locus as defined by genomic sequencing (**B**). Different alleles of the IGHV1-69/IGHV1-69D genes are primarily associated to different upstream region sequence variants (**C**). Commonly inferred alleles of IGHV2-70 are commonly associated to the same upstream region, although some instances of IGHV2-70*15 are associated to a different upstream region (**D**), in which case it also occurs in a different genomic context (**A**). In addition, in the transcriptome of some individuals, rare reads of additional variants of IGHV2-70 were identified (**E**), alleles that had not been inferred, one of which also encoded unusual residues including a cysteine in framework 3. One such allele has also been identified by genomic sequencing (**B,** **E**). Some of these alleles were associated to upstream regions different from those of commonly expressed alleles of IGHV2-70 (**D**). IGHV2-70*04/70D*04 T116C represents a 5’ and 3’-extended version of IGHV2-70*05, an allele that is currently only partly defined in the IMGT database (GenBank accession number: Z27502).

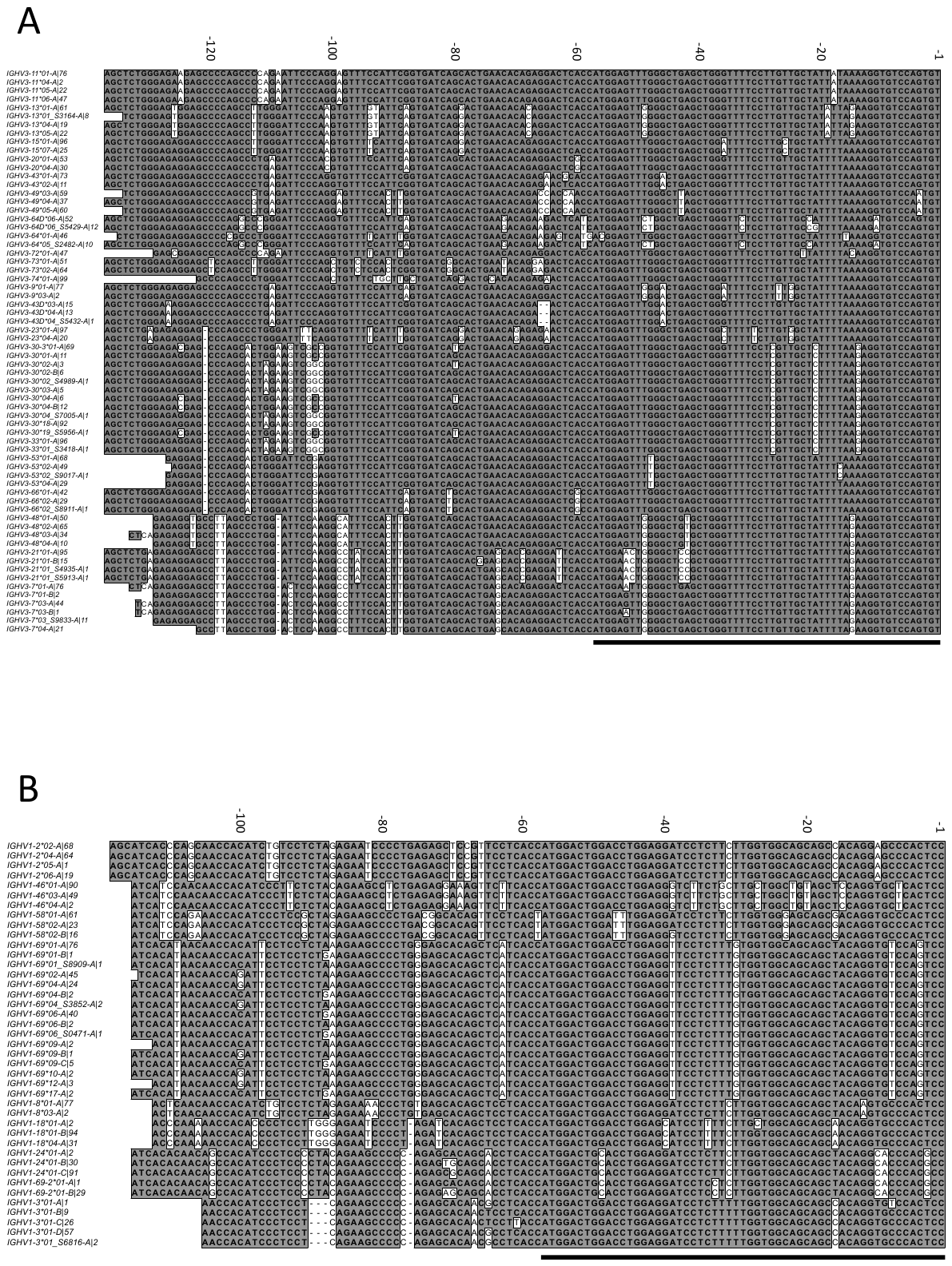

**Supplementary Figure 8.** Inferred upstream regions of genes/alleles of IGHV3 (**A**) and IGHV1 (**B**), grouped according to the presence of possible indels in the 5’UTR. The signal-peptide-encoding leader sequences are indicated by a solid line.

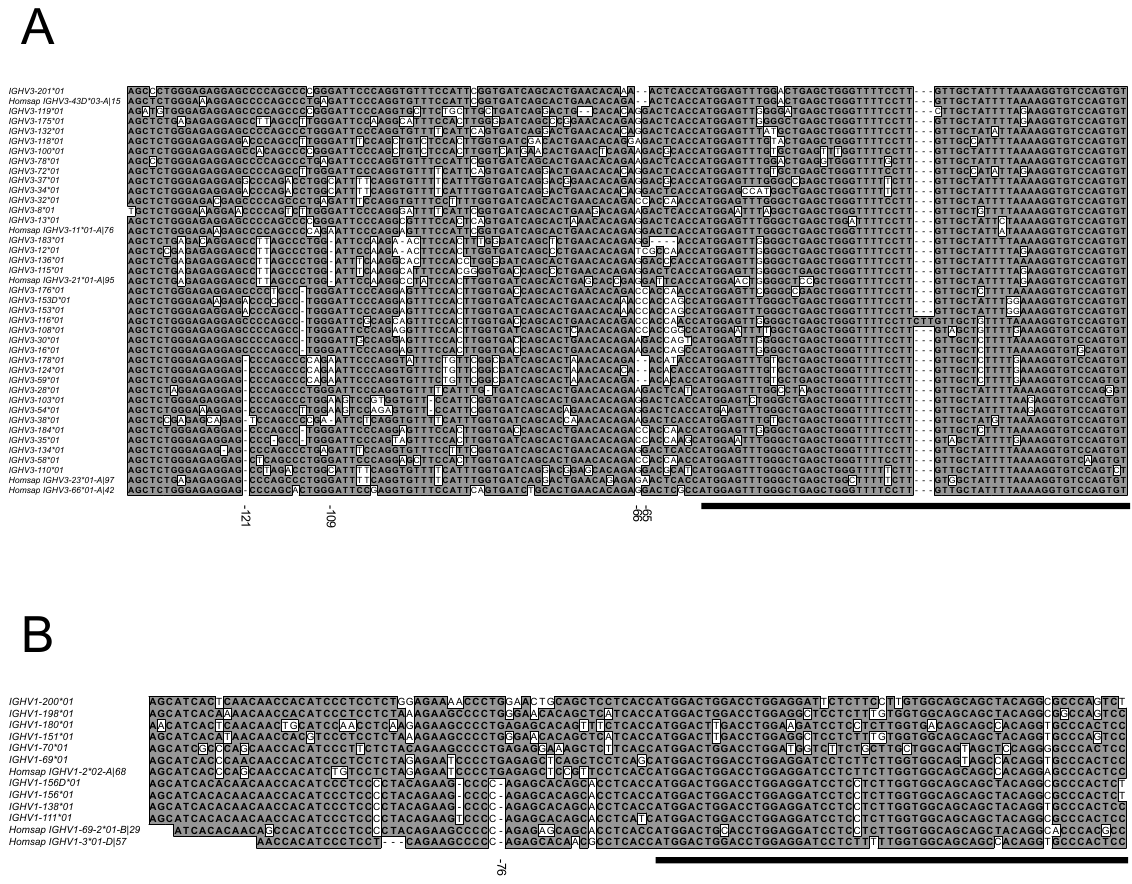

**Supplementary Figure 9.** Sequences of upstream regions (5’UTR-leader sequences excluding the intron found in the leader sequence) of functional genes of Rhesus monkey subgroups IGHV3 (**A**) and IGHV1 (**B**) as defined by IMGT entry IMGT000064 (<http://www.imgt.org/ligmdb/view?id=IMGT000064>). Examples of the corresponding inferred upstream regions of human alleles (all with the prefix “Homsap”) representing sequences of different lengths or showing evidence of insertion/deletion events are included. The sequences encoding the gene products’ signal sequences are underlined.

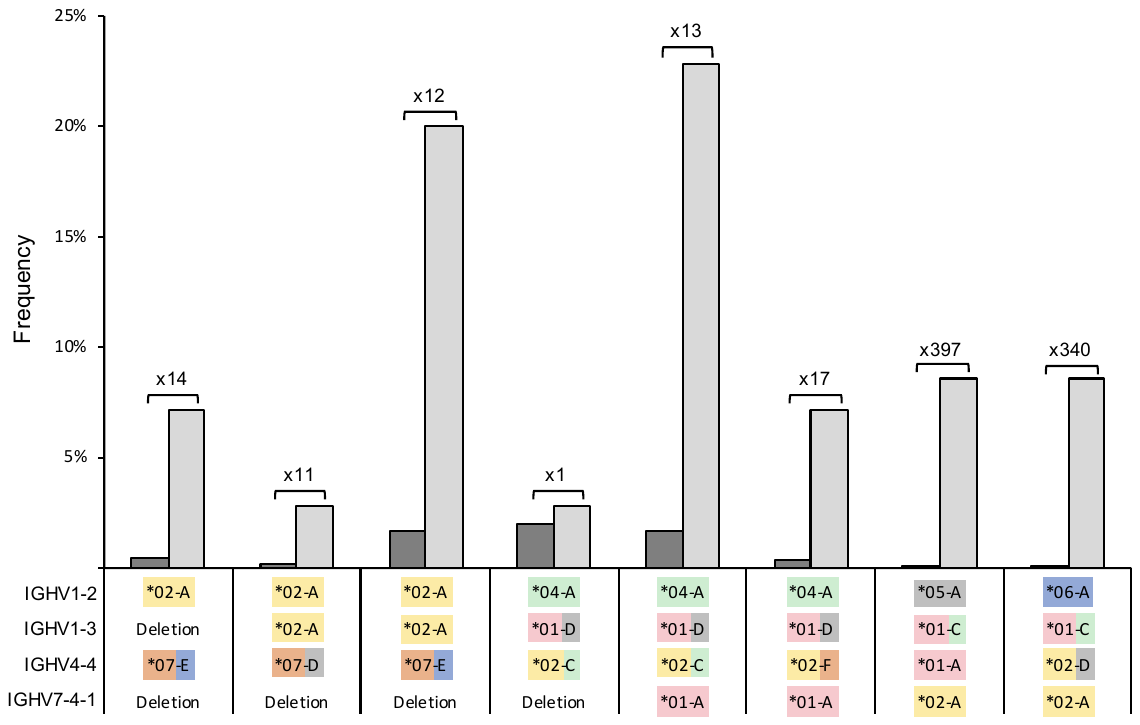

**Supplementary Figure 10.** Frequency of different gene-allele-upstream region combinations (light grey) in the 35 haplotypable individuals in the studied data set, together with theoretical frequencies (dark grey) (assuming random association of alleles). Values above bars indicate the ratio between the theoretical and actual frequency in the given 70 haplotypes. Gene combinations present in <2 haplotypes are not shown.
